# Nuclear phylogenomics clarifies the family-level backbone and gene-tree conflict in Zingiberales

**DOI:** 10.64898/2026.06.25.734679

**Authors:** Jiaxin Wang, Qihong Zhu, Canmei Chen, Yixin Luo, Jian He

## Abstract

Zingiberales includes eight morphologically distinctive families, but its family-level backbone has remained unstable, especially around Musaceae, Heliconiaceae, Lowiaceae, and Strelitziaceae. We analysed 1566 low-copy nuclear genes from 52 samples, representing all eight families and *Pontederia crassipes* as outgroup. Concatenated maximum likelihood and multispecies coalescent analyses recovered the same backbone: ((Zingiberaceae, Costaceae), (Cannaceae, Marantaceae)) is sister to (Musaceae, (Heliconiaceae, (Lowiaceae, Strelitziaceae))). Penalized-likelihood dating placed the sampled crown group in the Late Cretaceous, with several deep family-level divergences occurring on short internodes. Analysis of 1248 rerooted gene trees showed that conflict is concentrated on these deep branches and in several shallow clades. HyDe tests of empirical and simulated matrices, each including 62,475 triples, did not support widespread ancient hybridization among the major family-level lineages after filtering against the simulated null model. The nuclear data recover a stable Zingiberales backbone, and the long-standing instability of several deep nodes is best explained by rapid early divergence and extensive incomplete lineage sorting.

## Introduction

Zingiberales is a morphologically distinctive order of commelinid monocots comprising eight families: Musaceae, Heliconiaceae, Strelitziaceae, Lowiaceae, Cannaceae, Marantaceae, Costaceae, and Zingiberaceae (APG IV, 2016). These families are readily recognized by combinations of habit, leaf sheath structure, inflorescence architecture, floral symmetry, number of fertile stamens, and development of petaloid staminodes (Kress, 1990; Kress and Specht, 2006). Family limits in the order are therefore relatively clear, but the family-level backbone has been harder to resolve. The most persistent uncertainty concerns the early branching order of Musaceae, Heliconiaceae, Lowiaceae, and Strelitziaceae, and the relationship of this assemblage to the more highly modified ginger clade.

Molecular studies over the past two decades have progressively narrowed the problem. Analyses combining morphology with plastid rbcL and atpB and nuclear 18S rDNA, complete plastid genomes, multiplexed exon capture, and low-copy nuclear genes have repeatedly recovered several comparatively stable sister-group relationships, including Zingiberaceae + Costaceae, Cannaceae + Marantaceae, and Lowiaceae + Strelitziaceae (Kress et al., 2001; Barrett et al., 2014a; Barrett et al., 2014b; Sass et al., 2016; Givnish et al., 2018; Carlsen et al., 2018). By contrast, the placement of Musaceae, the sister-group relationship of Heliconiaceae, and the relationship between the banana-bird-of-paradise clade and the ginger clade have varied among data sets and analytical strategies. This pattern suggests that the remaining uncertainty is concentrated on a small number of deep, short internodes rather than reflecting inadequate family sampling or unstable family circumscriptions.

Topological conflict on such short internodes can arise from different evolutionary processes. Rapid successive divergences can carry ancestral polymorphism across speciation events, producing incomplete lineage sorting and allowing different gene trees to support different relationships (Pamilo and Nei, 1988; Maddison, 1997; Degnan and Rosenberg, 2009). Ancient hybridization or introgression can also generate gene-tree conflict, but should leave a more directional signal of excess allele sharing between particular lineages (Solis-Lemus and Ane, 2016; Blischak et al., 2018). Resolving the Zingiberales backbone therefore requires a species tree, explicit measurement of gene-tree conflict, and tests for possible reticulate signals.

Here we assembled a nuclear phylogenomic data set representing all eight families of Zingiberales. Using 1566 low-copy nuclear genes, we inferred concatenated and multispecies coalescent trees, estimated divergence times for major nodes, quantified conflict between gene trees and the species tree, and used HyDe to test candidate hybridization signals in both empirical and simulated matrices. We address three questions: what is the family-level nuclear backbone of Zingiberales; where are the nodes that have caused repeated instability in previous studies; and are these conflicts more consistent with incomplete lineage sorting, ancient hybridization, or a combination of both processes?

## Material and methods

### Sampling and sequence resources

The data set included 52 samples, comprising 51 ingroup samples from Zingiberales and *Pontederia crassipes* as outgroup. All eight families of Zingiberales were represented: Musaceae (17 samples), Zingiberaceae (13), Cannaceae (5), Marantaceae (5), Strelitziaceae (3), Lowiaceae (3), Costaceae (3), and Heliconiaceae (2). The data set combined 30 genome-derived samples and 22 transcriptome-derived samples. Sample names, data type, original source, accession information, and CDS recovery statistics are provided in Table S1 and Table S2.

Sampling was designed to resolve the family-level backbone and to include several lineages in which gene-tree conflict was expected. Musaceae sampling included *Ensete, Musella*, and *Musa*, with independent genomic or transcriptomic accessions retained for several species. Zingiberaceae sampling covered *Alpinia, Wurfbainia, Etlingera, Curcuma, Hedychium, Boesenbergia, Zingiber*, and *Kaempferia*. The remaining families were represented by the best available public resources, including *Orchidantha* in Lowiaceae, *Strelitzia* in Strelitziaceae, *Canna* in Cannaceae, *Maranta, Thalia*, and *Ischnosiphon* in Marantaceae, *Costus* in Costaceae, and *Heliconia* in Heliconiaceae.

### Orthology inference and low-copy gene filtering

For genome-derived samples, annotated CDS and protein sequences were used directly. For transcriptome-derived samples, CDS and protein sequences inferred with TransDecoder were used. Sequence headers were standardized before analysis to remove spaces, path separators, and accession-format inconsistencies that could interfere with downstream parsing.

Orthologous groups were inferred with Proteinortho (Lechner et al., 2011). Proteinortho output was filtered with custom Python scripts. We retained clusters that met three criteria: the proportion of missing species was no greater than 0.5; the Proteinortho Alg.-Conn. value was at least 0.5; and each sample contributed no more than two candidate low-copy sequences to a cluster. If two candidate sequences were present for a sample, the longer CDS was retained. Clusters containing more than two candidate sequences for any sample were removed to reduce the risk of including paralogous sequences. This procedure retained 1566 low-copy nuclear genes.

### Alignments, filtering, and concatenation

Each CDS gene was aligned separately with MAFFT v7 (Katoh and Standley, 2013). Columns with more than 20% missing data or indels were removed. The filtering script also removed sequences consisting entirely of gaps, question marks, or ambiguous bases, and padded shorter sequences where necessary to maintain equal alignment lengths.

Filtered single-gene alignments were concatenated with a custom script. Missing genes for a sample were filled with question marks, allowing all retained genes to be included in the supermatrix. The final concatenated matrix included 52 samples and 1,868,754 nucleotide sites. RAxML identified 1,050,539 distinct site patterns, and 21.43% of cells in the matrix were gaps or completely undetermined characters.

### Phylogenetic analyses

The concatenated maximum likelihood tree was inferred with RAxML v8.2.12 (Stamatakis, 2014) under the GTRGAMMA model. The analysis used the rapid bootstrap and best-tree search option with the command raxmlHPC-PTHREADS -T 150 -n result -s result.fasta -m GTRGAMMA -p 12345 -f a -N 11 -x 12345. Bootstrap values were mapped onto the best maximum likelihood tree. In Figure 1, only bootstrap values below 100 are shown; unlabelled internal nodes received full support in the plotted tree.

**Figure 1.**
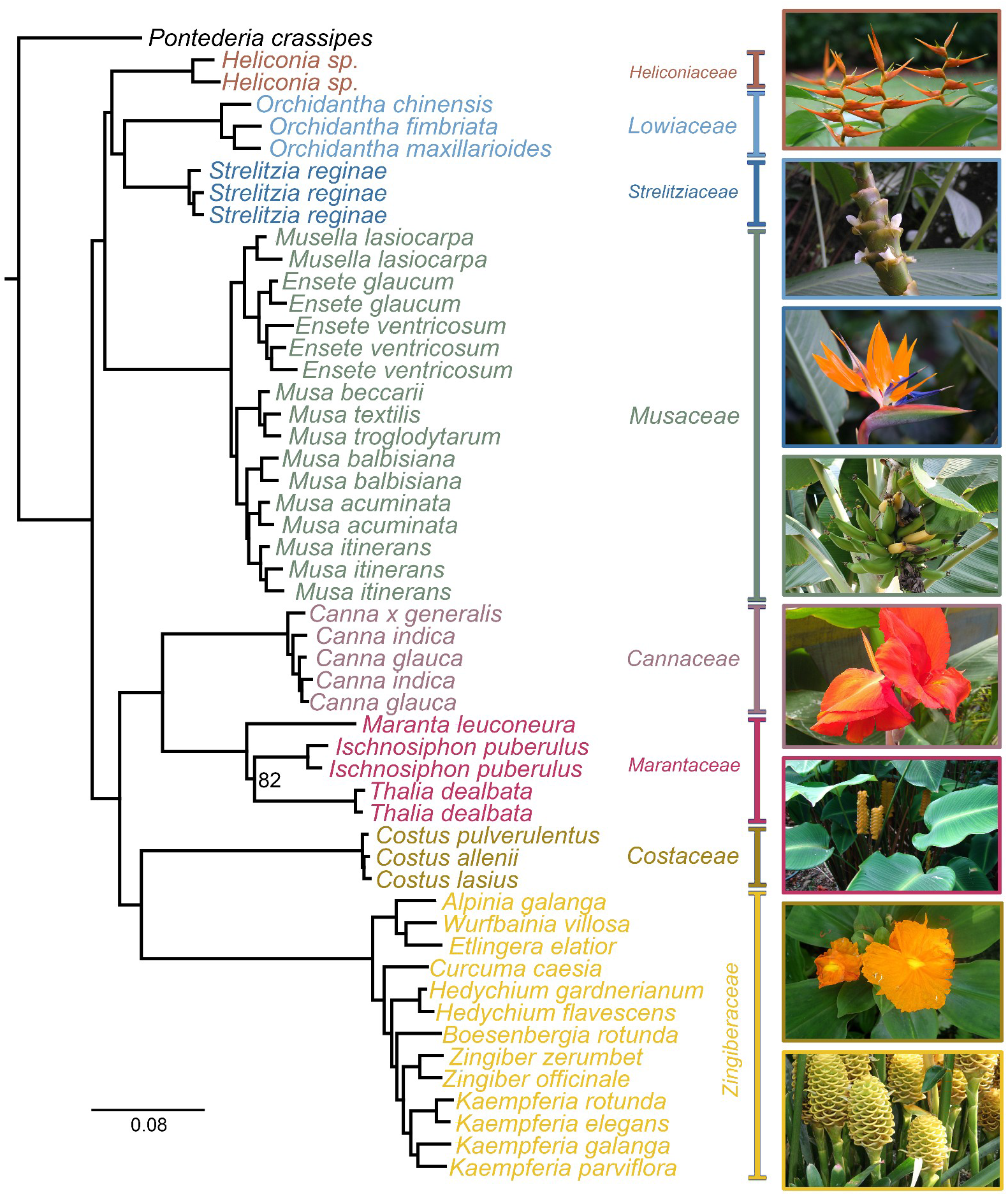
Maximum likelihood phylogeny of Zingiberales inferred from the concatenated nuclear matrix. The tree includes 51 ingroup samples of Zingiberales and *Pontederia crassipes* as the outgroup. Branch lengths are proportional to substitutions per site. Numbers at nodes are bootstrap support values below 100; unlabeled internal nodes have full bootstrap support. Coloured bars indicate family assignment.

Single-gene maximum likelihood trees were also inferred with RAxML under the GTRGAMMA model and were used for multispecies coalescent analyses and gene-tree conflict analyses. Species trees were inferred with ASTRAL-MP within the ASTRAL summary-coalescent framework (Mirarab et al., 2014; Zhang et al., 2018; Yin et al., 2019). In addition to the full set of 1566 gene trees, eight filtered data sets were generated to test sensitivity to gene selection: mean bootstrap support >= 60 (1560 genes), mean bootstrap support >= 70 (1497 genes), missing data <= 10% (239 genes), missing data <= 20% (830 genes), alignment length >= 500 bp (1394 genes), alignment length >= 1000 bp (861 genes), parsimony-informative sites >= 200 (1482 genes), and parsimony-informative sites >= 400 (1040 genes).

### Divergence time estimation

Divergence times were estimated using the coalescent species-tree topology as the backbone. Branch lengths were first estimated on the fixed topology with IQ-TREE (Nguyen et al., 2015; Minh et al., 2020) under the GTR+F+G4 model using the concatenated matrix. Penalized-likelihood dating was then performed with treePL (Smith and O’Meara, 2012). Calibration constraints were set as follows: the root node at 99.0-99.1 Ma, a Musaceae-related split at 43.0-43.1 Ma, and the Zingiberaceae crown node at 65.0-65.1 Ma, based on fossil and metacalibrated monocot age evidence relevant to Zingiberales and its close relatives (Manchester and Kress, 1993; Magallon et al., 2015). Dating uncertainty was assessed using 100 bootstrap-resampled alignments generated by resampling sites from the concatenated matrix. For each bootstrap replicate, branch lengths were re-estimated on the same fixed topology and the resulting tree was dated with treePL using thorough optimization; the smoothing parameter was selected by cross-validation for each run. The dated bootstrap trees were summarized as a maximum-clade-credibility chronogram, which was used as the dated species tree in Figure 2.

**Figure 2.**
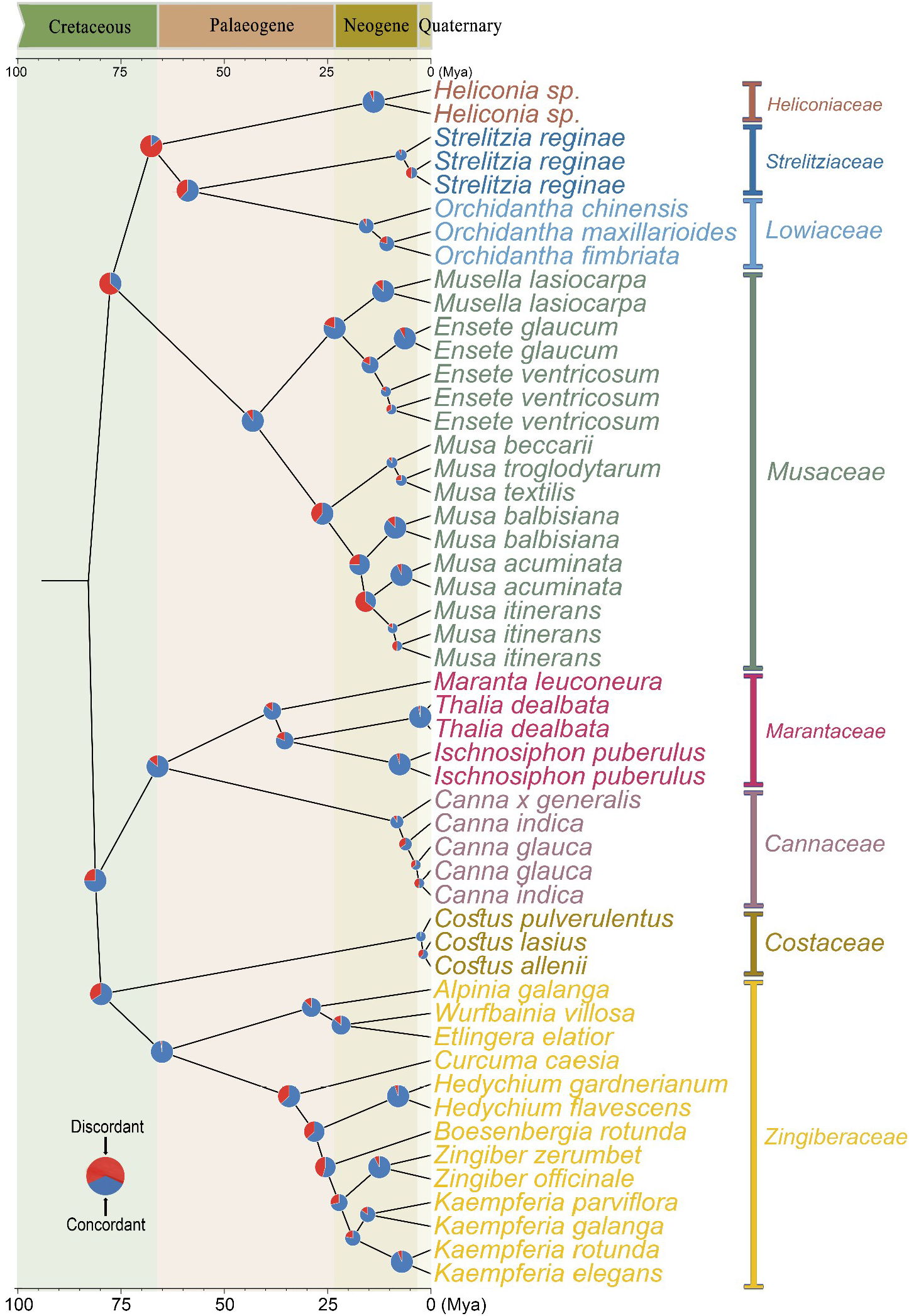
Dated coalescent species tree of Zingiberales with gene-tree conflict mapped to internal nodes. Branch lengths are scaled to time before present (Ma). Node pie charts summarize 1248 rerooted gene trees: blue indicates gene trees concordant with the species-tree bipartition, and red indicates discordant gene trees. Coloured bars indicate family-level lineages; geological periods are shown above and behind the tree.

### Gene-tree conflict analysis

Conflict between gene trees and the dated species tree was quantified by comparing bipartitions. Gene trees containing the outgroup *Pontederia crassipes* were first rerooted with that taxon. After rerooting and filtering, 1248 gene trees were retained for node-level conflict analysis. For each internal node of the species tree, gene trees were classified as concordant when they supported the species-tree bipartition, discordant when they supported an alternative bipartition, or uninformative when taxon sampling or resolution was insufficient. Counts of concordant and discordant gene trees were plotted as node pie charts on the dated tree (Fig. 2).

### HyDe analyses and false-positive filtering

Potential hybridization or introgression was tested with HyDe (Blischak et al., 2018) using the nuclear matrix. A total of 62,475 triples were tested in the empirical matrix. For each triple, HyDe reported candidate parents, the candidate hybrid lineage, Z score, P value, and gamma value. To associate HyDe results with species-tree nodes, we generated VisualHyDe-style heat maps for 50 internal nodes.

Because incomplete lineage sorting, topology, uneven information content, and missing-data structure can produce nominally significant HyDe results, we generated a simulated null matrix with the same topology and matrix structure and analysed it using the same pipeline. Biological interpretation was based primarily on the false-positive-filtered results, with the unfiltered empirical and simulated matrices used as controls for background signal.

## Results

### Nuclear gene data set

The final data set included 52 samples representing all eight families of Zingiberales and one outgroup. CDS recovery varied among samples, with the number of non-redundant CDS ranging from 9680 to 44,760 and an average of 22,007.2. After Proteinortho clustering and low-copy filtering, 1566 nuclear genes were retained for phylogenetic analyses. The concatenated matrix contained 1,868,754 nucleotide sites and sufficient information for family-level phylogenetic inference, divergence dating, and gene-tree conflict analysis.

### Concatenated nuclear genes recover a stable family-level backbone

The RAxML tree inferred from the concatenated matrix recovered a clear family-level structure in Zingiberales (Fig. 1). All eight families formed the expected family-level or sampled family-level lineages. Zingiberaceae and Costaceae were sister groups and together were sister to Cannaceae + Marantaceae. In the other major lineage, Musaceae was sister to a clade comprising Heliconiaceae, Lowiaceae, and Strelitziaceae; Heliconiaceae was sister to Lowiaceae + Strelitziaceae. Apart from a small number of shallow nodes, the family-level backbone and major intrafamilial relationships received high support.

### Coalescent analyses and gene filtering support the same backbone

The ASTRAL-MP species tree inferred from all 1566 gene trees was congruent with the concatenated maximum likelihood tree at the family-level backbone. The eight filtered coalescent data sets also supported the same main relationship: ((Zingiberaceae, Costaceae), (Cannaceae, Marantaceae)) is sister to (Musaceae, (Heliconiaceae, (Lowiaceae, Strelitziaceae))). Differences among filtered data sets were limited mainly to local arrangements or support values on shallow or short branches, rather than to the overall family-level backbone. The nine coalescent species trees and their node supports are shown in Figure S1.

### Divergence time estimates

The treePL analysis placed the sampled crown group of Zingiberales in the Late Cretaceous, at approximately 82.8 Ma (Fig. 2). The split between the banana-bird-of-paradise clade and the ginger-canna-arrowroot-costus clade occurred within a narrow Late Cretaceous time window. Under the fossil constraints used here, the crown age of Zingiberaceae was estimated at 65.0 Ma and the crown age of Musaceae at 43.0 Ma. Other dated nodes included the crowns of Cannaceae at approximately 8.1 Ma and Marantaceae at approximately 38.3 Ma. These time estimates indicate that several key family-level divergences occurred along short internodes, providing a temporal context for the gene-tree conflict observed below.

#### Gene-tree conflict is concentrated on deep short branches and some shallow clades

Gene-tree conflict analysis retained 1248 rerooted gene trees. The numbers of concordant, discordant, and uninformative gene trees varied markedly among nodes (Fig. 2). Several family crown nodes, including those of Zingiberaceae, Costaceae, Cannaceae, Marantaceae, and Musaceae, were supported by many gene trees. By contrast, deeper backbone nodes showed higher proportions of discordant and uninformative gene trees.

Conflict was particularly pronounced around the banana-bird-of-paradise clade. Deep nodes connecting Heliconiaceae, Lowiaceae, Strelitziaceae, and Musaceae had many discordant and uninformative gene trees, indicating that this part of the nuclear genome does not record lineage divergence as a single uniform history. Local conflict was also detected within *Musa* and around several shallow nodes in Zingiberaceae, including near *Kaempferia, Zingiber, Boesenbergia, Hedychium*, and *Curcuma*. These results localize the sources of phylogenetic instability to a small number of deep short branches and to some rapidly diverging intrafamilial lineages.

### HyDe results and false-positive filtering

HyDe tested 62,475 triples in both the empirical matrix and the simulated null matrix. The unfiltered empirical heat maps contained 18,806 non-empty cells across 36 nodes. The simulated matrix also produced many nominal signals, indicating substantial background false positives in the raw HyDe output. After filtering against the simulated matrix, 2220 non-empty cells were retained across 27 nodes in the empirical results (Figs S2-S4).

Filtering changed the interpretation of HyDe results substantially. The unfiltered heat maps should not be read as a list of hybridization events. Only signals present in the empirical matrix and absent from comparable patterns in the simulated null model are appropriate as candidate introgression signals. Overall, conflict on the deep family-level backbone did not show a strong, widespread signal of interfamily hybridization sufficient to replace an incomplete-lineage-sorting explanation. Some shallow nodes and local lineages retained directional HyDe signals and represent better targets for denser future sampling within genera or species complexes.

## Discussion

### A resolved family-level framework for Zingiberales

The concatenated maximum likelihood and multispecies coalescent analyses recovered the same family-level backbone. Musaceae is the earliest-diverging lineage within the banana-bird-of-paradise clade, Heliconiaceae is placed next, and Lowiaceae and Strelitziaceae are sister groups. This clade as a whole is sister to the ginger clade, here recovered as ((Zingiberaceae, Costaceae), (Cannaceae, Marantaceae)). The same framework was also supported by coalescent trees inferred from eight filtered gene sets. This result is consistent with the main signal recovered in some low-copy nuclear and broad monocot plastome studies (Givnish et al., 2018; Carlsen et al., 2018), and it further narrows the historically unstable portion of the backbone to consecutive short internodes near Musaceae and Heliconiaceae.

This topology separates stable family-level relationships from the nodes that have driven earlier instability. Zingiberaceae + Costaceae, Cannaceae + Marantaceae, and Lowiaceae + Strelitziaceae can be treated as repeated and comparatively stable sister-group relationships across morphology-molecular, plastome, exon-capture, and nuclear data sets (Kress et al., 2001; Barrett et al., 2014a; Barrett et al., 2014b; Sass et al., 2016; Carlsen et al., 2018). The placement of Musaceae and the sister-group relationship of Heliconiaceae correspond to the most difficult deep nodes. The resolved backbone provides a firmer basis for comparing characters between the banana-bird-of-paradise clade and the ginger clade.

The topology also helps polarize floral character evolution in the order. Musaceae, Heliconiaceae, Lowiaceae, and Strelitziaceae generally retain more fertile stamens and a floral organization closer to early-diverging states in Zingiberales. Zingiberaceae, Costaceae, Cannaceae, and Marantaceae show more extensive stamen reduction, petaloid staminode development, floral organ fusion, and changes in floral symmetry (Kress, 1990; Kress and Specht, 2006). Within this framework, the highly modified flowers of the ginger clade can be interpreted as a set of linked character transitions following establishment of that lineage. The sister-group relationship of Zingiberaceae and Costaceae provides a basis for comparing the role of staminodes in floral display, whereas Cannaceae + Marantaceae provides a direct framework for analysing asymmetry of the androecium and the evolution of specialized pollination mechanisms (Bartlett and Specht, 2011; Yu et al., 2020).

The resolved relationships within the banana-bird-of-paradise clade also clarify comparisons of pollination-related and floral display traits. The basal position of Musaceae indicates that the elaborate floral displays of Heliconiaceae, Lowiaceae, and Strelitziaceae should be interpreted on later-diverging branches. Heliconiaceae has conspicuous bracts, bird pollination, and diverse inflorescence orientations, traits closely associated with pollinator interactions (Iles et al., 2017). Lowiaceae and Strelitziaceae are sister groups but differ markedly in floral display and ecological strategy (Cron et al., 2012). The family-level backbone makes the evolutionary history of floral structure, pollination systems, and ecological display strategies in early Zingiberales more directly testable.

### Gene-tree conflict explains long-standing backbone instability

The instability of the Zingiberales backbone is closely associated with topological conflict among nuclear gene trees. When deep short internodes and gene-tree conflict occur together, different marker sets can sample different historical signals, as expected under the multispecies coalescent and as widely observed in phylogenomic analyses of ancient radiations (Edwards, 2009; Degnan and Rosenberg, 2009; Smith et al., 2015; Salichos and Rokas, 2013). A single locus, or a completely linked plastid genome, represents only one part of the genomic history and may support a topology different from the main species-tree signal recovered from the nuclear genome.

The node-level counts match this interpretation. Several family crown nodes are supported by many gene trees, showing that the data contain strong phylogenetic signal. Deep nodes connecting major family-level lineages, however, have higher proportions of discordant and uninformative gene trees. This pattern is consistent with coalescent expectations for short internal branches, where alternative gene-tree topologies can remain common and, in some parameter regions, may even be more probable than the gene tree matching the species tree (Rosenberg, 2002; Degnan and Rosenberg, 2006, 2009). In the banana-bird-of-paradise clade, fewer gene trees support the species-tree bipartition than conflict with it, and many gene trees cannot resolve the node. This pattern explains why relationships among Musaceae, Heliconiaceae, Lowiaceae, and Strelitziaceae have varied among previous studies: these early nodes preserve multiple gene histories within the nuclear genome.

High support in a concatenated analysis should be interpreted together with evidence from individual gene trees. Concatenation can recover a well-resolved tree from the dominant signal, but it does not show the proportion of genes supporting alternative topologies and can become misleading when gene-tree discordance is high (Kubatko and Degnan, 2007; Ane et al., 2007; Edwards, 2009). For an ancient rapid radiation such as Zingiberales, concatenated trees, coalescent species trees, and gene-tree conflict analyses provide complementary evidence. Gene-tree conflict records the coexistence of multiple gene histories during the early diversification of the order.

#### Rapid early divergence, incomplete lineage sorting, and limited evidence for interfamily hybridization

Divergence time estimation provides a temporal explanation for the observed conflict. The sampled crown group of Zingiberales was placed in the Late Cretaceous, and several major family-level divergences occurred within a relatively narrow time interval. The split between the ginger clade and the banana-bird-of-paradise clade, the early divergences within the latter clade, and the divergences among Zingiberaceae, Costaceae, Cannaceae, and Marantaceae all correspond to short internodes. Such rapid successive divergence is a setting in which incomplete lineage sorting is expected to be common (Pamilo and Nei, 1988; Maddison, 1997; Degnan and Rosenberg, 2009).

Incomplete lineage sorting explains how the species tree can be clear while individual gene trees remain in conflict. When lineages diverge in quick succession, ancestral polymorphisms may fail to sort completely before being carried into descendant lineages. Different genomic regions can therefore have different coalescent histories and support different gene-tree topologies; the same process can also create hemiplasy, in which a character or molecular signal appears synapomorphic on a gene tree but does not match the species tree (Avise and Robinson, 2008; Degnan and Rosenberg, 2009). The high conflict at basal family-level nodes in Zingiberales fits this process and explains why studies based on fewer markers can recover different relationships near short internodes.

We also tested for hybridization because introgression can generate both gene-tree conflict and directional allele sharing. Distinguishing introgression from incomplete lineage sorting is a recurring problem in phylogenomics, and tests for reticulate signal are most informative when interpreted alongside gene-tree conflict and appropriate null expectations (Solis-Lemus and Ane, 2016; Folk et al., 2018; Blischak et al., 2018). After comparison with the simulated matrix and filtering of false-positive patterns, HyDe did not reveal a strong signal of interfamily hybridization among the major lineages of Zingiberales. The deep family-level conflict is best interpreted primarily as a consequence of rapid early divergence and incomplete lineage sorting. Several local signals warrant denser sampling at generic and species levels, but they do not alter the interpretation of the family-level backbone.

The nuclear topology, divergence times, gene-tree conflict, and filtered HyDe results explain the long-standing instability of Zingiberales systematics. The family-level backbone has a stable dominant signal, but early short internodes retain substantial genomic conflict. This combination explains why Musaceae and Heliconiaceae have shifted among previous studies and shows why analyses of ancient rapid radiations should report the species tree alongside the underlying gene-tree conflict and tests of possible reticulate evolution (Edwards, 2009; Smith et al., 2015; Parins-Fukuchi et al., 2021).

## Data availability

Orthogroup matrices, the concatenated matrix, species trees, gene trees, HyDe outputs, and analysis scripts are available from Zenodo at https://doi.org/10.5281/zenodo.20908440.

## Supplementary figure legends

Figure S1. ASTRAL multispecies coalescent trees inferred from all gene trees and from eight filtered gene sets. Node pie charts show ASTRAL node support, with blue indicating support and red indicating 1 - support. For visualization, terminal branches are displayed with a length of 1 when terminal branch lengths were absent or non-informative in the ASTRAL output; internal branches retain the relative lengths from the original ASTRAL trees.

Figure S2. HyDe node heat maps after false-positive filtering against the simulated null matrix. Each page corresponds to an internal node of the species tree. Heat-map colour indicates gamma values for retained significant parent-hybrid-parent triples.

Figure S3. Unfiltered HyDe node heat maps from the empirical nuclear matrix. These heat maps show raw empirical signals before false-positive filtering and should be interpreted together with Figures S2 and S4.

Figure S4. HyDe node heat maps from the simulated null matrix. These heat maps show background HyDe patterns expected under a model without true hybridization and were used to filter the empirical results shown in Figure S2.

## Supporting Information

Table S1. Taxon sampling, data sources, data types, and NCBI accession numbers for Zingiberales and the outgroup sampled in this study.

Table S2. CDS recovery statistics for each sample after redundancy reduction.

