## Supplementary material for "Nuclear phylogenomics clarifies the family-level backbone and gene-tree conflict in Zingiberales": Figure S1

### Coalescent species tree from all gene trees

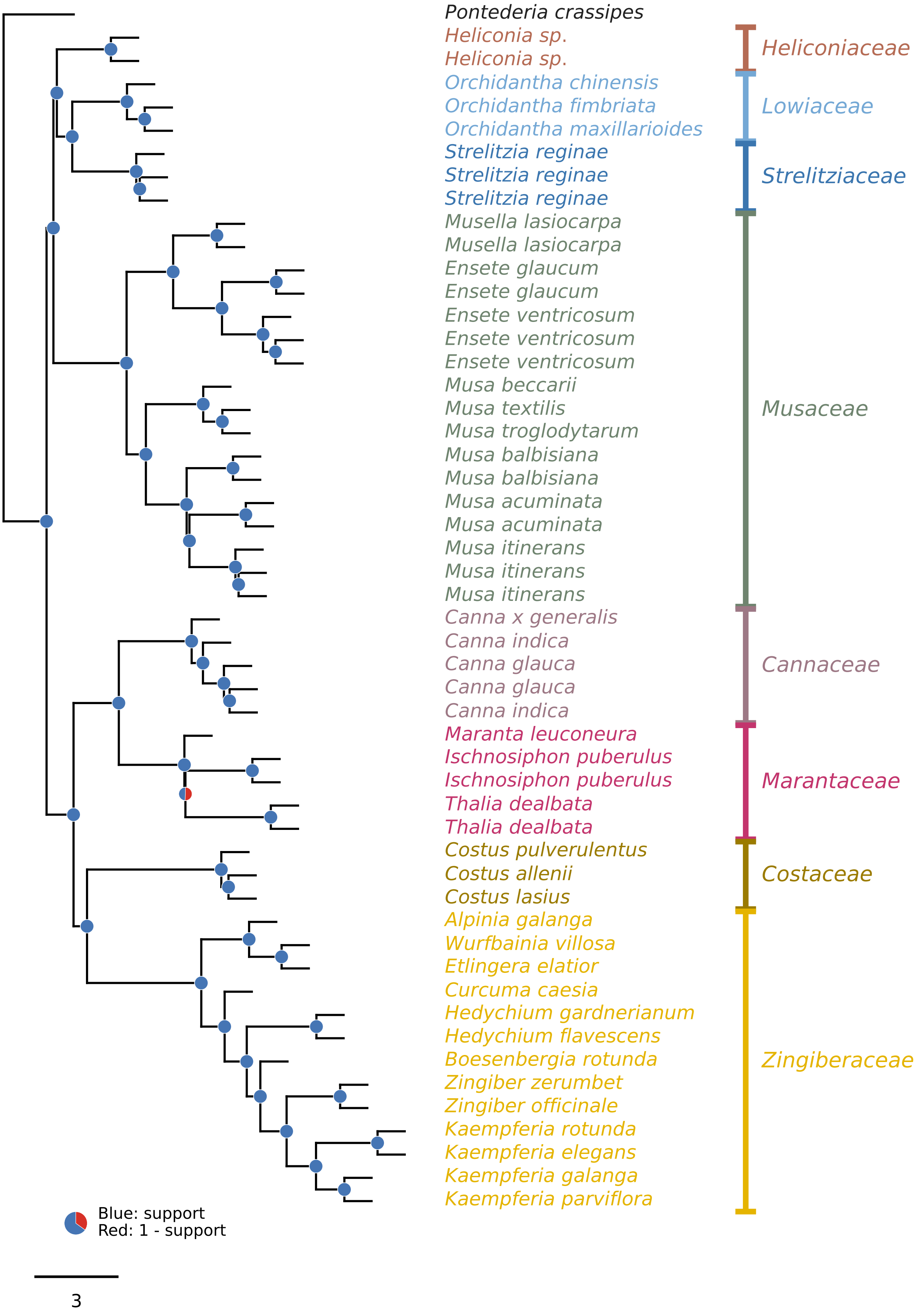

### Coalescent species tree from genes with bootstrap >= 60

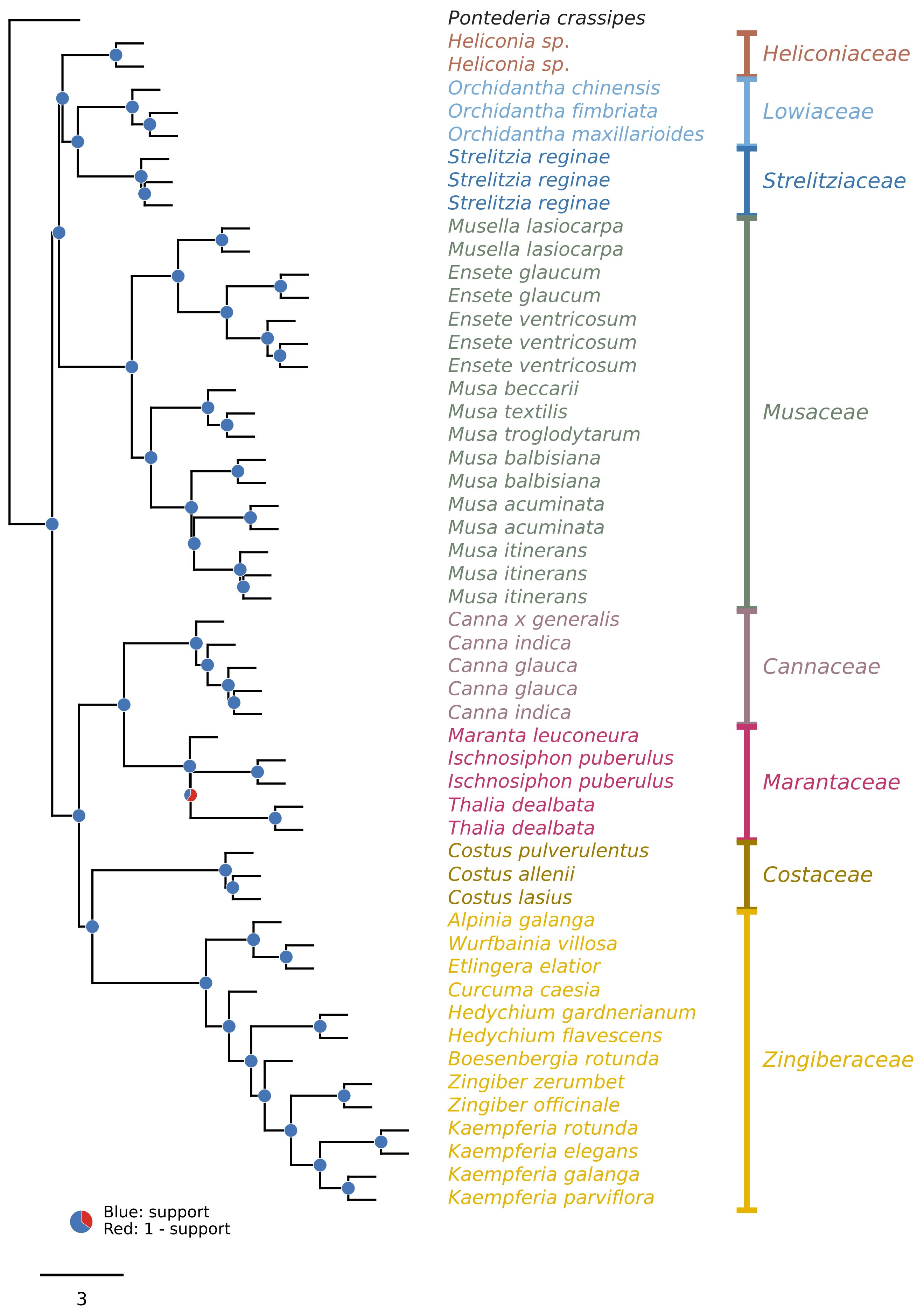

**Coalescent species tree from genes with bootstrap >= 70**

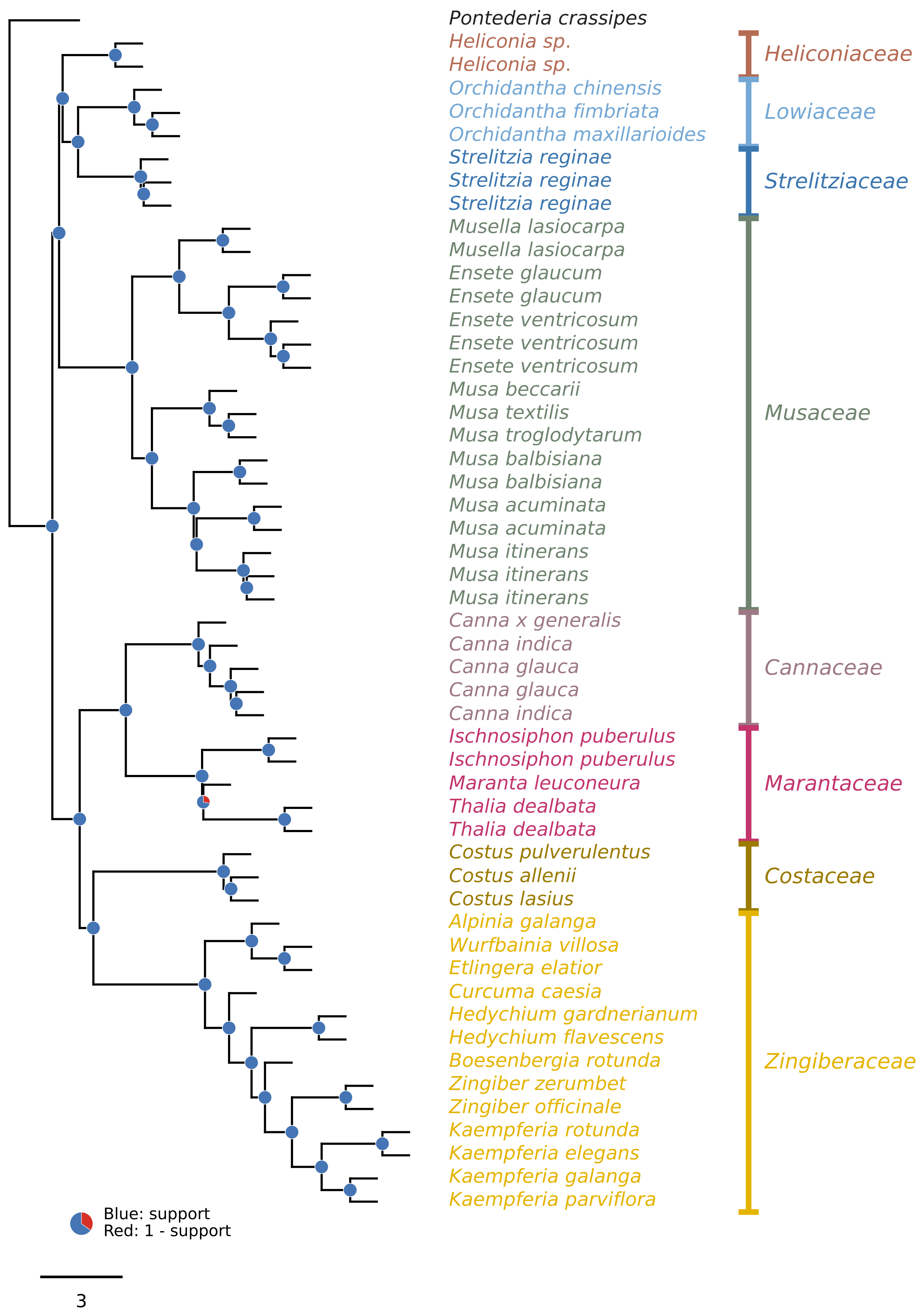

### Coalescent species tree from genes with missing data <= 10%

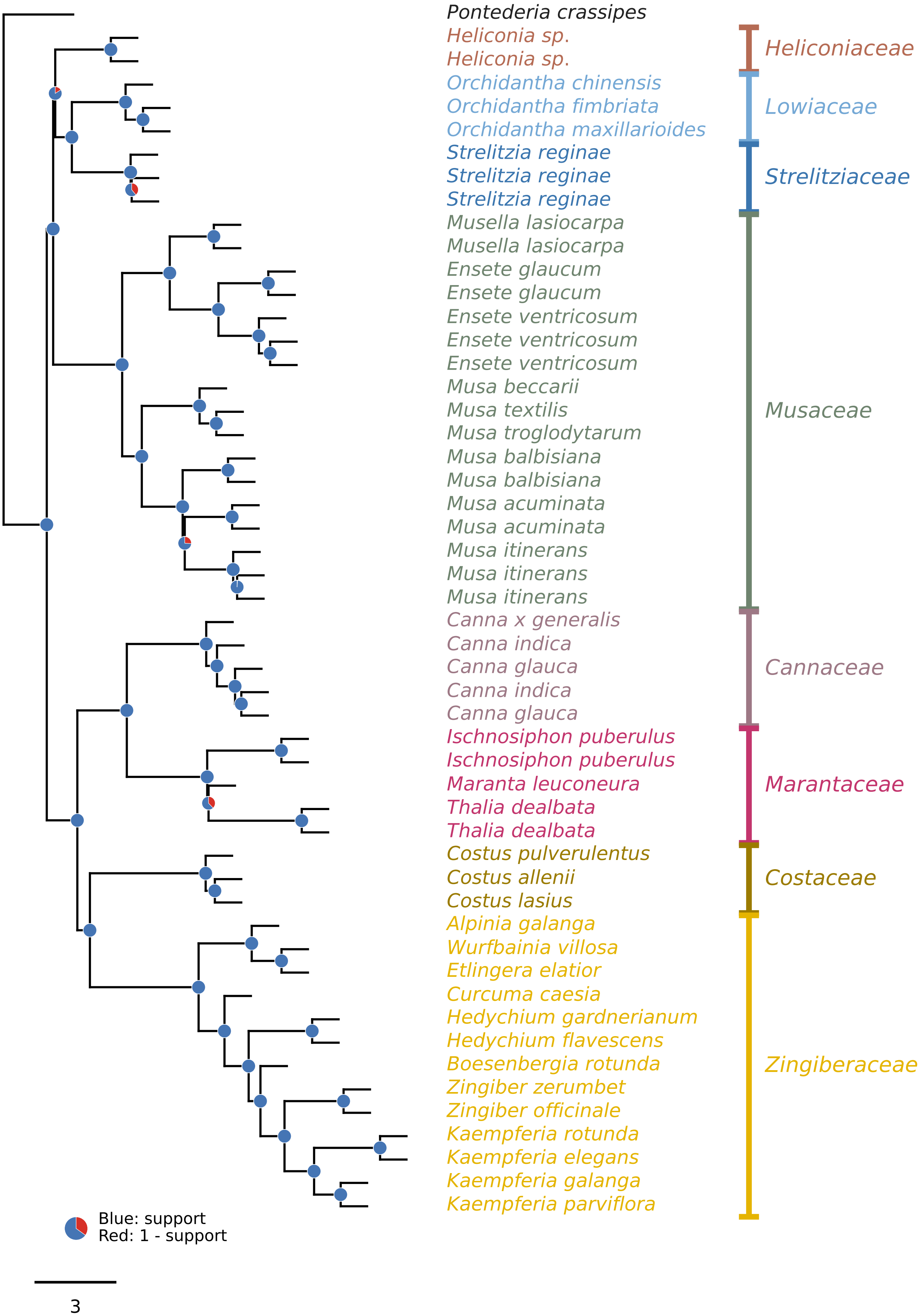

**Coalescent species tree from genes with missing data <= 20%**

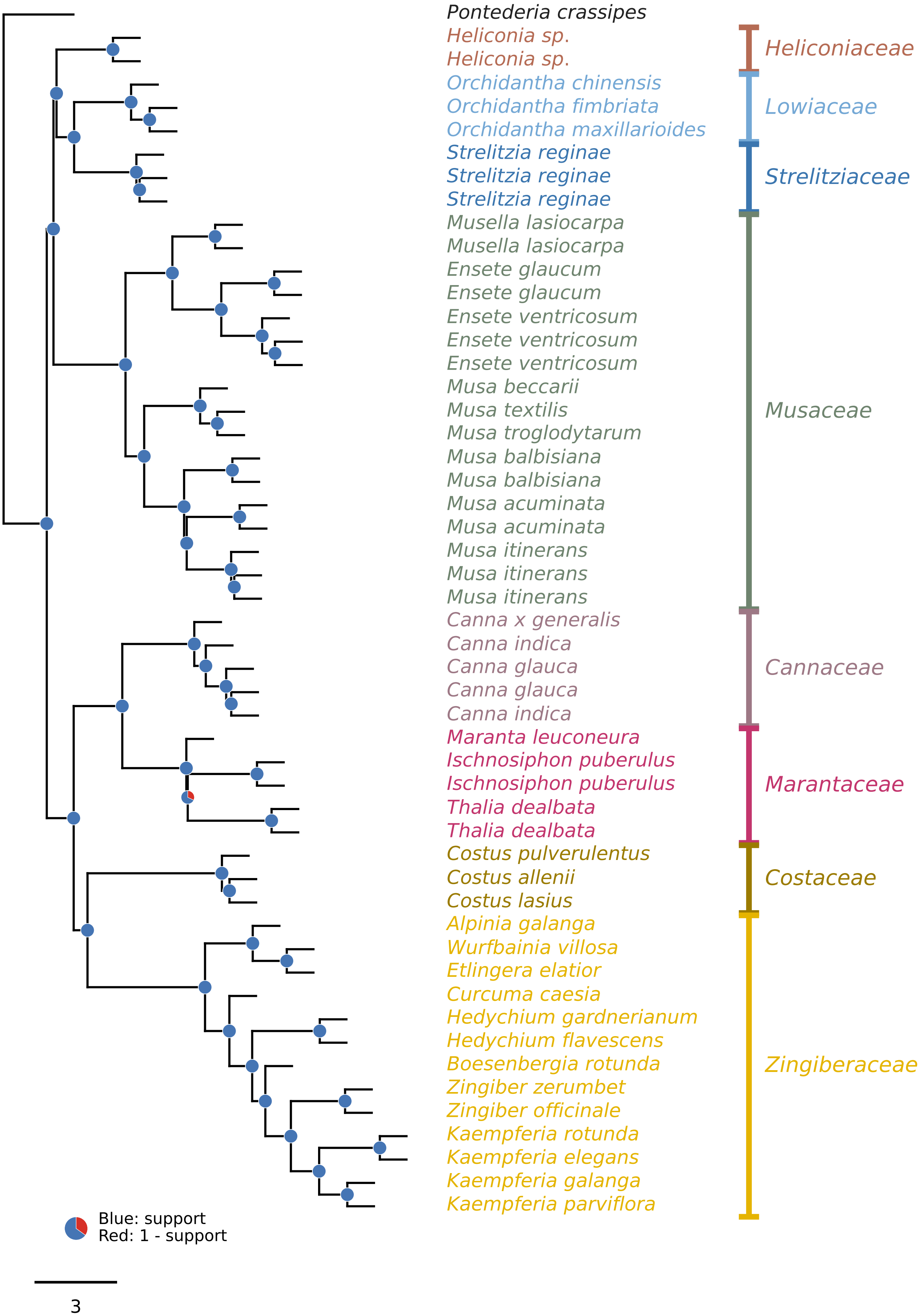

### Coalescent species tree from genes with PI sites >= 200

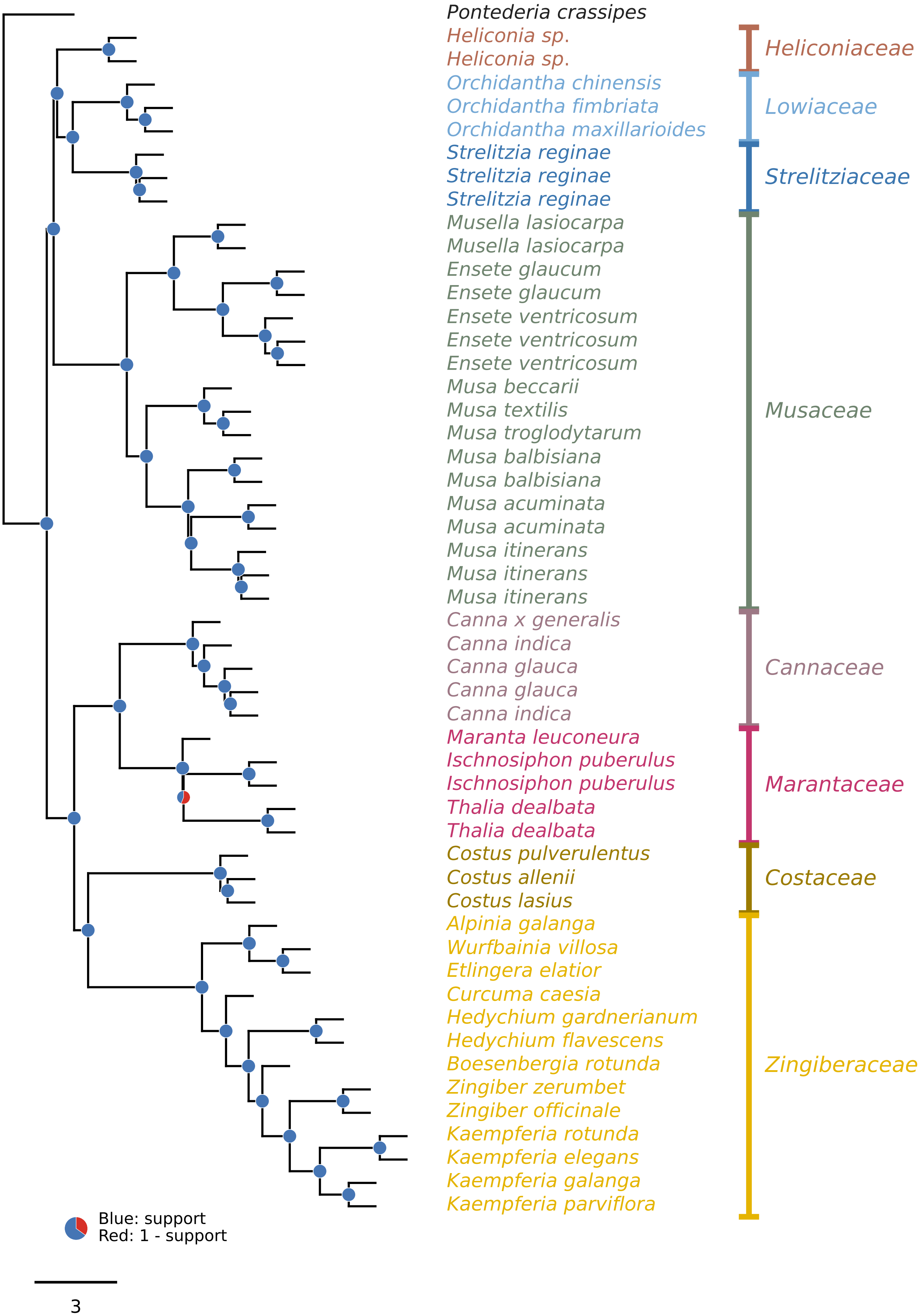

### Coalescent species tree from genes with PI sites >= 400

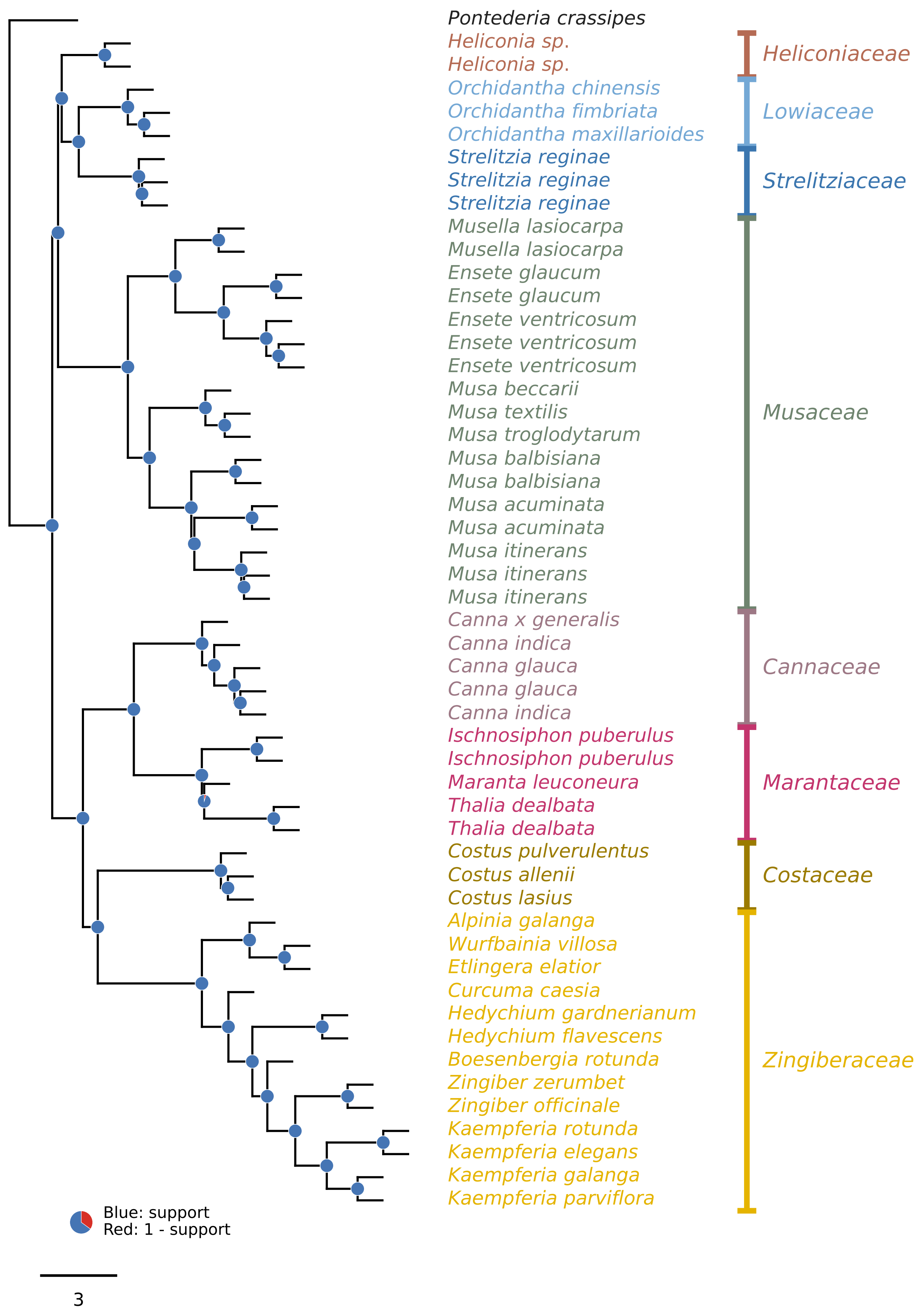

**Coalescent species tree from genes with length >= 500 bp**

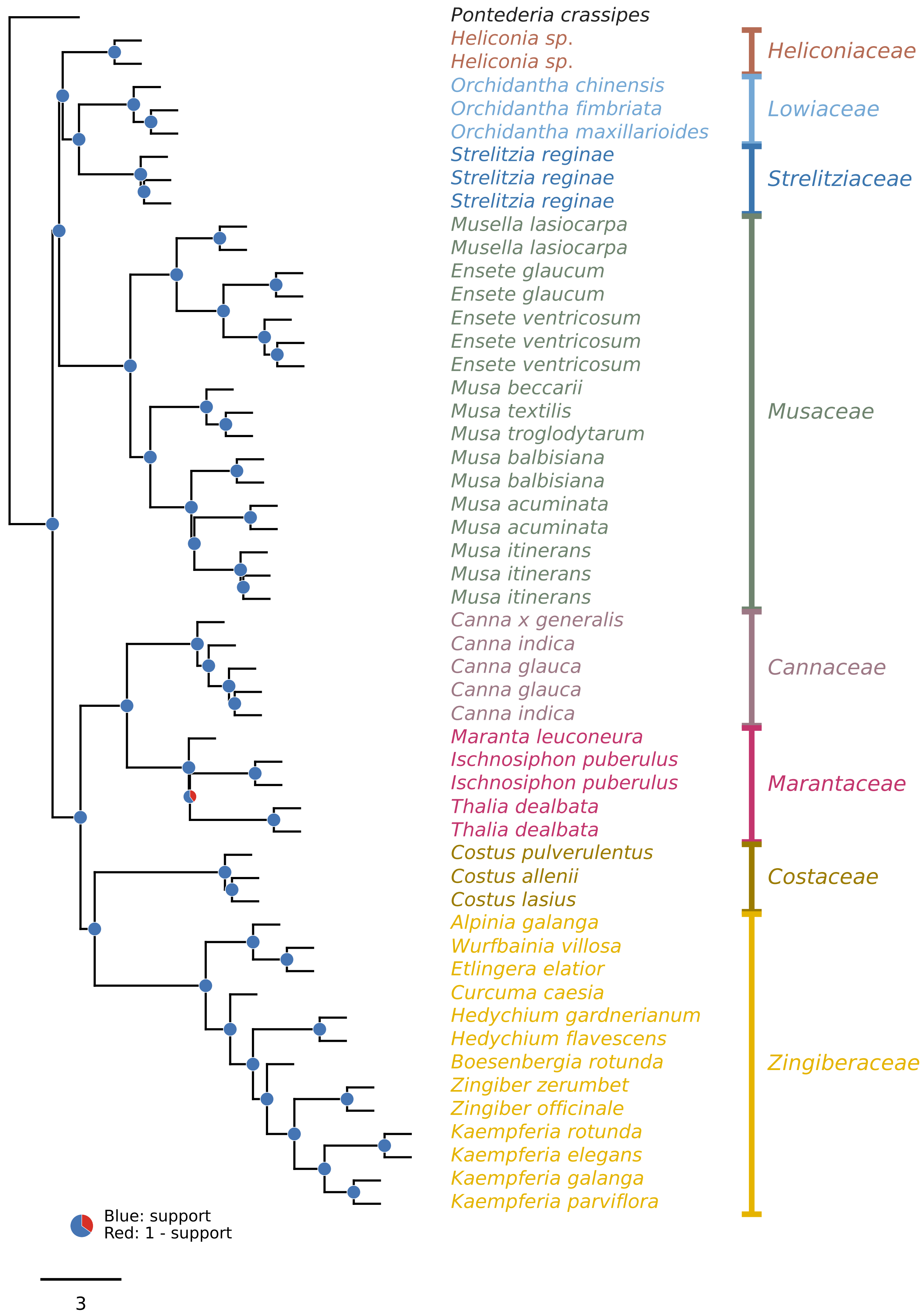

**Coalescent species tree from genes with length >= 1000 bp**

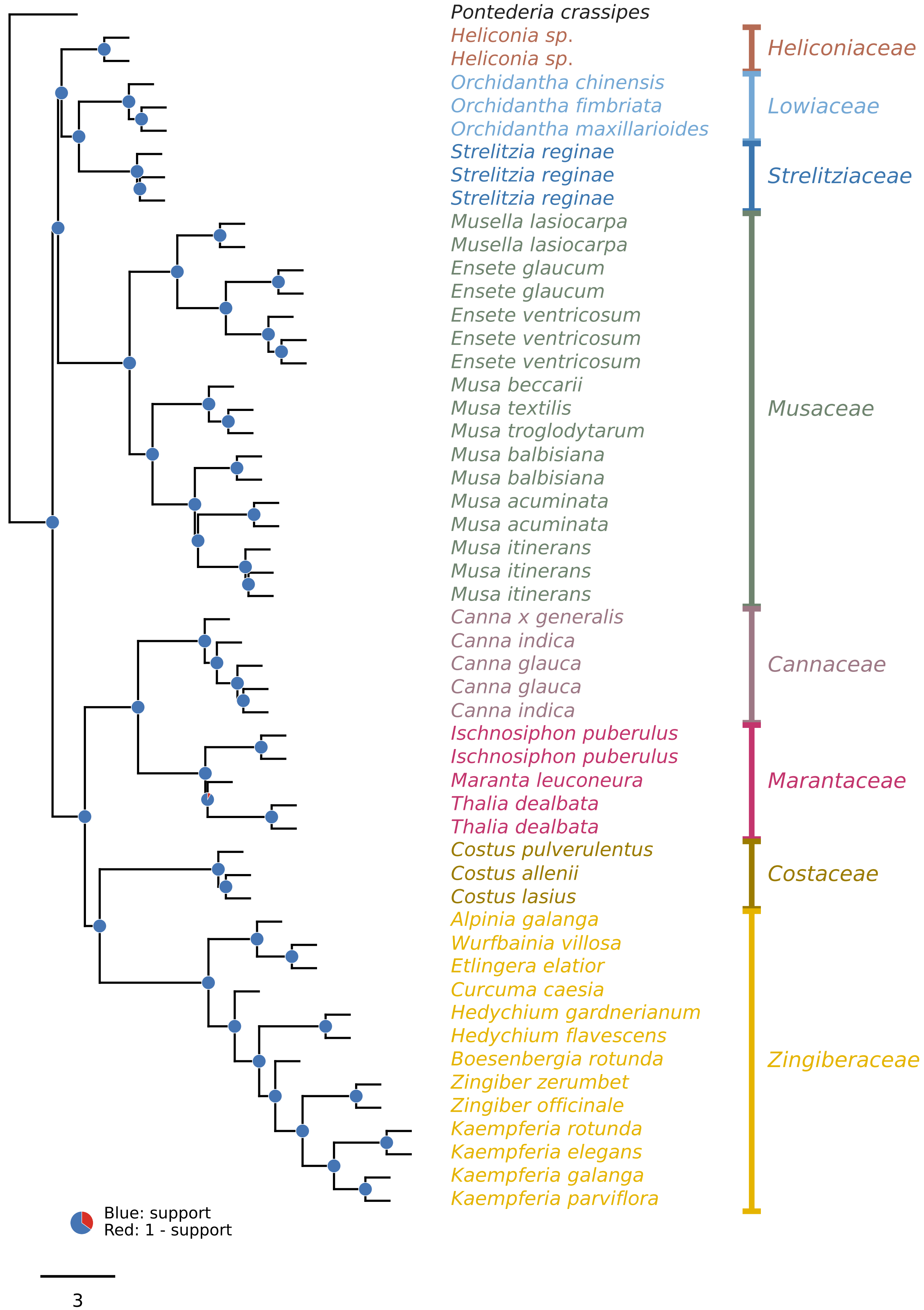
