## Supplementary material for "Nuclear phylogenomics clarifies the family-level backbone and gene-tree conflict in Zingiberales": Figure S2

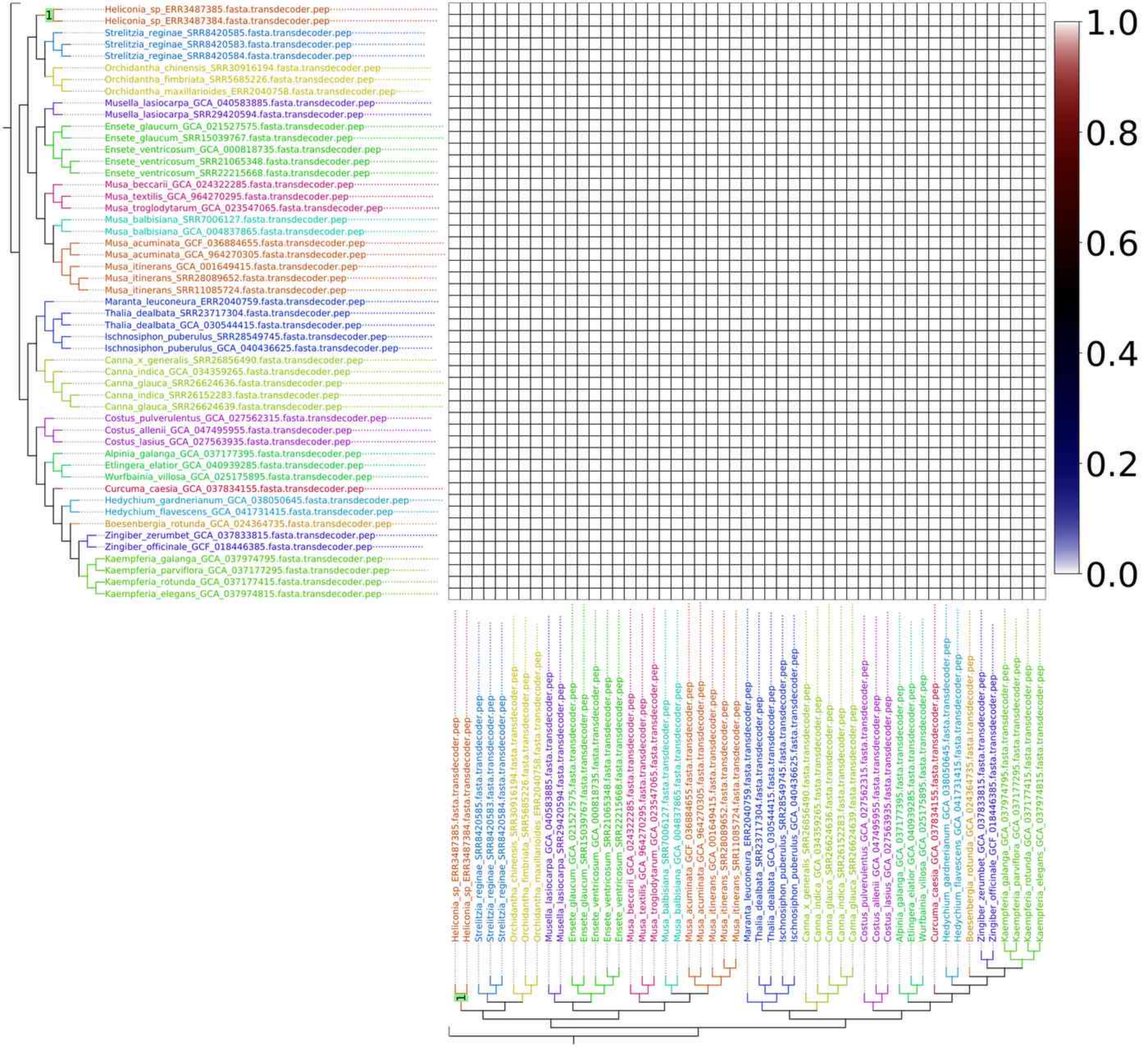
1
Heliconia_sp_ERR3487385.fasta.transdecoder.pep
Heliconia_sp_ERR3487384.fasta.transdecoder.pep
Strelitzia_reginae_SRR8420585.fasta.transdecoder.pep
Strelitzia_reginae_SRR8420583.fasta.transdecoder.pep
Strelitzia_reginae_SRR8420584.fasta.transdecoder.pep
Orchidantha_chinensis_SRR30916194.fasta.transdecoder.pep
Orchidantha_fimbriata_SRR5685226.fasta.transdecoder.pep
Orchidantha_maxillarioides_ERR2040758.fasta.transdecoder.pep
Musella_lasiocarpa_GCA_040583885.fasta.transdecoder.pep
Musella_lasiocarpa_SRR29420594.fasta.transdecoder.pep
Ensete_glaucum_GCA_021527575.fasta.transdecoder.pep
Ensete_glaucum_SRR15039767.fasta.transdecoder.pep
Ensete_ventricosum_GCA_000818735.fasta.transdecoder.pep
Ensete_ventricosum_SRR21065348.fasta.transdecoder.pep
Ensete_ventricosum_SRR22215668.fasta.transdecoder.pep
Musa_beccarii_GCA_024322285.fasta.transdecoder.pep
Musa_textilis_GCA_964270295.fasta.transdecoder.pep
Musa_troglodytarum_GCA_023547065.fasta.transdecoder.pep
Musa_balbisiana_SRR7006127.fasta.transdecoder.pep
Musa_balbisiana_GCA_004837865.fasta.transdecoder.pep
Musa_acuminata_GCF_036884655.fasta.transdecoder.pep
Musa_acuminata_GCA_964270305.fasta.transdecoder.pep
Musa_itinerans_GCA_001649415.fasta.transdecoder.pep
Musa_itinerans_SRR28089652.fasta.transdecoder.pep
Musa_itinerans_SRR11085724.fasta.transdecoder.pep
Maranta_leuconeura_ERR2040759.fasta.transdecoder.pep
Thalia_dealbata_SRR23717304.fasta.transdecoder.pep
Thalia_dealbata_GCA_030544415.fasta.transdecoder.pep
Ischnosiphon_puberulus_SRR28549745.fasta.transdecoder.pep
Ischnosiphon_puberulus_GCA_040436625.fasta.transdecoder.pep
Canna_x_generalis_SRR26856490.fasta.transdecoder.pep
Canna_indica_GCA_034359265.fasta.transdecoder.pep
Canna_glauca_SRR26624636.fasta.transdecoder.pep
Canna_indica_SRR26152283.fasta.transdecoder.pep
Canna_glauca_SRR26624639.fasta.transdecoder.pep
Costus_pulverulentus_GCA_027562315.fasta.transdecoder.pep
Costus_allenii_GCA_047495955.fasta.transdecoder.pep
Costus_lasius_GCA_027563935.fasta.transdecoder.pep
Alpinia_galanga_GCA_037177395.fasta.transdecoder.pep
Etlingera_elatior_GCA_040939285.fasta.transdecoder.pep
Wurfbainia_villosa_GCA_025175895.fasta.transdecoder.pep
Curcuma_caesia_GCA_037834155.fasta.transdecoder.pep
Hedychium_gardnerianum_GCA_038050645.fasta.transdecoder.pep
Hedychium_flavescens_GCA_041731415.fasta.transdecoder.pep
Boesenbergia_rotunda_GCA_024364735.fasta.transdecoder.pep
Zingiber_zerumbet_GCA_037833815.fasta.transdecoder.pep
Zingiber_officinale_GCF_018446385.fasta.transdecoder.pep
Kaempferia_galanga_GCA_037974795.fasta.transdecoder.pep
Kaempferia_parviflora_GCA_037177295.fasta.transdecoder.pep
Kaempferia_rotunda_GCA_037177415.fasta.transdecoder.pep
Kaempferia_elegans_GCA_037974815.fasta.transdecoder.pep
1.0
0.8
0.6
0.4
0.2
0.0

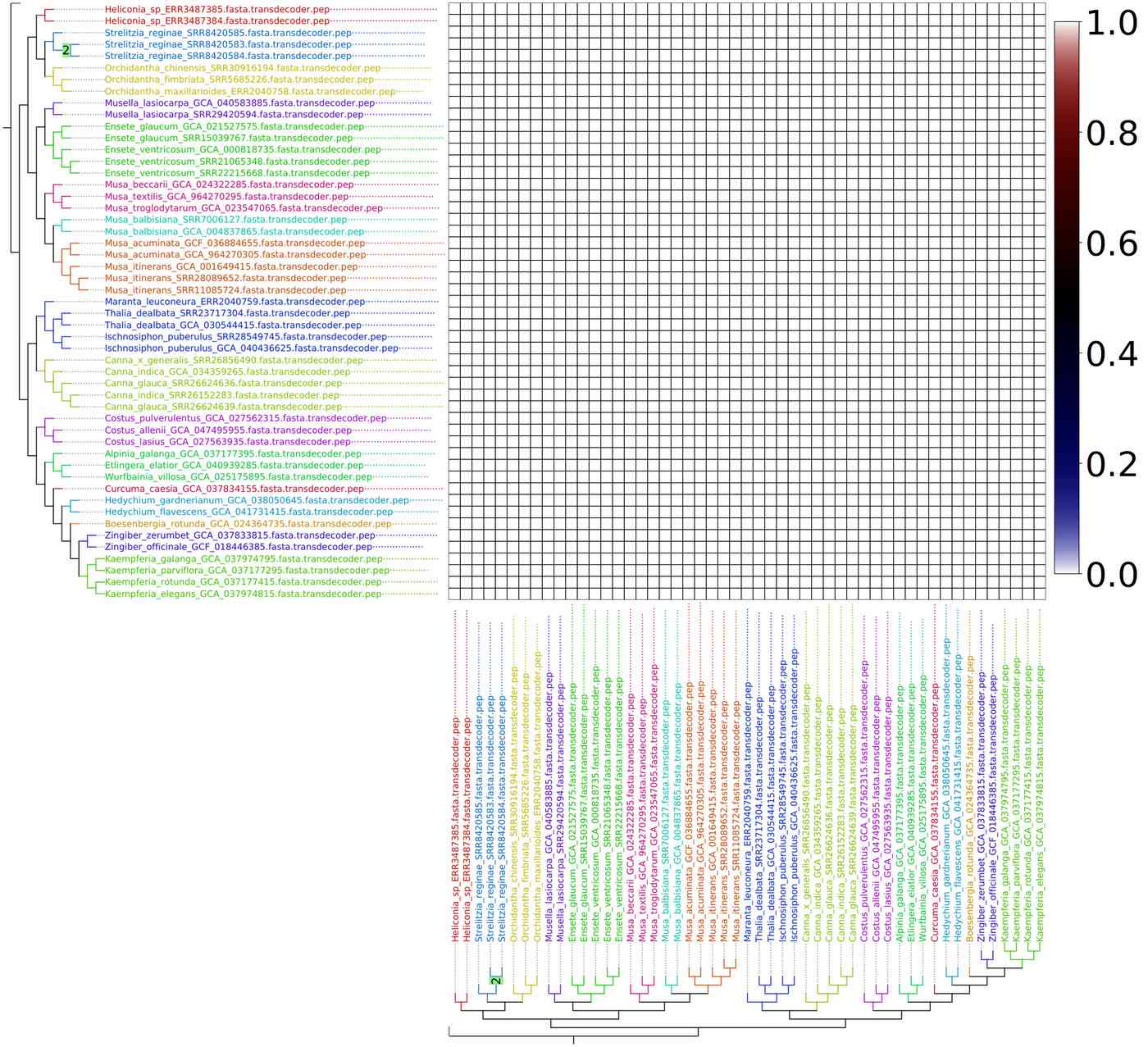
Heliconia_sp_ERR3487385.fasta.transdecoder.pep
Heliconia_sp_ERR3487384.fasta.transdecoder.pep
Strelitzia_reginae_SRR8420585.fasta.transdecoder.pep
Strelitzia_reginae_SRR8420583.fasta.transdecoder.pep
Strelitzia_reginae_SRR8420584.fasta.transdecoder.pep
Orchidantha_chinensis_SRR30916194.fasta.transdecoder.pep
Orchidantha_fimbriata_SRR5685226.fasta.transdecoder.pep
Orchidantha_maxillarioides_ERR2040758.fasta.transdecoder.pep
Musella_lasiocarpa_GCA_040583885.fasta.transdecoder.pep
Musella_lasiocarpa_SRR29420594.fasta.transdecoder.pep
Ensete_glaucum_GCA_021527575.fasta.transdecoder.pep
Ensete_glaucum_SRR15039767.fasta.transdecoder.pep
Ensete_ventricosum_GCA_000818735.fasta.transdecoder.pep
Ensete_ventricosum_SRR21065348.fasta.transdecoder.pep
Ensete_ventricosum_SRR22215668.fasta.transdecoder.pep
Musa_beccarii_GCA_024322285.fasta.transdecoder.pep
Musa_textilis_GCA_964270295.fasta.transdecoder.pep
Musa_troglodytarum_GCA_023547065.fasta.transdecoder.pep
Musa_balbisiana_SRR7006127.fasta.transdecoder.pep
Musa_balbisiana_GCA_004837865.fasta.transdecoder.pep
Musa_acuminata_GCF_036884655.fasta.transdecoder.pep
Musa_acuminata_GCA_964270305.fasta.transdecoder.pep
Musa_itinerans_GCA_001649415.fasta.transdecoder.pep
Musa_itinerans_SRR28089652.fasta.transdecoder.pep
Musa_itinerans_SRR11085724.fasta.transdecoder.pep
Maranta_leuconeura_ERR2040759.fasta.transdecoder.pep
Thalia_dealbata_SRR23717304.fasta.transdecoder.pep
Thalia_dealbata_GCA_030544415.fasta.transdecoder.pep
Ischnosiphon_puberulus_SRR28549745.fasta.transdecoder.pep
Ischnosiphon_puberulus_GCA_040436625.fasta.transdecoder.pep
Canna_x_generalis_SRR26856490.fasta.transdecoder.pep
Canna_indica_GCA_034359265.fasta.transdecoder.pep
Canna_glauca_SRR26624636.fasta.transdecoder.pep
Canna_indica_SRR26152283.fasta.transdecoder.pep
Canna_glauca_SRR26624639.fasta.transdecoder.pep
Costus_pulverulentus_GCA_027562315.fasta.transdecoder.pep
Costus_allenii_GCA_047495955.fasta.transdecoder.pep
Costus_lasius_GCA_027563935.fasta.transdecoder.pep
Alpinia_galanga_GCA_037177395.fasta.transdecoder.pep
Etlingera_elatior_GCA_040939285.fasta.transdecoder.pep
Wurfbainia_villosa_GCA_025175895.fasta.transdecoder.pep
Curcuma_caesia_GCA_037834155.fasta.transdecoder.pep
Hedychium_gardnerianum_GCA_038050645.fasta.transdecoder.pep
Hedychium_flavescens_GCA_041731415.fasta.transdecoder.pep
Boesenbergia_rotunda_GCA_024364735.fasta.transdecoder.pep
Zingiber_zerumbet_GCA_037833815.fasta.transdecoder.pep
Zingiber_officinale_GCF_018446385.fasta.transdecoder.pep
Kaempferia_galanga_GCA_037974795.fasta.transdecoder.pep
Kaempferia_parviflora_GCA_037177295.fasta.transdecoder.pep
Kaempferia_rotunda_GCA_037177415.fasta.transdecoder.pep
Kaempferia_elegans_GCA_037974815.fasta.transdecoder.pep
2
1.0
0.8
0.6
0.4
0.2
0.0

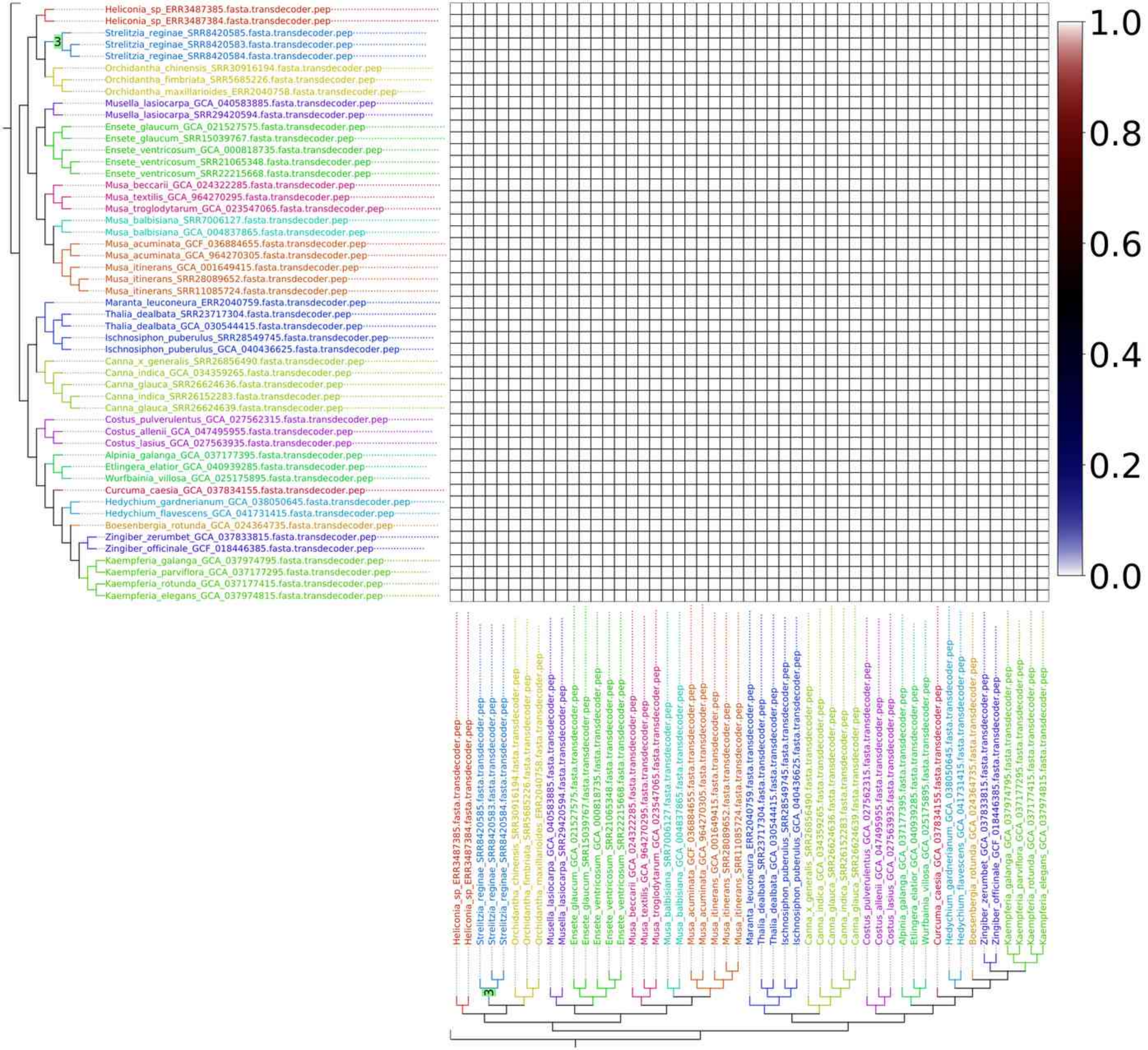
1.0
0.8
0.6
0.4
0.2
0.0
3
Heliconia_sp_ERR3487385.fasta.transdecoder.pep
Heliconia_sp_ERR3487384.fasta.transdecoder.pep
Strelitzia_reginae_SRR8420585.fasta.transdecoder.pep
Strelitzia_reginae_SRR8420583.fasta.transdecoder.pep
Strelitzia_reginae_SRR8420584.fasta.transdecoder.pep
Orchidantha_chinensis_SRR30916194.fasta.transdecoder.pep
Orchidantha_fimbriata_SRR5685226.fasta.transdecoder.pep
Orchidantha_maxillarioides_ERR2040758.fasta.transdecoder.pep
Musella_lasiocarpa_GCA_040583885.fasta.transdecoder.pep
Musella_lasiocarpa_SRR29420594.fasta.transdecoder.pep
Ensete_glaucum_GCA_021527575.fasta.transdecoder.pep
Ensete_glaucum_SRR15039767.fasta.transdecoder.pep
Ensete_ventricosum_GCA_000818735.fasta.transdecoder.pep
Ensete_ventricosum_SRR21065348.fasta.transdecoder.pep
Ensete_ventricosum_SRR22215668.fasta.transdecoder.pep
Musa_beccarii_GCA_024322285.fasta.transdecoder.pep
Musa_textilis_GCA_964270295.fasta.transdecoder.pep
Musa_troglodytarum_GCA_023547065.fasta.transdecoder.pep
Musa_balbisiana_SRR7006127.fasta.transdecoder.pep
Musa_balbisiana_GCA_004837865.fasta.transdecoder.pep
Musa_acuminata_GCF_036884655.fasta.transdecoder.pep
Musa_acuminata_GCA_964270305.fasta.transdecoder.pep
Musa_itinerans_GCA_001649415.fasta.transdecoder.pep
Musa_itinerans_SRR28089652.fasta.transdecoder.pep
Musa_itinerans_SRR11085724.fasta.transdecoder.pep
Maranta_leuconeura_ERR2040759.fasta.transdecoder.pep
Thalia_dealbata_SRR23717304.fasta.transdecoder.pep
Thalia_dealbata_GCA_030544415.fasta.transdecoder.pep
Ischnosiphon_puberulus_SRR28549745.fasta.transdecoder.pep
Ischnosiphon_puberulus_GCA_040436625.fasta.transdecoder.pep
Canna_x_generalis_SRR26856490.fasta.transdecoder.pep
Canna_indica_GCA_034359265.fasta.transdecoder.pep
Canna_glauca_SRR26624636.fasta.transdecoder.pep
Canna_indica_SRR26152283.fasta.transdecoder.pep
Canna_glauca_SRR26624639.fasta.transdecoder.pep
Costus_pulverulentus_GCA_027562315.fasta.transdecoder.pep
Costus_allenii_GCA_047495955.fasta.transdecoder.pep
Costus_lasius_GCA_027563935.fasta.transdecoder.pep
Alpinia_galanga_GCA_037177395.fasta.transdecoder.pep
Etlingera_elatior_GCA_040939285.fasta.transdecoder.pep
Wurfbainia_villosa_GCA_025175895.fasta.transdecoder.pep
Curcuma_caesia_GCA_037834155.fasta.transdecoder.pep
Hedychium_gardnerianum_GCA_038050645.fasta.transdecoder.pep
Hedychium_flavescens_GCA_041731415.fasta.transdecoder.pep
Boesenbergia_rotunda_GCA_024364735.fasta.transdecoder.pep
Zingiber_zerumbet_GCA_037833815.fasta.transdecoder.pep
Zingiber_officinale_GCF_018446385.fasta.transdecoder.pep
Kaempferia_galanga_GCA_037974795.fasta.transdecoder.pep
Kaempferia_parviflora_GCA_037177295.fasta.transdecoder.pep
Kaempferia_rotunda_GCA_037177415.fasta.transdecoder.pep
Kaempferia_elegans_GCA_037974815.fasta.transdecoder.pep

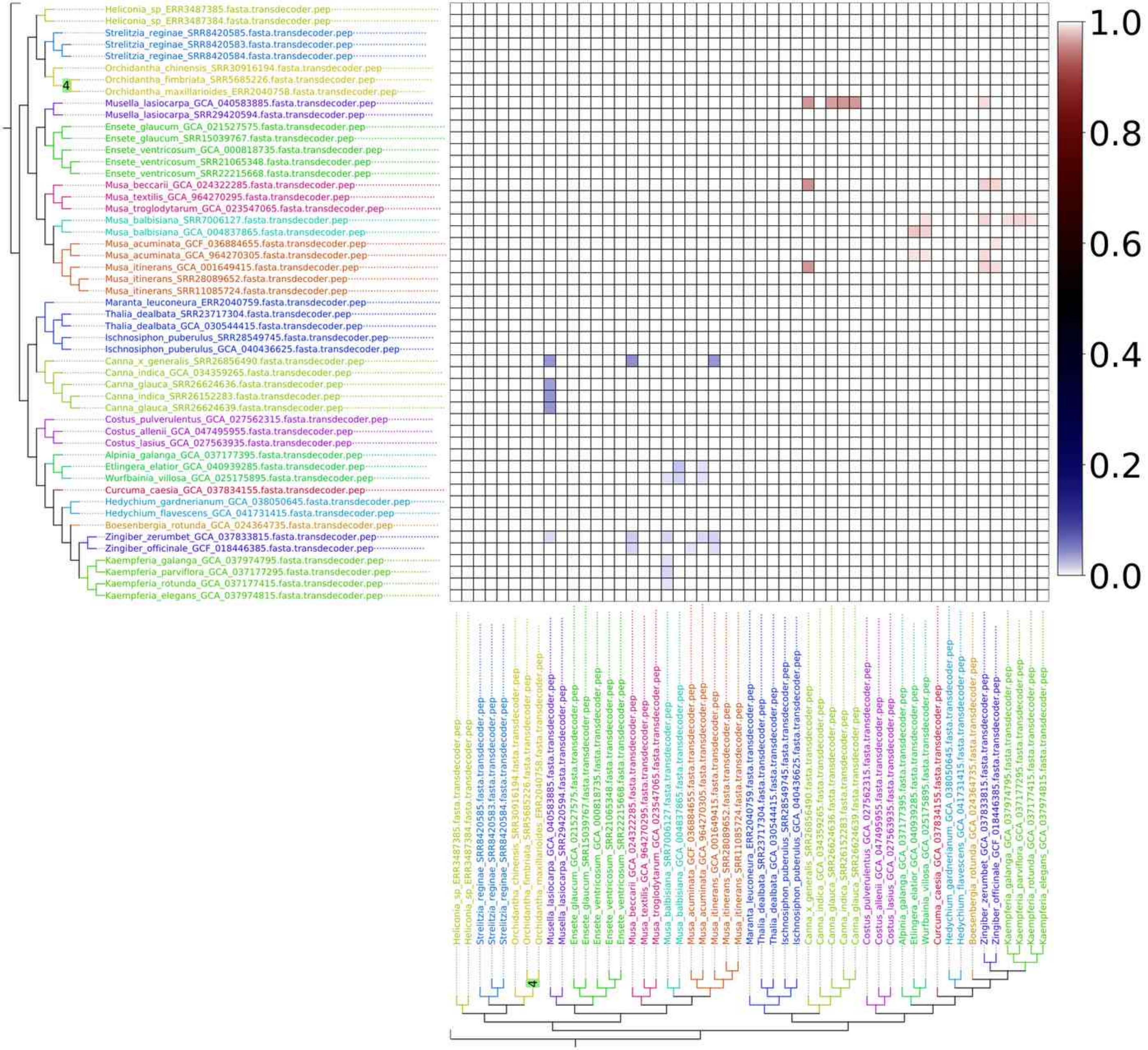
Heliconia_sp_ERR3487385.fasta.transdecoder.pep
Heliconia_sp_ERR3487384.fasta.transdecoder.pep
Strelitzia_reginae_SRR8420585.fasta.transdecoder.pep
Strelitzia_reginae_SRR8420583.fasta.transdecoder.pep
Strelitzia_reginae_SRR8420584.fasta.transdecoder.pep
Orchidantha_chinensis_SRR30916194.fasta.transdecoder.pep
Orchidantha_fimbriata_SRR5685226.fasta.transdecoder.pep
4
Orchidantha_maxillarioides_ERR2040758.fasta.transdecoder.pep
Musella_lasiocarpa_GCA_040583885.fasta.transdecoder.pep
Musella_lasiocarpa_SRR29420594.fasta.transdecoder.pep
Ensete_glaucum_GCA_021527575.fasta.transdecoder.pep
Ensete_glaucum_SRR15039767.fasta.transdecoder.pep
Ensete_ventricosum_GCA_000818735.fasta.transdecoder.pep
Ensete_ventricosum_SRR21065348.fasta.transdecoder.pep
Ensete_ventricosum_SRR22215668.fasta.transdecoder.pep
Musa_beccarii_GCA_024322285.fasta.transdecoder.pep
Musa_textilis_GCA_964270295.fasta.transdecoder.pep
Musa_troglodytarum_GCA_023547065.fasta.transdecoder.pep
Musa_balbisiana_SRR7006127.fasta.transdecoder.pep
Musa_balbisiana_GCA_004837865.fasta.transdecoder.pep
Musa_acuminata_GCF_036884655.fasta.transdecoder.pep
Musa_acuminata_GCA_964270305.fasta.transdecoder.pep
Musa_itinerans_GCA_001649415.fasta.transdecoder.pep
Musa_itinerans_SRR28089652.fasta.transdecoder.pep
Musa_itinerans_SRR11085724.fasta.transdecoder.pep
Maranta_leuconeura_ERR2040759.fasta.transdecoder.pep
Thalia_dealbata_SRR23717304.fasta.transdecoder.pep
Thalia_dealbata_GCA_030544415.fasta.transdecoder.pep
Ischnosiphon_puberulus_SRR28549745.fasta.transdecoder.pep
Ischnosiphon_puberulus_GCA_040436625.fasta.transdecoder.pep
Canna_x_generalis_SRR26856490.fasta.transdecoder.pep
Canna_indica_GCA_034359265.fasta.transdecoder.pep
Canna_glauca_SRR26624636.fasta.transdecoder.pep
Canna_indica_SRR26152283.fasta.transdecoder.pep
Canna_glauca_SRR26624639.fasta.transdecoder.pep
Costus_pulverulentus_GCA_027562315.fasta.transdecoder.pep
Costus_allenii_GCA_047495955.fasta.transdecoder.pep
Costus_lasius_GCA_027563935.fasta.transdecoder.pep
Alpinia_galanga_GCA_037177395.fasta.transdecoder.pep
Etlingera_elatior_GCA_040939285.fasta.transdecoder.pep
Wurfbainia_villosa_GCA_025175895.fasta.transdecoder.pep
Curcuma_caesia_GCA_037834155.fasta.transdecoder.pep
Hedychium_gardnerianum_GCA_038050645.fasta.transdecoder.pep
Hedychium_flavescens_GCA_041731415.fasta.transdecoder.pep
Boesenbergia_rotunda_GCA_024364735.fasta.transdecoder.pep
Zingiber_zerumbet_GCA_037833815.fasta.transdecoder.pep
Zingiber_officinale_GCF_018446385.fasta.transdecoder.pep
Kaempferia_galanga_GCA_037974795.fasta.transdecoder.pep
Kaempferia_parviflora_GCA_037177295.fasta.transdecoder.pep
Kaempferia_rotunda_GCA_037177415.fasta.transdecoder.pep
Kaempferia_elegans_GCA_037974815.fasta.transdecoder.pep
1.0
0.8
0.6
0.4
0.2
0.0

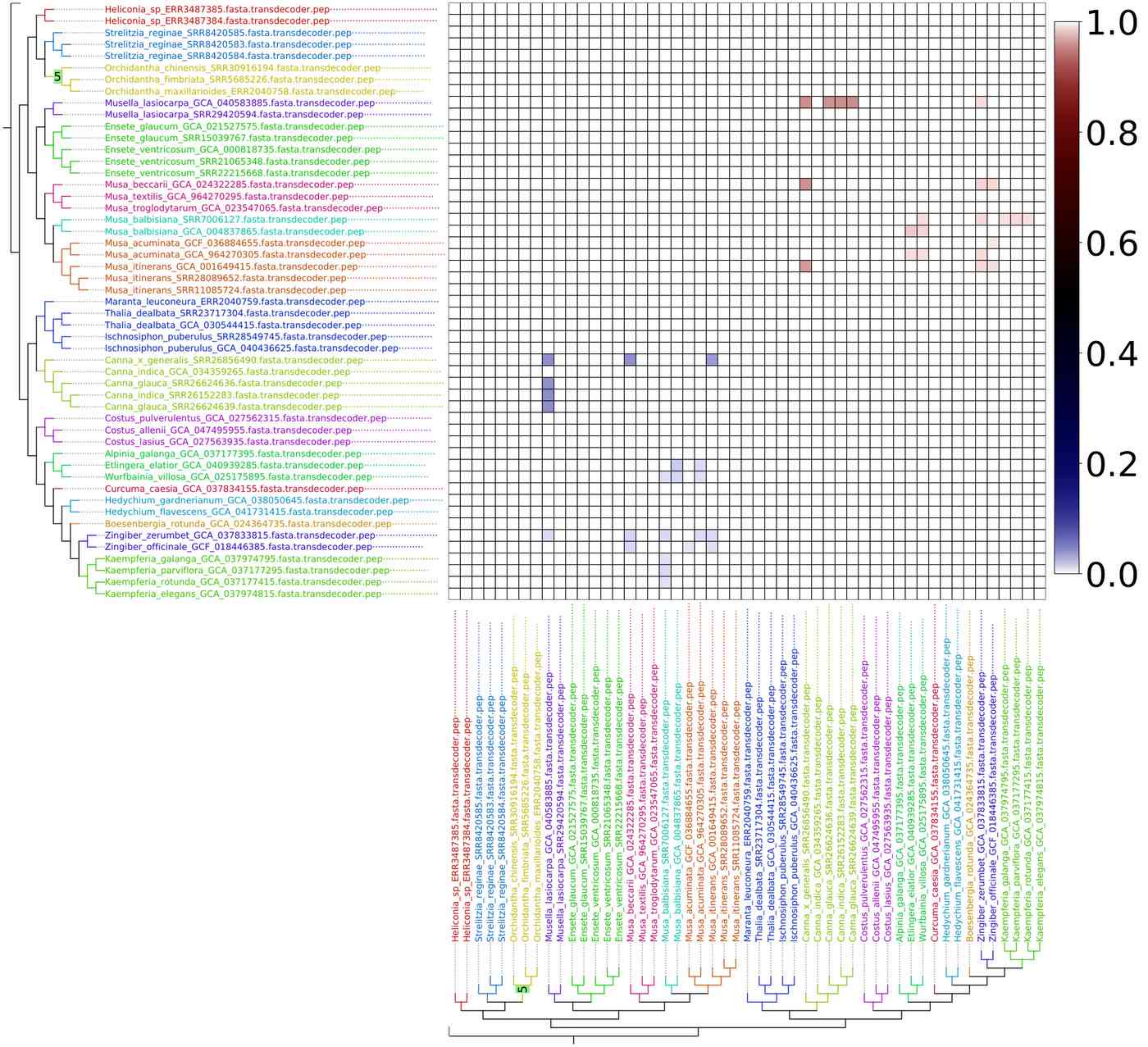
Heliconia_sp_ERR3487385.fasta.transdecoder.pep
Heliconia_sp_ERR3487384.fasta.transdecoder.pep
Strelitzia_reginae_SRR8420585.fasta.transdecoder.pep
Strelitzia_reginae_SRR8420583.fasta.transdecoder.pep
Strelitzia_reginae_SRR8420584.fasta.transdecoder.pep
Orchidantha_chinensis_SRR30916194.fasta.transdecoder.pep
Orchidantha_fimbriata_SRR5685226.fasta.transdecoder.pep
Orchidantha_maxillarioides_ERR2040758.fasta.transdecoder.pep
Musella_lasiocarpa_GCA_040583885.fasta.transdecoder.pep
Musella_lasiocarpa_SRR29420594.fasta.transdecoder.pep
Ensete_glaucum_GCA_021527575.fasta.transdecoder.pep
Ensete_glaucum_SRR15039767.fasta.transdecoder.pep
Ensete_ventricosum_GCA_000818735.fasta.transdecoder.pep
Ensete_ventricosum_SRR21065348.fasta.transdecoder.pep
Ensete_ventricosum_SRR22215668.fasta.transdecoder.pep
Musa_beccarii_GCA_024322285.fasta.transdecoder.pep
Musa_textilis_GCA_964270295.fasta.transdecoder.pep
Musa_troglodytarum_GCA_023547065.fasta.transdecoder.pep
Musa_balbisiana_SRR7006127.fasta.transdecoder.pep
Musa_balbisiana_GCA_004837865.fasta.transdecoder.pep
Musa_acuminata_GCF_036884655.fasta.transdecoder.pep
Musa_acuminata_GCA_964270305.fasta.transdecoder.pep
Musa_itinerans_GCA_001649415.fasta.transdecoder.pep
Musa_itinerans_SRR28089652.fasta.transdecoder.pep
Musa_itinerans_SRR11085724.fasta.transdecoder.pep
Maranta_leuconeura_ERR2040759.fasta.transdecoder.pep
Thalia_dealbata_SRR23717304.fasta.transdecoder.pep
Thalia_dealbata_GCA_030544415.fasta.transdecoder.pep
Ischnosiphon_puberulus_SRR28549745.fasta.transdecoder.pep
Ischnosiphon_puberulus_GCA_040436625.fasta.transdecoder.pep
Canna_x_generalis_SRR26856490.fasta.transdecoder.pep
Canna_indica_GCA_034359265.fasta.transdecoder.pep
Canna_glauca_SRR26624636.fasta.transdecoder.pep
Canna_indica_SRR26152283.fasta.transdecoder.pep
Canna_glauca_SRR26624639.fasta.transdecoder.pep
Costus_pulverulentus_GCA_027562315.fasta.transdecoder.pep
Costus_allenii_GCA_047495955.fasta.transdecoder.pep
Costus_lasius_GCA_027563935.fasta.transdecoder.pep
Alpinia_galanga_GCA_037177395.fasta.transdecoder.pep
Etlingera_elatior_GCA_040939285.fasta.transdecoder.pep
Wurfbainia_villosa_GCA_025175895.fasta.transdecoder.pep
Curcuma_caesia_GCA_037834155.fasta.transdecoder.pep
Hedychium_gardnerianum_GCA_038050645.fasta.transdecoder.pep
Hedychium_flavescens_GCA_041731415.fasta.transdecoder.pep
Boesenbergia_rotunda_GCA_024364735.fasta.transdecoder.pep
Zingiber_zerumbet_GCA_037833815.fasta.transdecoder.pep
Zingiber_officinale_GCF_018446385.fasta.transdecoder.pep
Kaempferia_galanga_GCA_037974795.fasta.transdecoder.pep
Kaempferia_parviflora_GCA_037177295.fasta.transdecoder.pep
Kaempferia_rotunda_GCA_037177415.fasta.transdecoder.pep
Kaempferia_elegans_GCA_037974815.fasta.transdecoder.pep
5
1.0
0.8
0.6
0.4
0.2
0.0

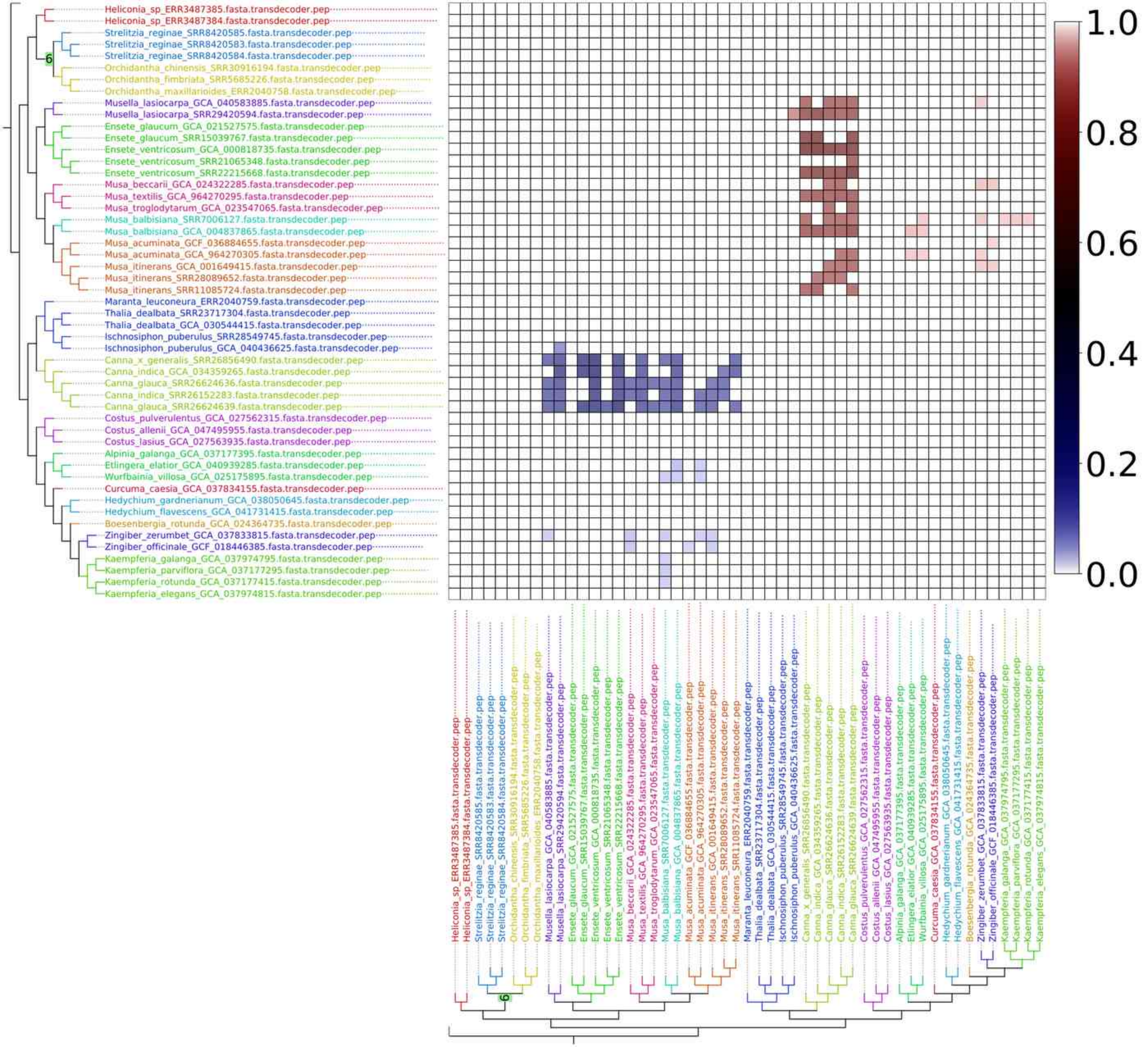
Heliconia_sp_ERR3487385.fasta.transdecoder.pep
Heliconia_sp_ERR3487384.fasta.transdecoder.pep
Strelitzia_reginae_SRR8420585.fasta.transdecoder.pep
Strelitzia_reginae_SRR8420583.fasta.transdecoder.pep
Strelitzia_reginae_SRR8420584.fasta.transdecoder.pep
Orchidantha_chinensis_SRR30916194.fasta.transdecoder.pep
Orchidantha_fimbriata_SRR5685226.fasta.transdecoder.pep
Orchidantha_maxillarioides_ERR2040758.fasta.transdecoder.pep
Musella_lasiocarpa_GCA_040583885.fasta.transdecoder.pep
Musella_lasiocarpa_SRR29420594.fasta.transdecoder.pep
Ensete_glaucum_GCA_021527575.fasta.transdecoder.pep
Ensete_glaucum_SRR15039767.fasta.transdecoder.pep
Ensete_ventricosum_GCA_000818735.fasta.transdecoder.pep
Ensete_ventricosum_SRR21065348.fasta.transdecoder.pep
Ensete_ventricosum_SRR22215668.fasta.transdecoder.pep
Musa_beccarii_GCA_024322285.fasta.transdecoder.pep
Musa_textilis_GCA_964270295.fasta.transdecoder.pep
Musa_troglodytarum_GCA_023547065.fasta.transdecoder.pep
Musa_balbisiana_SRR7006127.fasta.transdecoder.pep
Musa_balbisiana_GCA_004837865.fasta.transdecoder.pep
Musa_acuminata_GCF_036884655.fasta.transdecoder.pep
Musa_acuminata_GCA_964270305.fasta.transdecoder.pep
Musa_itinerans_GCA_001649415.fasta.transdecoder.pep
Musa_itinerans_SRR28089652.fasta.transdecoder.pep
Musa_itinerans_SRR11085724.fasta.transdecoder.pep
Maranta_leuconeura_ERR2040759.fasta.transdecoder.pep
Thalia_dealbata_SRR23717304.fasta.transdecoder.pep
Thalia_dealbata_GCA_030544415.fasta.transdecoder.pep
Ischnosiphon_puberulus_SRR28549745.fasta.transdecoder.pep
Ischnosiphon_puberulus_GCA_040436625.fasta.transdecoder.pep
Canna_x_generalis_SRR26856490.fasta.transdecoder.pep
Canna_indica_GCA_034359265.fasta.transdecoder.pep
Canna_glauca_SRR26624636.fasta.transdecoder.pep
Canna_indica_SRR26152283.fasta.transdecoder.pep
Canna_glauca_SRR26624639.fasta.transdecoder.pep
Costus_pulverulentus_GCA_027562315.fasta.transdecoder.pep
Costus_allenii_GCA_047495955.fasta.transdecoder.pep
Costus_lasius_GCA_027563935.fasta.transdecoder.pep
Alpinia_galanga_GCA_037177395.fasta.transdecoder.pep
Etlingera_elatior_GCA_040939285.fasta.transdecoder.pep
Wurfbainia_villosa_GCA_025175895.fasta.transdecoder.pep
Curcuma_caesia_GCA_037834155.fasta.transdecoder.pep
Hedychium_gardnerianum_GCA_038050645.fasta.transdecoder.pep
Hedychium_flavescens_GCA_041731415.fasta.transdecoder.pep
Boesenbergia_rotunda_GCA_024364735.fasta.transdecoder.pep
Zingiber_zerumbet_GCA_037833815.fasta.transdecoder.pep
Zingiber_officinale_GCF_018446385.fasta.transdecoder.pep
Kaempferia_galanga_GCA_037974795.fasta.transdecoder.pep
Kaempferia_parviflora_GCA_037177295.fasta.transdecoder.pep
Kaempferia_rotunda_GCA_037177415.fasta.transdecoder.pep
Kaempferia_elegans_GCA_037974815.fasta.transdecoder.pep
6
1.0
0.8
0.6
0.4
0.2
0.0

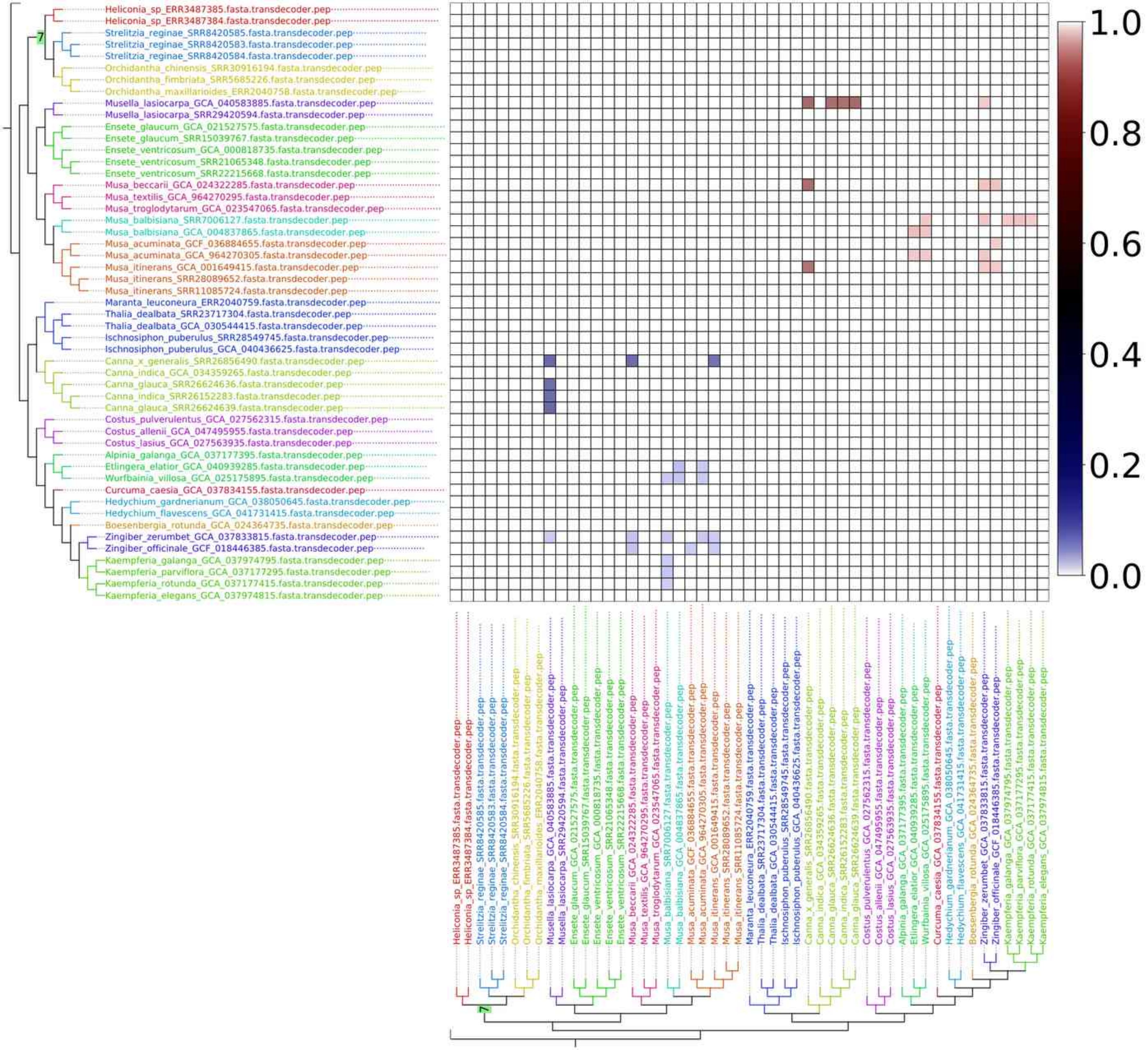
Heliconia_sp_ERR3487385.fasta.transdecoder.pep
Heliconia_sp_ERR3487384.fasta.transdecoder.pep
Strelitzia_reginae_SRR8420585.fasta.transdecoder.pep
Strelitzia_reginae_SRR8420583.fasta.transdecoder.pep
Strelitzia_reginae_SRR8420584.fasta.transdecoder.pep
Orchidantha_chinensis_SRR30916194.fasta.transdecoder.pep
Orchidantha_fimbriata_SRR5685226.fasta.transdecoder.pep
Orchidantha_maxillarioides_ERR2040758.fasta.transdecoder.pep
Musella_lasiocarpa_GCA_040583885.fasta.transdecoder.pep
Musella_lasiocarpa_SRR29420594.fasta.transdecoder.pep
Ensete_glaucum_GCA_021527575.fasta.transdecoder.pep
Ensete_glaucum_SRR15039767.fasta.transdecoder.pep
Ensete_ventricosum_GCA_000818735.fasta.transdecoder.pep
Ensete_ventricosum_SRR21065348.fasta.transdecoder.pep
Ensete_ventricosum_SRR22215668.fasta.transdecoder.pep
Musa_beccarii_GCA_024322285.fasta.transdecoder.pep
Musa_textilis_GCA_964270295.fasta.transdecoder.pep
Musa_troglodytarum_GCA_023547065.fasta.transdecoder.pep
Musa_balbisiana_SRR7006127.fasta.transdecoder.pep
Musa_balbisiana_GCA_004837865.fasta.transdecoder.pep
Musa_acuminata_GCF_036884655.fasta.transdecoder.pep
Musa_acuminata_GCA_964270305.fasta.transdecoder.pep
Musa_itinerans_GCA_001649415.fasta.transdecoder.pep
Musa_itinerans_SRR28089652.fasta.transdecoder.pep
Musa_itinerans_SRR11085724.fasta.transdecoder.pep
Maranta_leuconeura_ERR2040759.fasta.transdecoder.pep
Thalia_dealbata_SRR23717304.fasta.transdecoder.pep
Thalia_dealbata_GCA_030544415.fasta.transdecoder.pep
Ischnosiphon_puberulus_SRR28549745.fasta.transdecoder.pep
Ischnosiphon_puberulus_GCA_040436625.fasta.transdecoder.pep
Canna_x_generalis_SRR26856490.fasta.transdecoder.pep
Canna_indica_GCA_034359265.fasta.transdecoder.pep
Canna_glauca_SRR26624636.fasta.transdecoder.pep
Canna_indica_SRR26152283.fasta.transdecoder.pep
Canna_glauca_SRR26624639.fasta.transdecoder.pep
Costus_pulverulentus_GCA_027562315.fasta.transdecoder.pep
Costus_allenii_GCA_047495955.fasta.transdecoder.pep
Costus_lasius_GCA_027563935.fasta.transdecoder.pep
Alpinia_galanga_GCA_037177395.fasta.transdecoder.pep
Etlingera_elatior_GCA_040939285.fasta.transdecoder.pep
Wurfbainia_villosa_GCA_025175895.fasta.transdecoder.pep
Curcuma_caesia_GCA_037834155.fasta.transdecoder.pep
Hedychium_gardnerianum_GCA_038050645.fasta.transdecoder.pep
Hedychium_flavescens_GCA_041731415.fasta.transdecoder.pep
Boesenbergia_rotunda_GCA_024364735.fasta.transdecoder.pep
Zingiber_zerumbet_GCA_037833815.fasta.transdecoder.pep
Zingiber_officinale_GCF_018446385.fasta.transdecoder.pep
Kaempferia_galanga_GCA_037974795.fasta.transdecoder.pep
Kaempferia_parviflora_GCA_037177295.fasta.transdecoder.pep
Kaempferia_rotunda_GCA_037177415.fasta.transdecoder.pep
Kaempferia_elegans_GCA_037974815.fasta.transdecoder.pep
7
1.0
0.8
0.6
0.4
0.2
0.0

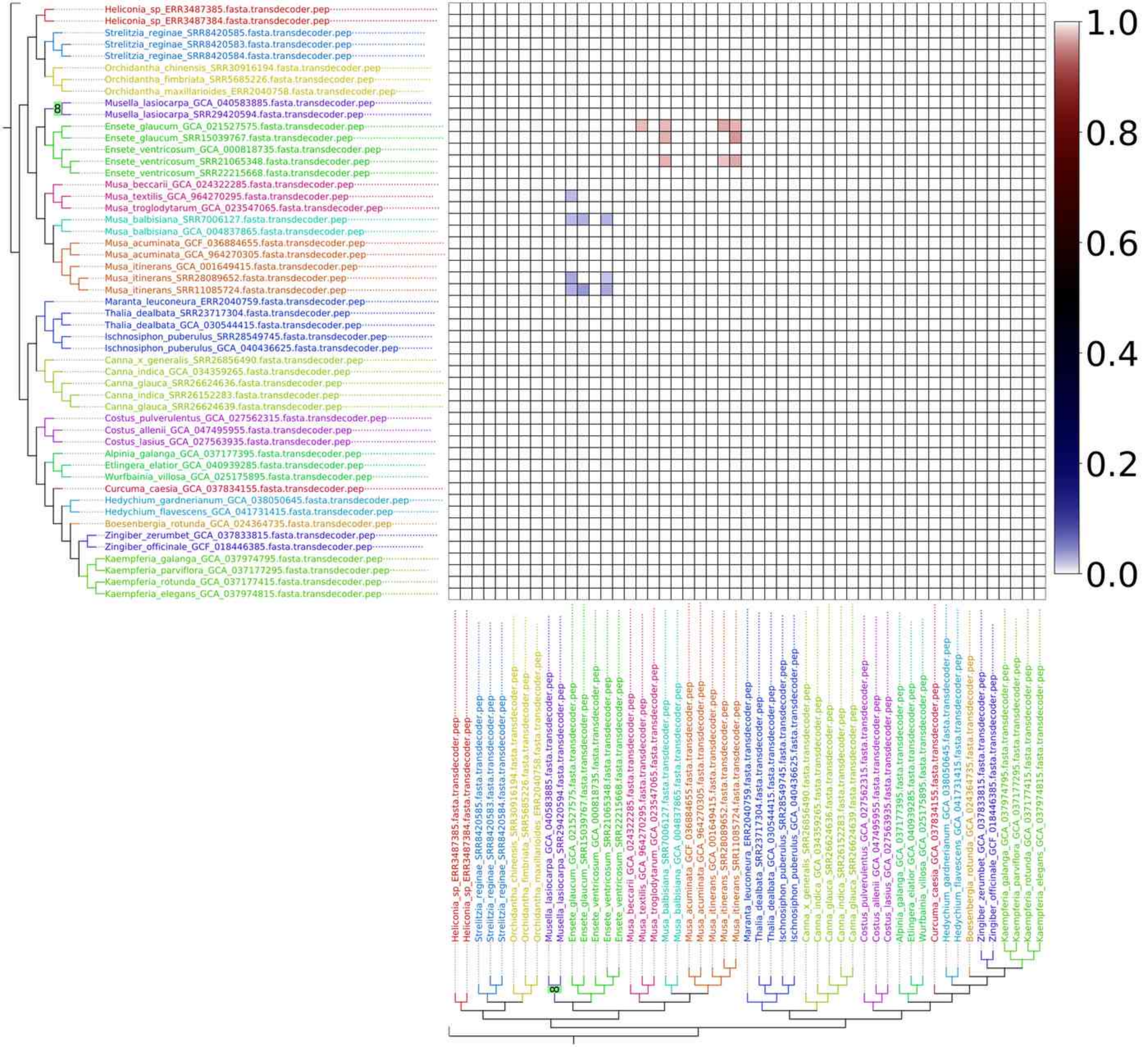
Heliconia_sp_ERR3487385.fasta.transdecoder.pep
Heliconia_sp_ERR3487384.fasta.transdecoder.pep
Strelitzia_reginae_SRR8420585.fasta.transdecoder.pep
Strelitzia_reginae_SRR8420583.fasta.transdecoder.pep
Strelitzia_reginae_SRR8420584.fasta.transdecoder.pep
Orchidantha_chinensis_SRR30916194.fasta.transdecoder.pep
Orchidantha_fimbriata_SRR5685226.fasta.transdecoder.pep
Orchidantha_maxillarioides_ERR2040758.fasta.transdecoder.pep
Musella_lasiocarpa_GCA_040583885.fasta.transdecoder.pep
Musella_lasiocarpa_SRR29420594.fasta.transdecoder.pep
Ensete_glaucum_GCA_021527575.fasta.transdecoder.pep
Ensete_glaucum_SRR15039767.fasta.transdecoder.pep
Ensete_ventricosum_GCA_000818735.fasta.transdecoder.pep
Ensete_ventricosum_SRR21065348.fasta.transdecoder.pep
Ensete_ventricosum_SRR22215668.fasta.transdecoder.pep
Musa_beccarii_GCA_024322285.fasta.transdecoder.pep
Musa_textilis_GCA_964270295.fasta.transdecoder.pep
Musa_troglodytarum_GCA_023547065.fasta.transdecoder.pep
Musa_balbisiana_SRR7006127.fasta.transdecoder.pep
Musa_balbisiana_GCA_004837865.fasta.transdecoder.pep
Musa_acuminata_GCF_036884655.fasta.transdecoder.pep
Musa_acuminata_GCA_964270305.fasta.transdecoder.pep
Musa_itinerans_GCA_001649415.fasta.transdecoder.pep
Musa_itinerans_SRR28089652.fasta.transdecoder.pep
Musa_itinerans_SRR11085724.fasta.transdecoder.pep
Maranta_leuconeura_ERR2040759.fasta.transdecoder.pep
Thalia_dealbata_SRR23717304.fasta.transdecoder.pep
Thalia_dealbata_GCA_030544415.fasta.transdecoder.pep
Ischnosiphon_puberulus_SRR28549745.fasta.transdecoder.pep
Ischnosiphon_puberulus_GCA_040436625.fasta.transdecoder.pep
Canna_x_generalis_SRR26856490.fasta.transdecoder.pep
Canna_indica_GCA_034359265.fasta.transdecoder.pep
Canna_glauca_SRR26624636.fasta.transdecoder.pep
Canna_indica_SRR26152283.fasta.transdecoder.pep
Canna_glauca_SRR26624639.fasta.transdecoder.pep
Costus_pulverulentus_GCA_027562315.fasta.transdecoder.pep
Costus_allenii_GCA_047495955.fasta.transdecoder.pep
Costus_lasius_GCA_027563935.fasta.transdecoder.pep
Alpinia_galanga_GCA_037177395.fasta.transdecoder.pep
Etlingera_elatior_GCA_040939285.fasta.transdecoder.pep
Wurfbainia_villosa_GCA_025175895.fasta.transdecoder.pep
Curcuma_caesia_GCA_037834155.fasta.transdecoder.pep
Hedychium_gardnerianum_GCA_038050645.fasta.transdecoder.pep
Hedychium_flavescens_GCA_041731415.fasta.transdecoder.pep
Boesenbergia_rotunda_GCA_024364735.fasta.transdecoder.pep
Zingiber_zerumbet_GCA_037833815.fasta.transdecoder.pep
Zingiber_officinale_GCF_018446385.fasta.transdecoder.pep
Kaempferia_galanga_GCA_037974795.fasta.transdecoder.pep
Kaempferia_parviflora_GCA_037177295.fasta.transdecoder.pep
Kaempferia_rotunda_GCA_037177415.fasta.transdecoder.pep
Kaempferia_elegans_GCA_037974815.fasta.transdecoder.pep
8
1.0
0.8
0.6
0.4
0.2
0.0

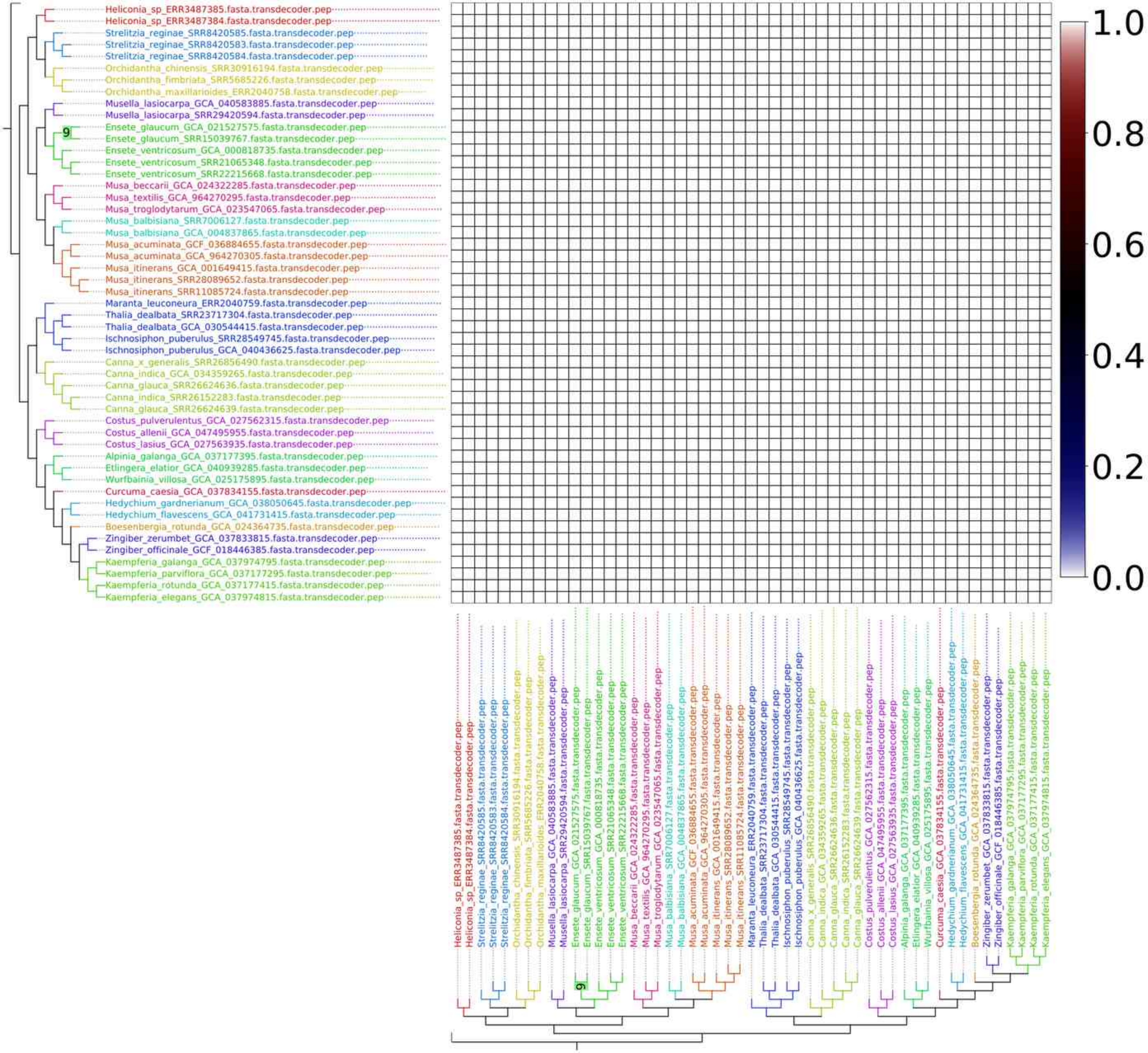
Heliconia_sp_ERR3487385.fasta.transdecoder.pep
Heliconia_sp_ERR3487384.fasta.transdecoder.pep
Strelitzia_reginae_SRR8420585.fasta.transdecoder.pep
Strelitzia_reginae_SRR8420583.fasta.transdecoder.pep
Strelitzia_reginae_SRR8420584.fasta.transdecoder.pep
Orchidantha_chinensis_SRR30916194.fasta.transdecoder.pep
Orchidantha_fimbriata_SRR5685226.fasta.transdecoder.pep
Orchidantha_maxillarioides_ERR2040758.fasta.transdecoder.pep
Musella_lasiocarpa_GCA_040583885.fasta.transdecoder.pep
Musella_lasiocarpa_SRR29420594.fasta.transdecoder.pep
9
Ensete_glaucum_GCA_021527575.fasta.transdecoder.pep
Ensete_glaucum_SRR15039767.fasta.transdecoder.pep
Ensete_ventricosum_GCA_000818735.fasta.transdecoder.pep
Ensete_ventricosum_SRR21065348.fasta.transdecoder.pep
Ensete_ventricosum_SRR22215668.fasta.transdecoder.pep
Musa_beccarii_GCA_024322285.fasta.transdecoder.pep
Musa_textilis_GCA_964270295.fasta.transdecoder.pep
Musa_troglodytarum_GCA_023547065.fasta.transdecoder.pep
Musa_balbisiana_SRR7006127.fasta.transdecoder.pep
Musa_balbisiana_GCA_004837865.fasta.transdecoder.pep
Musa_acuminata_GCF_036884655.fasta.transdecoder.pep
Musa_acuminata_GCA_964270305.fasta.transdecoder.pep
Musa_itinerans_GCA_001649415.fasta.transdecoder.pep
Musa_itinerans_SRR28089652.fasta.transdecoder.pep
Musa_itinerans_SRR11085724.fasta.transdecoder.pep
Maranta_leuconeura_ERR2040759.fasta.transdecoder.pep
Thalia_dealbata_SRR23717304.fasta.transdecoder.pep
Thalia_dealbata_GCA_030544415.fasta.transdecoder.pep
Ischnosiphon_puberulus_SRR28549745.fasta.transdecoder.pep
Ischnosiphon_puberulus_GCA_040436625.fasta.transdecoder.pep
Canna_x_generalis_SRR26856490.fasta.transdecoder.pep
Canna_indica_GCA_034359265.fasta.transdecoder.pep
Canna_glauca_SRR26624636.fasta.transdecoder.pep
Canna_indica_SRR26152283.fasta.transdecoder.pep
Canna_glauca_SRR26624639.fasta.transdecoder.pep
Costus_pulverulentus_GCA_027562315.fasta.transdecoder.pep
Costus_allenii_GCA_047495955.fasta.transdecoder.pep
Costus_lasius_GCA_027563935.fasta.transdecoder.pep
Alpinia_galanga_GCA_037177395.fasta.transdecoder.pep
Etlingera_elatior_GCA_040939285.fasta.transdecoder.pep
Wurfbainia_villosa_GCA_025175895.fasta.transdecoder.pep
Curcuma_caesia_GCA_037834155.fasta.transdecoder.pep
Hedychium_gardnerianum_GCA_038050645.fasta.transdecoder.pep
Hedychium_flavescens_GCA_041731415.fasta.transdecoder.pep
Boesenbergia_rotunda_GCA_024364735.fasta.transdecoder.pep
Zingiber_zerumbet_GCA_037833815.fasta.transdecoder.pep
Zingiber_officinale_GCF_018446385.fasta.transdecoder.pep
Kaempferia_galanga_GCA_037974795.fasta.transdecoder.pep
Kaempferia_parviflora_GCA_037177295.fasta.transdecoder.pep
Kaempferia_rotunda_GCA_037177415.fasta.transdecoder.pep
Kaempferia_elegans_GCA_037974815.fasta.transdecoder.pep
1.0
0.8
0.6
0.4
0.2
0.0

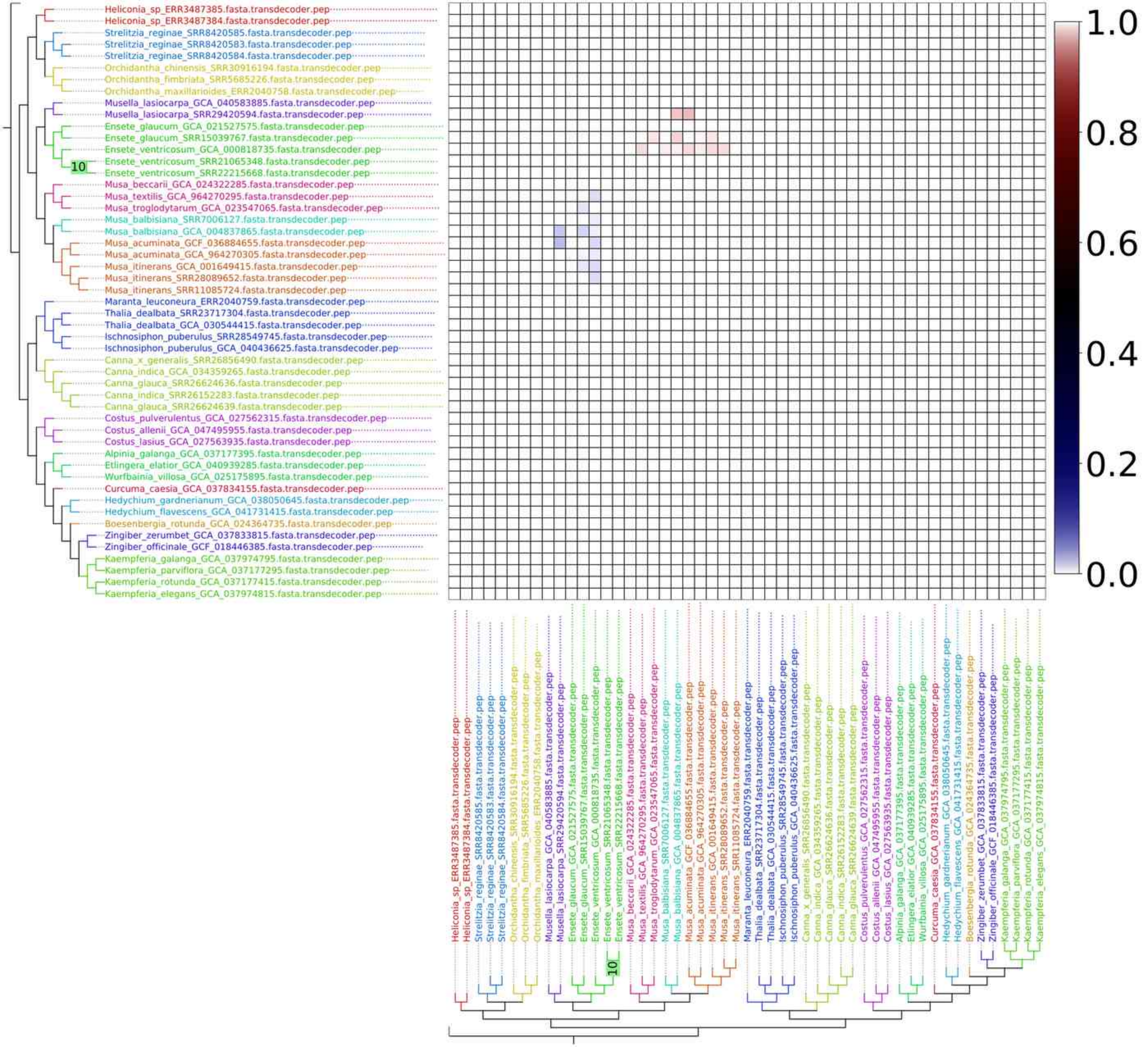

Heliconia_sp_ERR3487385.fasta.transdecoder.pep
Heliconia_sp_ERR3487384.fasta.transdecoder.pep
Strelitzia_reginae_SRR8420585.fasta.transdecoder.pep
Strelitzia_reginae_SRR8420583.fasta.transdecoder.pep
Strelitzia_reginae_SRR8420584.fasta.transdecoder.pep
Orchidantha_chinensis_SRR30916194.fasta.transdecoder.pep
Orchidantha_fimbriata_SRR5685226.fasta.transdecoder.pep
Orchidantha_maxillarioides_ERR2040758.fasta.transdecoder.pep
Musella_lasiocarpa_GCA_040583885.fasta.transdecoder.pep
Musella_lasiocarpa_SRR29420594.fasta.transdecoder.pep
Ensete_glaucum_GCA_021527575.fasta.transdecoder.pep
Ensete_glaucum_SRR15039767.fasta.transdecoder.pep
Ensete_ventricosum_GCA_000818735.fasta.transdecoder.pep
Ensete_ventricosum_SRR21065348.fasta.transdecoder.pep
Ensete_ventricosum_SRR22215668.fasta.transdecoder.pep
10
Musa_beccarii_GCA_024322285.fasta.transdecoder.pep
Musa_textilis_GCA_964270295.fasta.transdecoder.pep
Musa_troglodytarum_GCA_023547065.fasta.transdecoder.pep
Musa_balbisiana_SRR7006127.fasta.transdecoder.pep
Musa_balbisiana_GCA_004837865.fasta.transdecoder.pep
Musa_acuminata_GCF_036884655.fasta.transdecoder.pep
Musa_acuminata_GCA_964270305.fasta.transdecoder.pep
Musa_itinerans_GCA_001649415.fasta.transdecoder.pep
Musa_itinerans_SRR28089652.fasta.transdecoder.pep
Musa_itinerans_SRR11085724.fasta.transdecoder.pep
Maranta_leuconeura_ERR2040759.fasta.transdecoder.pep
Thalia_dealbata_SRR23717304.fasta.transdecoder.pep
Thalia_dealbata_GCA_030544415.fasta.transdecoder.pep
Ischnosiphon_puberulus_SRR28549745.fasta.transdecoder.pep
Ischnosiphon_puberulus_GCA_040436625.fasta.transdecoder.pep
Canna_x_generalis_SRR26856490.fasta.transdecoder.pep
Canna_indica_GCA_034359265.fasta.transdecoder.pep
Canna_glauca_SRR26624636.fasta.transdecoder.pep
Canna_indica_SRR26152283.fasta.transdecoder.pep
Canna_glauca_SRR26624639.fasta.transdecoder.pep
Costus_pulverulentus_GCA_027562315.fasta.transdecoder.pep
Costus_allenii_GCA_047495955.fasta.transdecoder.pep
Costus_lasius_GCA_027563935.fasta.transdecoder.pep
Alpinia_galanga_GCA_037177395.fasta.transdecoder.pep
Etlingera_elatior_GCA_040939285.fasta.transdecoder.pep
Wurfbainia_villosa_GCA_025175895.fasta.transdecoder.pep
Curcuma_caesia_GCA_037834155.fasta.transdecoder.pep
Hedychium_gardnerianum_GCA_038050645.fasta.transdecoder.pep
Hedychium_flavescens_GCA_041731415.fasta.transdecoder.pep
Boesenbergia_rotunda_GCA_024364735.fasta.transdecoder.pep
Zingiber_zerumbet_GCA_037833815.fasta.transdecoder.pep
Zingiber_officinale_GCF_018446385.fasta.transdecoder.pep
Kaempferia_galanga_GCA_037974795.fasta.transdecoder.pep
Kaempferia_parviflora_GCA_037177295.fasta.transdecoder.pep
Kaempferia_rotunda_GCA_037177415.fasta.transdecoder.pep
Kaempferia_elegans_GCA_037974815.fasta.transdecoder.pep
1.0
0.8
0.6
0.4
0.2
0.0

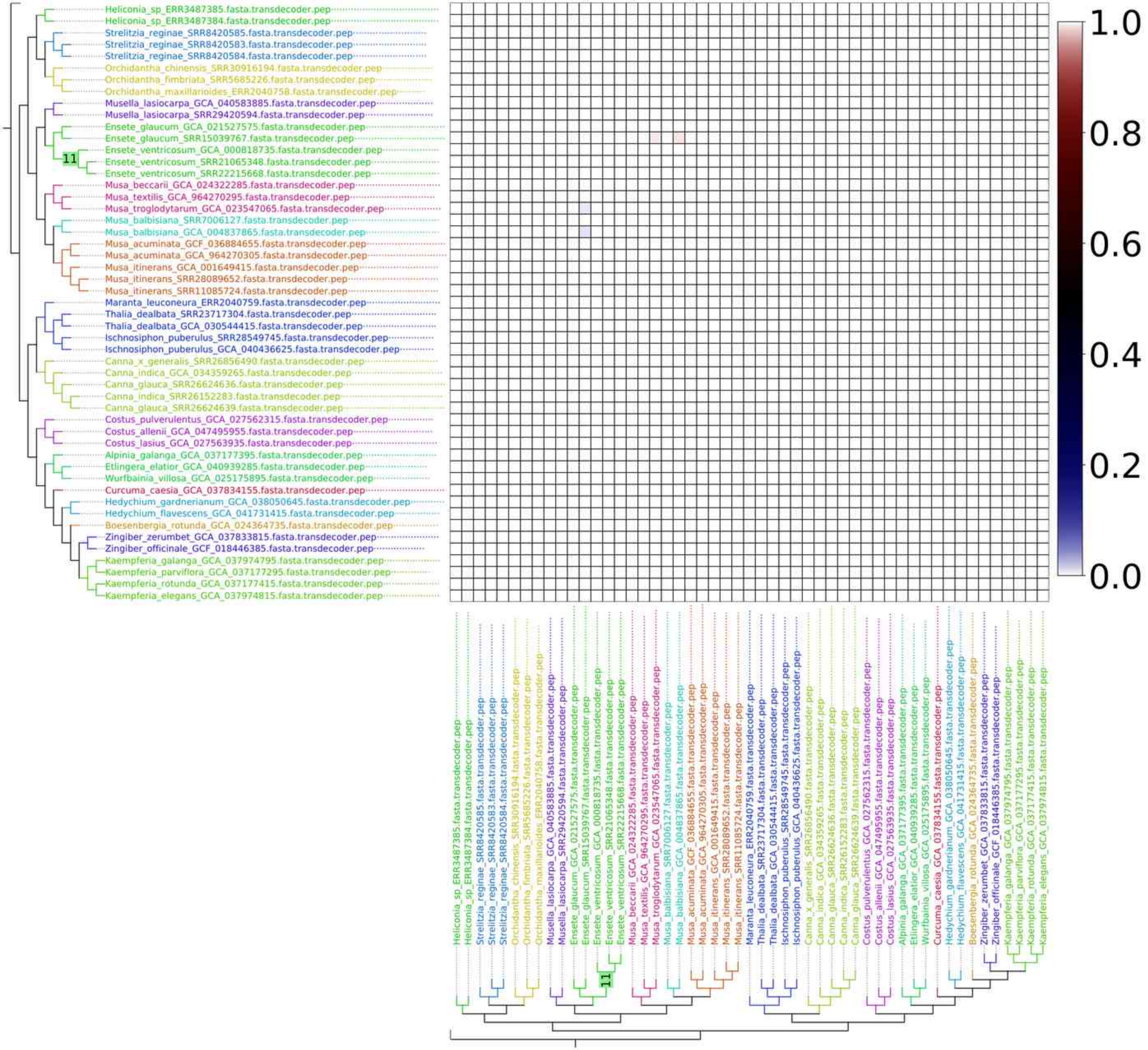
Heliconia_sp_ERR3487385.fasta.transdecoder.pep
Heliconia_sp_ERR3487384.fasta.transdecoder.pep
Strelitzia_reginae_SRR8420585.fasta.transdecoder.pep
Strelitzia_reginae_SRR8420583.fasta.transdecoder.pep
Strelitzia_reginae_SRR8420584.fasta.transdecoder.pep
Orchidantha_chinensis_SRR30916194.fasta.transdecoder.pep
Orchidantha_fimbriata_SRR5685226.fasta.transdecoder.pep
Orchidantha_maxillarioides_ERR2040758.fasta.transdecoder.pep
Musella_lasiocarpa_GCA_040583885.fasta.transdecoder.pep
Musella_lasiocarpa_SRR29420594.fasta.transdecoder.pep
Ensete_glaucum_GCA_021527575.fasta.transdecoder.pep
Ensete_glaucum_SRR15039767.fasta.transdecoder.pep
Ensete_ventricosum_GCA_000818735.fasta.transdecoder.pep
Ensete_ventricosum_SRR21065348.fasta.transdecoder.pep
Ensete_ventricosum_SRR22215668.fasta.transdecoder.pep
Musa_beccarii_GCA_024322285.fasta.transdecoder.pep
Musa_textilis_GCA_964270295.fasta.transdecoder.pep
Musa_troglodytarum_GCA_023547065.fasta.transdecoder.pep
Musa_balbisiana_SRR7006127.fasta.transdecoder.pep
Musa_balbisiana_GCA_004837865.fasta.transdecoder.pep
Musa_acuminata_GCF_036884655.fasta.transdecoder.pep
Musa_acuminata_GCA_964270305.fasta.transdecoder.pep
Musa_itinerans_GCA_001649415.fasta.transdecoder.pep
Musa_itinerans_SRR28089652.fasta.transdecoder.pep
Musa_itinerans_SRR11085724.fasta.transdecoder.pep
Maranta_leuconeura_ERR2040759.fasta.transdecoder.pep
Thalia_dealbata_SRR23717304.fasta.transdecoder.pep
Thalia_dealbata_GCA_030544415.fasta.transdecoder.pep
Ischnosiphon_puberulus_SRR28549745.fasta.transdecoder.pep
Ischnosiphon_puberulus_GCA_040436625.fasta.transdecoder.pep
Canna_x_generalis_SRR26856490.fasta.transdecoder.pep
Canna_indica_GCA_034359265.fasta.transdecoder.pep
Canna_glauca_SRR26624636.fasta.transdecoder.pep
Canna_indica_SRR26152283.fasta.transdecoder.pep
Canna_glauca_SRR26624639.fasta.transdecoder.pep
Costus_pulverulentus_GCA_027562315.fasta.transdecoder.pep
Costus_allenii_GCA_047495955.fasta.transdecoder.pep
Costus_lasius_GCA_027563935.fasta.transdecoder.pep
Alpinia_galanga_GCA_037177395.fasta.transdecoder.pep
Etlingera_elatior_GCA_040939285.fasta.transdecoder.pep
Wurfbainia_villosa_GCA_025175895.fasta.transdecoder.pep
Curcuma_caesia_GCA_037834155.fasta.transdecoder.pep
Hedychium_gardnerianum_GCA_038050645.fasta.transdecoder.pep
Hedychium_flavescens_GCA_041731415.fasta.transdecoder.pep
Boesenbergia_rotunda_GCA_024364735.fasta.transdecoder.pep
Zingiber_zerumbet_GCA_037833815.fasta.transdecoder.pep
Zingiber_officinale_GCF_018446385.fasta.transdecoder.pep
Kaempferia_galanga_GCA_037974795.fasta.transdecoder.pep
Kaempferia_parviflora_GCA_037177295.fasta.transdecoder.pep
Kaempferia_rotunda_GCA_037177415.fasta.transdecoder.pep
Kaempferia_elegans_GCA_037974815.fasta.transdecoder.pep
11
1.0
0.8
0.6
0.4
0.2
0.0

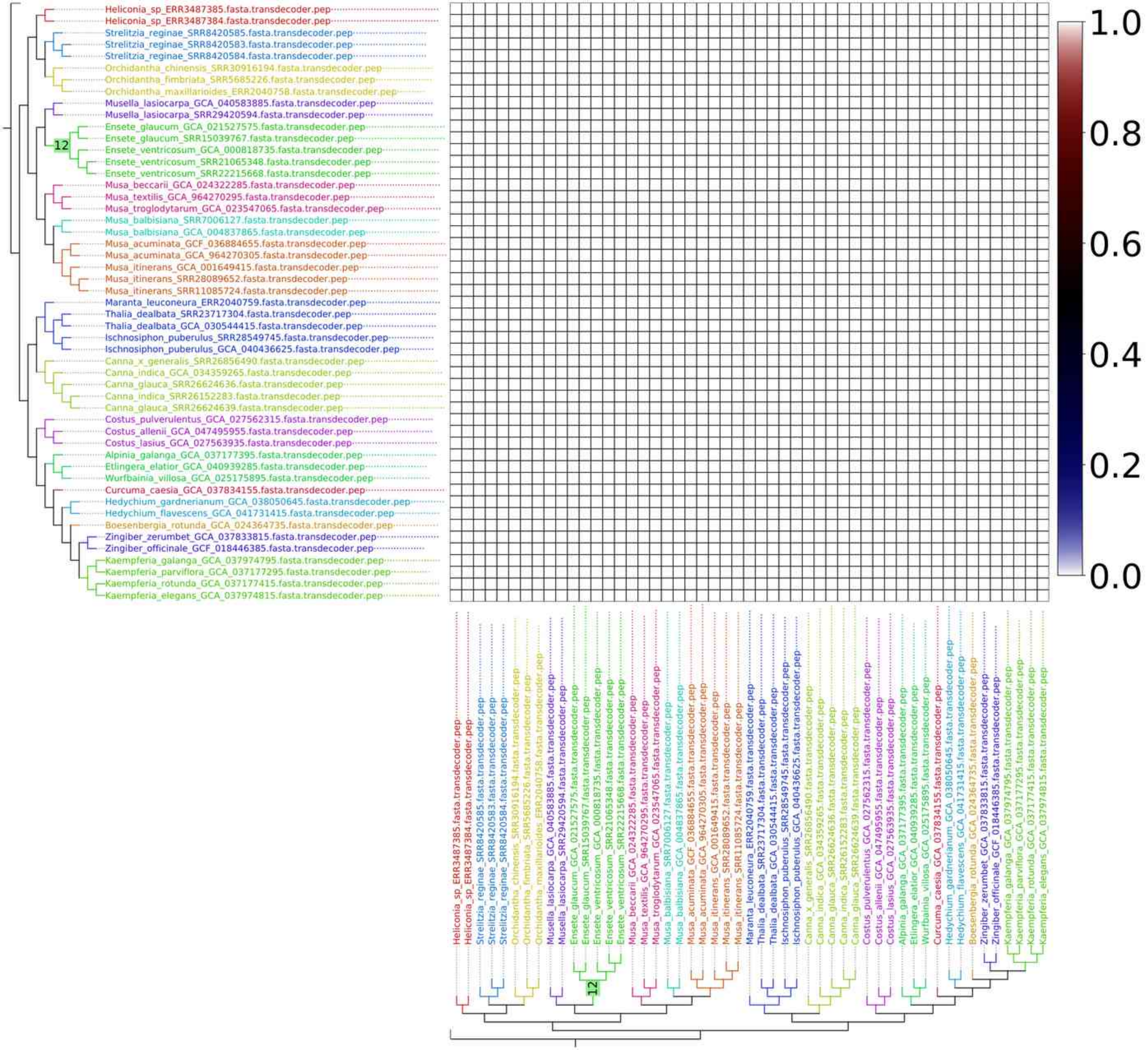
Heliconia_sp_ERR3487385.fasta.transdecoder.pep
Heliconia_sp_ERR3487384.fasta.transdecoder.pep
Strelitzia_reginae_SRR8420585.fasta.transdecoder.pep
Strelitzia_reginae_SRR8420583.fasta.transdecoder.pep
Strelitzia_reginae_SRR8420584.fasta.transdecoder.pep
Orchidantha_chinensis_SRR30916194.fasta.transdecoder.pep
Orchidantha_fimbriata_SRR5685226.fasta.transdecoder.pep
Orchidantha_maxillarioides_ERR2040758.fasta.transdecoder.pep
Musella_lasiocarpa_GCA_040583885.fasta.transdecoder.pep
Musella_lasiocarpa_SRR29420594.fasta.transdecoder.pep
Ensete_glaucum_GCA_021527575.fasta.transdecoder.pep
Ensete_glaucum_SRR15039767.fasta.transdecoder.pep
Ensete_ventricosum_GCA_000818735.fasta.transdecoder.pep
Ensete_ventricosum_SRR21065348.fasta.transdecoder.pep
Ensete_ventricosum_SRR22215668.fasta.transdecoder.pep
Musa_beccarii_GCA_024322285.fasta.transdecoder.pep
Musa_textilis_GCA_964270295.fasta.transdecoder.pep
Musa_troglodytarum_GCA_023547065.fasta.transdecoder.pep
Musa_balbisiana_SRR7006127.fasta.transdecoder.pep
Musa_balbisiana_GCA_004837865.fasta.transdecoder.pep
Musa_acuminata_GCF_036884655.fasta.transdecoder.pep
Musa_acuminata_GCA_964270305.fasta.transdecoder.pep
Musa_itinerans_GCA_001649415.fasta.transdecoder.pep
Musa_itinerans_SRR28089652.fasta.transdecoder.pep
Musa_itinerans_SRR11085724.fasta.transdecoder.pep
Maranta_leuconeura_ERR2040759.fasta.transdecoder.pep
Thalia_dealbata_SRR23717304.fasta.transdecoder.pep
Thalia_dealbata_GCA_030544415.fasta.transdecoder.pep
Ischnosiphon_puberulus_SRR28549745.fasta.transdecoder.pep
Ischnosiphon_puberulus_GCA_040436625.fasta.transdecoder.pep
Canna_x_generalis_SRR26856490.fasta.transdecoder.pep
Canna_indica_GCA_034359265.fasta.transdecoder.pep
Canna_glauca_SRR26624636.fasta.transdecoder.pep
Canna_indica_SRR26152283.fasta.transdecoder.pep
Canna_glauca_SRR26624639.fasta.transdecoder.pep
Costus_pulverulentus_GCA_027562315.fasta.transdecoder.pep
Costus_allenii_GCA_047495955.fasta.transdecoder.pep
Costus_lasius_GCA_027563935.fasta.transdecoder.pep
Alpinia_galanga_GCA_037177395.fasta.transdecoder.pep
Etlingera_elatior_GCA_040939285.fasta.transdecoder.pep
Wurfbainia_villosa_GCA_025175895.fasta.transdecoder.pep
Curcuma_caesia_GCA_037834155.fasta.transdecoder.pep
Hedychium_gardnerianum_GCA_038050645.fasta.transdecoder.pep
Hedychium_flavescens_GCA_041731415.fasta.transdecoder.pep
Boesenbergia_rotunda_GCA_024364735.fasta.transdecoder.pep
Zingiber_zerumbet_GCA_037833815.fasta.transdecoder.pep
Zingiber_officinale_GCF_018446385.fasta.transdecoder.pep
Kaempferia_galanga_GCA_037974795.fasta.transdecoder.pep
Kaempferia_parviflora_GCA_037177295.fasta.transdecoder.pep
Kaempferia_rotunda_GCA_037177415.fasta.transdecoder.pep
Kaempferia_elegans_GCA_037974815.fasta.transdecoder.pep
12
1.0
0.8
0.6
0.4
0.2
0.0

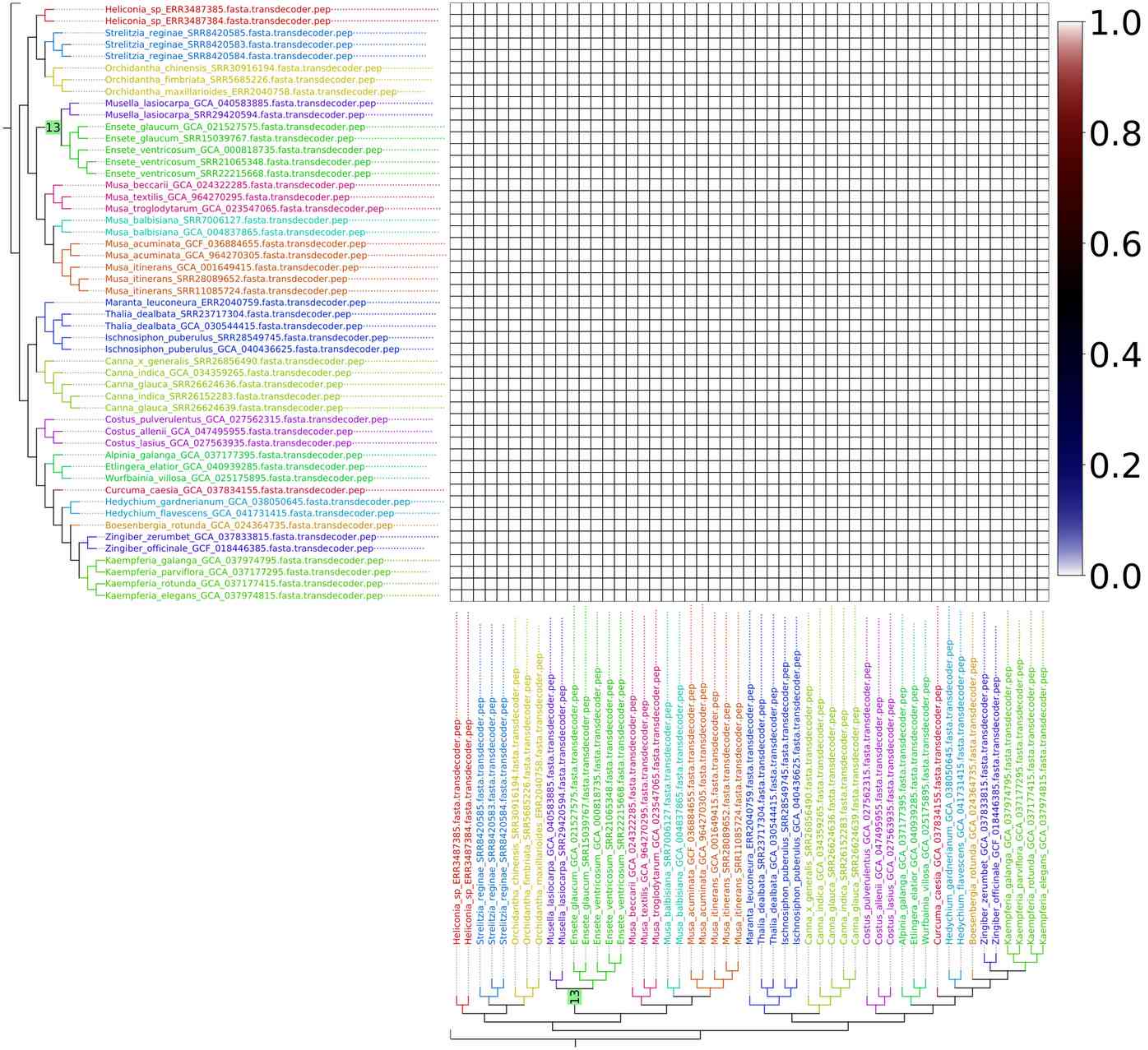
Heliconia_sp_ERR3487385.fasta.transdecoder.pep
Heliconia_sp_ERR3487384.fasta.transdecoder.pep
Strelitzia_reginae_SRR8420585.fasta.transdecoder.pep
Strelitzia_reginae_SRR8420583.fasta.transdecoder.pep
Strelitzia_reginae_SRR8420584.fasta.transdecoder.pep
Orchidantha_chinensis_SRR30916194.fasta.transdecoder.pep
Orchidantha_fimbriata_SRR5685226.fasta.transdecoder.pep
Orchidantha_maxillarioides_ERR2040758.fasta.transdecoder.pep
Musella_lasiocarpa_GCA_040583885.fasta.transdecoder.pep
Musella_lasiocarpa_SRR29420594.fasta.transdecoder.pep
Ensete_glaucum_GCA_021527575.fasta.transdecoder.pep
Ensete_glaucum_SRR15039767.fasta.transdecoder.pep
Ensete_ventricosum_GCA_000818735.fasta.transdecoder.pep
Ensete_ventricosum_SRR21065348.fasta.transdecoder.pep
Ensete_ventricosum_SRR22215668.fasta.transdecoder.pep
Musa_beccarii_GCA_024322285.fasta.transdecoder.pep
Musa_textilis_GCA_964270295.fasta.transdecoder.pep
Musa_troglodytarum_GCA_023547065.fasta.transdecoder.pep
Musa_balbisiana_SRR7006127.fasta.transdecoder.pep
Musa_balbisiana_GCA_004837865.fasta.transdecoder.pep
Musa_acuminata_GCF_036884655.fasta.transdecoder.pep
Musa_acuminata_GCA_964270305.fasta.transdecoder.pep
Musa_itinerans_GCA_001649415.fasta.transdecoder.pep
Musa_itinerans_SRR28089652.fasta.transdecoder.pep
Musa_itinerans_SRR11085724.fasta.transdecoder.pep
Maranta_leuconeura_ERR2040759.fasta.transdecoder.pep
Thalia_dealbata_SRR23717304.fasta.transdecoder.pep
Thalia_dealbata_GCA_030544415.fasta.transdecoder.pep
Ischnosiphon_puberulus_SRR28549745.fasta.transdecoder.pep
Ischnosiphon_puberulus_GCA_040436625.fasta.transdecoder.pep
Canna_x_generalis_SRR26856490.fasta.transdecoder.pep
Canna_indica_GCA_034359265.fasta.transdecoder.pep
Canna_glauca_SRR26624636.fasta.transdecoder.pep
Canna_indica_SRR26152283.fasta.transdecoder.pep
Canna_glauca_SRR26624639.fasta.transdecoder.pep
Costus_pulverulentus_GCA_027562315.fasta.transdecoder.pep
Costus_allenii_GCA_047495955.fasta.transdecoder.pep
Costus_lasius_GCA_027563935.fasta.transdecoder.pep
Alpinia_galanga_GCA_037177395.fasta.transdecoder.pep
Etlingera_elatior_GCA_040939285.fasta.transdecoder.pep
Wurfbainia_villosa_GCA_025175895.fasta.transdecoder.pep
Curcuma_caesia_GCA_037834155.fasta.transdecoder.pep
Hedychium_gardnerianum_GCA_038050645.fasta.transdecoder.pep
Hedychium_flavescens_GCA_041731415.fasta.transdecoder.pep
Boesenbergia_rotunda_GCA_024364735.fasta.transdecoder.pep
Zingiber_zerumbet_GCA_037833815.fasta.transdecoder.pep
Zingiber_officinale_GCF_018446385.fasta.transdecoder.pep
Kaempferia_galanga_GCA_037974795.fasta.transdecoder.pep
Kaempferia_parviflora_GCA_037177295.fasta.transdecoder.pep
Kaempferia_rotunda_GCA_037177415.fasta.transdecoder.pep
Kaempferia_elegans_GCA_037974815.fasta.transdecoder.pep
13
1.0
0.8
0.6
0.4
0.2
0.0

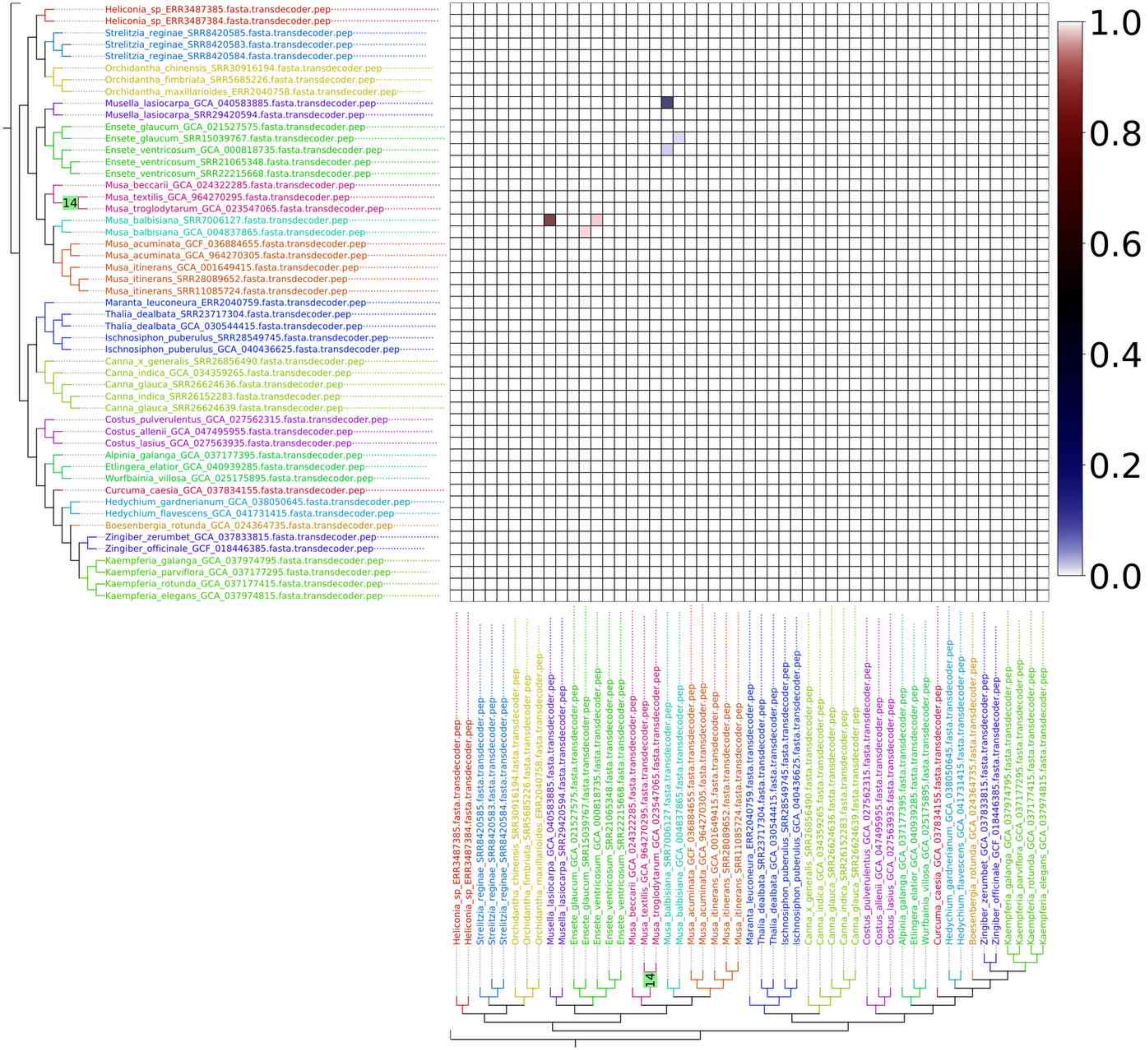
Heliconia_sp_ERR3487385.fasta.transdecoder.pep
Heliconia_sp_ERR3487384.fasta.transdecoder.pep
Strelitzia_reginae_SRR8420585.fasta.transdecoder.pep
Strelitzia_reginae_SRR8420583.fasta.transdecoder.pep
Strelitzia_reginae_SRR8420584.fasta.transdecoder.pep
Orchidantha_chinensis_SRR30916194.fasta.transdecoder.pep
Orchidantha_fimbriata_SRR5685226.fasta.transdecoder.pep
Orchidantha_maxillarioides_ERR2040758.fasta.transdecoder.pep
Musella_lasiocarpa_GCA_040583885.fasta.transdecoder.pep
Musella_lasiocarpa_SRR29420594.fasta.transdecoder.pep
Ensete_glaucum_GCA_021527575.fasta.transdecoder.pep
Ensete_glaucum_SRR15039767.fasta.transdecoder.pep
Ensete_ventricosum_GCA_000818735.fasta.transdecoder.pep
Ensete_ventricosum_SRR21065348.fasta.transdecoder.pep
Ensete_ventricosum_SRR22215668.fasta.transdecoder.pep
Musa_beccarii_GCA_024322285.fasta.transdecoder.pep
Musa_textilis_GCA_964270295.fasta.transdecoder.pep
Musa_troglodytarum_GCA_023547065.fasta.transdecoder.pep
Musa_balbisiana_SRR7006127.fasta.transdecoder.pep
Musa_balbisiana_GCA_004837865.fasta.transdecoder.pep
Musa_acuminata_GCF_036884655.fasta.transdecoder.pep
Musa_acuminata_GCA_964270305.fasta.transdecoder.pep
Musa_itinerans_GCA_001649415.fasta.transdecoder.pep
Musa_itinerans_SRR28089652.fasta.transdecoder.pep
Musa_itinerans_SRR11085724.fasta.transdecoder.pep
Maranta_leuconeura_ERR2040759.fasta.transdecoder.pep
Thalia_dealbata_SRR23717304.fasta.transdecoder.pep
Thalia_dealbata_GCA_030544415.fasta.transdecoder.pep
Ischnosiphon_puberulus_SRR28549745.fasta.transdecoder.pep
Ischnosiphon_puberulus_GCA_040436625.fasta.transdecoder.pep
Canna_x_generalis_SRR26856490.fasta.transdecoder.pep
Canna_indica_GCA_034359265.fasta.transdecoder.pep
Canna_glauca_SRR26624636.fasta.transdecoder.pep
Canna_indica_SRR26152283.fasta.transdecoder.pep
Canna_glauca_SRR26624639.fasta.transdecoder.pep
Costus_pulverulentus_GCA_027562315.fasta.transdecoder.pep
Costus_allenii_GCA_047495955.fasta.transdecoder.pep
Costus_lasius_GCA_027563935.fasta.transdecoder.pep
Alpinia_galanga_GCA_037177395.fasta.transdecoder.pep
Etlingera_elatior_GCA_040939285.fasta.transdecoder.pep
Wurfbainia_villosa_GCA_025175895.fasta.transdecoder.pep
Curcuma_caesia_GCA_037834155.fasta.transdecoder.pep
Hedychium_gardnerianum_GCA_038050645.fasta.transdecoder.pep
Hedychium_flavescens_GCA_041731415.fasta.transdecoder.pep
Boesenbergia_rotunda_GCA_024364735.fasta.transdecoder.pep
Zingiber_zerumbet_GCA_037833815.fasta.transdecoder.pep
Zingiber_officinale_GCF_018446385.fasta.transdecoder.pep
Kaempferia_galanga_GCA_037974795.fasta.transdecoder.pep
Kaempferia_parviflora_GCA_037177295.fasta.transdecoder.pep
Kaempferia_rotunda_GCA_037177415.fasta.transdecoder.pep
Kaempferia_elegans_GCA_037974815.fasta.transdecoder.pep
14
1.0
0.8
0.6
0.4
0.2
0.0

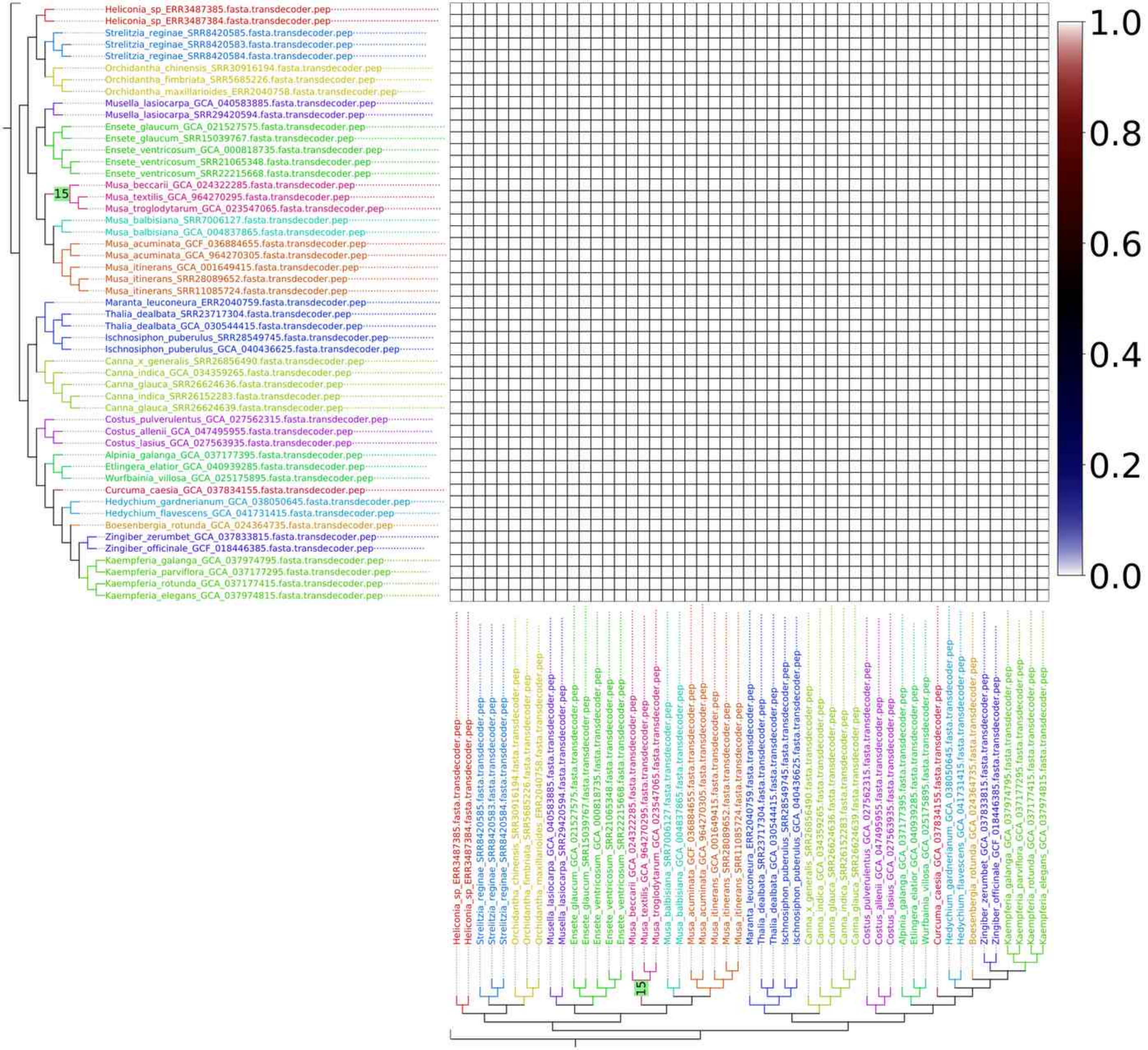
Heliconia_sp_ERR3487385.fasta.transdecoder.pep
Heliconia_sp_ERR3487384.fasta.transdecoder.pep
Strelitzia_reginae_SRR8420585.fasta.transdecoder.pep
Strelitzia_reginae_SRR8420583.fasta.transdecoder.pep
Strelitzia_reginae_SRR8420584.fasta.transdecoder.pep
Orchidantha_chinensis_SRR30916194.fasta.transdecoder.pep
Orchidantha_fimbriata_SRR5685226.fasta.transdecoder.pep
Orchidantha_maxillarioides_ERR2040758.fasta.transdecoder.pep
Musella_lasiocarpa_GCA_040583885.fasta.transdecoder.pep
Musella_lasiocarpa_SRR29420594.fasta.transdecoder.pep
Ensete_glaucum_GCA_021527575.fasta.transdecoder.pep
Ensete_glaucum_SRR15039767.fasta.transdecoder.pep
Ensete_ventricosum_GCA_000818735.fasta.transdecoder.pep
Ensete_ventricosum_SRR21065348.fasta.transdecoder.pep
Ensete_ventricosum_SRR22215668.fasta.transdecoder.pep
15
Musa_beccarii_GCA_024322285.fasta.transdecoder.pep
Musa_textilis_GCA_964270295.fasta.transdecoder.pep
Musa_troglodytarum_GCA_023547065.fasta.transdecoder.pep
Musa_balbisiana_SRR7006127.fasta.transdecoder.pep
Musa_balbisiana_GCA_004837865.fasta.transdecoder.pep
Musa_acuminata_GCF_036884655.fasta.transdecoder.pep
Musa_acuminata_GCA_964270305.fasta.transdecoder.pep
Musa_itinerans_GCA_001649415.fasta.transdecoder.pep
Musa_itinerans_SRR28089652.fasta.transdecoder.pep
Musa_itinerans_SRR11085724.fasta.transdecoder.pep
Maranta_leuconeura_ERR2040759.fasta.transdecoder.pep
Thalia_dealbata_SRR23717304.fasta.transdecoder.pep
Thalia_dealbata_GCA_030544415.fasta.transdecoder.pep
Ischnosiphon_puberulus_SRR28549745.fasta.transdecoder.pep
Ischnosiphon_puberulus_GCA_040436625.fasta.transdecoder.pep
Canna_x_generalis_SRR26856490.fasta.transdecoder.pep
Canna_indica_GCA_034359265.fasta.transdecoder.pep
Canna_glauca_SRR26624636.fasta.transdecoder.pep
Canna_indica_SRR26152283.fasta.transdecoder.pep
Canna_glauca_SRR26624639.fasta.transdecoder.pep
Costus_pulverulentus_GCA_027562315.fasta.transdecoder.pep
Costus_allenii_GCA_047495955.fasta.transdecoder.pep
Costus_lasius_GCA_027563935.fasta.transdecoder.pep
Alpinia_galanga_GCA_037177395.fasta.transdecoder.pep
Etlingera_elatior_GCA_040939285.fasta.transdecoder.pep
Wurfbainia_villosa_GCA_025175895.fasta.transdecoder.pep
Curcuma_caesia_GCA_037834155.fasta.transdecoder.pep
Hedychium_gardnerianum_GCA_038050645.fasta.transdecoder.pep
Hedychium_flavescens_GCA_041731415.fasta.transdecoder.pep
Boesenbergia_rotunda_GCA_024364735.fasta.transdecoder.pep
Zingiber_zerumbet_GCA_037833815.fasta.transdecoder.pep
Zingiber_officinale_GCF_018446385.fasta.transdecoder.pep
Kaempferia_galanga_GCA_037974795.fasta.transdecoder.pep
Kaempferia_parviflora_GCA_037177295.fasta.transdecoder.pep
Kaempferia_rotunda_GCA_037177415.fasta.transdecoder.pep
Kaempferia_elegans_GCA_037974815.fasta.transdecoder.pep
1.0
0.8
0.6
0.4
0.2
0.0

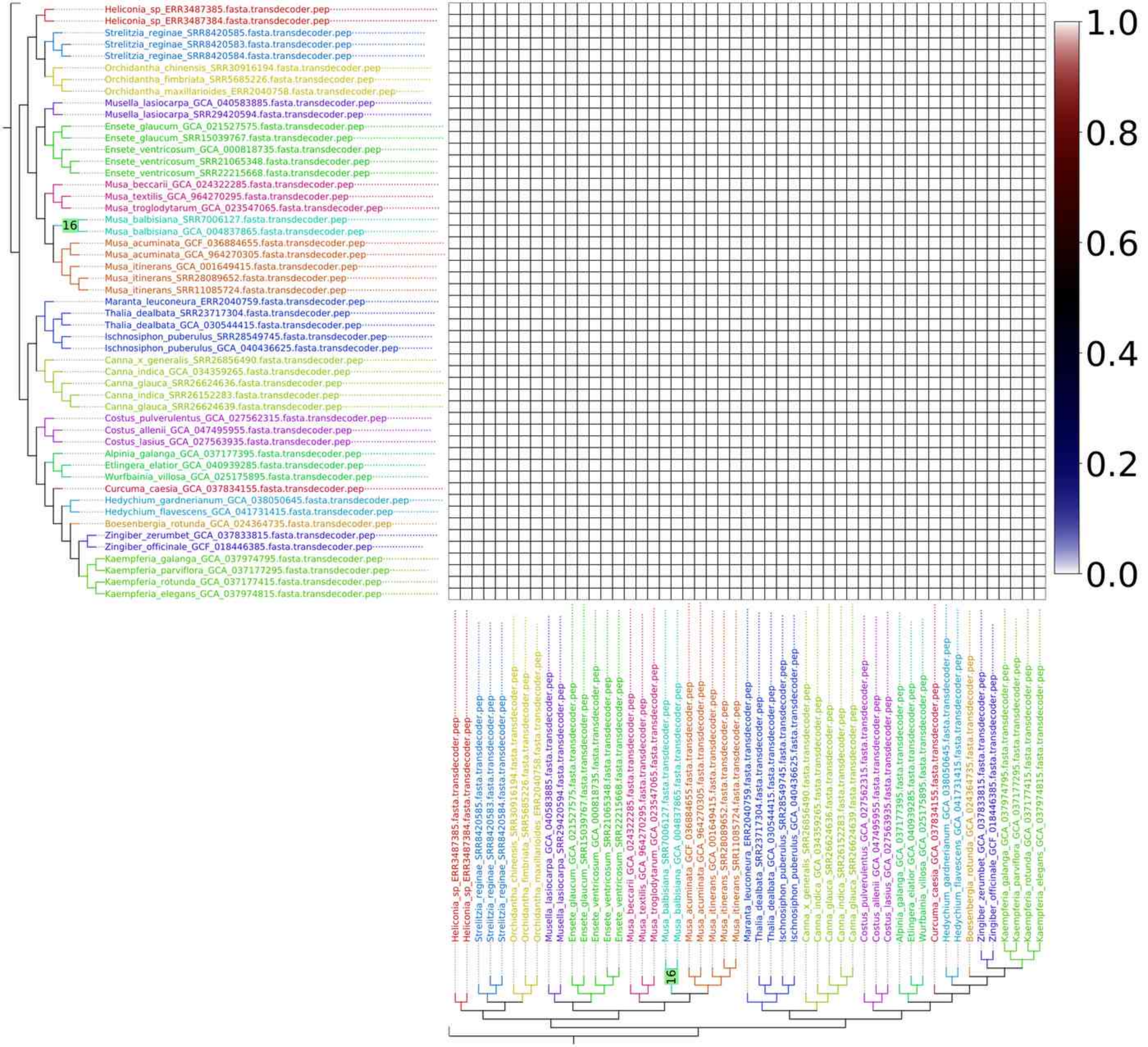
Heliconia_sp_ERR3487385.fasta.transdecoder.pep
Heliconia_sp_ERR3487384.fasta.transdecoder.pep
Strelitzia_reginae_SRR8420585.fasta.transdecoder.pep
Strelitzia_reginae_SRR8420583.fasta.transdecoder.pep
Strelitzia_reginae_SRR8420584.fasta.transdecoder.pep
Orchidantha_chinensis_SRR30916194.fasta.transdecoder.pep
Orchidantha_fimbriata_SRR5685226.fasta.transdecoder.pep
Orchidantha_maxillarioides_ERR2040758.fasta.transdecoder.pep
Musella_lasiocarpa_GCA_040583885.fasta.transdecoder.pep
Musella_lasiocarpa_SRR29420594.fasta.transdecoder.pep
Ensete_glaucum_GCA_021527575.fasta.transdecoder.pep
Ensete_glaucum_SRR15039767.fasta.transdecoder.pep
Ensete_ventricosum_GCA_000818735.fasta.transdecoder.pep
Ensete_ventricosum_SRR21065348.fasta.transdecoder.pep
Ensete_ventricosum_SRR22215668.fasta.transdecoder.pep
Musa_beccarii_GCA_024322285.fasta.transdecoder.pep
Musa_textilis_GCA_964270295.fasta.transdecoder.pep
Musa_troglodytarum_GCA_023547065.fasta.transdecoder.pep
Musa_balbisiana_SRR7006127.fasta.transdecoder.pep
16
Musa_balbisiana_GCA_004837865.fasta.transdecoder.pep
Musa_acuminata_GCF_036884655.fasta.transdecoder.pep
Musa_acuminata_GCA_964270305.fasta.transdecoder.pep
Musa_itinerans_GCA_001649415.fasta.transdecoder.pep
Musa_itinerans_SRR28089652.fasta.transdecoder.pep
Musa_itinerans_SRR11085724.fasta.transdecoder.pep
Maranta_leuconeura_ERR2040759.fasta.transdecoder.pep
Thalia_dealbata_SRR23717304.fasta.transdecoder.pep
Thalia_dealbata_GCA_030544415.fasta.transdecoder.pep
Ischnosiphon_puberulus_SRR28549745.fasta.transdecoder.pep
Ischnosiphon_puberulus_GCA_040436625.fasta.transdecoder.pep
Canna_x_generalis_SRR26856490.fasta.transdecoder.pep
Canna_indica_GCA_034359265.fasta.transdecoder.pep
Canna_glauca_SRR26624636.fasta.transdecoder.pep
Canna_indica_SRR26152283.fasta.transdecoder.pep
Canna_glauca_SRR26624639.fasta.transdecoder.pep
Costus_pulverulentus_GCA_027562315.fasta.transdecoder.pep
Costus_allenii_GCA_047495955.fasta.transdecoder.pep
Costus_lasius_GCA_027563935.fasta.transdecoder.pep
Alpinia_galanga_GCA_037177395.fasta.transdecoder.pep
Etlingera_elatior_GCA_040939285.fasta.transdecoder.pep
Wurfbainia_villosa_GCA_025175895.fasta.transdecoder.pep
Curcuma_caesia_GCA_037834155.fasta.transdecoder.pep
Hedychium_gardnerianum_GCA_038050645.fasta.transdecoder.pep
Hedychium_flavescens_GCA_041731415.fasta.transdecoder.pep
Boesenbergia_rotunda_GCA_024364735.fasta.transdecoder.pep
Zingiber_zerumbet_GCA_037833815.fasta.transdecoder.pep
Zingiber_officinale_GCF_018446385.fasta.transdecoder.pep
Kaempferia_galanga_GCA_037974795.fasta.transdecoder.pep
Kaempferia_parviflora_GCA_037177295.fasta.transdecoder.pep
Kaempferia_rotunda_GCA_037177415.fasta.transdecoder.pep
Kaempferia_elegans_GCA_037974815.fasta.transdecoder.pep
1.0
0.8
0.6
0.4
0.2
0.0

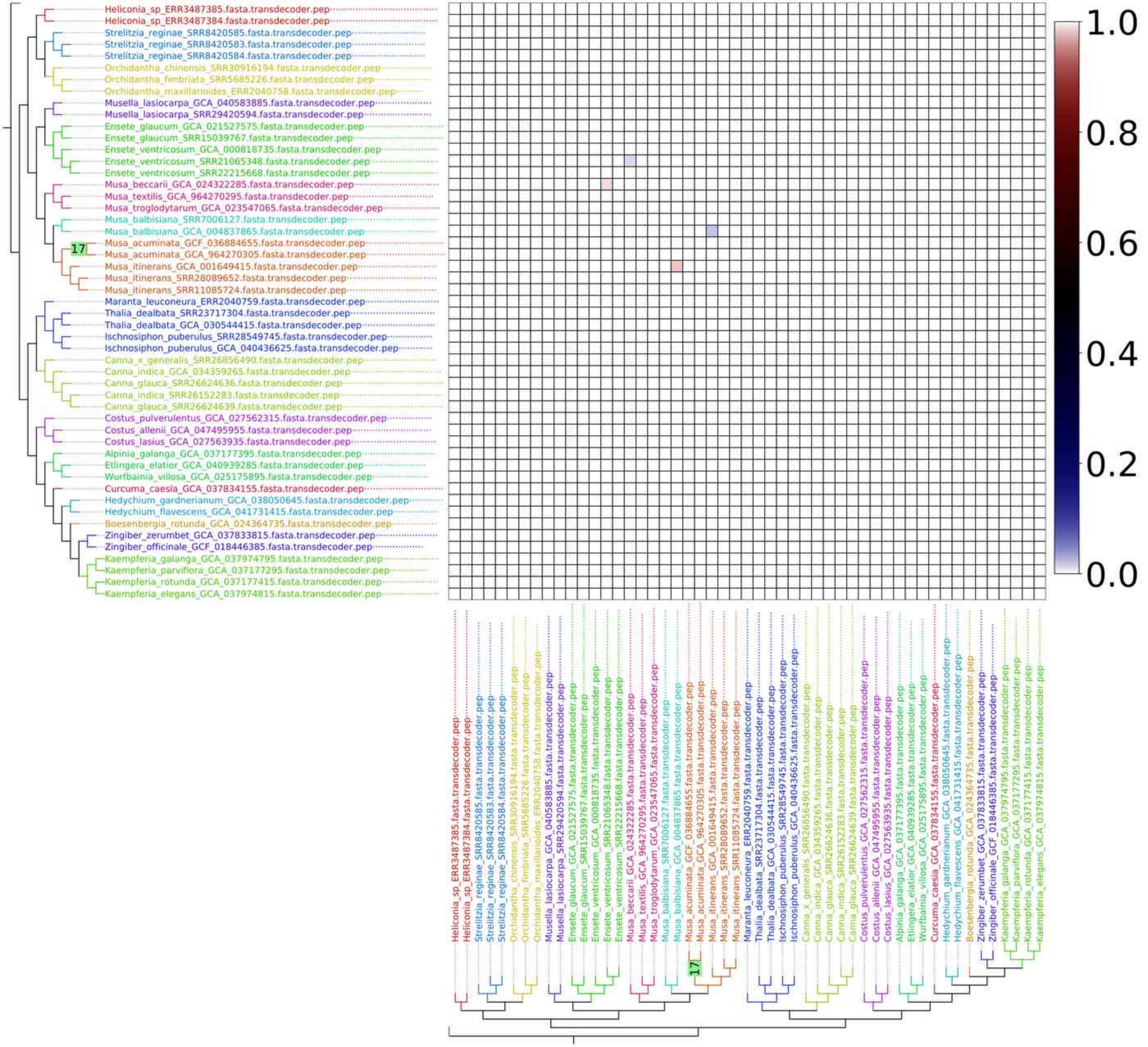
Heliconia_sp_ERR3487385.fasta.transdecoder.pep
Heliconia_sp_ERR3487384.fasta.transdecoder.pep
Strelitzia_reginae_SRR8420585.fasta.transdecoder.pep
Strelitzia_reginae_SRR8420583.fasta.transdecoder.pep
Strelitzia_reginae_SRR8420584.fasta.transdecoder.pep
Orchidantha_chinensis_SRR30916194.fasta.transdecoder.pep
Orchidantha_fimbriata_SRR5685226.fasta.transdecoder.pep
Orchidantha_maxillarioides_ERR2040758.fasta.transdecoder.pep
Musella_lasiocarpa_GCA_040583885.fasta.transdecoder.pep
Musella_lasiocarpa_SRR29420594.fasta.transdecoder.pep
Ensete_glaucum_GCA_021527575.fasta.transdecoder.pep
Ensete_glaucum_SRR15039767.fasta.transdecoder.pep
Ensete_ventricosum_GCA_000818735.fasta.transdecoder.pep
Ensete_ventricosum_SRR21065348.fasta.transdecoder.pep
Ensete_ventricosum_SRR22215668.fasta.transdecoder.pep
Musa_beccarii_GCA_024322285.fasta.transdecoder.pep
Musa_textilis_GCA_964270295.fasta.transdecoder.pep
Musa_troglodytarum_GCA_023547065.fasta.transdecoder.pep
Musa_balbisiana_SRR7006127.fasta.transdecoder.pep
Musa_balbisiana_GCA_004837865.fasta.transdecoder.pep
17
Musa_acuminata_GCF_036884655.fasta.transdecoder.pep
Musa_acuminata_GCA_964270305.fasta.transdecoder.pep
Musa_itinerans_GCA_001649415.fasta.transdecoder.pep
Musa_itinerans_SRR28089652.fasta.transdecoder.pep
Musa_itinerans_SRR11085724.fasta.transdecoder.pep
Maranta_leuconeura_ERR2040759.fasta.transdecoder.pep
Thalia_dealbata_SRR23717304.fasta.transdecoder.pep
Thalia_dealbata_GCA_030544415.fasta.transdecoder.pep
Ischnosiphon_puberulus_SRR28549745.fasta.transdecoder.pep
Ischnosiphon_puberulus_GCA_040436625.fasta.transdecoder.pep
Canna_x_generalis_SRR26856490.fasta.transdecoder.pep
Canna_indica_GCA_034359265.fasta.transdecoder.pep
Canna_glauca_SRR26624636.fasta.transdecoder.pep
Canna_indica_SRR26152283.fasta.transdecoder.pep
Canna_glauca_SRR26624639.fasta.transdecoder.pep
Costus_pulverulentus_GCA_027562315.fasta.transdecoder.pep
Costus_allenii_GCA_047495955.fasta.transdecoder.pep
Costus_lasius_GCA_027563935.fasta.transdecoder.pep
Alpinia_galanga_GCA_037177395.fasta.transdecoder.pep
Etlingera_elatior_GCA_040939285.fasta.transdecoder.pep
Wurfbainia_villosa_GCA_025175895.fasta.transdecoder.pep
Curcuma_caesia_GCA_037834155.fasta.transdecoder.pep
Hedychium_gardnerianum_GCA_038050645.fasta.transdecoder.pep
Hedychium_flavescens_GCA_041731415.fasta.transdecoder.pep
Boesenbergia_rotunda_GCA_024364735.fasta.transdecoder.pep
Zingiber_zerumbet_GCA_037833815.fasta.transdecoder.pep
Zingiber_officinale_GCF_018446385.fasta.transdecoder.pep
Kaempferia_galanga_GCA_037974795.fasta.transdecoder.pep
Kaempferia_parviflora_GCA_037177295.fasta.transdecoder.pep
Kaempferia_rotunda_GCA_037177415.fasta.transdecoder.pep
Kaempferia_elegans_GCA_037974815.fasta.transdecoder.pep
1.0
0.8
0.6
0.4
0.2
0.0

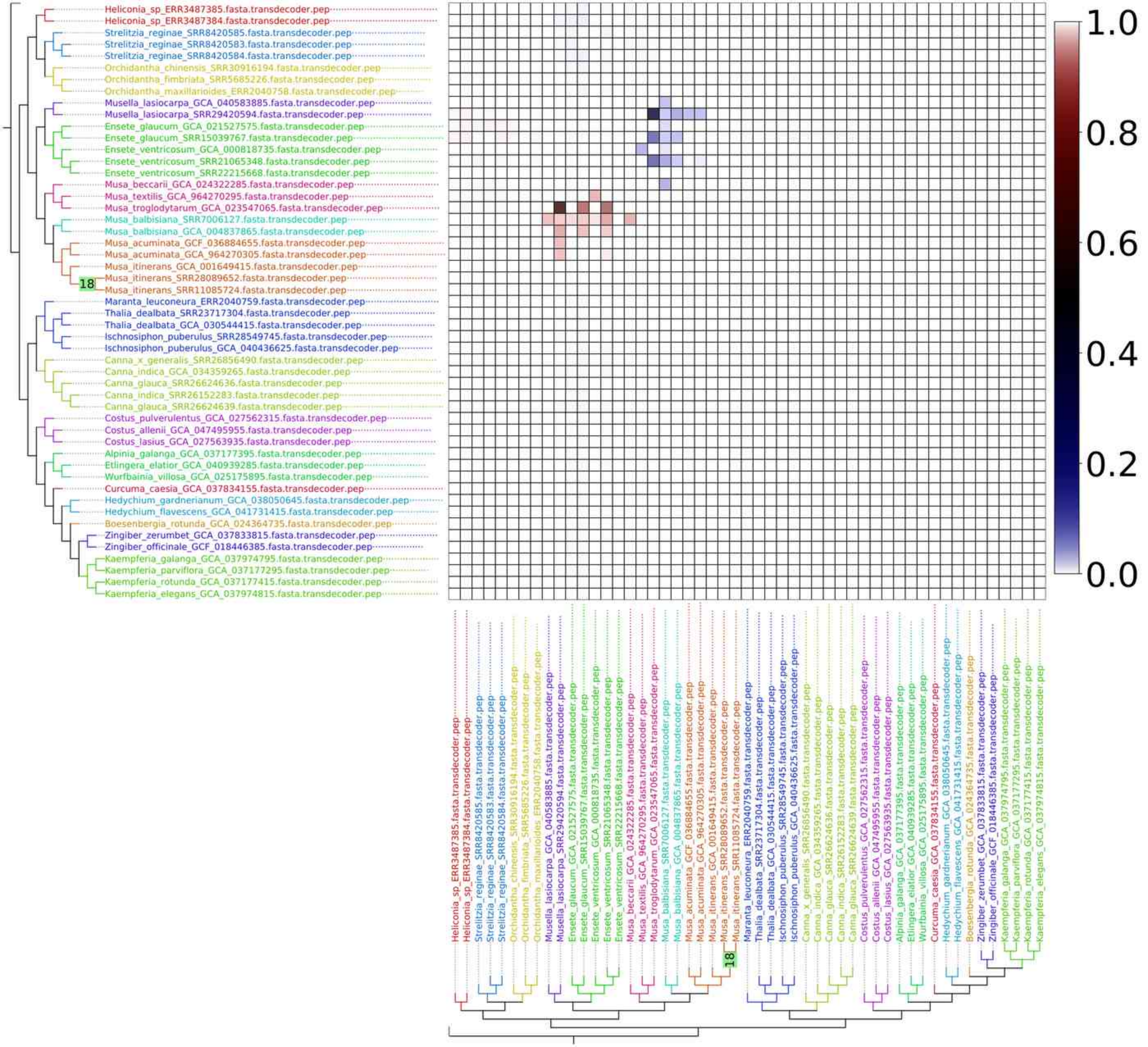
Heliconia_sp_ERR3487385.fasta.transdecoder.pep
Heliconia_sp_ERR3487384.fasta.transdecoder.pep
Strelitzia_reginae_SRR8420585.fasta.transdecoder.pep
Strelitzia_reginae_SRR8420583.fasta.transdecoder.pep
Strelitzia_reginae_SRR8420584.fasta.transdecoder.pep
Orchidantha_chinensis_SRR30916194.fasta.transdecoder.pep
Orchidantha_fimbriata_SRR5685226.fasta.transdecoder.pep
Orchidantha_maxillarioides_ERR2040758.fasta.transdecoder.pep
Musella_lasiocarpa_GCA_040583885.fasta.transdecoder.pep
Musella_lasiocarpa_SRR29420594.fasta.transdecoder.pep
Ensete_glaucum_GCA_021527575.fasta.transdecoder.pep
Ensete_glaucum_SRR15039767.fasta.transdecoder.pep
Ensete_ventricosum_GCA_000818735.fasta.transdecoder.pep
Ensete_ventricosum_SRR21065348.fasta.transdecoder.pep
Ensete_ventricosum_SRR22215668.fasta.transdecoder.pep
Musa_beccarii_GCA_024322285.fasta.transdecoder.pep
Musa_textilis_GCA_964270295.fasta.transdecoder.pep
Musa_troglodytarum_GCA_023547065.fasta.transdecoder.pep
Musa_balbisiana_SRR7006127.fasta.transdecoder.pep
Musa_balbisiana_GCA_004837865.fasta.transdecoder.pep
Musa_acuminata_GCF_036884655.fasta.transdecoder.pep
Musa_acuminata_GCA_964270305.fasta.transdecoder.pep
Musa_itinerans_GCA_001649415.fasta.transdecoder.pep
18
Musa_itinerans_SRR28089652.fasta.transdecoder.pep
Musa_itinerans_SRR11085724.fasta.transdecoder.pep
Maranta_leuconeura_ERR2040759.fasta.transdecoder.pep
Thalia_dealbata_SRR23717304.fasta.transdecoder.pep
Thalia_dealbata_GCA_030544415.fasta.transdecoder.pep
Ischnosiphon_puberulus_SRR28549745.fasta.transdecoder.pep
Ischnosiphon_puberulus_GCA_040436625.fasta.transdecoder.pep
Canna_x_generalis_SRR26856490.fasta.transdecoder.pep
Canna_indica_GCA_034359265.fasta.transdecoder.pep
Canna_glauca_SRR26624636.fasta.transdecoder.pep
Canna_indica_SRR26152283.fasta.transdecoder.pep
Canna_glauca_SRR26624639.fasta.transdecoder.pep
Costus_pulverulentus_GCA_027562315.fasta.transdecoder.pep
Costus_allenii_GCA_047495955.fasta.transdecoder.pep
Costus_lasius_GCA_027563935.fasta.transdecoder.pep
Alpinia_galanga_GCA_037177395.fasta.transdecoder.pep
Etlingera_elatior_GCA_040939285.fasta.transdecoder.pep
Wurfbainia_villosa_GCA_025175895.fasta.transdecoder.pep
Curcuma_caesia_GCA_037834155.fasta.transdecoder.pep
Hedychium_gardnerianum_GCA_038050645.fasta.transdecoder.pep
Hedychium_flavescens_GCA_041731415.fasta.transdecoder.pep
Boesenbergia_rotunda_GCA_024364735.fasta.transdecoder.pep
Zingiber_zerumbet_GCA_037833815.fasta.transdecoder.pep
Zingiber_officinale_GCF_018446385.fasta.transdecoder.pep
Kaempferia_galanga_GCA_037974795.fasta.transdecoder.pep
Kaempferia_parviflora_GCA_037177295.fasta.transdecoder.pep
Kaempferia_rotunda_GCA_037177415.fasta.transdecoder.pep
Kaempferia_elegans_GCA_037974815.fasta.transdecoder.pep
1.0
0.8
0.6
0.4
0.2
0.0

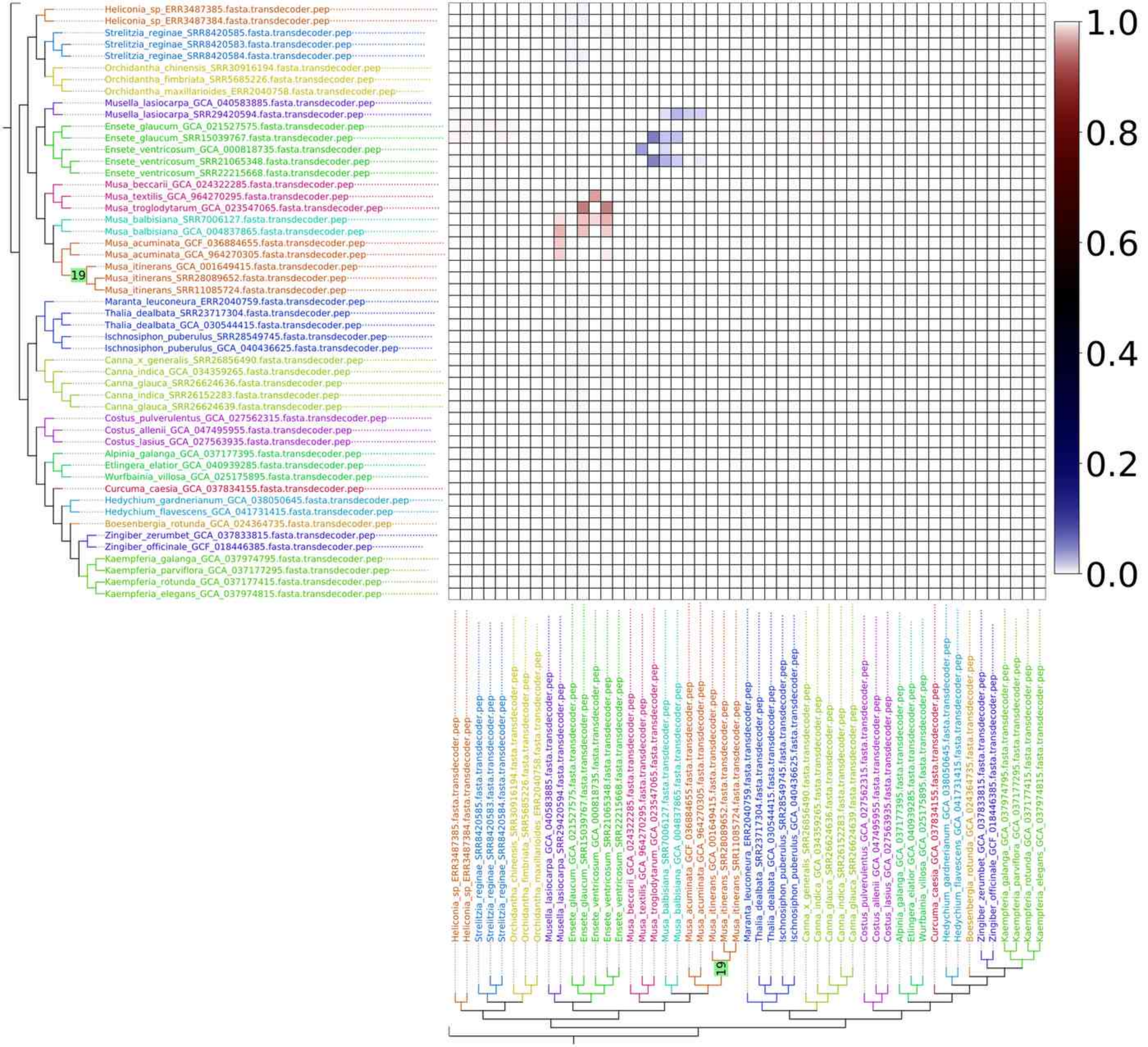
Heliconia_sp_ERR3487385.fasta.transdecoder.pep
Heliconia_sp_ERR3487384.fasta.transdecoder.pep
Strelitzia_reginae_SRR8420585.fasta.transdecoder.pep
Strelitzia_reginae_SRR8420583.fasta.transdecoder.pep
Strelitzia_reginae_SRR8420584.fasta.transdecoder.pep
Orchidantha_chinensis_SRR30916194.fasta.transdecoder.pep
Orchidantha_fimbriata_SRR5685226.fasta.transdecoder.pep
Orchidantha_maxillarioides_ERR2040758.fasta.transdecoder.pep
Musella_lasiocarpa_GCA_040583885.fasta.transdecoder.pep
Musella_lasiocarpa_SRR29420594.fasta.transdecoder.pep
Ensete_glaucum_GCA_021527575.fasta.transdecoder.pep
Ensete_glaucum_SRR15039767.fasta.transdecoder.pep
Ensete_ventricosum_GCA_000818735.fasta.transdecoder.pep
Ensete_ventricosum_SRR21065348.fasta.transdecoder.pep
Ensete_ventricosum_SRR22215668.fasta.transdecoder.pep
Musa_beccarii_GCA_024322285.fasta.transdecoder.pep
Musa_textilis_GCA_964270295.fasta.transdecoder.pep
Musa_troglodytarum_GCA_023547065.fasta.transdecoder.pep
Musa_balbisiana_SRR7006127.fasta.transdecoder.pep
Musa_balbisiana_GCA_004837865.fasta.transdecoder.pep
Musa_acuminata_GCF_036884655.fasta.transdecoder.pep
Musa_acuminata_GCA_964270305.fasta.transdecoder.pep
Musa_itinerans_GCA_001649415.fasta.transdecoder.pep
19
Musa_itinerans_SRR28089652.fasta.transdecoder.pep
Musa_itinerans_SRR11085724.fasta.transdecoder.pep
Maranta_leuconeura_ERR2040759.fasta.transdecoder.pep
Thalia_dealbata_SRR23717304.fasta.transdecoder.pep
Thalia_dealbata_GCA_030544415.fasta.transdecoder.pep
Ischnosiphon_puberulus_SRR28549745.fasta.transdecoder.pep
Ischnosiphon_puberulus_GCA_040436625.fasta.transdecoder.pep
Canna_x_generalis_SRR26856490.fasta.transdecoder.pep
Canna_indica_GCA_034359265.fasta.transdecoder.pep
Canna_glauca_SRR26624636.fasta.transdecoder.pep
Canna_indica_SRR26152283.fasta.transdecoder.pep
Canna_glauca_SRR26624639.fasta.transdecoder.pep
Costus_pulverulentus_GCA_027562315.fasta.transdecoder.pep
Costus_allenii_GCA_047495955.fasta.transdecoder.pep
Costus_lasius_GCA_027563935.fasta.transdecoder.pep
Alpinia_galanga_GCA_037177395.fasta.transdecoder.pep
Etlingera_elatior_GCA_040939285.fasta.transdecoder.pep
Wurfbainia_villosa_GCA_025175895.fasta.transdecoder.pep
Curcuma_caesia_GCA_037834155.fasta.transdecoder.pep
Hedychium_gardnerianum_GCA_038050645.fasta.transdecoder.pep
Hedychium_flavescens_GCA_041731415.fasta.transdecoder.pep
Boesenbergia_rotunda_GCA_024364735.fasta.transdecoder.pep
Zingiber_zerumbet_GCA_037833815.fasta.transdecoder.pep
Zingiber_officinale_GCF_018446385.fasta.transdecoder.pep
Kaempferia_galanga_GCA_037974795.fasta.transdecoder.pep
Kaempferia_parviflora_GCA_037177295.fasta.transdecoder.pep
Kaempferia_rotunda_GCA_037177415.fasta.transdecoder.pep
Kaempferia_elegans_GCA_037974815.fasta.transdecoder.pep
1.0
0.8
0.6
0.4
0.2
0.0

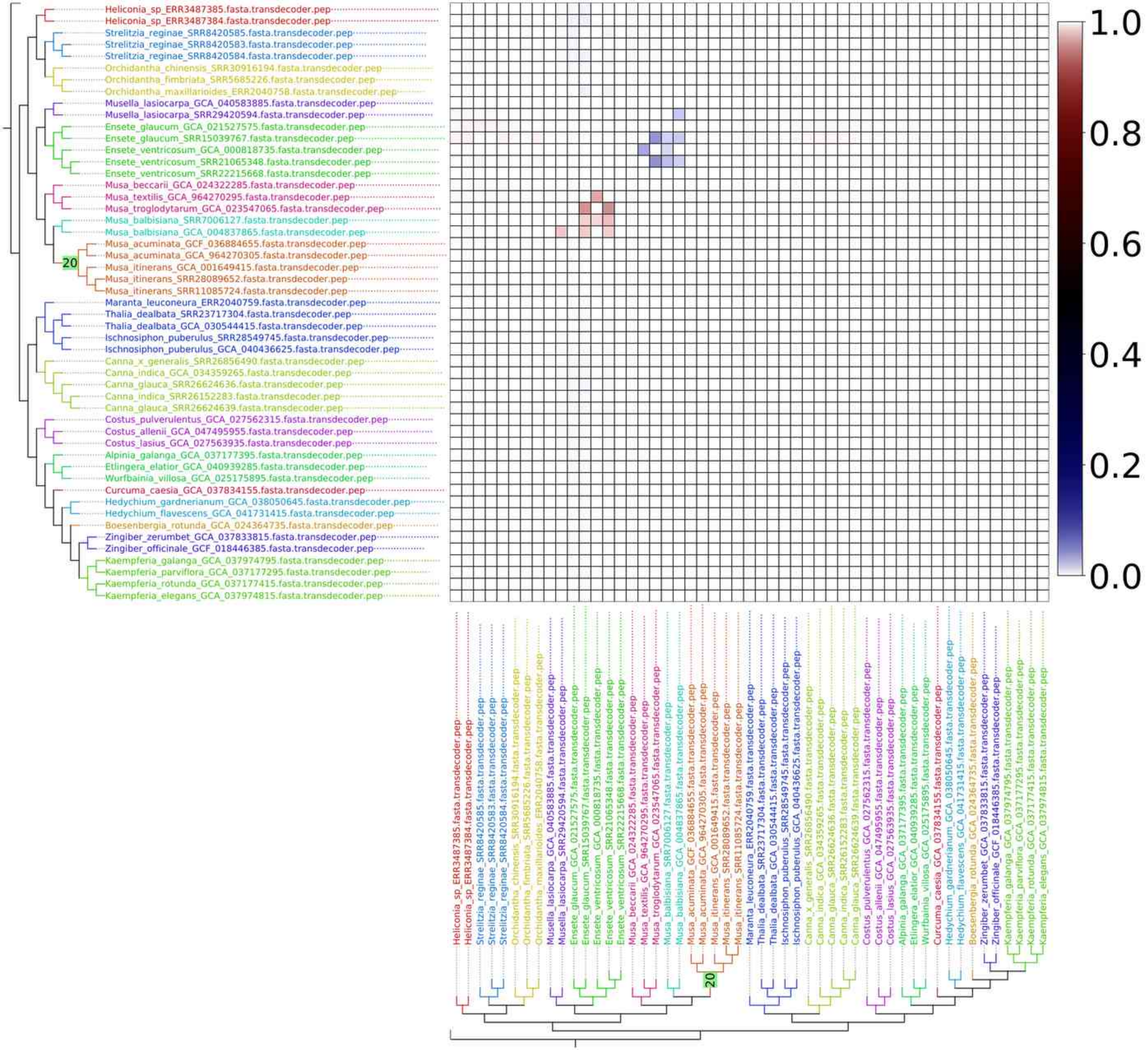

Heliconia_sp_ERR3487385.fasta.transdecoder.pep
Heliconia_sp_ERR3487384.fasta.transdecoder.pep
Strelitzia_reginae_SRR8420585.fasta.transdecoder.pep
Strelitzia_reginae_SRR8420583.fasta.transdecoder.pep
Strelitzia_reginae_SRR8420584.fasta.transdecoder.pep
Orchidantha_chinensis_SRR30916194.fasta.transdecoder.pep
Orchidantha_fimbriata_SRR5685226.fasta.transdecoder.pep
Orchidantha_maxillarioides_ERR2040758.fasta.transdecoder.pep
Musella_lasiocarpa_GCA_040583885.fasta.transdecoder.pep
Musella_lasiocarpa_SRR29420594.fasta.transdecoder.pep
Ensete_glaucum_GCA_021527575.fasta.transdecoder.pep
Ensete_glaucum_SRR15039767.fasta.transdecoder.pep
Ensete_ventricosum_GCA_000818735.fasta.transdecoder.pep
Ensete_ventricosum_SRR21065348.fasta.transdecoder.pep
Ensete_ventricosum_SRR22215668.fasta.transdecoder.pep
Musa_beccarii_GCA_024322285.fasta.transdecoder.pep
Musa_textilis_GCA_964270295.fasta.transdecoder.pep
Musa_troglodytarum_GCA_023547065.fasta.transdecoder.pep
Musa_balbisiana_SRR7006127.fasta.transdecoder.pep
Musa_balbisiana_GCA_004837865.fasta.transdecoder.pep
Musa_acuminata_GCF_036884655.fasta.transdecoder.pep
Musa_acuminata_GCA_964270305.fasta.transdecoder.pep
Musa_itinerans_GCA_001649415.fasta.transdecoder.pep
Musa_itinerans_SRR28089652.fasta.transdecoder.pep
Musa_itinerans_SRR11085724.fasta.transdecoder.pep
Maranta_leuconeura_ERR2040759.fasta.transdecoder.pep
Thalia_dealbata_SRR23717304.fasta.transdecoder.pep
Thalia_dealbata_GCA_030544415.fasta.transdecoder.pep
Ischnosiphon_puberulus_SRR28549745.fasta.transdecoder.pep
Ischnosiphon_puberulus_GCA_040436625.fasta.transdecoder.pep
Canna_x_generalis_SRR26856490.fasta.transdecoder.pep
Canna_indica_GCA_034359265.fasta.transdecoder.pep
Canna_glauca_SRR26624636.fasta.transdecoder.pep
Canna_indica_SRR26152283.fasta.transdecoder.pep
Canna_glauca_SRR26624639.fasta.transdecoder.pep
Costus_pulverulentus_GCA_027562315.fasta.transdecoder.pep
Costus_allenii_GCA_047495955.fasta.transdecoder.pep
Costus_lasius_GCA_027563935.fasta.transdecoder.pep
Alpinia_galanga_GCA_037177395.fasta.transdecoder.pep
Etlingera_elatior_GCA_040939285.fasta.transdecoder.pep
Wurfbainia_villosa_GCA_025175895.fasta.transdecoder.pep
Curcuma_caesia_GCA_037834155.fasta.transdecoder.pep
Hedychium_gardnerianum_GCA_038050645.fasta.transdecoder.pep
Hedychium_flavescens_GCA_041731415.fasta.transdecoder.pep
Boesenbergia_rotunda_GCA_024364735.fasta.transdecoder.pep
Zingiber_zerumbet_GCA_037833815.fasta.transdecoder.pep
Zingiber_officinale_GCF_018446385.fasta.transdecoder.pep
Kaempferia_galanga_GCA_037974795.fasta.transdecoder.pep
Kaempferia_parviflora_GCA_037177295.fasta.transdecoder.pep
Kaempferia_rotunda_GCA_037177415.fasta.transdecoder.pep
Kaempferia_elegans_GCA_037974815.fasta.transdecoder.pep
20
1.0
0.8
0.6
0.4
0.2
0.0

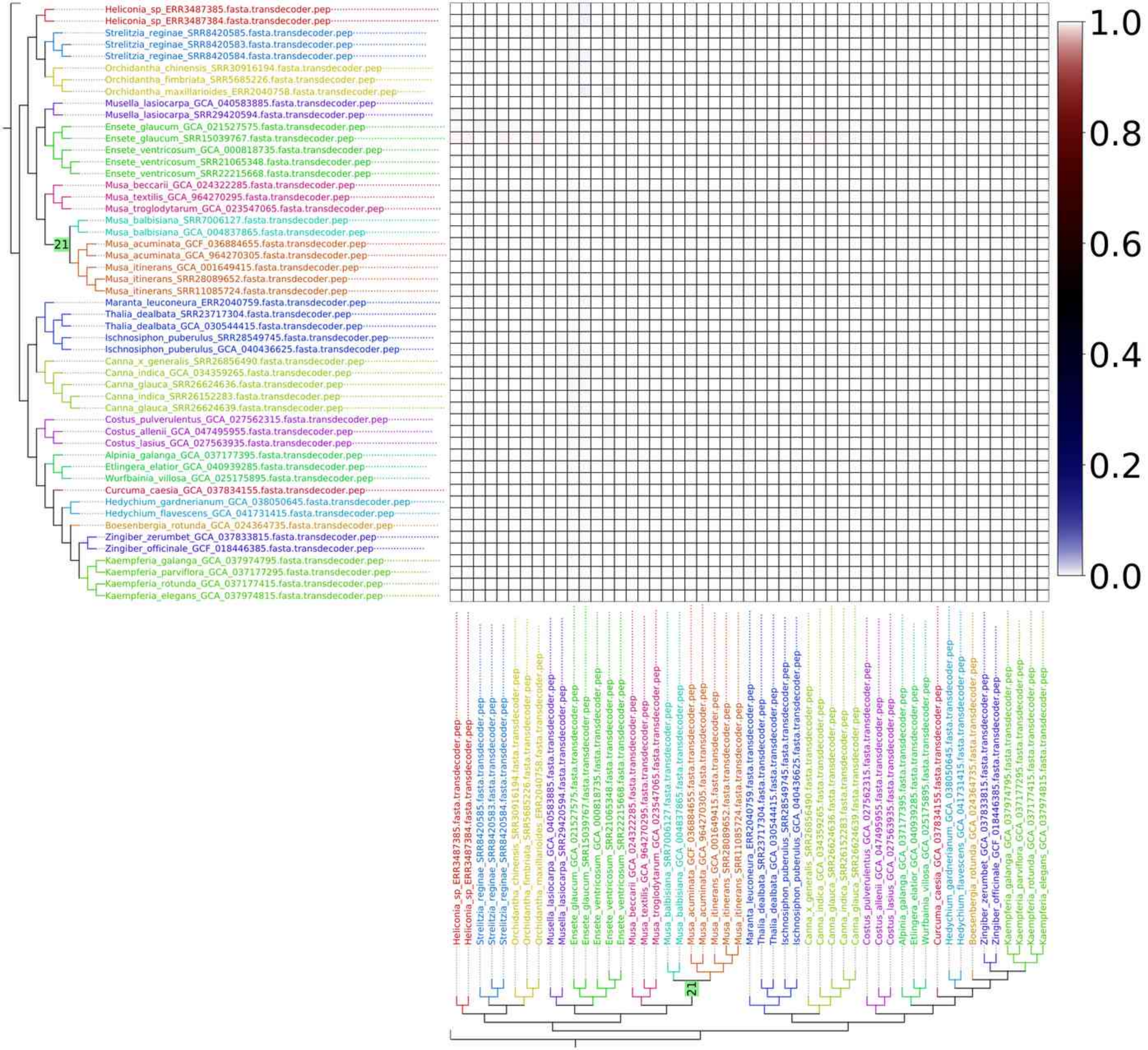
Heliconia_sp_ERR3487385.fasta.transdecoder.pep
Heliconia_sp_ERR3487384.fasta.transdecoder.pep
Strelitzia_reginae_SRR8420585.fasta.transdecoder.pep
Strelitzia_reginae_SRR8420583.fasta.transdecoder.pep
Strelitzia_reginae_SRR8420584.fasta.transdecoder.pep
Orchidantha_chinensis_SRR30916194.fasta.transdecoder.pep
Orchidantha_fimbriata_SRR5685226.fasta.transdecoder.pep
Orchidantha_maxillarioides_ERR2040758.fasta.transdecoder.pep
Musella_lasiocarpa_GCA_040583885.fasta.transdecoder.pep
Musella_lasiocarpa_SRR29420594.fasta.transdecoder.pep
Ensete_glaucum_GCA_021527575.fasta.transdecoder.pep
Ensete_glaucum_SRR15039767.fasta.transdecoder.pep
Ensete_ventricosum_GCA_000818735.fasta.transdecoder.pep
Ensete_ventricosum_SRR21065348.fasta.transdecoder.pep
Ensete_ventricosum_SRR22215668.fasta.transdecoder.pep
Musa_beccarii_GCA_024322285.fasta.transdecoder.pep
Musa_textilis_GCA_964270295.fasta.transdecoder.pep
Musa_troglodytarum_GCA_023547065.fasta.transdecoder.pep
Musa_balbisiana_SRR7006127.fasta.transdecoder.pep
Musa_balbisiana_GCA_004837865.fasta.transdecoder.pep
21
Musa_acuminata_GCF_036884655.fasta.transdecoder.pep
Musa_acuminata_GCA_964270305.fasta.transdecoder.pep
Musa_itinerans_GCA_001649415.fasta.transdecoder.pep
Musa_itinerans_SRR28089652.fasta.transdecoder.pep
Musa_itinerans_SRR11085724.fasta.transdecoder.pep
Maranta_leuconeura_ERR2040759.fasta.transdecoder.pep
Thalia_dealbata_SRR23717304.fasta.transdecoder.pep
Thalia_dealbata_GCA_030544415.fasta.transdecoder.pep
Ischnosiphon_puberulus_SRR28549745.fasta.transdecoder.pep
Ischnosiphon_puberulus_GCA_040436625.fasta.transdecoder.pep
Canna_x_generalis_SRR26856490.fasta.transdecoder.pep
Canna_indica_GCA_034359265.fasta.transdecoder.pep
Canna_glauca_SRR26624636.fasta.transdecoder.pep
Canna_indica_SRR26152283.fasta.transdecoder.pep
Canna_glauca_SRR26624639.fasta.transdecoder.pep
Costus_pulverulentus_GCA_027562315.fasta.transdecoder.pep
Costus_allenii_GCA_047495955.fasta.transdecoder.pep
Costus_lasius_GCA_027563935.fasta.transdecoder.pep
Alpinia_galanga_GCA_037177395.fasta.transdecoder.pep
Etlingera_elatior_GCA_040939285.fasta.transdecoder.pep
Wurfbainia_villosa_GCA_025175895.fasta.transdecoder.pep
Curcuma_caesia_GCA_037834155.fasta.transdecoder.pep
Hedychium_gardnerianum_GCA_038050645.fasta.transdecoder.pep
Hedychium_flavescens_GCA_041731415.fasta.transdecoder.pep
Boesenbergia_rotunda_GCA_024364735.fasta.transdecoder.pep
Zingiber_zerumbet_GCA_037833815.fasta.transdecoder.pep
Zingiber_officinale_GCF_018446385.fasta.transdecoder.pep
Kaempferia_galanga_GCA_037974795.fasta.transdecoder.pep
Kaempferia_parviflora_GCA_037177295.fasta.transdecoder.pep
Kaempferia_rotunda_GCA_037177415.fasta.transdecoder.pep
Kaempferia_elegans_GCA_037974815.fasta.transdecoder.pep
1.0
0.8
0.6
0.4
0.2
0.0

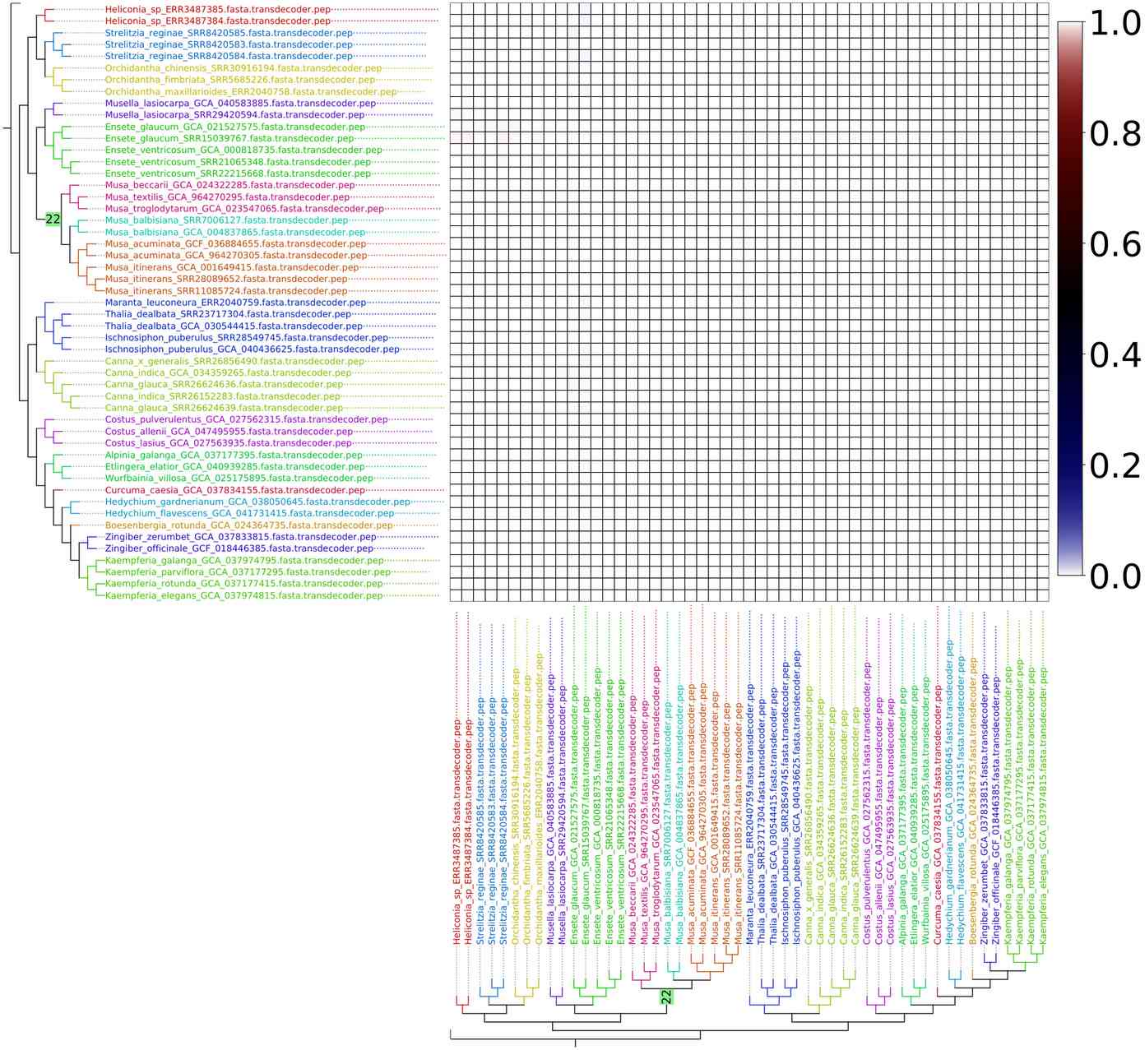
Heliconia_sp_ERR3487385.fasta.transdecoder.pep
Heliconia_sp_ERR3487384.fasta.transdecoder.pep
Strelitzia_reginae_SRR8420585.fasta.transdecoder.pep
Strelitzia_reginae_SRR8420583.fasta.transdecoder.pep
Strelitzia_reginae_SRR8420584.fasta.transdecoder.pep
Orchidantha_chinensis_SRR30916194.fasta.transdecoder.pep
Orchidantha_fimbriata_SRR5685226.fasta.transdecoder.pep
Orchidantha_maxillarioides_ERR2040758.fasta.transdecoder.pep
Musella_lasiocarpa_GCA_040583885.fasta.transdecoder.pep
Musella_lasiocarpa_SRR29420594.fasta.transdecoder.pep
Ensete_glaucum_GCA_021527575.fasta.transdecoder.pep
Ensete_glaucum_SRR15039767.fasta.transdecoder.pep
Ensete_ventricosum_GCA_000818735.fasta.transdecoder.pep
Ensete_ventricosum_SRR21065348.fasta.transdecoder.pep
Ensete_ventricosum_SRR22215668.fasta.transdecoder.pep
Musa_beccarii_GCA_024322285.fasta.transdecoder.pep
Musa_textilis_GCA_964270295.fasta.transdecoder.pep
Musa_troglodytarum_GCA_023547065.fasta.transdecoder.pep
22
Musa_balbisiana_SRR7006127.fasta.transdecoder.pep
Musa_balbisiana_GCA_004837865.fasta.transdecoder.pep
Musa_acuminata_GCF_036884655.fasta.transdecoder.pep
Musa_acuminata_GCA_964270305.fasta.transdecoder.pep
Musa_itinerans_GCA_001649415.fasta.transdecoder.pep
Musa_itinerans_SRR28089652.fasta.transdecoder.pep
Musa_itinerans_SRR11085724.fasta.transdecoder.pep
Maranta_leuconeura_ERR2040759.fasta.transdecoder.pep
Thalia_dealbata_SRR23717304.fasta.transdecoder.pep
Thalia_dealbata_GCA_030544415.fasta.transdecoder.pep
Ischnosiphon_puberulus_SRR28549745.fasta.transdecoder.pep
Ischnosiphon_puberulus_GCA_040436625.fasta.transdecoder.pep
Canna_x_generalis_SRR26856490.fasta.transdecoder.pep
Canna_indica_GCA_034359265.fasta.transdecoder.pep
Canna_glauca_SRR26624636.fasta.transdecoder.pep
Canna_indica_SRR26152283.fasta.transdecoder.pep
Canna_glauca_SRR26624639.fasta.transdecoder.pep
Costus_pulverulentus_GCA_027562315.fasta.transdecoder.pep
Costus_allenii_GCA_047495955.fasta.transdecoder.pep
Costus_lasius_GCA_027563935.fasta.transdecoder.pep
Alpinia_galanga_GCA_037177395.fasta.transdecoder.pep
Etlingera_elatior_GCA_040939285.fasta.transdecoder.pep
Wurfbainia_villosa_GCA_025175895.fasta.transdecoder.pep
Curcuma_caesia_GCA_037834155.fasta.transdecoder.pep
Hedychium_gardnerianum_GCA_038050645.fasta.transdecoder.pep
Hedychium_flavescens_GCA_041731415.fasta.transdecoder.pep
Boesenbergia_rotunda_GCA_024364735.fasta.transdecoder.pep
Zingiber_zerumbet_GCA_037833815.fasta.transdecoder.pep
Zingiber_officinale_GCF_018446385.fasta.transdecoder.pep
Kaempferia_galanga_GCA_037974795.fasta.transdecoder.pep
Kaempferia_parviflora_GCA_037177295.fasta.transdecoder.pep
Kaempferia_rotunda_GCA_037177415.fasta.transdecoder.pep
Kaempferia_elegans_GCA_037974815.fasta.transdecoder.pep
1.0
0.8
0.6
0.4
0.2
0.0

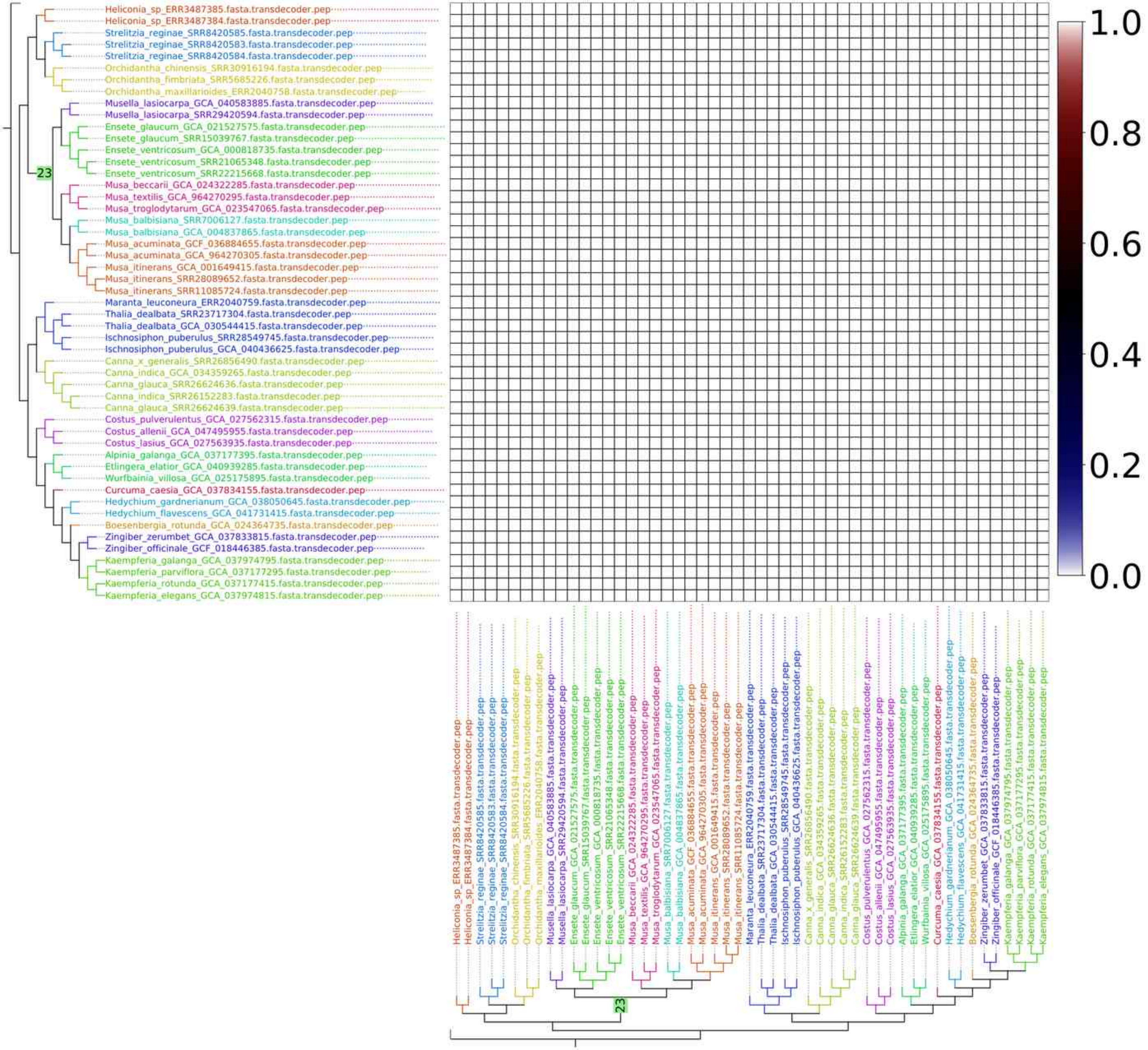
Heliconia_sp_ERR3487385.fasta.transdecoder.pep
Heliconia_sp_ERR3487384.fasta.transdecoder.pep
Strelitzia_reginae_SRR8420585.fasta.transdecoder.pep
Strelitzia_reginae_SRR8420583.fasta.transdecoder.pep
Strelitzia_reginae_SRR8420584.fasta.transdecoder.pep
Orchidantha_chinensis_SRR30916194.fasta.transdecoder.pep
Orchidantha_fimbriata_SRR5685226.fasta.transdecoder.pep
Orchidantha_maxillarioides_ERR2040758.fasta.transdecoder.pep
Musella_lasiocarpa_GCA_040583885.fasta.transdecoder.pep
Musella_lasiocarpa_SRR29420594.fasta.transdecoder.pep
Ensete_glaucum_GCA_021527575.fasta.transdecoder.pep
Ensete_glaucum_SRR15039767.fasta.transdecoder.pep
Ensete_ventricosum_GCA_000818735.fasta.transdecoder.pep
Ensete_ventricosum_SRR21065348.fasta.transdecoder.pep
Ensete_ventricosum_SRR22215668.fasta.transdecoder.pep
Musa_beccarii_GCA_024322285.fasta.transdecoder.pep
Musa_textilis_GCA_964270295.fasta.transdecoder.pep
Musa_troglodytarum_GCA_023547065.fasta.transdecoder.pep
Musa_balbisiana_SRR7006127.fasta.transdecoder.pep
Musa_balbisiana_GCA_004837865.fasta.transdecoder.pep
Musa_acuminata_GCF_036884655.fasta.transdecoder.pep
Musa_acuminata_GCA_964270305.fasta.transdecoder.pep
Musa_itinerans_GCA_001649415.fasta.transdecoder.pep
Musa_itinerans_SRR28089652.fasta.transdecoder.pep
Musa_itinerans_SRR11085724.fasta.transdecoder.pep
Maranta_leuconeura_ERR2040759.fasta.transdecoder.pep
Thalia_dealbata_SRR23717304.fasta.transdecoder.pep
Thalia_dealbata_GCA_030544415.fasta.transdecoder.pep
Ischnosiphon_puberulus_SRR28549745.fasta.transdecoder.pep
Ischnosiphon_puberulus_GCA_040436625.fasta.transdecoder.pep
Canna_x_generalis_SRR26856490.fasta.transdecoder.pep
Canna_indica_GCA_034359265.fasta.transdecoder.pep
Canna_glauca_SRR26624636.fasta.transdecoder.pep
Canna_indica_SRR26152283.fasta.transdecoder.pep
Canna_glauca_SRR26624639.fasta.transdecoder.pep
Costus_pulverulentus_GCA_027562315.fasta.transdecoder.pep
Costus_allenii_GCA_047495955.fasta.transdecoder.pep
Costus_lasius_GCA_027563935.fasta.transdecoder.pep
Alpinia_galanga_GCA_037177395.fasta.transdecoder.pep
Etlingera_elatior_GCA_040939285.fasta.transdecoder.pep
Wurfbainia_villosa_GCA_025175895.fasta.transdecoder.pep
Curcuma_caesia_GCA_037834155.fasta.transdecoder.pep
Hedychium_gardnerianum_GCA_038050645.fasta.transdecoder.pep
Hedychium_flavescens_GCA_041731415.fasta.transdecoder.pep
Boesenbergia_rotunda_GCA_024364735.fasta.transdecoder.pep
Zingiber_zerumbet_GCA_037833815.fasta.transdecoder.pep
Zingiber_officinale_GCF_018446385.fasta.transdecoder.pep
Kaempferia_galanga_GCA_037974795.fasta.transdecoder.pep
Kaempferia_parviflora_GCA_037177295.fasta.transdecoder.pep
Kaempferia_rotunda_GCA_037177415.fasta.transdecoder.pep
Kaempferia_elegans_GCA_037974815.fasta.transdecoder.pep
23
1.0
0.8
0.6
0.4
0.2
0.0

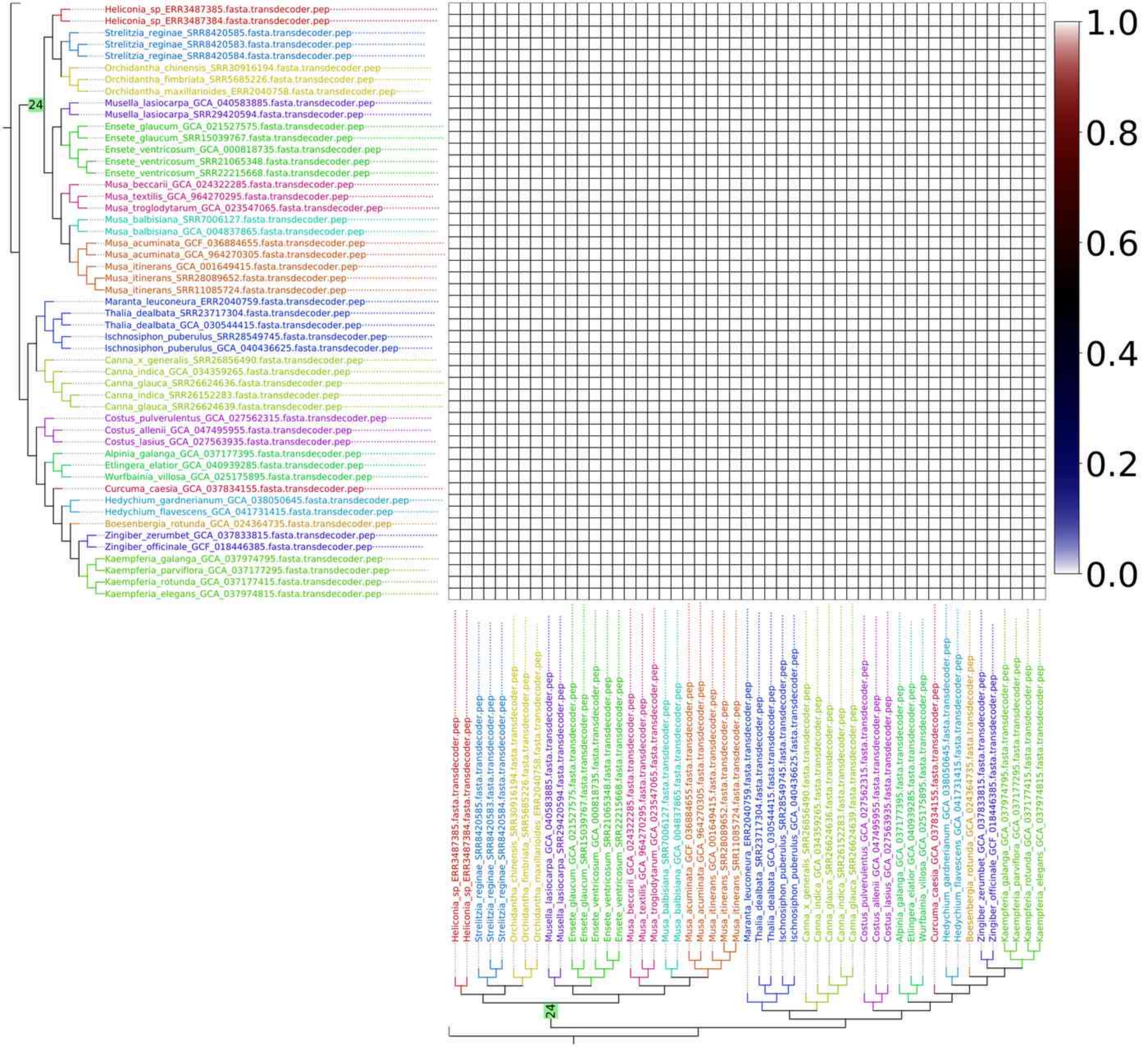
Heliconia_sp_ERR3487385.fasta.transdecoder.pep
Heliconia_sp_ERR3487384.fasta.transdecoder.pep
Strelitzia_reginae_SRR8420585.fasta.transdecoder.pep
Strelitzia_reginae_SRR8420583.fasta.transdecoder.pep
Strelitzia_reginae_SRR8420584.fasta.transdecoder.pep
Orchidantha_chinensis_SRR30916194.fasta.transdecoder.pep
Orchidantha_fimbriata_SRR5685226.fasta.transdecoder.pep
Orchidantha_maxillarioides_ERR2040758.fasta.transdecoder.pep
Musella_lasiocarpa_GCA_040583885.fasta.transdecoder.pep
Musella_lasiocarpa_SRR29420594.fasta.transdecoder.pep
Ensete_glaucum_GCA_021527575.fasta.transdecoder.pep
Ensete_glaucum_SRR15039767.fasta.transdecoder.pep
Ensete_ventricosum_GCA_000818735.fasta.transdecoder.pep
Ensete_ventricosum_SRR21065348.fasta.transdecoder.pep
Ensete_ventricosum_SRR22215668.fasta.transdecoder.pep
Musa_beccarii_GCA_024322285.fasta.transdecoder.pep
Musa_textilis_GCA_964270295.fasta.transdecoder.pep
Musa_troglodytarum_GCA_023547065.fasta.transdecoder.pep
Musa_balbisiana_SRR7006127.fasta.transdecoder.pep
Musa_balbisiana_GCA_004837865.fasta.transdecoder.pep
Musa_acuminata_GCF_036884655.fasta.transdecoder.pep
Musa_acuminata_GCA_964270305.fasta.transdecoder.pep
Musa_itinerans_GCA_001649415.fasta.transdecoder.pep
Musa_itinerans_SRR28089652.fasta.transdecoder.pep
Musa_itinerans_SRR11085724.fasta.transdecoder.pep
Maranta_leuconeura_ERR2040759.fasta.transdecoder.pep
Thalia_dealbata_SRR23717304.fasta.transdecoder.pep
Thalia_dealbata_GCA_030544415.fasta.transdecoder.pep
Ischnosiphon_puberulus_SRR28549745.fasta.transdecoder.pep
Ischnosiphon_puberulus_GCA_040436625.fasta.transdecoder.pep
Canna_x_generalis_SRR26856490.fasta.transdecoder.pep
Canna_indica_GCA_034359265.fasta.transdecoder.pep
Canna_glauca_SRR26624636.fasta.transdecoder.pep
Canna_indica_SRR26152283.fasta.transdecoder.pep
Canna_glauca_SRR26624639.fasta.transdecoder.pep
Costus_pulverulentus_GCA_027562315.fasta.transdecoder.pep
Costus_allenii_GCA_047495955.fasta.transdecoder.pep
Costus_lasius_GCA_027563935.fasta.transdecoder.pep
Alpinia_galanga_GCA_037177395.fasta.transdecoder.pep
Etlingera_elatior_GCA_040939285.fasta.transdecoder.pep
Wurfbainia_villosa_GCA_025175895.fasta.transdecoder.pep
Curcuma_caesia_GCA_037834155.fasta.transdecoder.pep
Hedychium_gardnerianum_GCA_038050645.fasta.transdecoder.pep
Hedychium_flavescens_GCA_041731415.fasta.transdecoder.pep
Boesenbergia_rotunda_GCA_024364735.fasta.transdecoder.pep
Zingiber_zerumbet_GCA_037833815.fasta.transdecoder.pep
Zingiber_officinale_GCF_018446385.fasta.transdecoder.pep
Kaempferia_galanga_GCA_037974795.fasta.transdecoder.pep
Kaempferia_parviflora_GCA_037177295.fasta.transdecoder.pep
Kaempferia_rotunda_GCA_037177415.fasta.transdecoder.pep
Kaempferia_elegans_GCA_037974815.fasta.transdecoder.pep
24
1.0
0.8
0.6
0.4
0.2
0.0

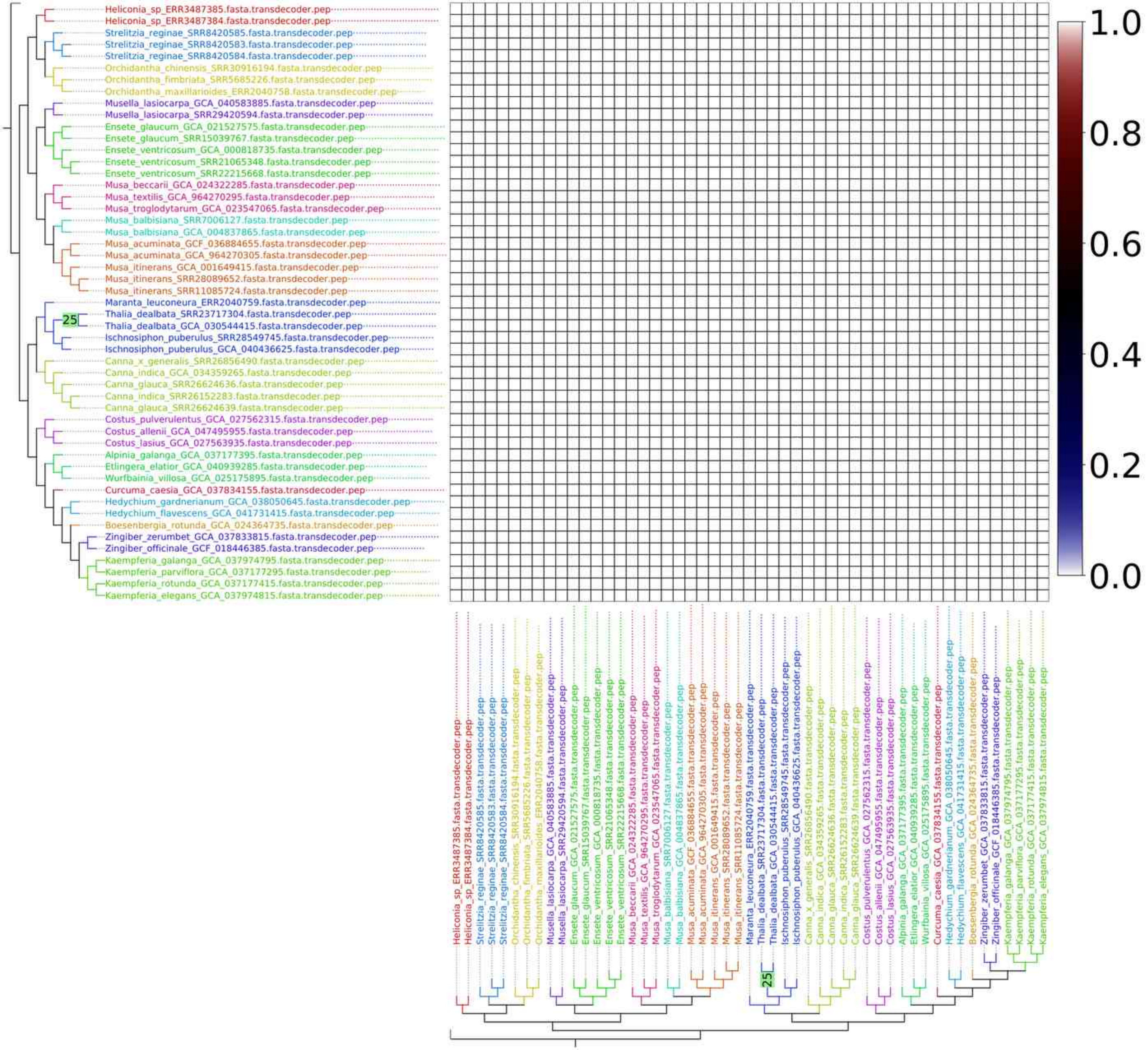
Heliconia_sp_ERR3487385.fasta.transdecoder.pep
Heliconia_sp_ERR3487384.fasta.transdecoder.pep
Strelitzia_reginae_SRR8420585.fasta.transdecoder.pep
Strelitzia_reginae_SRR8420583.fasta.transdecoder.pep
Strelitzia_reginae_SRR8420584.fasta.transdecoder.pep
Orchidantha_chinensis_SRR30916194.fasta.transdecoder.pep
Orchidantha_fimbriata_SRR5685226.fasta.transdecoder.pep
Orchidantha_maxillarioides_ERR2040758.fasta.transdecoder.pep
Musella_lasiocarpa_GCA_040583885.fasta.transdecoder.pep
Musella_lasiocarpa_SRR29420594.fasta.transdecoder.pep
Ensete_glaucum_GCA_021527575.fasta.transdecoder.pep
Ensete_glaucum_SRR15039767.fasta.transdecoder.pep
Ensete_ventricosum_GCA_000818735.fasta.transdecoder.pep
Ensete_ventricosum_SRR21065348.fasta.transdecoder.pep
Ensete_ventricosum_SRR22215668.fasta.transdecoder.pep
Musa_beccarii_GCA_024322285.fasta.transdecoder.pep
Musa_textilis_GCA_964270295.fasta.transdecoder.pep
Musa_troglodytarum_GCA_023547065.fasta.transdecoder.pep
Musa_balbisiana_SRR7006127.fasta.transdecoder.pep
Musa_balbisiana_GCA_004837865.fasta.transdecoder.pep
Musa_acuminata_GCF_036884655.fasta.transdecoder.pep
Musa_acuminata_GCA_964270305.fasta.transdecoder.pep
Musa_itinerans_GCA_001649415.fasta.transdecoder.pep
Musa_itinerans_SRR28089652.fasta.transdecoder.pep
Musa_itinerans_SRR11085724.fasta.transdecoder.pep
Maranta_leuconeura_ERR2040759.fasta.transdecoder.pep
25
Thalia_dealbata_SRR23717304.fasta.transdecoder.pep
Thalia_dealbata_GCA_030544415.fasta.transdecoder.pep
Ischnosiphon_puberulus_SRR28549745.fasta.transdecoder.pep
Ischnosiphon_puberulus_GCA_040436625.fasta.transdecoder.pep
Canna_x_generalis_SRR26856490.fasta.transdecoder.pep
Canna_indica_GCA_034359265.fasta.transdecoder.pep
Canna_glauca_SRR26624636.fasta.transdecoder.pep
Canna_indica_SRR26152283.fasta.transdecoder.pep
Canna_glauca_SRR26624639.fasta.transdecoder.pep
Costus_pulverulentus_GCA_027562315.fasta.transdecoder.pep
Costus_allenii_GCA_047495955.fasta.transdecoder.pep
Costus_lasius_GCA_027563935.fasta.transdecoder.pep
Alpinia_galanga_GCA_037177395.fasta.transdecoder.pep
Etlingera_elatior_GCA_040939285.fasta.transdecoder.pep
Wurfbainia_villosa_GCA_025175895.fasta.transdecoder.pep
Curcuma_caesia_GCA_037834155.fasta.transdecoder.pep
Hedychium_gardnerianum_GCA_038050645.fasta.transdecoder.pep
Hedychium_flavescens_GCA_041731415.fasta.transdecoder.pep
Boesenbergia_rotunda_GCA_024364735.fasta.transdecoder.pep
Zingiber_zerumbet_GCA_037833815.fasta.transdecoder.pep
Zingiber_officinale_GCF_018446385.fasta.transdecoder.pep
Kaempferia_galanga_GCA_037974795.fasta.transdecoder.pep
Kaempferia_parviflora_GCA_037177295.fasta.transdecoder.pep
Kaempferia_rotunda_GCA_037177415.fasta.transdecoder.pep
Kaempferia_elegans_GCA_037974815.fasta.transdecoder.pep
1.0
0.8
0.6
0.4
0.2
0.0

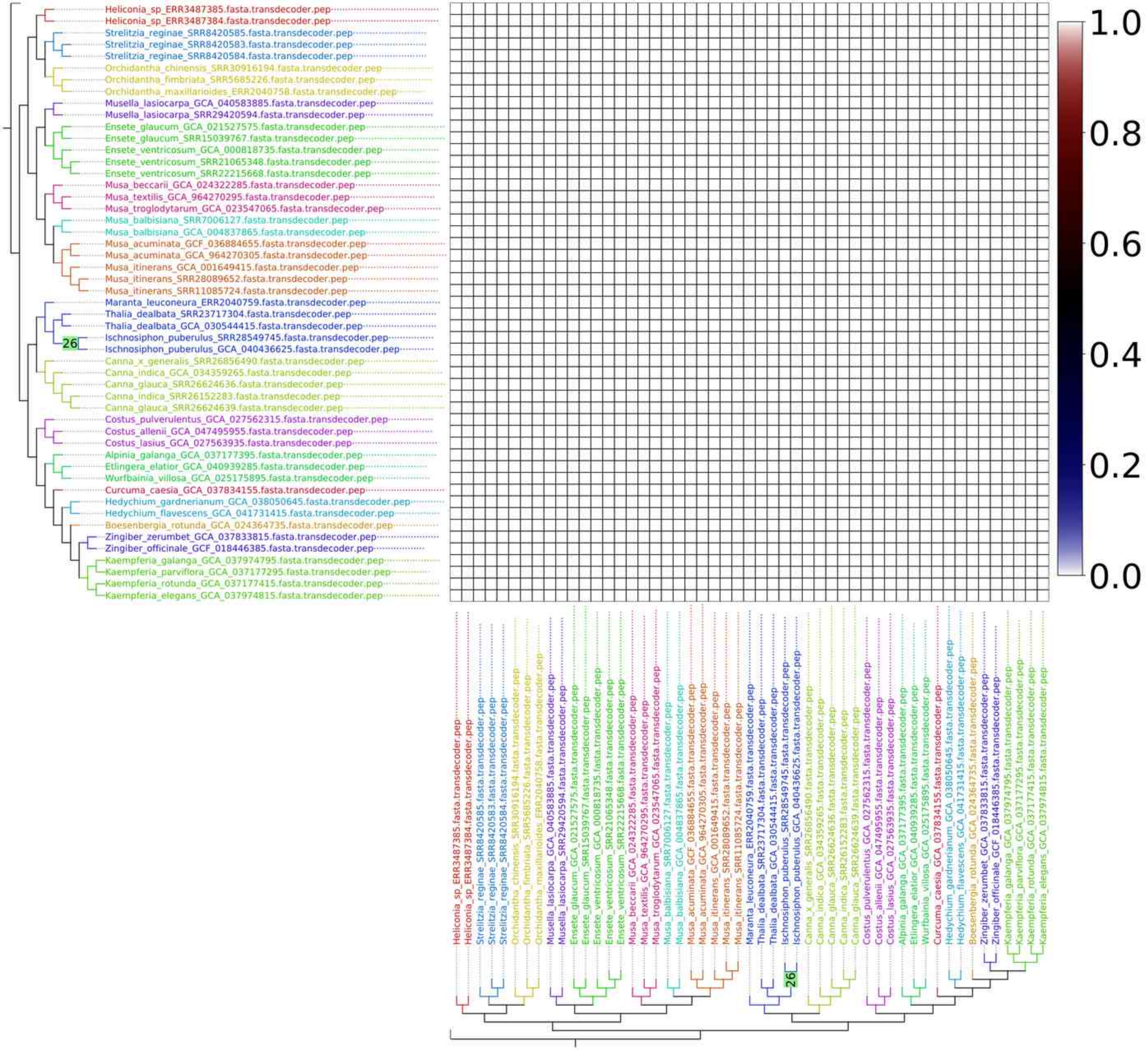
Heliconia_sp_ERR3487385.fasta.transdecoder.pep
Heliconia_sp_ERR3487384.fasta.transdecoder.pep
Strelitzia_reginae_SRR8420585.fasta.transdecoder.pep
Strelitzia_reginae_SRR8420583.fasta.transdecoder.pep
Strelitzia_reginae_SRR8420584.fasta.transdecoder.pep
Orchidantha_chinensis_SRR30916194.fasta.transdecoder.pep
Orchidantha_fimbriata_SRR5685226.fasta.transdecoder.pep
Orchidantha_maxillarioides_ERR2040758.fasta.transdecoder.pep
Musella_lasiocarpa_GCA_040583885.fasta.transdecoder.pep
Musella_lasiocarpa_SRR29420594.fasta.transdecoder.pep
Ensete_glaucum_GCA_021527575.fasta.transdecoder.pep
Ensete_glaucum_SRR15039767.fasta.transdecoder.pep
Ensete_ventricosum_GCA_000818735.fasta.transdecoder.pep
Ensete_ventricosum_SRR21065348.fasta.transdecoder.pep
Ensete_ventricosum_SRR22215668.fasta.transdecoder.pep
Musa_beccarii_GCA_024322285.fasta.transdecoder.pep
Musa_textilis_GCA_964270295.fasta.transdecoder.pep
Musa_troglodytarum_GCA_023547065.fasta.transdecoder.pep
Musa_balbisiana_SRR7006127.fasta.transdecoder.pep
Musa_balbisiana_GCA_004837865.fasta.transdecoder.pep
Musa_acuminata_GCF_036884655.fasta.transdecoder.pep
Musa_acuminata_GCA_964270305.fasta.transdecoder.pep
Musa_itinerans_GCA_001649415.fasta.transdecoder.pep
Musa_itinerans_SRR28089652.fasta.transdecoder.pep
Musa_itinerans_SRR11085724.fasta.transdecoder.pep
Maranta_leuconeura_ERR2040759.fasta.transdecoder.pep
Thalia_dealbata_SRR23717304.fasta.transdecoder.pep
Thalia_dealbata_GCA_030544415.fasta.transdecoder.pep
26
Ischnosiphon_puberulus_SRR28549745.fasta.transdecoder.pep
Ischnosiphon_puberulus_GCA_040436625.fasta.transdecoder.pep
Canna_x_generalis_SRR26856490.fasta.transdecoder.pep
Canna_indica_GCA_034359265.fasta.transdecoder.pep
Canna_glauca_SRR26624636.fasta.transdecoder.pep
Canna_indica_SRR26152283.fasta.transdecoder.pep
Canna_glauca_SRR26624639.fasta.transdecoder.pep
Costus_pulverulentus_GCA_027562315.fasta.transdecoder.pep
Costus_allenii_GCA_047495955.fasta.transdecoder.pep
Costus_lasius_GCA_027563935.fasta.transdecoder.pep
Alpinia_galanga_GCA_037177395.fasta.transdecoder.pep
Etlingera_elatior_GCA_040939285.fasta.transdecoder.pep
Wurfbainia_villosa_GCA_025175895.fasta.transdecoder.pep
Curcuma_caesia_GCA_037834155.fasta.transdecoder.pep
Hedychium_gardnerianum_GCA_038050645.fasta.transdecoder.pep
Hedychium_flavescens_GCA_041731415.fasta.transdecoder.pep
Boesenbergia_rotunda_GCA_024364735.fasta.transdecoder.pep
Zingiber_zerumbet_GCA_037833815.fasta.transdecoder.pep
Zingiber_officinale_GCF_018446385.fasta.transdecoder.pep
Kaempferia_galanga_GCA_037974795.fasta.transdecoder.pep
Kaempferia_parviflora_GCA_037177295.fasta.transdecoder.pep
Kaempferia_rotunda_GCA_037177415.fasta.transdecoder.pep
Kaempferia_elegans_GCA_037974815.fasta.transdecoder.pep
1.0
0.8
0.6
0.4
0.2
0.0

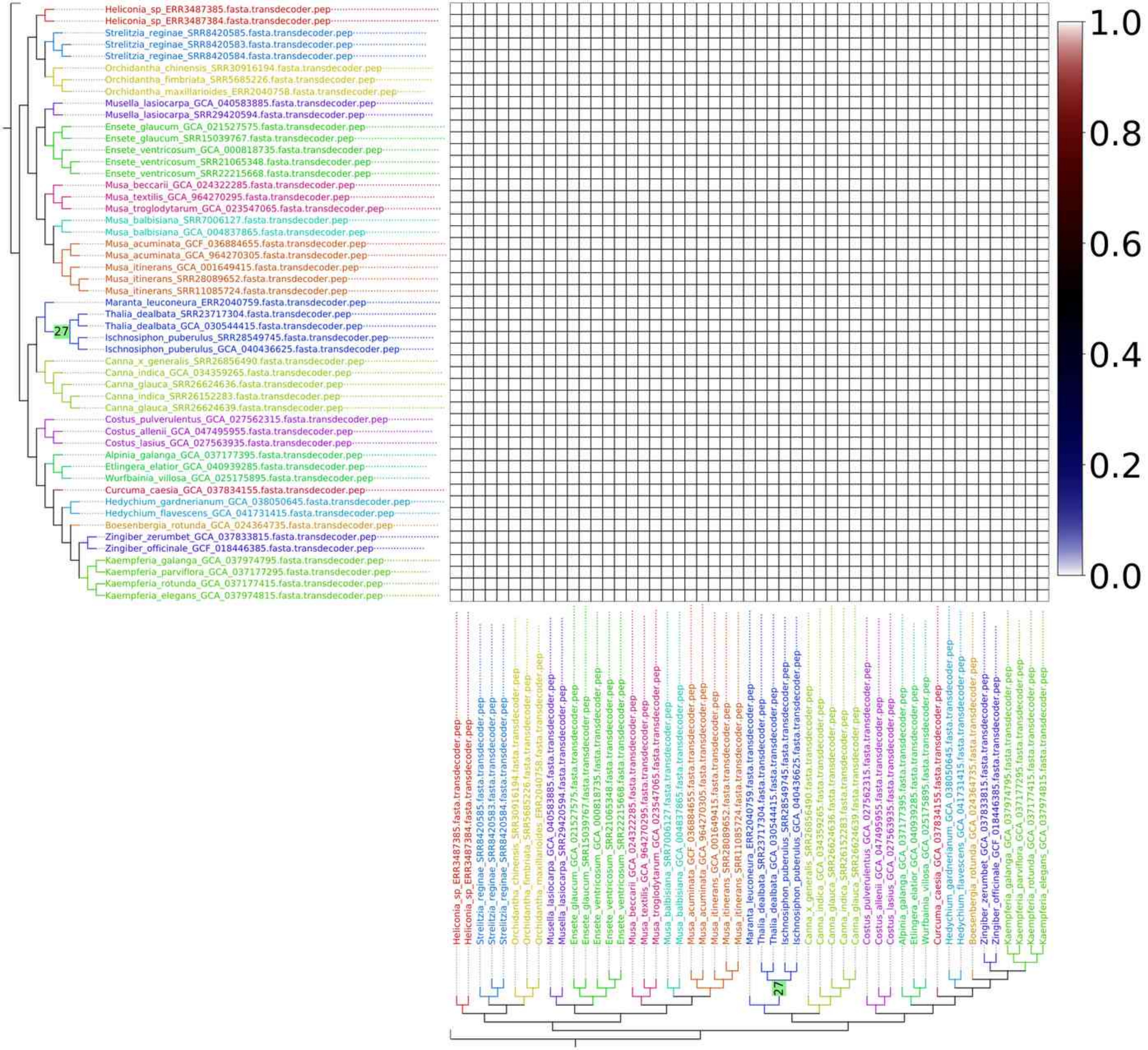

Heliconia_sp_ERR3487385.fasta.transdecoder.pep
Heliconia_sp_ERR3487384.fasta.transdecoder.pep
Strelitzia_reginae_SRR8420585.fasta.transdecoder.pep
Strelitzia_reginae_SRR8420583.fasta.transdecoder.pep
Strelitzia_reginae_SRR8420584.fasta.transdecoder.pep
Orchidantha_chinensis_SRR30916194.fasta.transdecoder.pep
Orchidantha_fimbriata_SRR5685226.fasta.transdecoder.pep
Orchidantha_maxillarioides_ERR2040758.fasta.transdecoder.pep
Musella_lasiocarpa_GCA_040583885.fasta.transdecoder.pep
Musella_lasiocarpa_SRR29420594.fasta.transdecoder.pep
Ensete_glaucum_GCA_021527575.fasta.transdecoder.pep
Ensete_glaucum_SRR15039767.fasta.transdecoder.pep
Ensete_ventricosum_GCA_000818735.fasta.transdecoder.pep
Ensete_ventricosum_SRR21065348.fasta.transdecoder.pep
Ensete_ventricosum_SRR22215668.fasta.transdecoder.pep
Musa_beccarii_GCA_024322285.fasta.transdecoder.pep
Musa_textilis_GCA_964270295.fasta.transdecoder.pep
Musa_troglodytarum_GCA_023547065.fasta.transdecoder.pep
Musa_balbisiana_SRR7006127.fasta.transdecoder.pep
Musa_balbisiana_GCA_004837865.fasta.transdecoder.pep
Musa_acuminata_GCF_036884655.fasta.transdecoder.pep
Musa_acuminata_GCA_964270305.fasta.transdecoder.pep
Musa_itinerans_GCA_001649415.fasta.transdecoder.pep
Musa_itinerans_SRR28089652.fasta.transdecoder.pep
Musa_itinerans_SRR11085724.fasta.transdecoder.pep
Maranta_leuconeura_ERR2040759.fasta.transdecoder.pep
Thalia_dealbata_SRR23717304.fasta.transdecoder.pep
Thalia_dealbata_GCA_030544415.fasta.transdecoder.pep
Ischnosiphon_puberulus_SRR28549745.fasta.transdecoder.pep
Ischnosiphon_puberulus_GCA_040436625.fasta.transdecoder.pep
Canna_x_generalis_SRR26856490.fasta.transdecoder.pep
Canna_indica_GCA_034359265.fasta.transdecoder.pep
Canna_glauca_SRR26624636.fasta.transdecoder.pep
Canna_indica_SRR26152283.fasta.transdecoder.pep
Canna_glauca_SRR26624639.fasta.transdecoder.pep
Costus_pulverulentus_GCA_027562315.fasta.transdecoder.pep
Costus_allenii_GCA_047495955.fasta.transdecoder.pep
Costus_lasius_GCA_027563935.fasta.transdecoder.pep
Alpinia_galanga_GCA_037177395.fasta.transdecoder.pep
Etlingera_elatior_GCA_040939285.fasta.transdecoder.pep
Wurfbainia_villosa_GCA_025175895.fasta.transdecoder.pep
Curcuma_caesia_GCA_037834155.fasta.transdecoder.pep
Hedychium_gardnerianum_GCA_038050645.fasta.transdecoder.pep
Hedychium_flavescens_GCA_041731415.fasta.transdecoder.pep
Boesenbergia_rotunda_GCA_024364735.fasta.transdecoder.pep
Zingiber_zerumbet_GCA_037833815.fasta.transdecoder.pep
Zingiber_officinale_GCF_018446385.fasta.transdecoder.pep
Kaempferia_galanga_GCA_037974795.fasta.transdecoder.pep
Kaempferia_parviflora_GCA_037177295.fasta.transdecoder.pep
Kaempferia_rotunda_GCA_037177415.fasta.transdecoder.pep
Kaempferia_elegans_GCA_037974815.fasta.transdecoder.pep
27
1.0
0.8
0.6
0.4
0.2
0.0

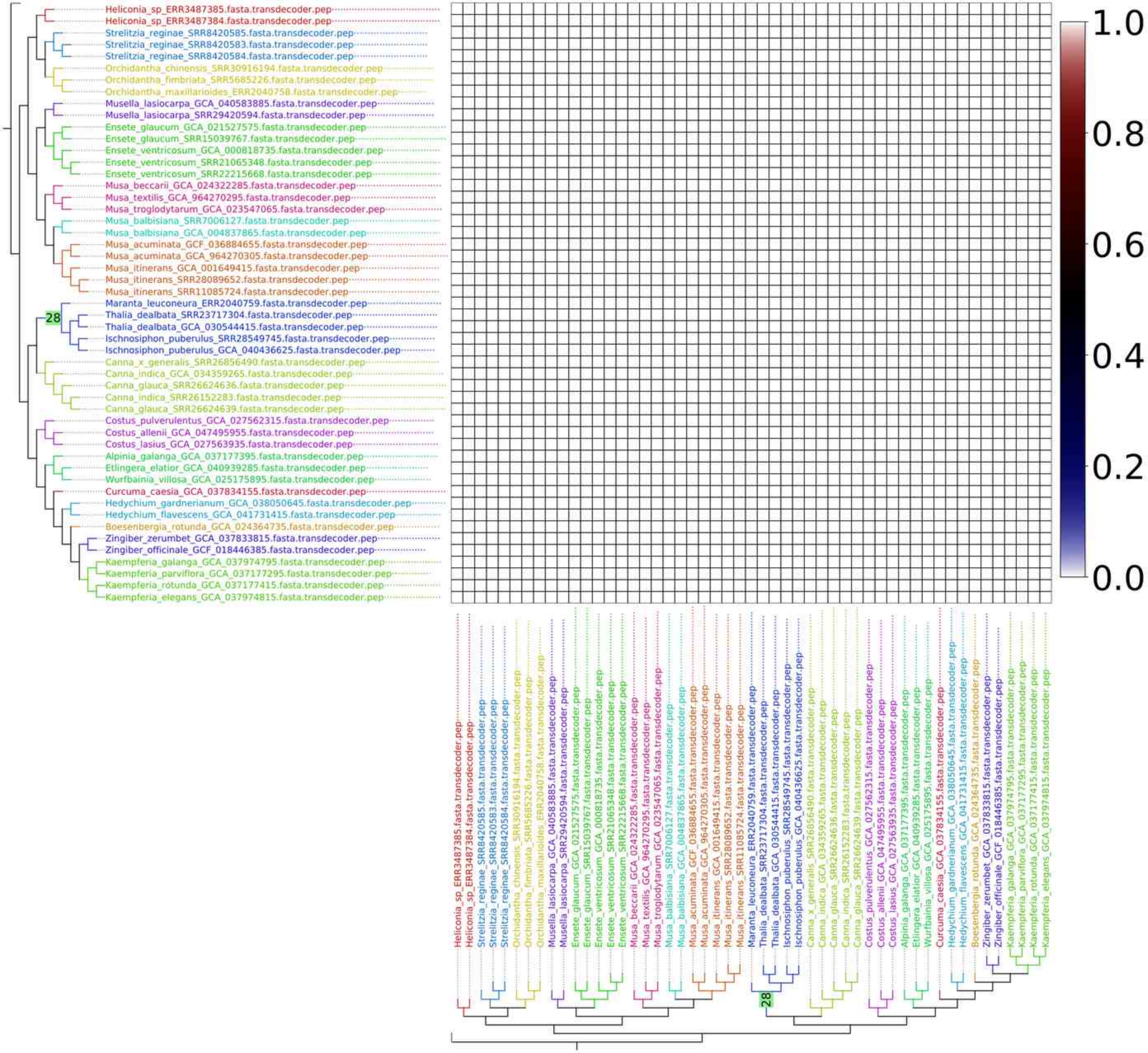

Heliconia_sp_ERR3487385.fasta.transdecoder.pep
Heliconia_sp_ERR3487384.fasta.transdecoder.pep
Strelitzia_reginae_SRR8420585.fasta.transdecoder.pep
Strelitzia_reginae_SRR8420583.fasta.transdecoder.pep
Strelitzia_reginae_SRR8420584.fasta.transdecoder.pep
Orchidantha_chinensis_SRR30916194.fasta.transdecoder.pep
Orchidantha_fimbriata_SRR5685226.fasta.transdecoder.pep
Orchidantha_maxillarioides_ERR2040758.fasta.transdecoder.pep
Musella_lasiocarpa_GCA_040583885.fasta.transdecoder.pep
Musella_lasiocarpa_SRR29420594.fasta.transdecoder.pep
Ensete_glaucum_GCA_021527575.fasta.transdecoder.pep
Ensete_glaucum_SRR15039767.fasta.transdecoder.pep
Ensete_ventricosum_GCA_000818735.fasta.transdecoder.pep
Ensete_ventricosum_SRR21065348.fasta.transdecoder.pep
Ensete_ventricosum_SRR22215668.fasta.transdecoder.pep
Musa_beccarii_GCA_024322285.fasta.transdecoder.pep
Musa_textilis_GCA_964270295.fasta.transdecoder.pep
Musa_troglodytarum_GCA_023547065.fasta.transdecoder.pep
Musa_balbisiana_SRR7006127.fasta.transdecoder.pep
Musa_balbisiana_GCA_004837865.fasta.transdecoder.pep
Musa_acuminata_GCF_036884655.fasta.transdecoder.pep
Musa_acuminata_GCA_964270305.fasta.transdecoder.pep
Musa_itinerans_GCA_001649415.fasta.transdecoder.pep
Musa_itinerans_SRR28089652.fasta.transdecoder.pep
Musa_itinerans_SRR11085724.fasta.transdecoder.pep
Maranta_leuconeura_ERR2040759.fasta.transdecoder.pep
28
Thalia_dealbata_SRR23717304.fasta.transdecoder.pep
Thalia_dealbata_GCA_030544415.fasta.transdecoder.pep
Ischnosiphon_puberulus_SRR28549745.fasta.transdecoder.pep
Ischnosiphon_puberulus_GCA_040436625.fasta.transdecoder.pep
Canna_x_generalis_SRR26856490.fasta.transdecoder.pep
Canna_indica_GCA_034359265.fasta.transdecoder.pep
Canna_glauca_SRR26624636.fasta.transdecoder.pep
Canna_indica_SRR26152283.fasta.transdecoder.pep
Canna_glauca_SRR26624639.fasta.transdecoder.pep
Costus_pulverulentus_GCA_027562315.fasta.transdecoder.pep
Costus_allenii_GCA_047495955.fasta.transdecoder.pep
Costus_lasius_GCA_027563935.fasta.transdecoder.pep
Alpinia_galanga_GCA_037177395.fasta.transdecoder.pep
Etlingera_elatior_GCA_040939285.fasta.transdecoder.pep
Wurfbainia_villosa_GCA_025175895.fasta.transdecoder.pep
Curcuma_caesia_GCA_037834155.fasta.transdecoder.pep
Hedychium_gardnerianum_GCA_038050645.fasta.transdecoder.pep
Hedychium_flavescens_GCA_041731415.fasta.transdecoder.pep
Boesenbergia_rotunda_GCA_024364735.fasta.transdecoder.pep
Zingiber_zerumbet_GCA_037833815.fasta.transdecoder.pep
Zingiber_officinale_GCF_018446385.fasta.transdecoder.pep
Kaempferia_galanga_GCA_037974795.fasta.transdecoder.pep
Kaempferia_parviflora_GCA_037177295.fasta.transdecoder.pep
Kaempferia_rotunda_GCA_037177415.fasta.transdecoder.pep
Kaempferia_elegans_GCA_037974815.fasta.transdecoder.pep
1.0
0.8
0.6
0.4
0.2
0.0

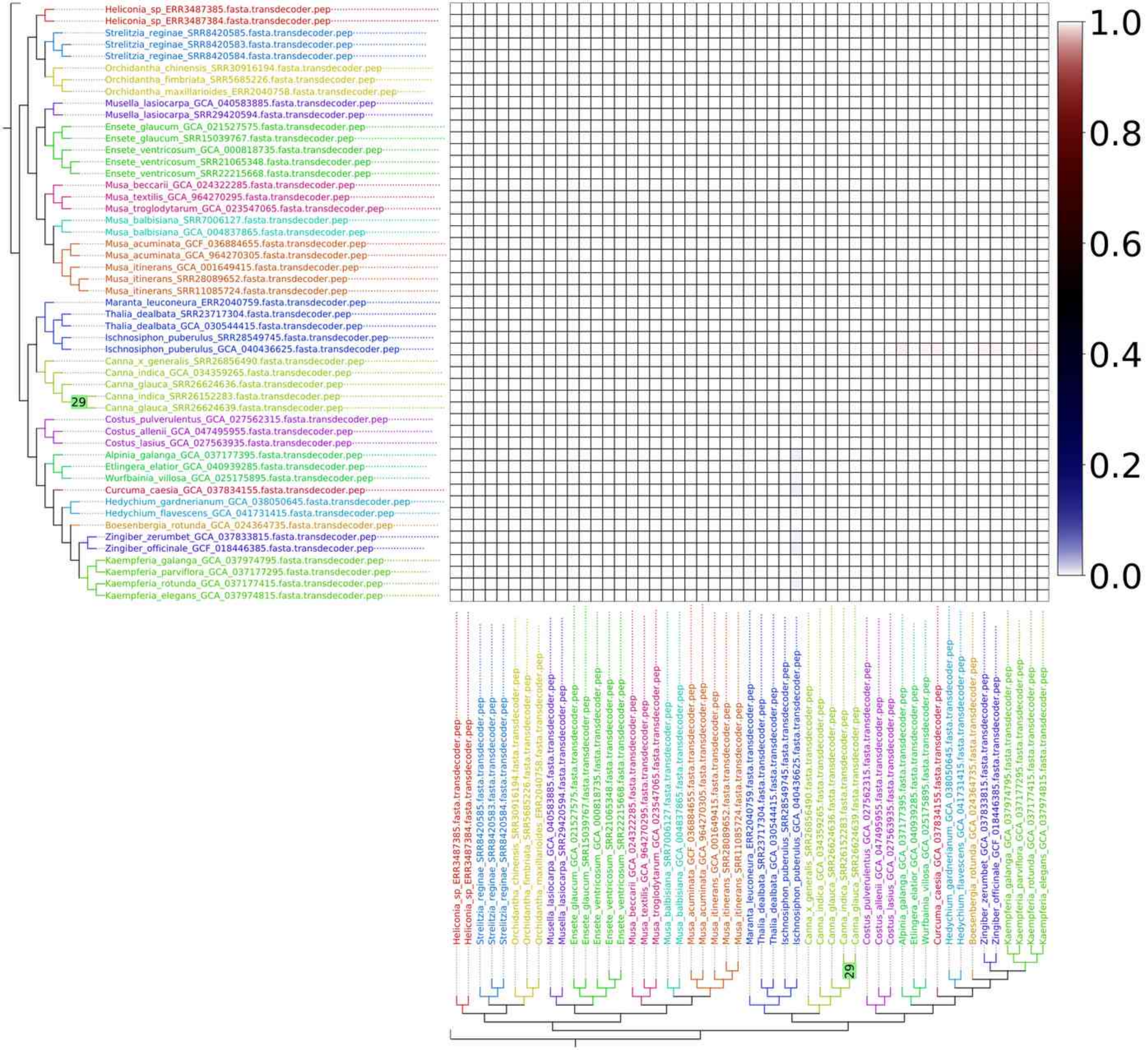
Heliconia_sp_ERR3487385.fasta.transdecoder.pep
Heliconia_sp_ERR3487384.fasta.transdecoder.pep
Strelitzia_reginae_SRR8420585.fasta.transdecoder.pep
Strelitzia_reginae_SRR8420583.fasta.transdecoder.pep
Strelitzia_reginae_SRR8420584.fasta.transdecoder.pep
Orchidantha_chinensis_SRR30916194.fasta.transdecoder.pep
Orchidantha_fimbriata_SRR5685226.fasta.transdecoder.pep
Orchidantha_maxillarioides_ERR2040758.fasta.transdecoder.pep
Musella_lasiocarpa_GCA_040583885.fasta.transdecoder.pep
Musella_lasiocarpa_SRR29420594.fasta.transdecoder.pep
Ensete_glaucum_GCA_021527575.fasta.transdecoder.pep
Ensete_glaucum_SRR15039767.fasta.transdecoder.pep
Ensete_ventricosum_GCA_000818735.fasta.transdecoder.pep
Ensete_ventricosum_SRR21065348.fasta.transdecoder.pep
Ensete_ventricosum_SRR22215668.fasta.transdecoder.pep
Musa_beccarii_GCA_024322285.fasta.transdecoder.pep
Musa_textilis_GCA_964270295.fasta.transdecoder.pep
Musa_troglodytarum_GCA_023547065.fasta.transdecoder.pep
Musa_balbisiana_SRR7006127.fasta.transdecoder.pep
Musa_balbisiana_GCA_004837865.fasta.transdecoder.pep
Musa_acuminata_GCF_036884655.fasta.transdecoder.pep
Musa_acuminata_GCA_964270305.fasta.transdecoder.pep
Musa_itinerans_GCA_001649415.fasta.transdecoder.pep
Musa_itinerans_SRR28089652.fasta.transdecoder.pep
Musa_itinerans_SRR11085724.fasta.transdecoder.pep
Maranta_leuconeura_ERR2040759.fasta.transdecoder.pep
Thalia_dealbata_SRR23717304.fasta.transdecoder.pep
Thalia_dealbata_GCA_030544415.fasta.transdecoder.pep
Ischnosiphon_puberulus_SRR28549745.fasta.transdecoder.pep
Ischnosiphon_puberulus_GCA_040436625.fasta.transdecoder.pep
Canna_x_generalis_SRR26856490.fasta.transdecoder.pep
Canna_indica_GCA_034359265.fasta.transdecoder.pep
Canna_glauca_SRR26624636.fasta.transdecoder.pep
Canna_indica_SRR26152283.fasta.transdecoder.pep
Canna_glauca_SRR26624639.fasta.transdecoder.pep
29
Costus_pulverulentus_GCA_027562315.fasta.transdecoder.pep
Costus_allenii_GCA_047495955.fasta.transdecoder.pep
Costus_lasius_GCA_027563935.fasta.transdecoder.pep
Alpinia_galanga_GCA_037177395.fasta.transdecoder.pep
Etlingera_elatior_GCA_040939285.fasta.transdecoder.pep
Wurfbainia_villosa_GCA_025175895.fasta.transdecoder.pep
Curcuma_caesia_GCA_037834155.fasta.transdecoder.pep
Hedychium_gardnerianum_GCA_038050645.fasta.transdecoder.pep
Hedychium_flavescens_GCA_041731415.fasta.transdecoder.pep
Boesenbergia_rotunda_GCA_024364735.fasta.transdecoder.pep
Zingiber_zerumbet_GCA_037833815.fasta.transdecoder.pep
Zingiber_officinale_GCF_018446385.fasta.transdecoder.pep
Kaempferia_galanga_GCA_037974795.fasta.transdecoder.pep
Kaempferia_parviflora_GCA_037177295.fasta.transdecoder.pep
Kaempferia_rotunda_GCA_037177415.fasta.transdecoder.pep
Kaempferia_elegans_GCA_037974815.fasta.transdecoder.pep
1.0
0.8
0.6
0.4
0.2
0.0

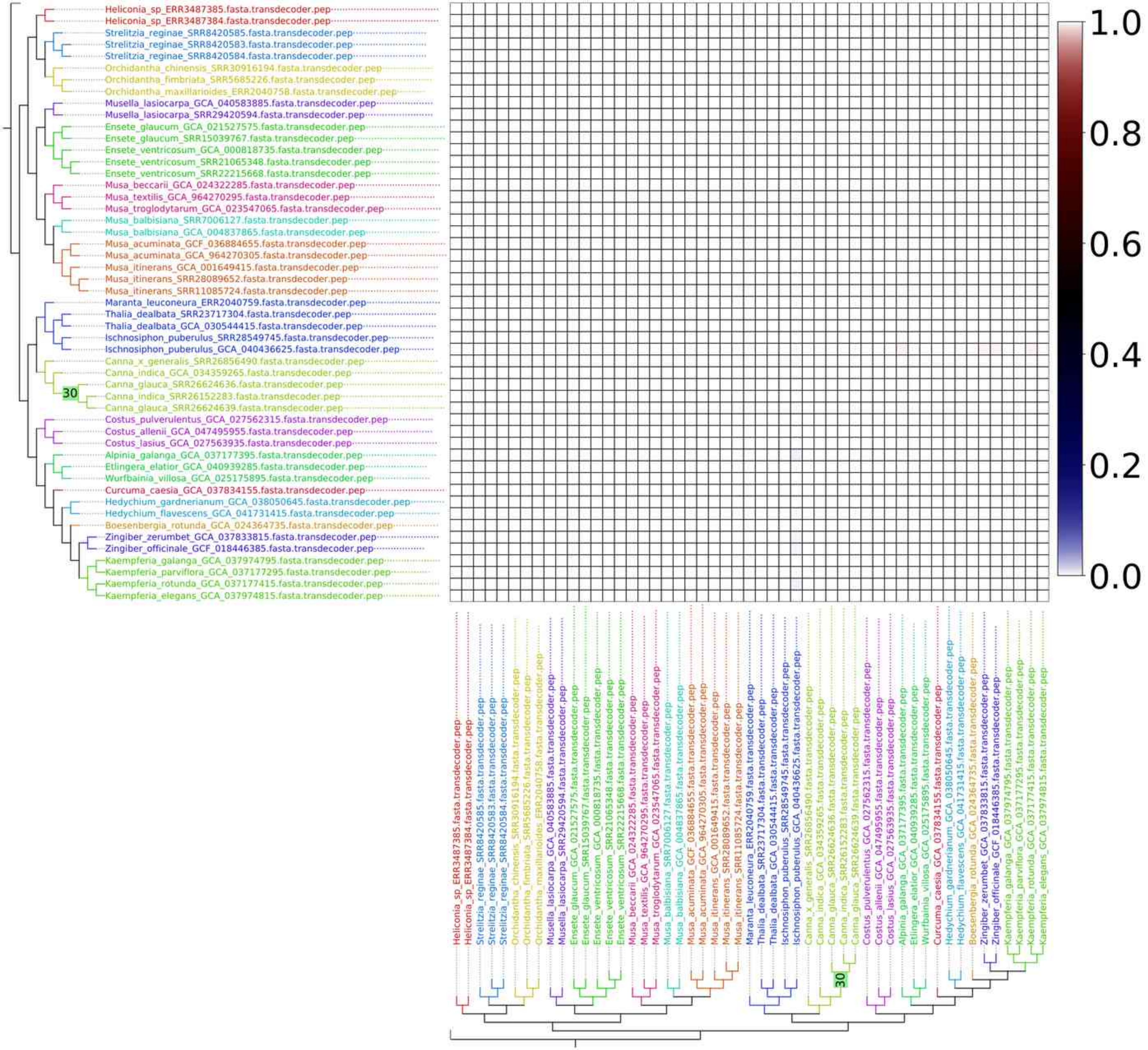
Heliconia_sp_ERR3487385.fasta.transdecoder.pep
Heliconia_sp_ERR3487384.fasta.transdecoder.pep
Strelitzia_reginae_SRR8420585.fasta.transdecoder.pep
Strelitzia_reginae_SRR8420583.fasta.transdecoder.pep
Strelitzia_reginae_SRR8420584.fasta.transdecoder.pep
Orchidantha_chinensis_SRR30916194.fasta.transdecoder.pep
Orchidantha_fimbriata_SRR5685226.fasta.transdecoder.pep
Orchidantha_maxillarioides_ERR2040758.fasta.transdecoder.pep
Musella_lasiocarpa_GCA_040583885.fasta.transdecoder.pep
Musella_lasiocarpa_SRR29420594.fasta.transdecoder.pep
Ensete_glaucum_GCA_021527575.fasta.transdecoder.pep
Ensete_glaucum_SRR15039767.fasta.transdecoder.pep
Ensete_ventricosum_GCA_000818735.fasta.transdecoder.pep
Ensete_ventricosum_SRR21065348.fasta.transdecoder.pep
Ensete_ventricosum_SRR22215668.fasta.transdecoder.pep
Musa_beccarii_GCA_024322285.fasta.transdecoder.pep
Musa_textilis_GCA_964270295.fasta.transdecoder.pep
Musa_troglodytarum_GCA_023547065.fasta.transdecoder.pep
Musa_balbisiana_SRR7006127.fasta.transdecoder.pep
Musa_balbisiana_GCA_004837865.fasta.transdecoder.pep
Musa_acuminata_GCF_036884655.fasta.transdecoder.pep
Musa_acuminata_GCA_964270305.fasta.transdecoder.pep
Musa_itinerans_GCA_001649415.fasta.transdecoder.pep
Musa_itinerans_SRR28089652.fasta.transdecoder.pep
Musa_itinerans_SRR11085724.fasta.transdecoder.pep
Maranta_leuconeura_ERR2040759.fasta.transdecoder.pep
Thalia_dealbata_SRR23717304.fasta.transdecoder.pep
Thalia_dealbata_GCA_030544415.fasta.transdecoder.pep
Ischnosiphon_puberulus_SRR28549745.fasta.transdecoder.pep
Ischnosiphon_puberulus_GCA_040436625.fasta.transdecoder.pep
Canna_x_generalis_SRR26856490.fasta.transdecoder.pep
Canna_indica_GCA_034359265.fasta.transdecoder.pep
Canna_glauca_SRR26624636.fasta.transdecoder.pep
30
Canna_indica_SRR26152283.fasta.transdecoder.pep
Canna_glauca_SRR26624639.fasta.transdecoder.pep
Costus_pulverulentus_GCA_027562315.fasta.transdecoder.pep
Costus_allenii_GCA_047495955.fasta.transdecoder.pep
Costus_lasius_GCA_027563935.fasta.transdecoder.pep
Alpinia_galanga_GCA_037177395.fasta.transdecoder.pep
Etlingera_elatior_GCA_040939285.fasta.transdecoder.pep
Wurfbainia_villosa_GCA_025175895.fasta.transdecoder.pep
Curcuma_caesia_GCA_037834155.fasta.transdecoder.pep
Hedychium_gardnerianum_GCA_038050645.fasta.transdecoder.pep
Hedychium_flavescens_GCA_041731415.fasta.transdecoder.pep
Boesenbergia_rotunda_GCA_024364735.fasta.transdecoder.pep
Zingiber_zerumbet_GCA_037833815.fasta.transdecoder.pep
Zingiber_officinale_GCF_018446385.fasta.transdecoder.pep
Kaempferia_galanga_GCA_037974795.fasta.transdecoder.pep
Kaempferia_parviflora_GCA_037177295.fasta.transdecoder.pep
Kaempferia_rotunda_GCA_037177415.fasta.transdecoder.pep
Kaempferia_elegans_GCA_037974815.fasta.transdecoder.pep
1.0
0.8
0.6
0.4
0.2
0.0

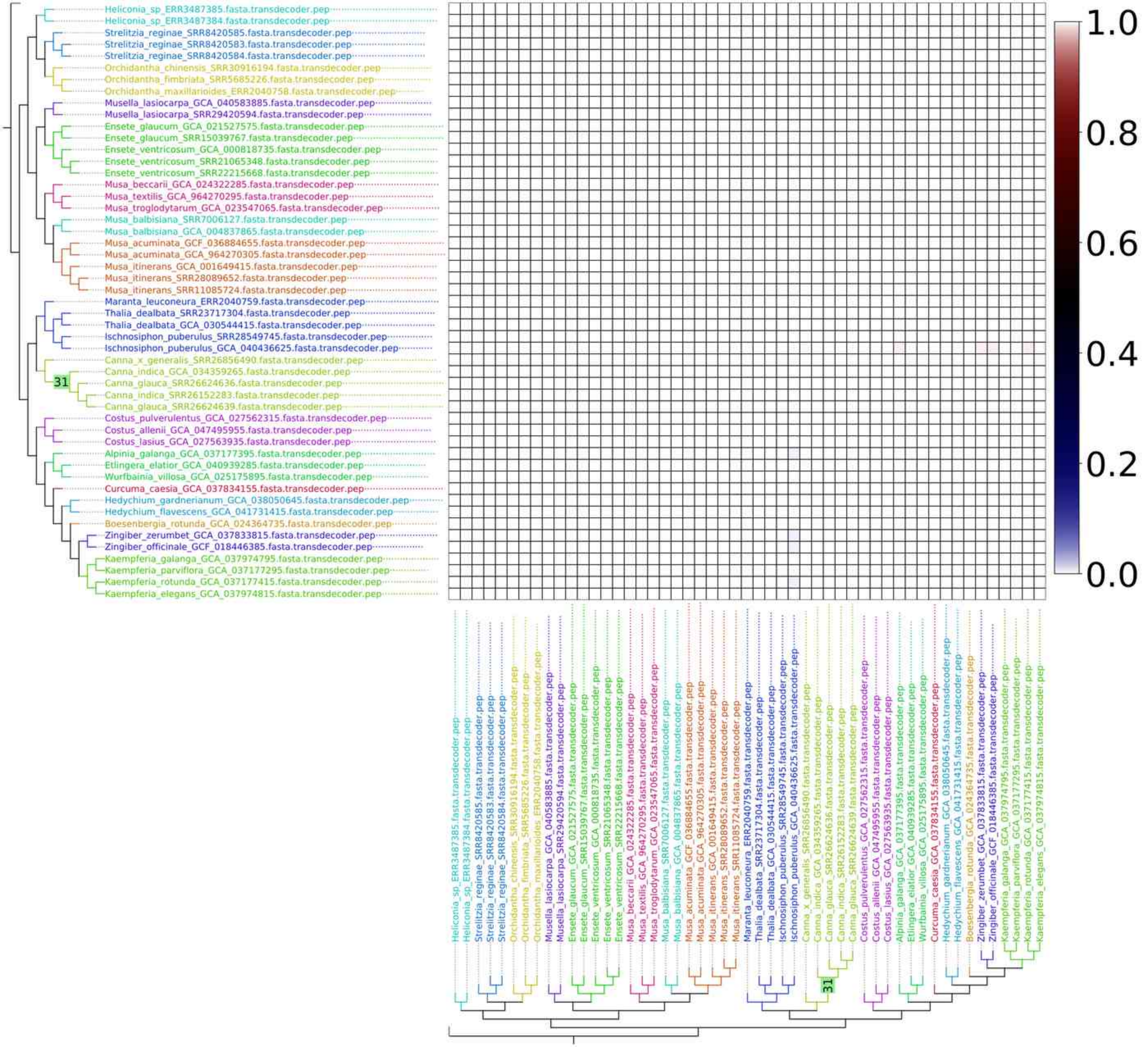
Heliconia_sp_ERR3487385.fasta.transdecoder.pep
Heliconia_sp_ERR3487384.fasta.transdecoder.pep
Strelitzia_reginae_SRR8420585.fasta.transdecoder.pep
Strelitzia_reginae_SRR8420583.fasta.transdecoder.pep
Strelitzia_reginae_SRR8420584.fasta.transdecoder.pep
Orchidantha_chinensis_SRR30916194.fasta.transdecoder.pep
Orchidantha_fimbriata_SRR5685226.fasta.transdecoder.pep
Orchidantha_maxillarioides_ERR2040758.fasta.transdecoder.pep
Musella_lasiocarpa_GCA_040583885.fasta.transdecoder.pep
Musella_lasiocarpa_SRR29420594.fasta.transdecoder.pep
Ensete_glaucum_GCA_021527575.fasta.transdecoder.pep
Ensete_glaucum_SRR15039767.fasta.transdecoder.pep
Ensete_ventricosum_GCA_000818735.fasta.transdecoder.pep
Ensete_ventricosum_SRR21065348.fasta.transdecoder.pep
Ensete_ventricosum_SRR22215668.fasta.transdecoder.pep
Musa_beccarii_GCA_024322285.fasta.transdecoder.pep
Musa_textilis_GCA_964270295.fasta.transdecoder.pep
Musa_troglodytarum_GCA_023547065.fasta.transdecoder.pep
Musa_balbisiana_SRR7006127.fasta.transdecoder.pep
Musa_balbisiana_GCA_004837865.fasta.transdecoder.pep
Musa_acuminata_GCF_036884655.fasta.transdecoder.pep
Musa_acuminata_GCA_964270305.fasta.transdecoder.pep
Musa_itinerans_GCA_001649415.fasta.transdecoder.pep
Musa_itinerans_SRR28089652.fasta.transdecoder.pep
Musa_itinerans_SRR11085724.fasta.transdecoder.pep
Maranta_leuconeura_ERR2040759.fasta.transdecoder.pep
Thalia_dealbata_SRR23717304.fasta.transdecoder.pep
Thalia_dealbata_GCA_030544415.fasta.transdecoder.pep
Ischnosiphon_puberulus_SRR28549745.fasta.transdecoder.pep
Ischnosiphon_puberulus_GCA_040436625.fasta.transdecoder.pep
Canna_x_generalis_SRR26856490.fasta.transdecoder.pep
Canna_indica_GCA_034359265.fasta.transdecoder.pep
Canna_glauca_SRR26624636.fasta.transdecoder.pep
Canna_indica_SRR26152283.fasta.transdecoder.pep
Canna_glauca_SRR26624639.fasta.transdecoder.pep
Costus_pulverulentus_GCA_027562315.fasta.transdecoder.pep
Costus_allenii_GCA_047495955.fasta.transdecoder.pep
Costus_lasius_GCA_027563935.fasta.transdecoder.pep
Alpinia_galanga_GCA_037177395.fasta.transdecoder.pep
Etlingera_elatior_GCA_040939285.fasta.transdecoder.pep
Wurfbainia_villosa_GCA_025175895.fasta.transdecoder.pep
Curcuma_caesia_GCA_037834155.fasta.transdecoder.pep
Hedychium_gardnerianum_GCA_038050645.fasta.transdecoder.pep
Hedychium_flavescens_GCA_041731415.fasta.transdecoder.pep
Boesenbergia_rotunda_GCA_024364735.fasta.transdecoder.pep
Zingiber_zerumbet_GCA_037833815.fasta.transdecoder.pep
Zingiber_officinale_GCF_018446385.fasta.transdecoder.pep
Kaempferia_galanga_GCA_037974795.fasta.transdecoder.pep
Kaempferia_parviflora_GCA_037177295.fasta.transdecoder.pep
Kaempferia_rotunda_GCA_037177415.fasta.transdecoder.pep
Kaempferia_elegans_GCA_037974815.fasta.transdecoder.pep
31
1.0
0.8
0.6
0.4
0.2
0.0

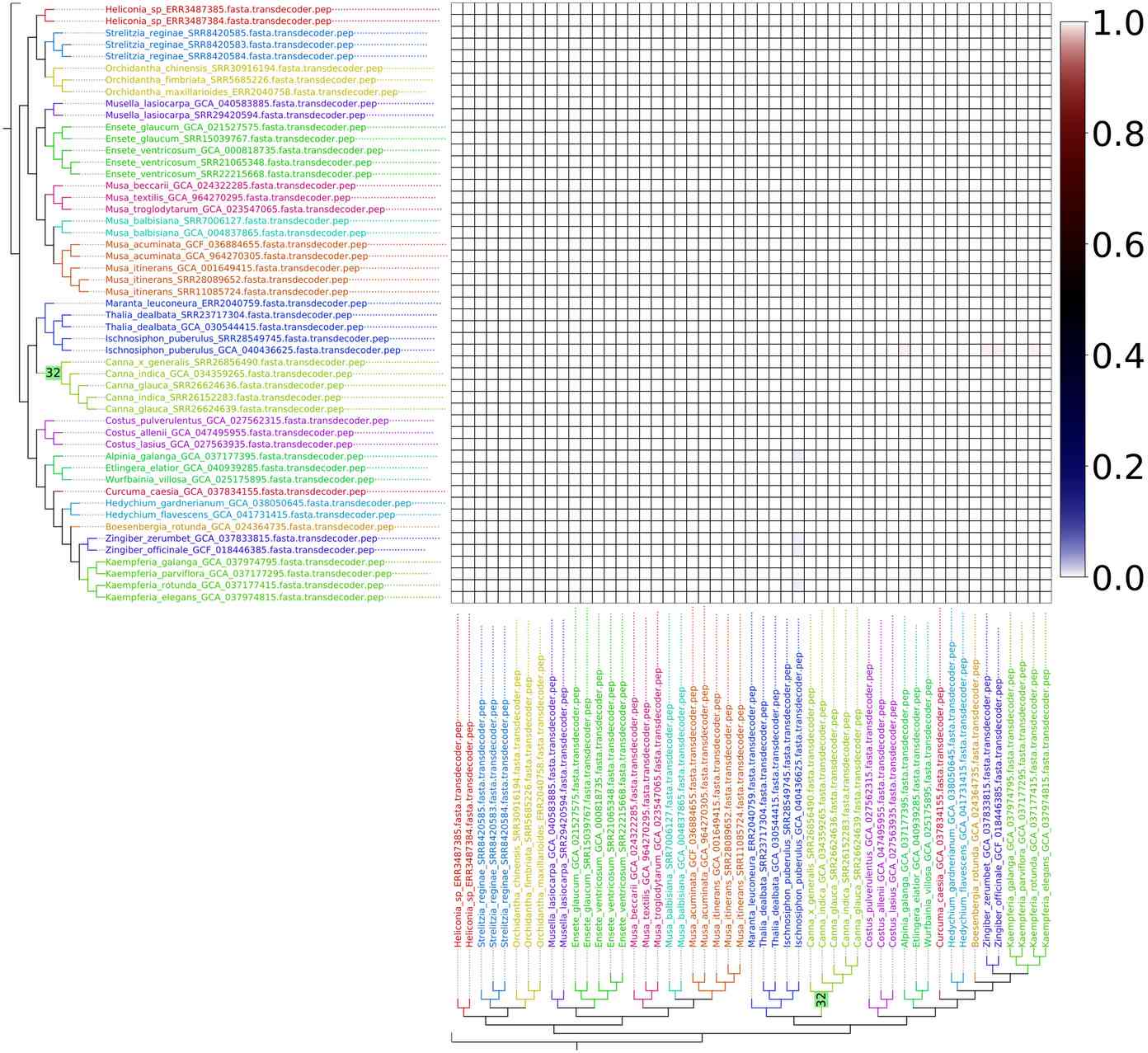
Heliconia_sp_ERR3487385.fasta.transdecoder.pep
Heliconia_sp_ERR3487384.fasta.transdecoder.pep
Strelitzia_reginae_SRR8420585.fasta.transdecoder.pep
Strelitzia_reginae_SRR8420583.fasta.transdecoder.pep
Strelitzia_reginae_SRR8420584.fasta.transdecoder.pep
Orchidantha_chinensis_SRR30916194.fasta.transdecoder.pep
Orchidantha_fimbriata_SRR5685226.fasta.transdecoder.pep
Orchidantha_maxillarioides_ERR2040758.fasta.transdecoder.pep
Musella_lasiocarpa_GCA_040583885.fasta.transdecoder.pep
Musella_lasiocarpa_SRR29420594.fasta.transdecoder.pep
Ensete_glaucum_GCA_021527575.fasta.transdecoder.pep
Ensete_glaucum_SRR15039767.fasta.transdecoder.pep
Ensete_ventricosum_GCA_000818735.fasta.transdecoder.pep
Ensete_ventricosum_SRR21065348.fasta.transdecoder.pep
Ensete_ventricosum_SRR22215668.fasta.transdecoder.pep
Musa_beccarii_GCA_024322285.fasta.transdecoder.pep
Musa_textilis_GCA_964270295.fasta.transdecoder.pep
Musa_troglodytarum_GCA_023547065.fasta.transdecoder.pep
Musa_balbisiana_SRR7006127.fasta.transdecoder.pep
Musa_balbisiana_GCA_004837865.fasta.transdecoder.pep
Musa_acuminata_GCF_036884655.fasta.transdecoder.pep
Musa_acuminata_GCA_964270305.fasta.transdecoder.pep
Musa_itinerans_GCA_001649415.fasta.transdecoder.pep
Musa_itinerans_SRR28089652.fasta.transdecoder.pep
Musa_itinerans_SRR11085724.fasta.transdecoder.pep
Maranta_leuconeura_ERR2040759.fasta.transdecoder.pep
Thalia_dealbata_SRR23717304.fasta.transdecoder.pep
Thalia_dealbata_GCA_030544415.fasta.transdecoder.pep
Ischnosiphon_puberulus_SRR28549745.fasta.transdecoder.pep
Ischnosiphon_puberulus_GCA_040436625.fasta.transdecoder.pep
Canna_x_generalis_SRR26856490.fasta.transdecoder.pep
Canna_indica_GCA_034359265.fasta.transdecoder.pep
Canna_glauca_SRR26624636.fasta.transdecoder.pep
Canna_indica_SRR26152283.fasta.transdecoder.pep
Canna_glauca_SRR26624639.fasta.transdecoder.pep
Costus_pulverulentus_GCA_027562315.fasta.transdecoder.pep
Costus_allenii_GCA_047495955.fasta.transdecoder.pep
Costus_lasius_GCA_027563935.fasta.transdecoder.pep
Alpinia_galanga_GCA_037177395.fasta.transdecoder.pep
Etlingera_elatior_GCA_040939285.fasta.transdecoder.pep
Wurfbainia_villosa_GCA_025175895.fasta.transdecoder.pep
Curcuma_caesia_GCA_037834155.fasta.transdecoder.pep
Hedychium_gardnerianum_GCA_038050645.fasta.transdecoder.pep
Hedychium_flavescens_GCA_041731415.fasta.transdecoder.pep
Boesenbergia_rotunda_GCA_024364735.fasta.transdecoder.pep
Zingiber_zerumbet_GCA_037833815.fasta.transdecoder.pep
Zingiber_officinale_GCF_018446385.fasta.transdecoder.pep
Kaempferia_galanga_GCA_037974795.fasta.transdecoder.pep
Kaempferia_parviflora_GCA_037177295.fasta.transdecoder.pep
Kaempferia_rotunda_GCA_037177415.fasta.transdecoder.pep
Kaempferia_elegans_GCA_037974815.fasta.transdecoder.pep
32
1.0
0.8
0.6
0.4
0.2
0.0

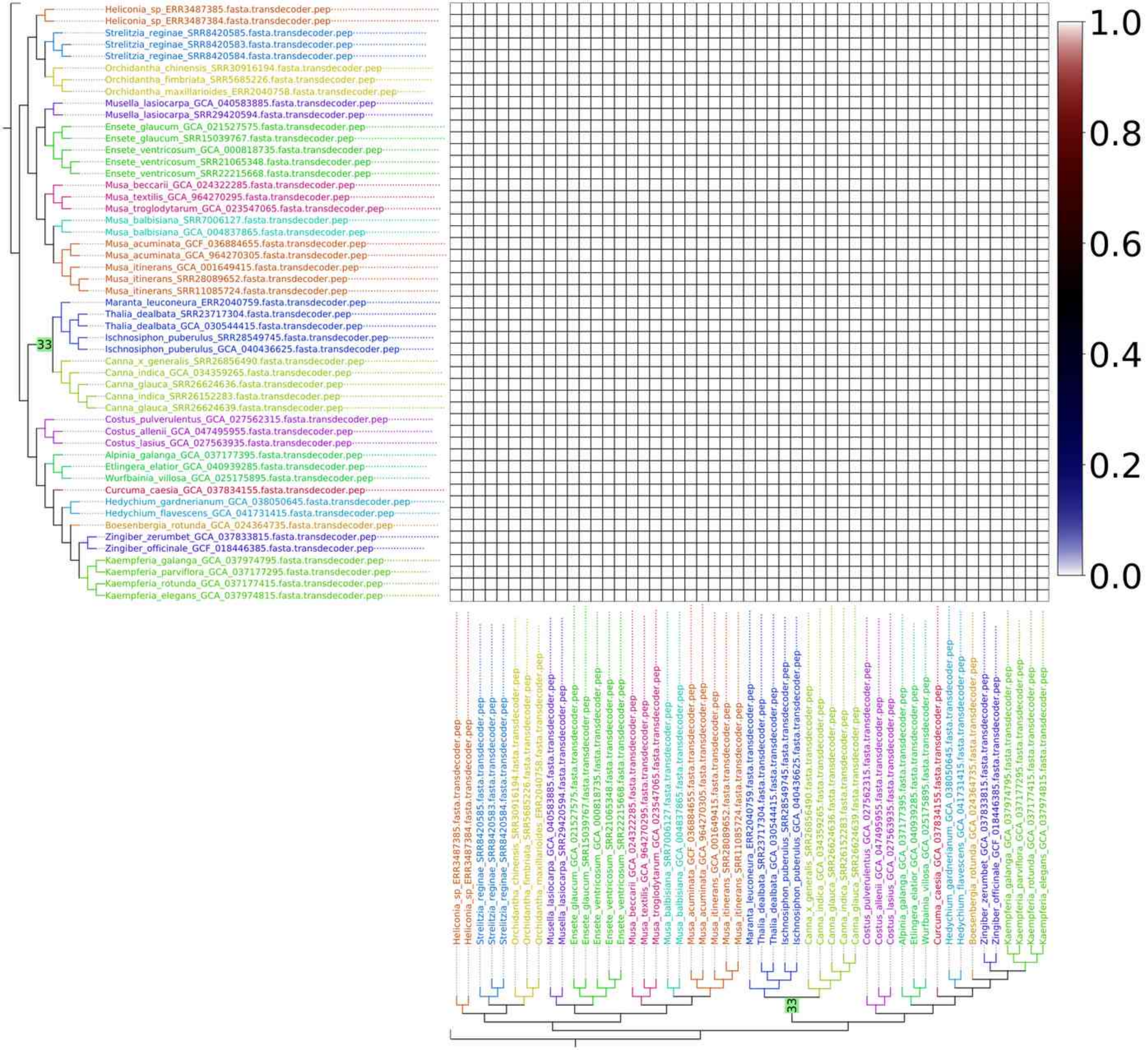
Heliconia_sp_ERR3487385.fasta.transdecoder.pep
Heliconia_sp_ERR3487384.fasta.transdecoder.pep
Strelitzia_reginae_SRR8420585.fasta.transdecoder.pep
Strelitzia_reginae_SRR8420583.fasta.transdecoder.pep
Strelitzia_reginae_SRR8420584.fasta.transdecoder.pep
Orchidantha_chinensis_SRR30916194.fasta.transdecoder.pep
Orchidantha_fimbriata_SRR5685226.fasta.transdecoder.pep
Orchidantha_maxillarioides_ERR2040758.fasta.transdecoder.pep
Musella_lasiocarpa_GCA_040583885.fasta.transdecoder.pep
Musella_lasiocarpa_SRR29420594.fasta.transdecoder.pep
Ensete_glaucum_GCA_021527575.fasta.transdecoder.pep
Ensete_glaucum_SRR15039767.fasta.transdecoder.pep
Ensete_ventricosum_GCA_000818735.fasta.transdecoder.pep
Ensete_ventricosum_SRR21065348.fasta.transdecoder.pep
Ensete_ventricosum_SRR22215668.fasta.transdecoder.pep
Musa_beccarii_GCA_024322285.fasta.transdecoder.pep
Musa_textilis_GCA_964270295.fasta.transdecoder.pep
Musa_troglodytarum_GCA_023547065.fasta.transdecoder.pep
Musa_balbisiana_SRR7006127.fasta.transdecoder.pep
Musa_balbisiana_GCA_004837865.fasta.transdecoder.pep
Musa_acuminata_GCF_036884655.fasta.transdecoder.pep
Musa_acuminata_GCA_964270305.fasta.transdecoder.pep
Musa_itinerans_GCA_001649415.fasta.transdecoder.pep
Musa_itinerans_SRR28089652.fasta.transdecoder.pep
Musa_itinerans_SRR11085724.fasta.transdecoder.pep
Maranta_leuconeura_ERR2040759.fasta.transdecoder.pep
Thalia_dealbata_SRR23717304.fasta.transdecoder.pep
Thalia_dealbata_GCA_030544415.fasta.transdecoder.pep
Ischnosiphon_puberulus_SRR28549745.fasta.transdecoder.pep
Ischnosiphon_puberulus_GCA_040436625.fasta.transdecoder.pep
Canna_x_generalis_SRR26856490.fasta.transdecoder.pep
Canna_indica_GCA_034359265.fasta.transdecoder.pep
Canna_glauca_SRR26624636.fasta.transdecoder.pep
Canna_indica_SRR26152283.fasta.transdecoder.pep
Canna_glauca_SRR26624639.fasta.transdecoder.pep
Costus_pulverulentus_GCA_027562315.fasta.transdecoder.pep
Costus_allenii_GCA_047495955.fasta.transdecoder.pep
Costus_lasius_GCA_027563935.fasta.transdecoder.pep
Alpinia_galanga_GCA_037177395.fasta.transdecoder.pep
Etlingera_elatior_GCA_040939285.fasta.transdecoder.pep
Wurfbainia_villosa_GCA_025175895.fasta.transdecoder.pep
Curcuma_caesia_GCA_037834155.fasta.transdecoder.pep
Hedychium_gardnerianum_GCA_038050645.fasta.transdecoder.pep
Hedychium_flavescens_GCA_041731415.fasta.transdecoder.pep
Boesenbergia_rotunda_GCA_024364735.fasta.transdecoder.pep
Zingiber_zerumbet_GCA_037833815.fasta.transdecoder.pep
Zingiber_officinale_GCF_018446385.fasta.transdecoder.pep
Kaempferia_galanga_GCA_037974795.fasta.transdecoder.pep
Kaempferia_parviflora_GCA_037177295.fasta.transdecoder.pep
Kaempferia_rotunda_GCA_037177415.fasta.transdecoder.pep
Kaempferia_elegans_GCA_037974815.fasta.transdecoder.pep
33
1.0
0.8
0.6
0.4
0.2
0.0

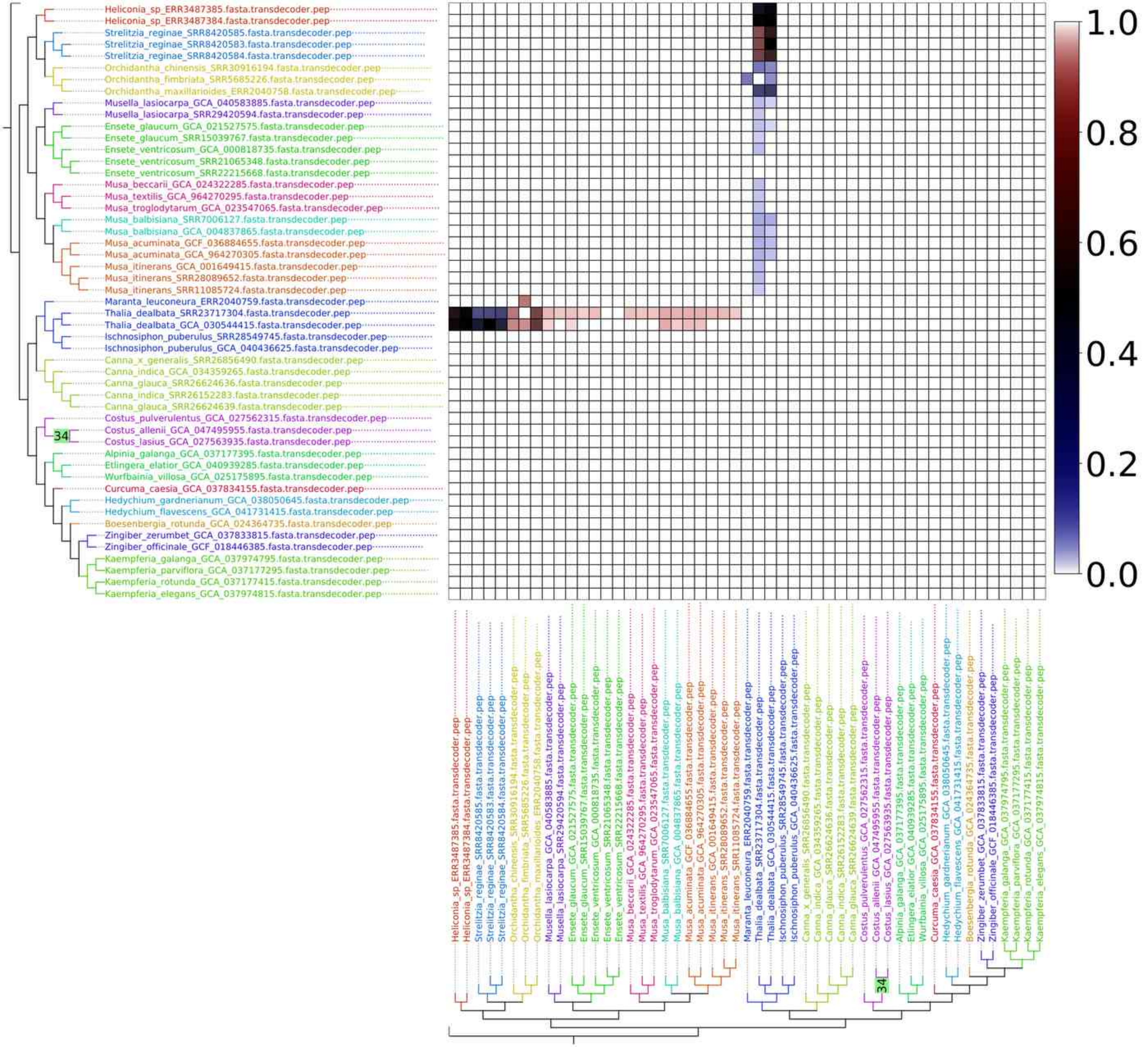
Heliconia_sp_ERR3487385.fasta.transdecoder.pep
Heliconia_sp_ERR3487384.fasta.transdecoder.pep
Strelitzia_reginae_SRR8420585.fasta.transdecoder.pep
Strelitzia_reginae_SRR8420583.fasta.transdecoder.pep
Strelitzia_reginae_SRR8420584.fasta.transdecoder.pep
Orchidantha_chinensis_SRR30916194.fasta.transdecoder.pep
Orchidantha_fimbriata_SRR5685226.fasta.transdecoder.pep
Orchidantha_maxillarioides_ERR2040758.fasta.transdecoder.pep
Musella_lasiocarpa_GCA_040583885.fasta.transdecoder.pep
Musella_lasiocarpa_SRR29420594.fasta.transdecoder.pep
Ensete_glaucum_GCA_021527575.fasta.transdecoder.pep
Ensete_glaucum_SRR15039767.fasta.transdecoder.pep
Ensete_ventricosum_GCA_000818735.fasta.transdecoder.pep
Ensete_ventricosum_SRR21065348.fasta.transdecoder.pep
Ensete_ventricosum_SRR22215668.fasta.transdecoder.pep
Musa_beccarii_GCA_024322285.fasta.transdecoder.pep
Musa_textilis_GCA_964270295.fasta.transdecoder.pep
Musa_troglodytarum_GCA_023547065.fasta.transdecoder.pep
Musa_balbisiana_SRR7006127.fasta.transdecoder.pep
Musa_balbisiana_GCA_004837865.fasta.transdecoder.pep
Musa_acuminata_GCF_036884655.fasta.transdecoder.pep
Musa_acuminata_GCA_964270305.fasta.transdecoder.pep
Musa_itinerans_GCA_001649415.fasta.transdecoder.pep
Musa_itinerans_SRR28089652.fasta.transdecoder.pep
Musa_itinerans_SRR11085724.fasta.transdecoder.pep
Maranta_leuconeura_ERR2040759.fasta.transdecoder.pep
Thalia_dealbata_SRR23717304.fasta.transdecoder.pep
Thalia_dealbata_GCA_030544415.fasta.transdecoder.pep
Ischnosiphon_puberulus_SRR28549745.fasta.transdecoder.pep
Ischnosiphon_puberulus_GCA_040436625.fasta.transdecoder.pep
Canna_x_generalis_SRR26856490.fasta.transdecoder.pep
Canna_indica_GCA_034359265.fasta.transdecoder.pep
Canna_glauca_SRR26624636.fasta.transdecoder.pep
Canna_indica_SRR26152283.fasta.transdecoder.pep
Canna_glauca_SRR26624639.fasta.transdecoder.pep
Costus_pulverulentus_GCA_027562315.fasta.transdecoder.pep
Costus_allenii_GCA_047495955.fasta.transdecoder.pep
34
Costus_lasius_GCA_027563935.fasta.transdecoder.pep
Alpinia_galanga_GCA_037177395.fasta.transdecoder.pep
Etlingera_elatior_GCA_040939285.fasta.transdecoder.pep
Wurfbainia_villosa_GCA_025175895.fasta.transdecoder.pep
Curcuma_caesia_GCA_037834155.fasta.transdecoder.pep
Hedychium_gardnerianum_GCA_038050645.fasta.transdecoder.pep
Hedychium_flavescens_GCA_041731415.fasta.transdecoder.pep
Boesenbergia_rotunda_GCA_024364735.fasta.transdecoder.pep
Zingiber_zerumbet_GCA_037833815.fasta.transdecoder.pep
Zingiber_officinale_GCF_018446385.fasta.transdecoder.pep
Kaempferia_galanga_GCA_037974795.fasta.transdecoder.pep
Kaempferia_parviflora_GCA_037177295.fasta.transdecoder.pep
Kaempferia_rotunda_GCA_037177415.fasta.transdecoder.pep
Kaempferia_elegans_GCA_037974815.fasta.transdecoder.pep
1.0
0.8
0.6
0.4
0.2
0.0

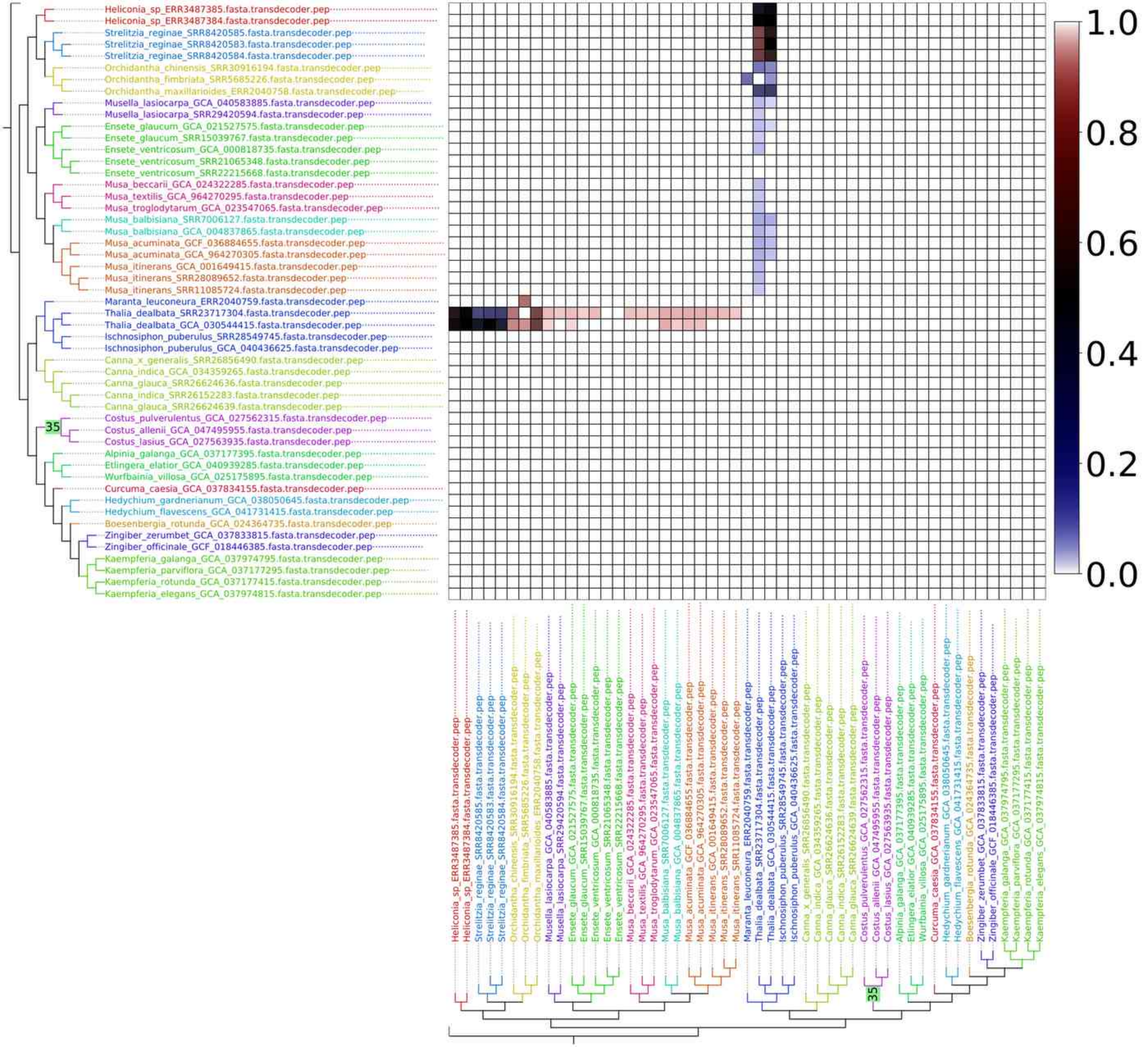
Heliconia_sp_ERR3487385.fasta.transdecoder.pep
Heliconia_sp_ERR3487384.fasta.transdecoder.pep
Strelitzia_reginae_SRR8420585.fasta.transdecoder.pep
Strelitzia_reginae_SRR8420583.fasta.transdecoder.pep
Strelitzia_reginae_SRR8420584.fasta.transdecoder.pep
Orchidantha_chinensis_SRR30916194.fasta.transdecoder.pep
Orchidantha_fimbriata_SRR5685226.fasta.transdecoder.pep
Orchidantha_maxillarioides_ERR2040758.fasta.transdecoder.pep
Musella_lasiocarpa_GCA_040583885.fasta.transdecoder.pep
Musella_lasiocarpa_SRR29420594.fasta.transdecoder.pep
Ensete_glaucum_GCA_021527575.fasta.transdecoder.pep
Ensete_glaucum_SRR15039767.fasta.transdecoder.pep
Ensete_ventricosum_GCA_000818735.fasta.transdecoder.pep
Ensete_ventricosum_SRR21065348.fasta.transdecoder.pep
Ensete_ventricosum_SRR22215668.fasta.transdecoder.pep
Musa_beccarii_GCA_024322285.fasta.transdecoder.pep
Musa_textilis_GCA_964270295.fasta.transdecoder.pep
Musa_troglodytarum_GCA_023547065.fasta.transdecoder.pep
Musa_balbisiana_SRR7006127.fasta.transdecoder.pep
Musa_balbisiana_GCA_004837865.fasta.transdecoder.pep
Musa_acuminata_GCF_036884655.fasta.transdecoder.pep
Musa_acuminata_GCA_964270305.fasta.transdecoder.pep
Musa_itinerans_GCA_001649415.fasta.transdecoder.pep
Musa_itinerans_SRR28089652.fasta.transdecoder.pep
Musa_itinerans_SRR11085724.fasta.transdecoder.pep
Maranta_leuconeura_ERR2040759.fasta.transdecoder.pep
Thalia_dealbata_SRR23717304.fasta.transdecoder.pep
Thalia_dealbata_GCA_030544415.fasta.transdecoder.pep
Ischnosiphon_puberulus_SRR28549745.fasta.transdecoder.pep
Ischnosiphon_puberulus_GCA_040436625.fasta.transdecoder.pep
Canna_x_generalis_SRR26856490.fasta.transdecoder.pep
Canna_indica_GCA_034359265.fasta.transdecoder.pep
Canna_glauca_SRR26624636.fasta.transdecoder.pep
Canna_indica_SRR26152283.fasta.transdecoder.pep
Canna_glauca_SRR26624639.fasta.transdecoder.pep
Costus_pulverulentus_GCA_027562315.fasta.transdecoder.pep
Costus_allenii_GCA_047495955.fasta.transdecoder.pep
Costus_lasius_GCA_027563935.fasta.transdecoder.pep
Alpinia_galanga_GCA_037177395.fasta.transdecoder.pep
Etlingera_elatior_GCA_040939285.fasta.transdecoder.pep
Wurfbainia_villosa_GCA_025175895.fasta.transdecoder.pep
Curcuma_caesia_GCA_037834155.fasta.transdecoder.pep
Hedychium_gardnerianum_GCA_038050645.fasta.transdecoder.pep
Hedychium_flavescens_GCA_041731415.fasta.transdecoder.pep
Boesenbergia_rotunda_GCA_024364735.fasta.transdecoder.pep
Zingiber_zerumbet_GCA_037833815.fasta.transdecoder.pep
Zingiber_officinale_GCF_018446385.fasta.transdecoder.pep
Kaempferia_galanga_GCA_037974795.fasta.transdecoder.pep
Kaempferia_parviflora_GCA_037177295.fasta.transdecoder.pep
Kaempferia_rotunda_GCA_037177415.fasta.transdecoder.pep
Kaempferia_elegans_GCA_037974815.fasta.transdecoder.pep
35
1.0
0.8
0.6
0.4
0.2
0.0

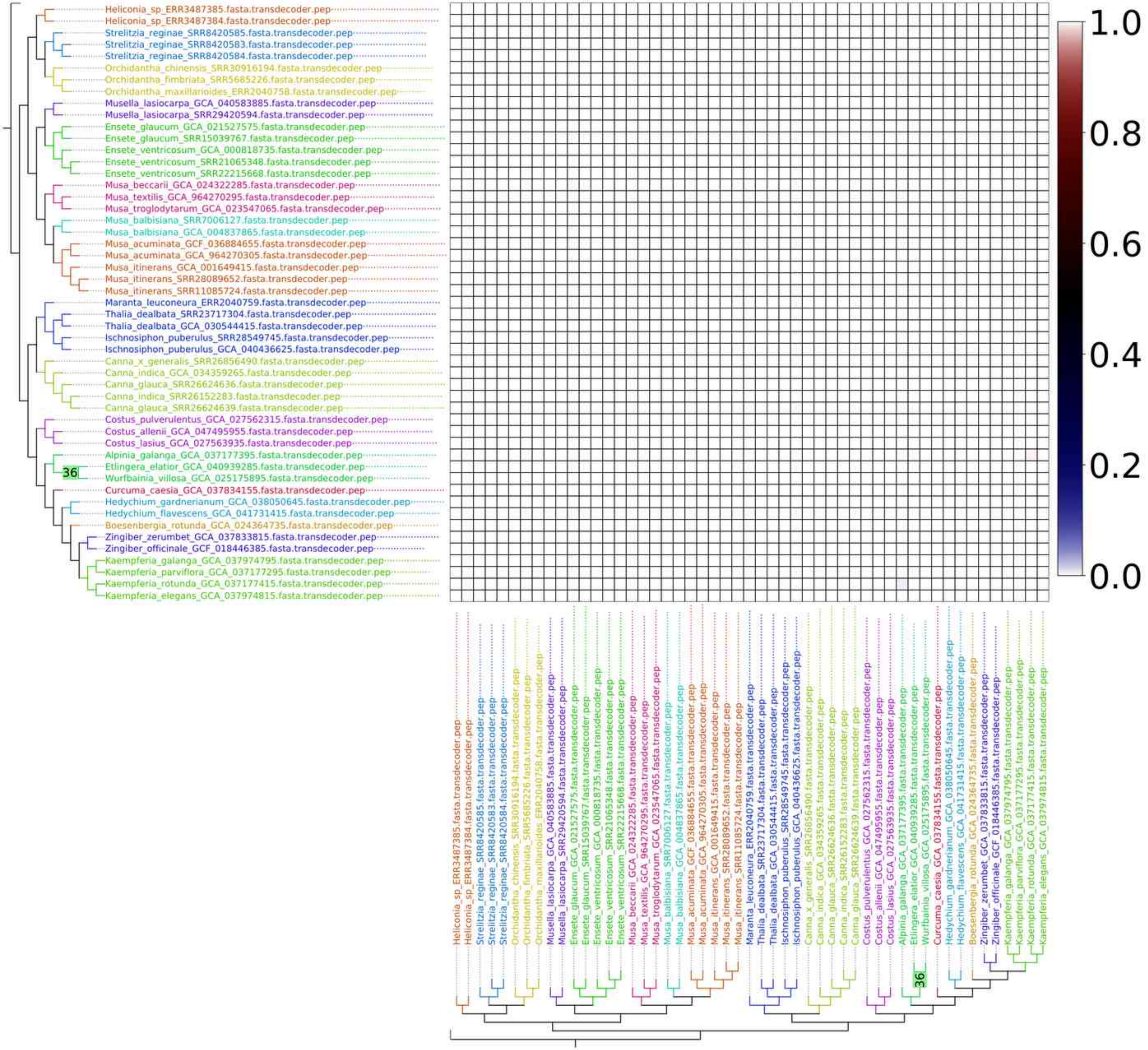
Heliconia_sp_ERR3487385.fasta.transdecoder.pep
Heliconia_sp_ERR3487384.fasta.transdecoder.pep
Strelitzia_reginae_SRR8420585.fasta.transdecoder.pep
Strelitzia_reginae_SRR8420583.fasta.transdecoder.pep
Strelitzia_reginae_SRR8420584.fasta.transdecoder.pep
Orchidantha_chinensis_SRR30916194.fasta.transdecoder.pep
Orchidantha_fimbriata_SRR5685226.fasta.transdecoder.pep
Orchidantha_maxillarioides_ERR2040758.fasta.transdecoder.pep
Musella_lasiocarpa_GCA_040583885.fasta.transdecoder.pep
Musella_lasiocarpa_SRR29420594.fasta.transdecoder.pep
Ensete_glaucum_GCA_021527575.fasta.transdecoder.pep
Ensete_glaucum_SRR15039767.fasta.transdecoder.pep
Ensete_ventricosum_GCA_000818735.fasta.transdecoder.pep
Ensete_ventricosum_SRR21065348.fasta.transdecoder.pep
Ensete_ventricosum_SRR22215668.fasta.transdecoder.pep
Musa_beccarii_GCA_024322285.fasta.transdecoder.pep
Musa_textilis_GCA_964270295.fasta.transdecoder.pep
Musa_troglodytarum_GCA_023547065.fasta.transdecoder.pep
Musa_balbisiana_SRR7006127.fasta.transdecoder.pep
Musa_balbisiana_GCA_004837865.fasta.transdecoder.pep
Musa_acuminata_GCF_036884655.fasta.transdecoder.pep
Musa_acuminata_GCA_964270305.fasta.transdecoder.pep
Musa_itinerans_GCA_001649415.fasta.transdecoder.pep
Musa_itinerans_SRR28089652.fasta.transdecoder.pep
Musa_itinerans_SRR11085724.fasta.transdecoder.pep
Maranta_leuconeura_ERR2040759.fasta.transdecoder.pep
Thalia_dealbata_SRR23717304.fasta.transdecoder.pep
Thalia_dealbata_GCA_030544415.fasta.transdecoder.pep
Ischnosiphon_puberulus_SRR28549745.fasta.transdecoder.pep
Ischnosiphon_puberulus_GCA_040436625.fasta.transdecoder.pep
Canna_x_generalis_SRR26856490.fasta.transdecoder.pep
Canna_indica_GCA_034359265.fasta.transdecoder.pep
Canna_glauca_SRR26624636.fasta.transdecoder.pep
Canna_indica_SRR26152283.fasta.transdecoder.pep
Canna_glauca_SRR26624639.fasta.transdecoder.pep
Costus_pulverulentus_GCA_027562315.fasta.transdecoder.pep
Costus_allenii_GCA_047495955.fasta.transdecoder.pep
Costus_lasius_GCA_027563935.fasta.transdecoder.pep
Alpinia_galanga_GCA_037177395.fasta.transdecoder.pep
Etlingera_elatior_GCA_040939285.fasta.transdecoder.pep
Wurfbainia_villosa_GCA_025175895.fasta.transdecoder.pep
Curcuma_caesia_GCA_037834155.fasta.transdecoder.pep
Hedychium_gardnerianum_GCA_038050645.fasta.transdecoder.pep
Hedychium_flavescens_GCA_041731415.fasta.transdecoder.pep
Boesenbergia_rotunda_GCA_024364735.fasta.transdecoder.pep
Zingiber_zerumbet_GCA_037833815.fasta.transdecoder.pep
Zingiber_officinale_GCF_018446385.fasta.transdecoder.pep
Kaempferia_galanga_GCA_037974795.fasta.transdecoder.pep
Kaempferia_parviflora_GCA_037177295.fasta.transdecoder.pep
Kaempferia_rotunda_GCA_037177415.fasta.transdecoder.pep
Kaempferia_elegans_GCA_037974815.fasta.transdecoder.pep
36
1.0
0.8
0.6
0.4
0.2
0.0

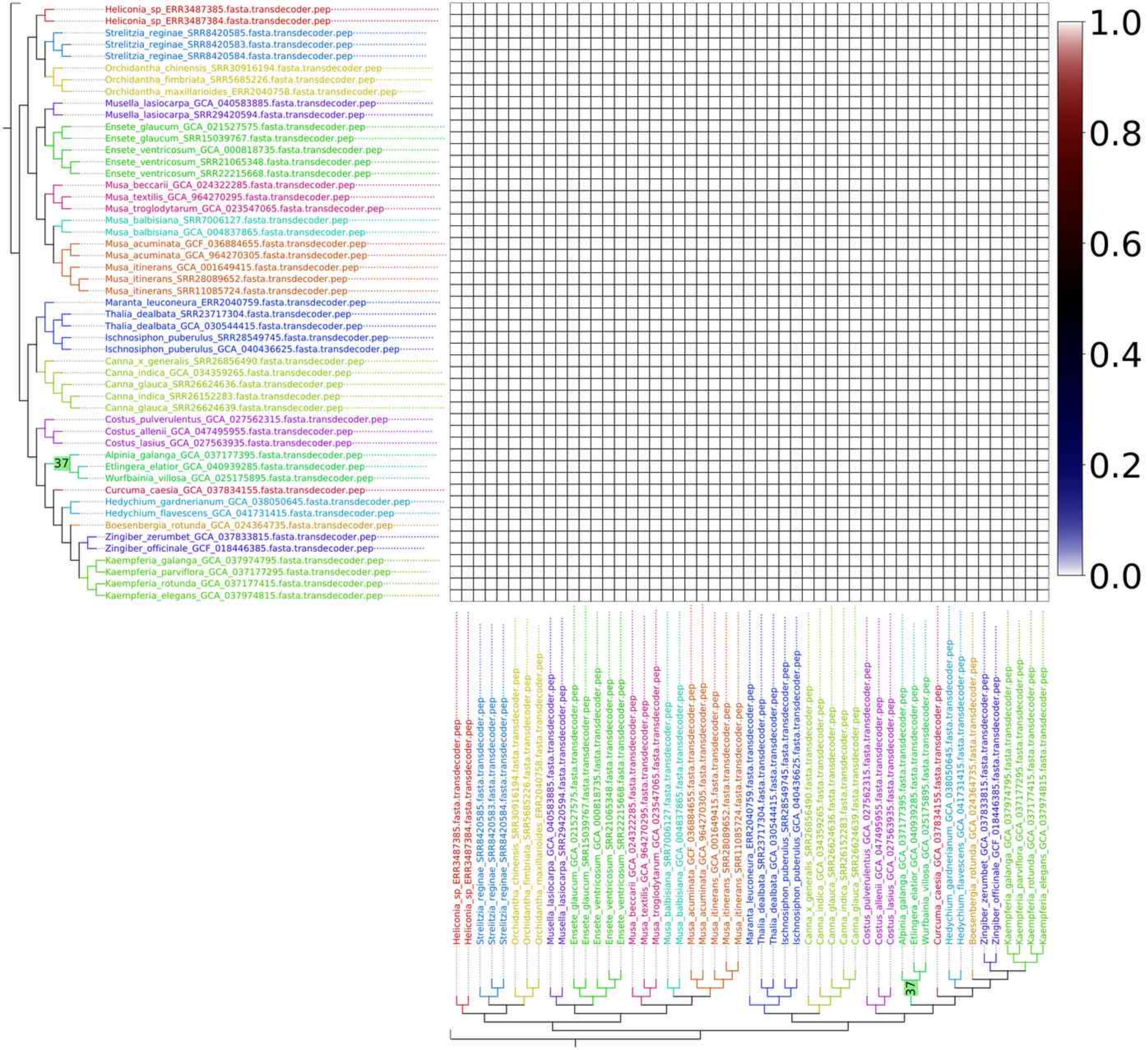
Heliconia_sp_ERR3487385.fasta.transdecoder.pep
Heliconia_sp_ERR3487384.fasta.transdecoder.pep
Strelitzia_reginae_SRR8420585.fasta.transdecoder.pep
Strelitzia_reginae_SRR8420583.fasta.transdecoder.pep
Strelitzia_reginae_SRR8420584.fasta.transdecoder.pep
Orchidantha_chinensis_SRR30916194.fasta.transdecoder.pep
Orchidantha_fimbriata_SRR5685226.fasta.transdecoder.pep
Orchidantha_maxillarioides_ERR2040758.fasta.transdecoder.pep
Musella_lasiocarpa_GCA_040583885.fasta.transdecoder.pep
Musella_lasiocarpa_SRR29420594.fasta.transdecoder.pep
Ensete_glaucum_GCA_021527575.fasta.transdecoder.pep
Ensete_glaucum_SRR15039767.fasta.transdecoder.pep
Ensete_ventricosum_GCA_000818735.fasta.transdecoder.pep
Ensete_ventricosum_SRR21065348.fasta.transdecoder.pep
Ensete_ventricosum_SRR22215668.fasta.transdecoder.pep
Musa_beccarii_GCA_024322285.fasta.transdecoder.pep
Musa_textilis_GCA_964270295.fasta.transdecoder.pep
Musa_troglodytarum_GCA_023547065.fasta.transdecoder.pep
Musa_balbisiana_SRR7006127.fasta.transdecoder.pep
Musa_balbisiana_GCA_004837865.fasta.transdecoder.pep
Musa_acuminata_GCF_036884655.fasta.transdecoder.pep
Musa_acuminata_GCA_964270305.fasta.transdecoder.pep
Musa_itinerans_GCA_001649415.fasta.transdecoder.pep
Musa_itinerans_SRR28089652.fasta.transdecoder.pep
Musa_itinerans_SRR11085724.fasta.transdecoder.pep
Maranta_leuconeura_ERR2040759.fasta.transdecoder.pep
Thalia_dealbata_SRR23717304.fasta.transdecoder.pep
Thalia_dealbata_GCA_030544415.fasta.transdecoder.pep
Ischnosiphon_puberulus_SRR28549745.fasta.transdecoder.pep
Ischnosiphon_puberulus_GCA_040436625.fasta.transdecoder.pep
Canna_x_generalis_SRR26856490.fasta.transdecoder.pep
Canna_indica_GCA_034359265.fasta.transdecoder.pep
Canna_glauca_SRR26624636.fasta.transdecoder.pep
Canna_indica_SRR26152283.fasta.transdecoder.pep
Canna_glauca_SRR26624639.fasta.transdecoder.pep
Costus_pulverulentus_GCA_027562315.fasta.transdecoder.pep
Costus_allenii_GCA_047495955.fasta.transdecoder.pep
Costus_lasius_GCA_027563935.fasta.transdecoder.pep
37
Alpinia_galanga_GCA_037177395.fasta.transdecoder.pep
Etlingera_elatior_GCA_040939285.fasta.transdecoder.pep
Wurfbainia_villosa_GCA_025175895.fasta.transdecoder.pep
Curcuma_caesia_GCA_037834155.fasta.transdecoder.pep
Hedychium_gardnerianum_GCA_038050645.fasta.transdecoder.pep
Hedychium_flavescens_GCA_041731415.fasta.transdecoder.pep
Boesenbergia_rotunda_GCA_024364735.fasta.transdecoder.pep
Zingiber_zerumbet_GCA_037833815.fasta.transdecoder.pep
Zingiber_officinale_GCF_018446385.fasta.transdecoder.pep
Kaempferia_galanga_GCA_037974795.fasta.transdecoder.pep
Kaempferia_parviflora_GCA_037177295.fasta.transdecoder.pep
Kaempferia_rotunda_GCA_037177415.fasta.transdecoder.pep
Kaempferia_elegans_GCA_037974815.fasta.transdecoder.pep
1.0
0.8
0.6
0.4
0.2
0.0

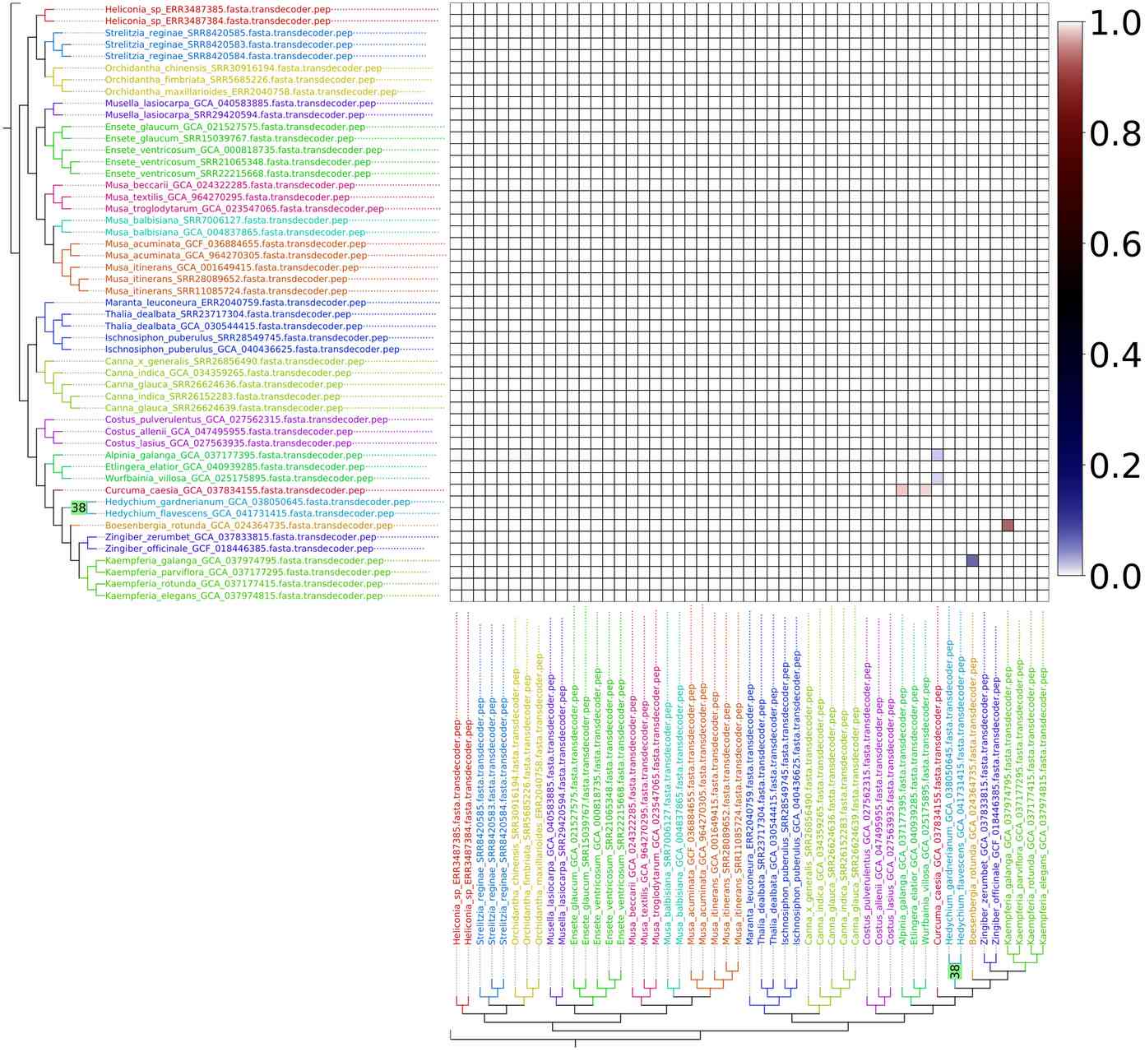
Heliconia_sp_ERR3487385.fasta.transdecoder.pep
Heliconia_sp_ERR3487384.fasta.transdecoder.pep
Strelitzia_reginae_SRR8420585.fasta.transdecoder.pep
Strelitzia_reginae_SRR8420583.fasta.transdecoder.pep
Strelitzia_reginae_SRR8420584.fasta.transdecoder.pep
Orchidantha_chinensis_SRR30916194.fasta.transdecoder.pep
Orchidantha_fimbriata_SRR5685226.fasta.transdecoder.pep
Orchidantha_maxillarioides_ERR2040758.fasta.transdecoder.pep
Musella_lasiocarpa_GCA_040583885.fasta.transdecoder.pep
Musella_lasiocarpa_SRR29420594.fasta.transdecoder.pep
Ensete_glaucum_GCA_021527575.fasta.transdecoder.pep
Ensete_glaucum_SRR15039767.fasta.transdecoder.pep
Ensete_ventricosum_GCA_000818735.fasta.transdecoder.pep
Ensete_ventricosum_SRR21065348.fasta.transdecoder.pep
Ensete_ventricosum_SRR22215668.fasta.transdecoder.pep
Musa_beccarii_GCA_024322285.fasta.transdecoder.pep
Musa_textilis_GCA_964270295.fasta.transdecoder.pep
Musa_troglodytarum_GCA_023547065.fasta.transdecoder.pep
Musa_balbisiana_SRR7006127.fasta.transdecoder.pep
Musa_balbisiana_GCA_004837865.fasta.transdecoder.pep
Musa_acuminata_GCF_036884655.fasta.transdecoder.pep
Musa_acuminata_GCA_964270305.fasta.transdecoder.pep
Musa_itinerans_GCA_001649415.fasta.transdecoder.pep
Musa_itinerans_SRR28089652.fasta.transdecoder.pep
Musa_itinerans_SRR11085724.fasta.transdecoder.pep
Maranta_leuconeura_ERR2040759.fasta.transdecoder.pep
Thalia_dealbata_SRR23717304.fasta.transdecoder.pep
Thalia_dealbata_GCA_030544415.fasta.transdecoder.pep
Ischnosiphon_puberulus_SRR28549745.fasta.transdecoder.pep
Ischnosiphon_puberulus_GCA_040436625.fasta.transdecoder.pep
Canna_x_generalis_SRR26856490.fasta.transdecoder.pep
Canna_indica_GCA_034359265.fasta.transdecoder.pep
Canna_glauca_SRR26624636.fasta.transdecoder.pep
Canna_indica_SRR26152283.fasta.transdecoder.pep
Canna_glauca_SRR26624639.fasta.transdecoder.pep
Costus_pulverulentus_GCA_027562315.fasta.transdecoder.pep
Costus_allenii_GCA_047495955.fasta.transdecoder.pep
Costus_lasius_GCA_027563935.fasta.transdecoder.pep
Alpinia_galanga_GCA_037177395.fasta.transdecoder.pep
Etlingera_elatior_GCA_040939285.fasta.transdecoder.pep
Wurfbainia_villosa_GCA_025175895.fasta.transdecoder.pep
Curcuma_caesia_GCA_037834155.fasta.transdecoder.pep
38
Hedychium_gardnerianum_GCA_038050645.fasta.transdecoder.pep
Hedychium_flavescens_GCA_041731415.fasta.transdecoder.pep
Boesenbergia_rotunda_GCA_024364735.fasta.transdecoder.pep
Zingiber_zerumbet_GCA_037833815.fasta.transdecoder.pep
Zingiber_officinale_GCF_018446385.fasta.transdecoder.pep
Kaempferia_galanga_GCA_037974795.fasta.transdecoder.pep
Kaempferia_parviflora_GCA_037177295.fasta.transdecoder.pep
Kaempferia_rotunda_GCA_037177415.fasta.transdecoder.pep
Kaempferia_elegans_GCA_037974815.fasta.transdecoder.pep
1.0
0.8
0.6
0.4
0.2
0.0

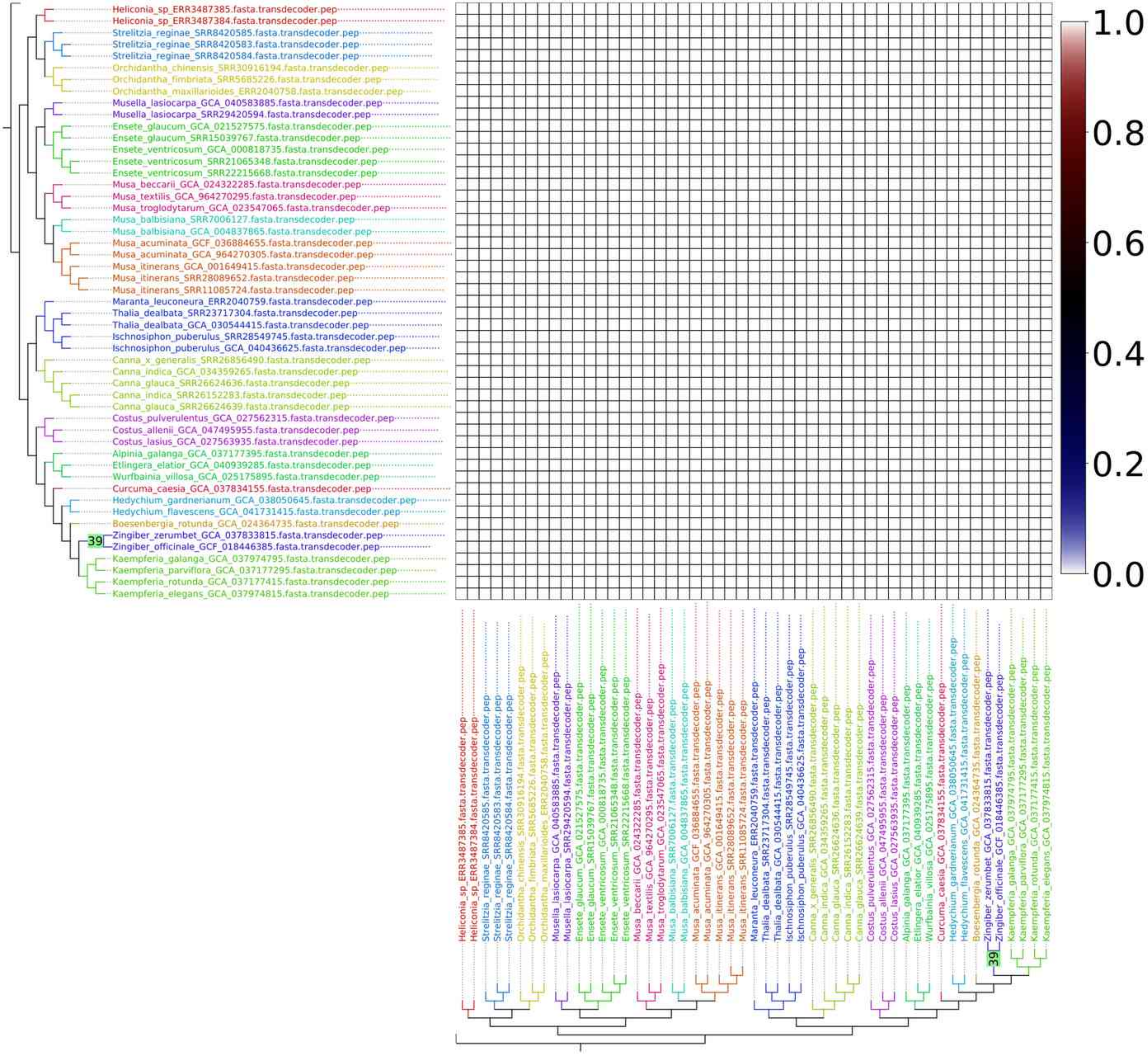
Heliconia_sp_ERR3487385.fasta.transdecoder.pep
Heliconia_sp_ERR3487384.fasta.transdecoder.pep
Strelitzia_reginae_SRR8420585.fasta.transdecoder.pep
Strelitzia_reginae_SRR8420583.fasta.transdecoder.pep
Strelitzia_reginae_SRR8420584.fasta.transdecoder.pep
Orchidantha_chinensis_SRR30916194.fasta.transdecoder.pep
Orchidantha_fimbriata_SRR5685226.fasta.transdecoder.pep
Orchidantha_maxillarioides_ERR2040758.fasta.transdecoder.pep
Musella_lasiocarpa_GCA_040583885.fasta.transdecoder.pep
Musella_lasiocarpa_SRR29420594.fasta.transdecoder.pep
Ensete_glaucum_GCA_021527575.fasta.transdecoder.pep
Ensete_glaucum_SRR15039767.fasta.transdecoder.pep
Ensete_ventricosum_GCA_000818735.fasta.transdecoder.pep
Ensete_ventricosum_SRR21065348.fasta.transdecoder.pep
Ensete_ventricosum_SRR22215668.fasta.transdecoder.pep
Musa_beccarii_GCA_024322285.fasta.transdecoder.pep
Musa_textilis_GCA_964270295.fasta.transdecoder.pep
Musa_troglodytarum_GCA_023547065.fasta.transdecoder.pep
Musa_balbisiana_SRR7006127.fasta.transdecoder.pep
Musa_balbisiana_GCA_004837865.fasta.transdecoder.pep
Musa_acuminata_GCF_036884655.fasta.transdecoder.pep
Musa_acuminata_GCA_964270305.fasta.transdecoder.pep
Musa_itinerans_GCA_001649415.fasta.transdecoder.pep
Musa_itinerans_SRR28089652.fasta.transdecoder.pep
Musa_itinerans_SRR11085724.fasta.transdecoder.pep
Maranta_leuconeura_ERR2040759.fasta.transdecoder.pep
Thalia_dealbata_SRR23717304.fasta.transdecoder.pep
Thalia_dealbata_GCA_030544415.fasta.transdecoder.pep
Ischnosiphon_puberulus_SRR28549745.fasta.transdecoder.pep
Ischnosiphon_puberulus_GCA_040436625.fasta.transdecoder.pep
Canna_x_generalis_SRR26856490.fasta.transdecoder.pep
Canna_indica_GCA_034359265.fasta.transdecoder.pep
Canna_glauca_SRR26624636.fasta.transdecoder.pep
Canna_indica_SRR26152283.fasta.transdecoder.pep
Canna_glauca_SRR26624639.fasta.transdecoder.pep
Costus_pulverulentus_GCA_027562315.fasta.transdecoder.pep
Costus_allenii_GCA_047495955.fasta.transdecoder.pep
Costus_lasius_GCA_027563935.fasta.transdecoder.pep
Alpinia_galanga_GCA_037177395.fasta.transdecoder.pep
Etlingera_elatior_GCA_040939285.fasta.transdecoder.pep
Wurfbainia_villosa_GCA_025175895.fasta.transdecoder.pep
Curcuma_caesia_GCA_037834155.fasta.transdecoder.pep
Hedychium_gardnerianum_GCA_038050645.fasta.transdecoder.pep
Hedychium_flavescens_GCA_041731415.fasta.transdecoder.pep
Boesenbergia_rotunda_GCA_024364735.fasta.transdecoder.pep
39
Zingiber_zerumbet_GCA_037833815.fasta.transdecoder.pep
Zingiber_officinale_GCF_018446385.fasta.transdecoder.pep
Kaempferia_galanga_GCA_037974795.fasta.transdecoder.pep
Kaempferia_parviflora_GCA_037177295.fasta.transdecoder.pep
Kaempferia_rotunda_GCA_037177415.fasta.transdecoder.pep
Kaempferia_elegans_GCA_037974815.fasta.transdecoder.pep
1.0
0.8
0.6
0.4
0.2
0.0

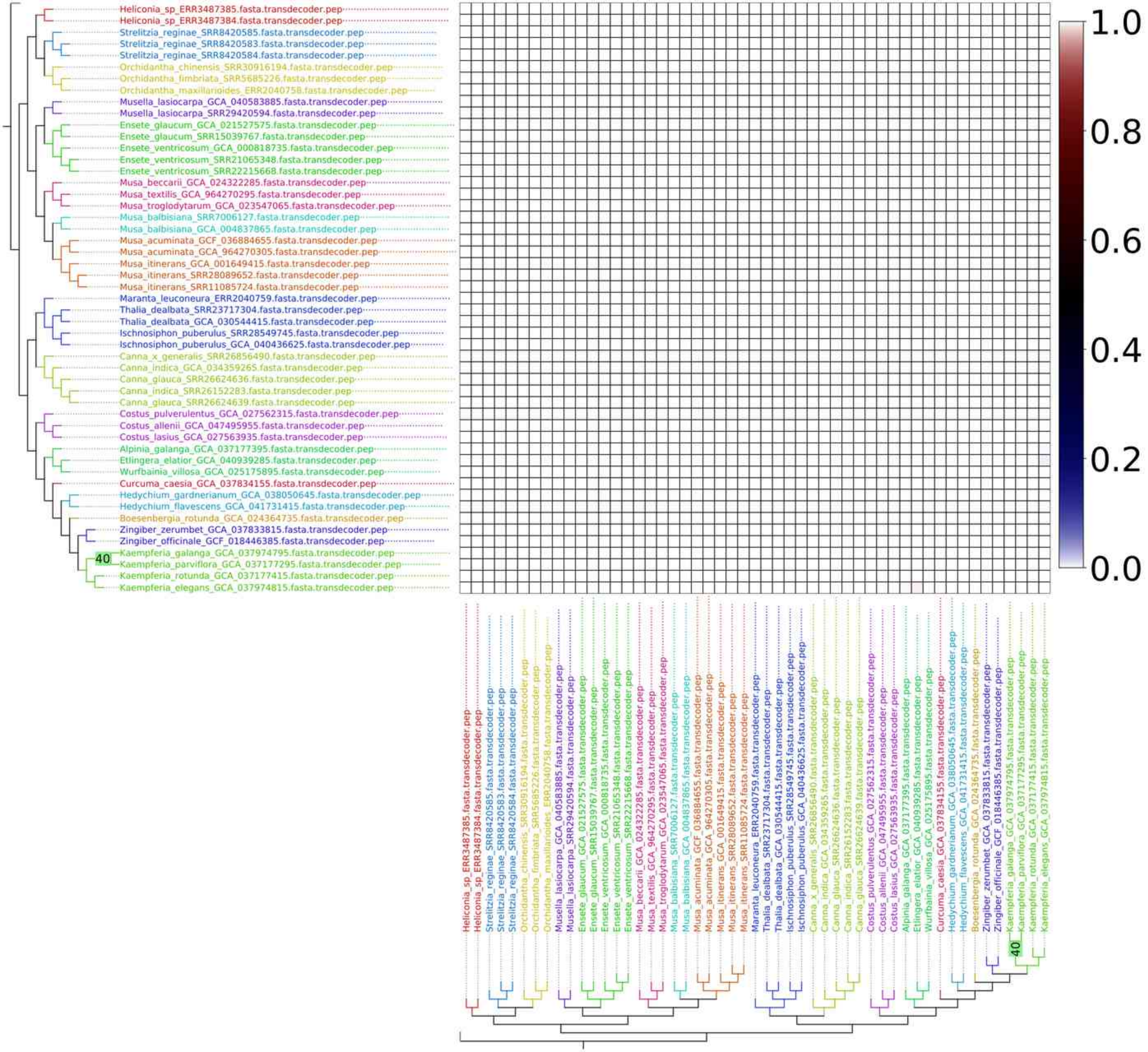
Heliconia_sp_ERR3487385.fasta.transdecoder.pep
Heliconia_sp_ERR3487384.fasta.transdecoder.pep
Strelitzia_reginae_SRR8420585.fasta.transdecoder.pep
Strelitzia_reginae_SRR8420583.fasta.transdecoder.pep
Strelitzia_reginae_SRR8420584.fasta.transdecoder.pep
Orchidantha_chinensis_SRR30916194.fasta.transdecoder.pep
Orchidantha_fimbriata_SRR5685226.fasta.transdecoder.pep
Orchidantha_maxillarioides_ERR2040758.fasta.transdecoder.pep
Musella_lasiocarpa_GCA_040583885.fasta.transdecoder.pep
Musella_lasiocarpa_SRR29420594.fasta.transdecoder.pep
Ensete_glaucum_GCA_021527575.fasta.transdecoder.pep
Ensete_glaucum_SRR15039767.fasta.transdecoder.pep
Ensete_ventricosum_GCA_000818735.fasta.transdecoder.pep
Ensete_ventricosum_SRR21065348.fasta.transdecoder.pep
Ensete_ventricosum_SRR22215668.fasta.transdecoder.pep
Musa_beccarii_GCA_024322285.fasta.transdecoder.pep
Musa_textilis_GCA_964270295.fasta.transdecoder.pep
Musa_troglodytarum_GCA_023547065.fasta.transdecoder.pep
Musa_balbisiana_SRR7006127.fasta.transdecoder.pep
Musa_balbisiana_GCA_004837865.fasta.transdecoder.pep
Musa_acuminata_GCF_036884655.fasta.transdecoder.pep
Musa_acuminata_GCA_964270305.fasta.transdecoder.pep
Musa_itinerans_GCA_001649415.fasta.transdecoder.pep
Musa_itinerans_SRR28089652.fasta.transdecoder.pep
Musa_itinerans_SRR11085724.fasta.transdecoder.pep
Maranta_leuconeura_ERR2040759.fasta.transdecoder.pep
Thalia_dealbata_SRR23717304.fasta.transdecoder.pep
Thalia_dealbata_GCA_030544415.fasta.transdecoder.pep
Ischnosiphon_puberulus_SRR28549745.fasta.transdecoder.pep
Ischnosiphon_puberulus_GCA_040436625.fasta.transdecoder.pep
Canna_x_generalis_SRR26856490.fasta.transdecoder.pep
Canna_indica_GCA_034359265.fasta.transdecoder.pep
Canna_glauca_SRR26624636.fasta.transdecoder.pep
Canna_indica_SRR26152283.fasta.transdecoder.pep
Canna_glauca_SRR26624639.fasta.transdecoder.pep
Costus_pulverulentus_GCA_027562315.fasta.transdecoder.pep
Costus_allenii_GCA_047495955.fasta.transdecoder.pep
Costus_lasius_GCA_027563935.fasta.transdecoder.pep
Alpinia_galanga_GCA_037177395.fasta.transdecoder.pep
Etlingera_elatior_GCA_040939285.fasta.transdecoder.pep
Wurfbainia_villosa_GCA_025175895.fasta.transdecoder.pep
Curcuma_caesia_GCA_037834155.fasta.transdecoder.pep
Hedychium_gardnerianum_GCA_038050645.fasta.transdecoder.pep
Hedychium_flavescens_GCA_041731415.fasta.transdecoder.pep
Boesenbergia_rotunda_GCA_024364735.fasta.transdecoder.pep
Zingiber_zerumbet_GCA_037833815.fasta.transdecoder.pep
Zingiber_officinale_GCF_018446385.fasta.transdecoder.pep
40
Kaempferia_galanga_GCA_037974795.fasta.transdecoder.pep
Kaempferia_parviflora_GCA_037177295.fasta.transdecoder.pep
Kaempferia_rotunda_GCA_037177415.fasta.transdecoder.pep
Kaempferia_elegans_GCA_037974815.fasta.transdecoder.pep
1.0
0.8
0.6
0.4
0.2
0.0

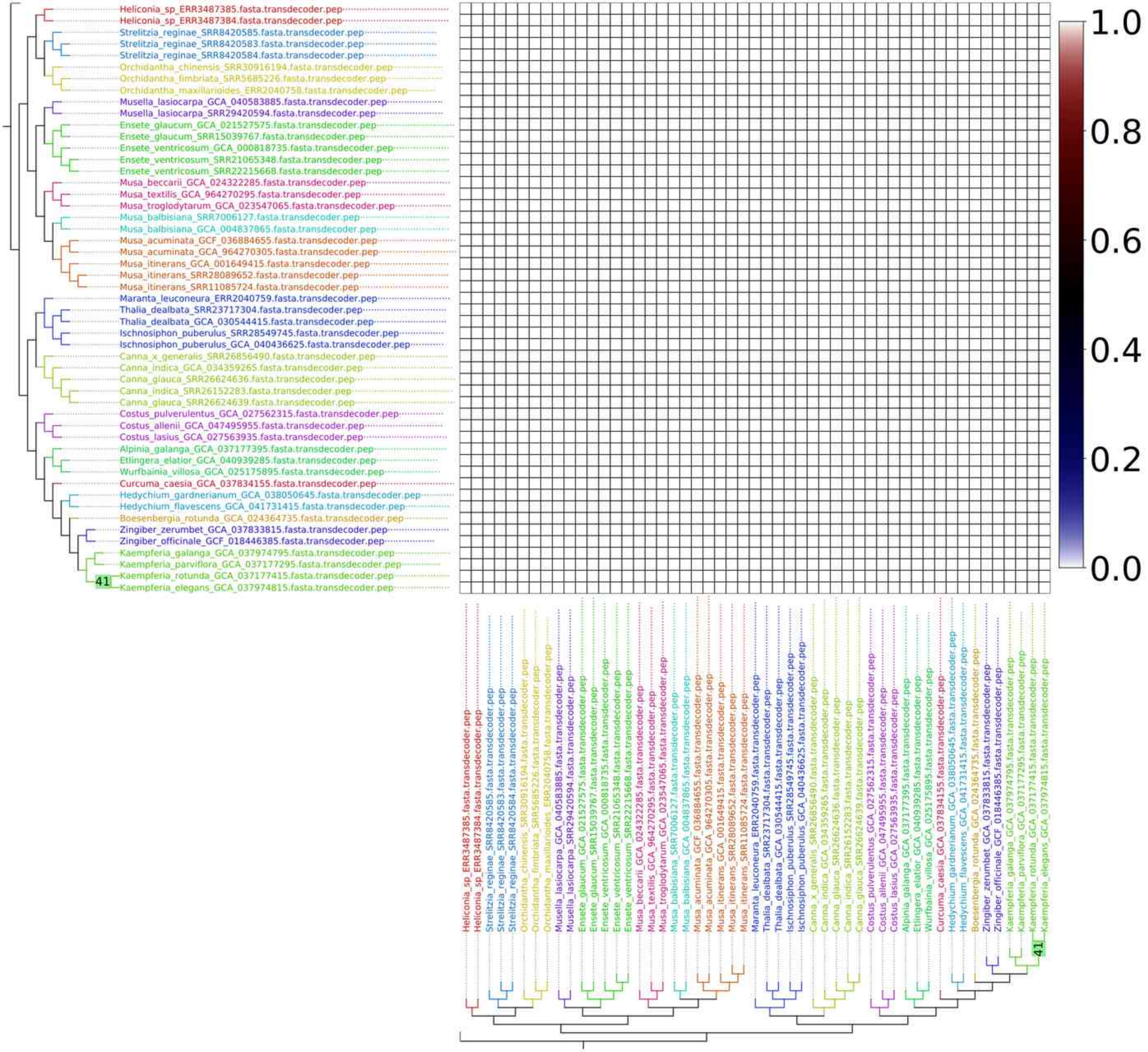

Heliconia_sp_ERR3487385.fasta.transdecoder.pep
Heliconia_sp_ERR3487384.fasta.transdecoder.pep
Strelitzia_reginae_SRR8420585.fasta.transdecoder.pep
Strelitzia_reginae_SRR8420583.fasta.transdecoder.pep
Strelitzia_reginae_SRR8420584.fasta.transdecoder.pep
Orchidantha_chinensis_SRR30916194.fasta.transdecoder.pep
Orchidantha_fimbriata_SRR5685226.fasta.transdecoder.pep
Orchidantha_maxillarioides_ERR2040758.fasta.transdecoder.pep
Musella_lasiocarpa_GCA_040583885.fasta.transdecoder.pep
Musella_lasiocarpa_SRR29420594.fasta.transdecoder.pep
Ensete_glaucum_GCA_021527575.fasta.transdecoder.pep
Ensete_glaucum_SRR15039767.fasta.transdecoder.pep
Ensete_ventricosum_GCA_000818735.fasta.transdecoder.pep
Ensete_ventricosum_SRR21065348.fasta.transdecoder.pep
Ensete_ventricosum_SRR22215668.fasta.transdecoder.pep
Musa_beccarii_GCA_024322285.fasta.transdecoder.pep
Musa_textilis_GCA_964270295.fasta.transdecoder.pep
Musa_troglodytarum_GCA_023547065.fasta.transdecoder.pep
Musa_balbisiana_SRR7006127.fasta.transdecoder.pep
Musa_balbisiana_GCA_004837865.fasta.transdecoder.pep
Musa_acuminata_GCF_036884655.fasta.transdecoder.pep
Musa_acuminata_GCA_964270305.fasta.transdecoder.pep
Musa_itinerans_GCA_001649415.fasta.transdecoder.pep
Musa_itinerans_SRR28089652.fasta.transdecoder.pep
Musa_itinerans_SRR11085724.fasta.transdecoder.pep
Maranta_leuconeura_ERR2040759.fasta.transdecoder.pep
Thalia_dealbata_SRR23717304.fasta.transdecoder.pep
Thalia_dealbata_GCA_030544415.fasta.transdecoder.pep
Ischnosiphon_puberulus_SRR28549745.fasta.transdecoder.pep
Ischnosiphon_puberulus_GCA_040436625.fasta.transdecoder.pep
Canna_x_generalis_SRR26856490.fasta.transdecoder.pep
Canna_indica_GCA_034359265.fasta.transdecoder.pep
Canna_glauca_SRR26624636.fasta.transdecoder.pep
Canna_indica_SRR26152283.fasta.transdecoder.pep
Canna_glauca_SRR26624639.fasta.transdecoder.pep
Costus_pulverulentus_GCA_027562315.fasta.transdecoder.pep
Costus_allenii_GCA_047495955.fasta.transdecoder.pep
Costus_lasius_GCA_027563935.fasta.transdecoder.pep
Alpinia_galanga_GCA_037177395.fasta.transdecoder.pep
Etlingera_elatior_GCA_040939285.fasta.transdecoder.pep
Wurfbainia_villosa_GCA_025175895.fasta.transdecoder.pep
Curcuma_caesia_GCA_037834155.fasta.transdecoder.pep
Hedychium_gardnerianum_GCA_038050645.fasta.transdecoder.pep
Hedychium_flavescens_GCA_041731415.fasta.transdecoder.pep
Boesenbergia_rotunda_GCA_024364735.fasta.transdecoder.pep
Zingiber_zerumbet_GCA_037833815.fasta.transdecoder.pep
Zingiber_officinale_GCF_018446385.fasta.transdecoder.pep
Kaempferia_galanga_GCA_037974795.fasta.transdecoder.pep
Kaempferia_parviflora_GCA_037177295.fasta.transdecoder.pep
Kaempferia_rotunda_GCA_037177415.fasta.transdecoder.pep
Kaempferia_elegans_GCA_037974815.fasta.transdecoder.pep
41
1.0
0.8
0.6
0.4
0.2
0.0

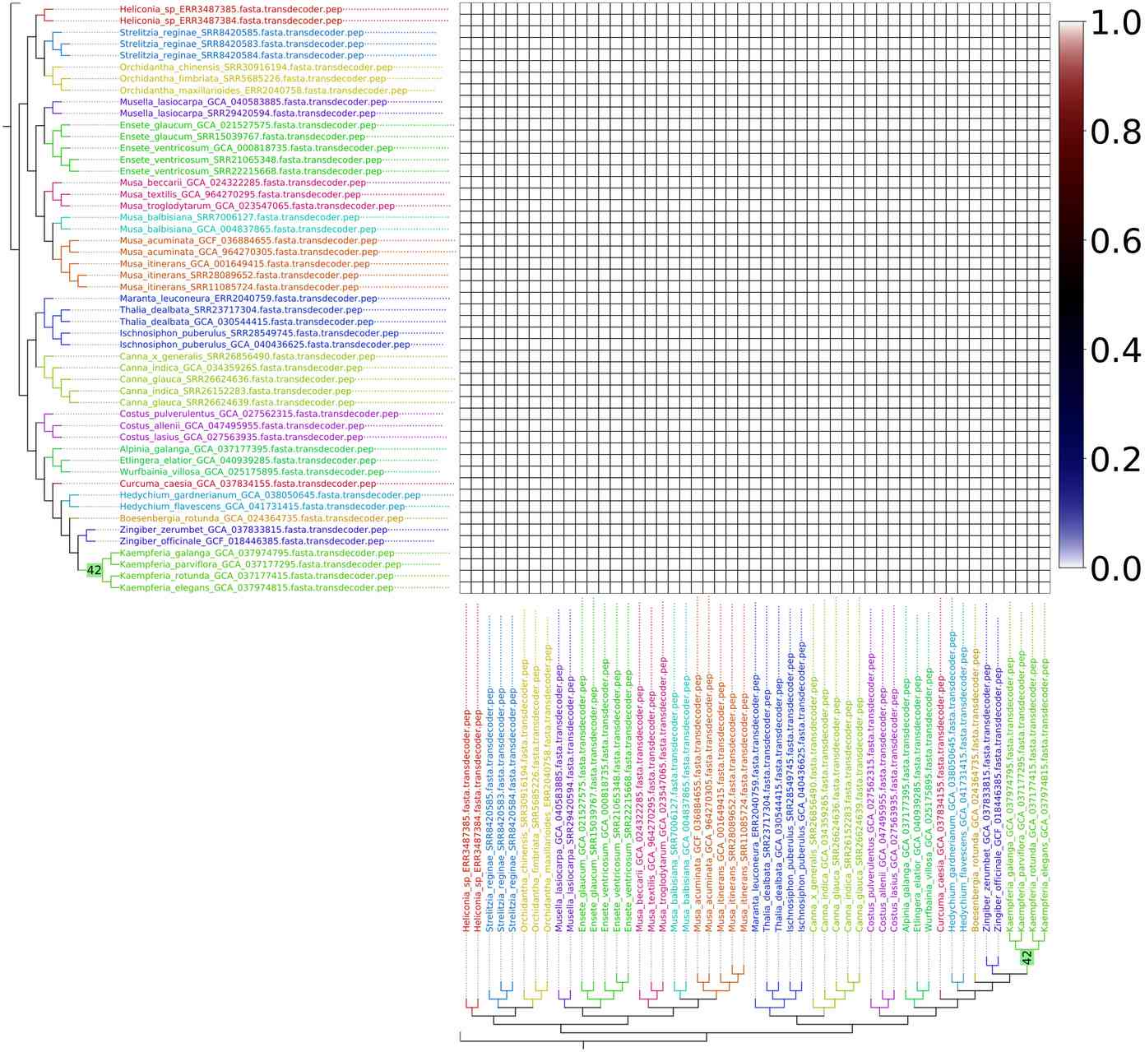
Heliconia_sp_ERR3487385.fasta.transdecoder.pep
Heliconia_sp_ERR3487384.fasta.transdecoder.pep
Strelitzia_reginae_SRR8420585.fasta.transdecoder.pep
Strelitzia_reginae_SRR8420583.fasta.transdecoder.pep
Strelitzia_reginae_SRR8420584.fasta.transdecoder.pep
Orchidantha_chinensis_SRR30916194.fasta.transdecoder.pep
Orchidantha_fimbriata_SRR5685226.fasta.transdecoder.pep
Orchidantha_maxillarioides_ERR2040758.fasta.transdecoder.pep
Musella_lasiocarpa_GCA_040583885.fasta.transdecoder.pep
Musella_lasiocarpa_SRR29420594.fasta.transdecoder.pep
Ensete_glaucum_GCA_021527575.fasta.transdecoder.pep
Ensete_glaucum_SRR15039767.fasta.transdecoder.pep
Ensete_ventricosum_GCA_000818735.fasta.transdecoder.pep
Ensete_ventricosum_SRR21065348.fasta.transdecoder.pep
Ensete_ventricosum_SRR22215668.fasta.transdecoder.pep
Musa_beccarii_GCA_024322285.fasta.transdecoder.pep
Musa_textilis_GCA_964270295.fasta.transdecoder.pep
Musa_troglodytarum_GCA_023547065.fasta.transdecoder.pep
Musa_balbisiana_SRR7006127.fasta.transdecoder.pep
Musa_balbisiana_GCA_004837865.fasta.transdecoder.pep
Musa_acuminata_GCF_036884655.fasta.transdecoder.pep
Musa_acuminata_GCA_964270305.fasta.transdecoder.pep
Musa_itinerans_GCA_001649415.fasta.transdecoder.pep
Musa_itinerans_SRR28089652.fasta.transdecoder.pep
Musa_itinerans_SRR11085724.fasta.transdecoder.pep
Maranta_leuconeura_ERR2040759.fasta.transdecoder.pep
Thalia_dealbata_SRR23717304.fasta.transdecoder.pep
Thalia_dealbata_GCA_030544415.fasta.transdecoder.pep
Ischnosiphon_puberulus_SRR28549745.fasta.transdecoder.pep
Ischnosiphon_puberulus_GCA_040436625.fasta.transdecoder.pep
Canna_x_generalis_SRR26856490.fasta.transdecoder.pep
Canna_indica_GCA_034359265.fasta.transdecoder.pep
Canna_glauca_SRR26624636.fasta.transdecoder.pep
Canna_indica_SRR26152283.fasta.transdecoder.pep
Canna_glauca_SRR26624639.fasta.transdecoder.pep
Costus_pulverulentus_GCA_027562315.fasta.transdecoder.pep
Costus_allenii_GCA_047495955.fasta.transdecoder.pep
Costus_lasius_GCA_027563935.fasta.transdecoder.pep
Alpinia_galanga_GCA_037177395.fasta.transdecoder.pep
Etlingera_elatior_GCA_040939285.fasta.transdecoder.pep
Wurfbainia_villosa_GCA_025175895.fasta.transdecoder.pep
Curcuma_caesia_GCA_037834155.fasta.transdecoder.pep
Hedychium_gardnerianum_GCA_038050645.fasta.transdecoder.pep
Hedychium_flavescens_GCA_041731415.fasta.transdecoder.pep
Boesenbergia_rotunda_GCA_024364735.fasta.transdecoder.pep
Zingiber_zerumbet_GCA_037833815.fasta.transdecoder.pep
Zingiber_officinale_GCF_018446385.fasta.transdecoder.pep
Kaempferia_galanga_GCA_037974795.fasta.transdecoder.pep
Kaempferia_parviflora_GCA_037177295.fasta.transdecoder.pep
Kaempferia_rotunda_GCA_037177415.fasta.transdecoder.pep
Kaempferia_elegans_GCA_037974815.fasta.transdecoder.pep
42
1.0
0.8
0.6
0.4
0.2
0.0

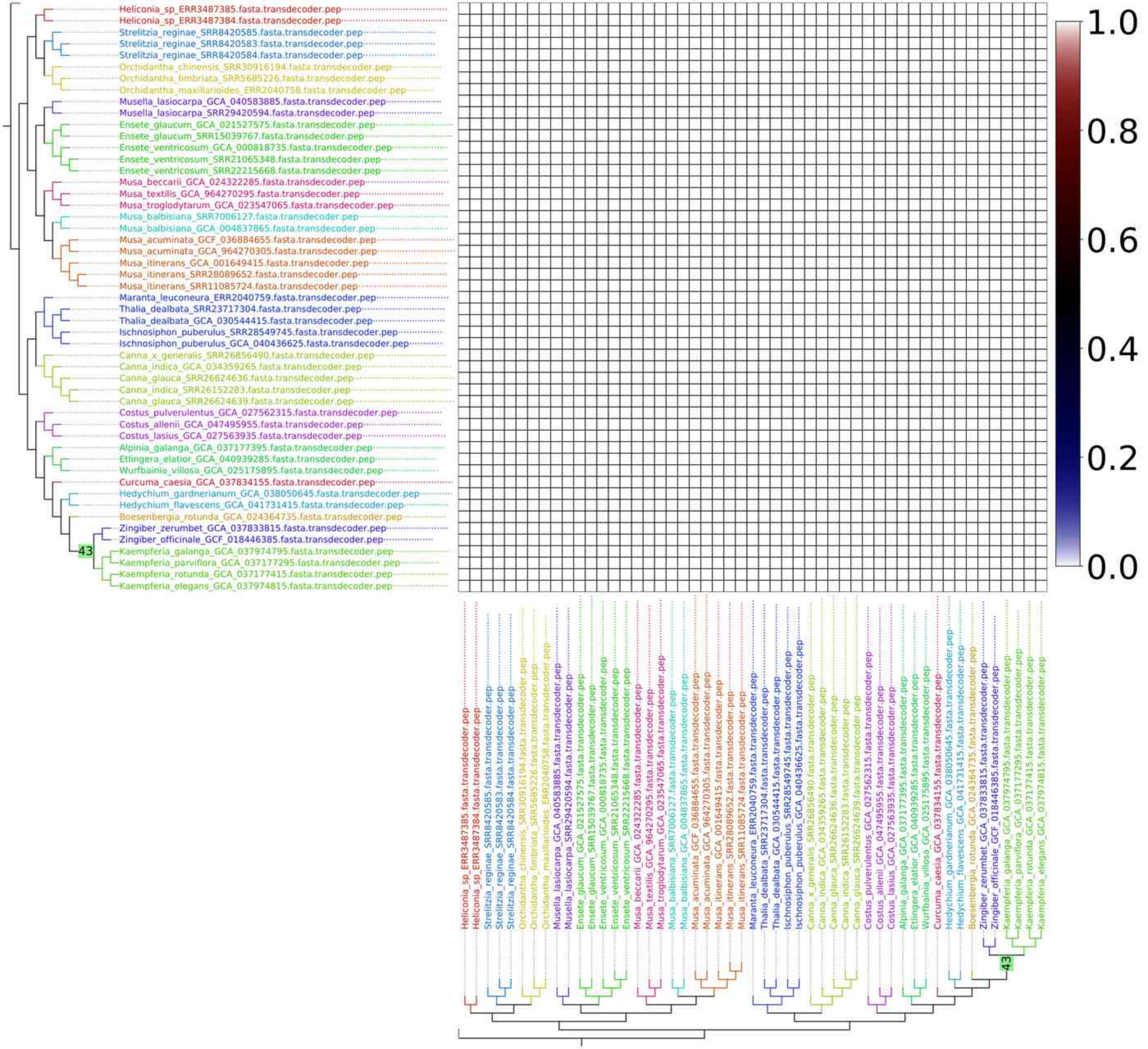

Heliconia_sp_ERR3487385.fasta.transdecoder.pep
Heliconia_sp_ERR3487384.fasta.transdecoder.pep
Strelitzia_reginae_SRR8420585.fasta.transdecoder.pep
Strelitzia_reginae_SRR8420583.fasta.transdecoder.pep
Strelitzia_reginae_SRR8420584.fasta.transdecoder.pep
Orchidantha_chinensis_SRR30916194.fasta.transdecoder.pep
Orchidantha_fimbriata_SRR5685226.fasta.transdecoder.pep
Orchidantha_maxillarioides_ERR2040758.fasta.transdecoder.pep
Musella_lasiocarpa_GCA_040583885.fasta.transdecoder.pep
Musella_lasiocarpa_SRR29420594.fasta.transdecoder.pep
Ensete_glaucum_GCA_021527575.fasta.transdecoder.pep
Ensete_glaucum_SRR15039767.fasta.transdecoder.pep
Ensete_ventricosum_GCA_000818735.fasta.transdecoder.pep
Ensete_ventricosum_SRR21065348.fasta.transdecoder.pep
Ensete_ventricosum_SRR22215668.fasta.transdecoder.pep
Musa_beccarii_GCA_024322285.fasta.transdecoder.pep
Musa_textilis_GCA_964270295.fasta.transdecoder.pep
Musa_troglodytarum_GCA_023547065.fasta.transdecoder.pep
Musa_balbisiana_SRR7006127.fasta.transdecoder.pep
Musa_balbisiana_GCA_004837865.fasta.transdecoder.pep
Musa_acuminata_GCF_036884655.fasta.transdecoder.pep
Musa_acuminata_GCA_964270305.fasta.transdecoder.pep
Musa_itinerans_GCA_001649415.fasta.transdecoder.pep
Musa_itinerans_SRR28089652.fasta.transdecoder.pep
Musa_itinerans_SRR11085724.fasta.transdecoder.pep
Maranta_leuconeura_ERR2040759.fasta.transdecoder.pep
Thalia_dealbata_SRR23717304.fasta.transdecoder.pep
Thalia_dealbata_GCA_030544415.fasta.transdecoder.pep
Ischnosiphon_puberulus_SRR28549745.fasta.transdecoder.pep
Ischnosiphon_puberulus_GCA_040436625.fasta.transdecoder.pep
Canna_x_generalis_SRR26856490.fasta.transdecoder.pep
Canna_indica_GCA_034359265.fasta.transdecoder.pep
Canna_glauca_SRR26624636.fasta.transdecoder.pep
Canna_indica_SRR26152283.fasta.transdecoder.pep
Canna_glauca_SRR26624639.fasta.transdecoder.pep
Costus_pulverulentus_GCA_027562315.fasta.transdecoder.pep
Costus_allenii_GCA_047495955.fasta.transdecoder.pep
Costus_lasius_GCA_027563935.fasta.transdecoder.pep
Alpinia_galanga_GCA_037177395.fasta.transdecoder.pep
Etlingera_elatior_GCA_040939285.fasta.transdecoder.pep
Wurfbainia_villosa_GCA_025175895.fasta.transdecoder.pep
Curcuma_caesia_GCA_037834155.fasta.transdecoder.pep
Hedychium_gardnerianum_GCA_038050645.fasta.transdecoder.pep
Hedychium_flavescens_GCA_041731415.fasta.transdecoder.pep
Boesenbergia_rotunda_GCA_024364735.fasta.transdecoder.pep
Zingiber_zerumbet_GCA_037833815.fasta.transdecoder.pep
Zingiber_officinale_GCF_018446385.fasta.transdecoder.pep
43
Kaempferia_galanga_GCA_037974795.fasta.transdecoder.pep
Kaempferia_parviflora_GCA_037177295.fasta.transdecoder.pep
Kaempferia_rotunda_GCA_037177415.fasta.transdecoder.pep
Kaempferia_elegans_GCA_037974815.fasta.transdecoder.pep
1.0
0.8
0.6
0.4
0.2
0.0

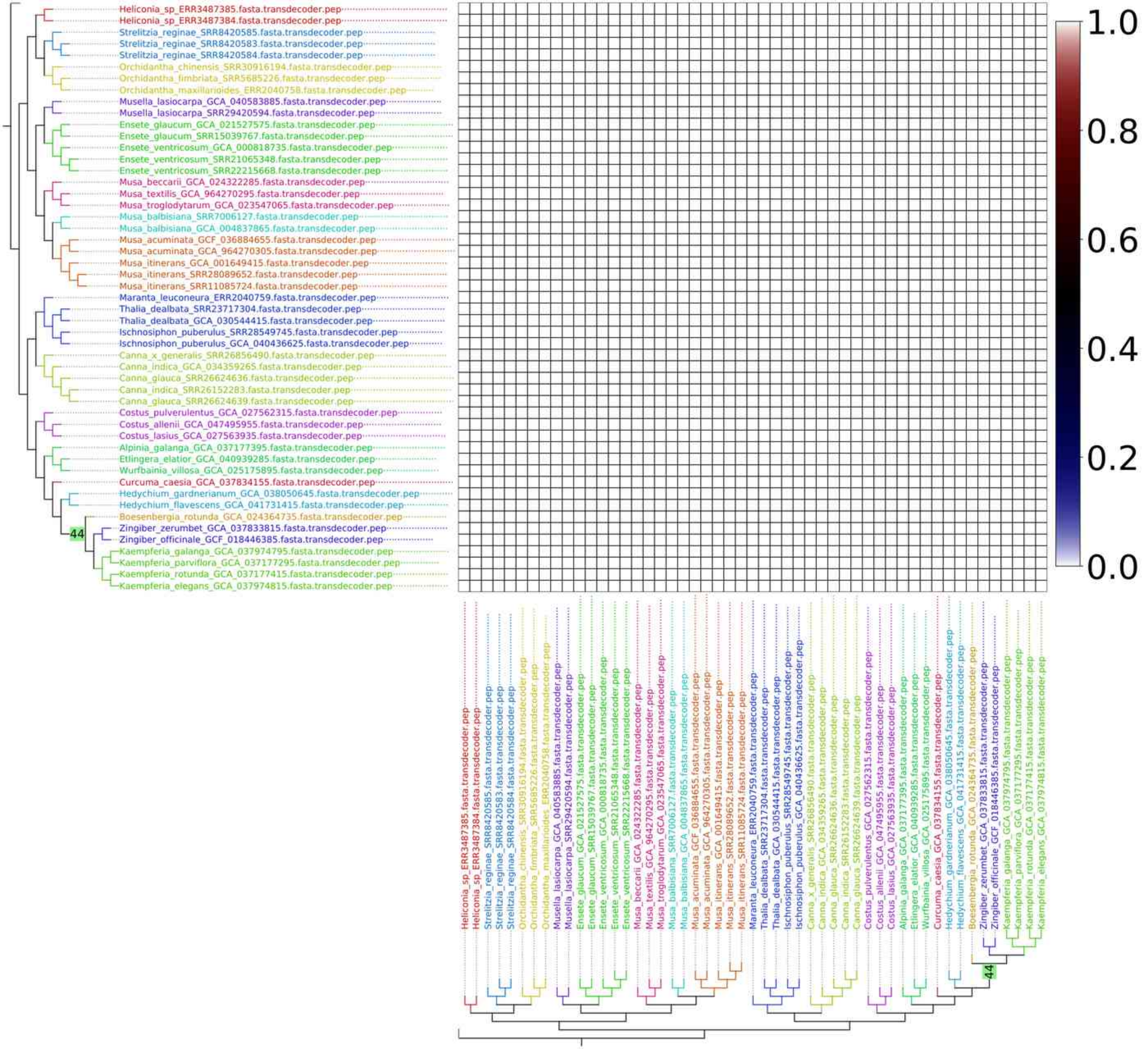
Heliconia_sp_ERR3487385.fasta.transdecoder.pep
Heliconia_sp_ERR3487384.fasta.transdecoder.pep
Strelitzia_reginae_SRR8420585.fasta.transdecoder.pep
Strelitzia_reginae_SRR8420583.fasta.transdecoder.pep
Strelitzia_reginae_SRR8420584.fasta.transdecoder.pep
Orchidantha_chinensis_SRR30916194.fasta.transdecoder.pep
Orchidantha_fimbriata_SRR5685226.fasta.transdecoder.pep
Orchidantha_maxillarioides_ERR2040758.fasta.transdecoder.pep
Musella_lasiocarpa_GCA_040583885.fasta.transdecoder.pep
Musella_lasiocarpa_SRR29420594.fasta.transdecoder.pep
Ensete_glaucum_GCA_021527575.fasta.transdecoder.pep
Ensete_glaucum_SRR15039767.fasta.transdecoder.pep
Ensete_ventricosum_GCA_000818735.fasta.transdecoder.pep
Ensete_ventricosum_SRR21065348.fasta.transdecoder.pep
Ensete_ventricosum_SRR22215668.fasta.transdecoder.pep
Musa_beccarii_GCA_024322285.fasta.transdecoder.pep
Musa_textilis_GCA_964270295.fasta.transdecoder.pep
Musa_troglodytarum_GCA_023547065.fasta.transdecoder.pep
Musa_balbisiana_SRR7006127.fasta.transdecoder.pep
Musa_balbisiana_GCA_004837865.fasta.transdecoder.pep
Musa_acuminata_GCF_036884655.fasta.transdecoder.pep
Musa_acuminata_GCA_964270305.fasta.transdecoder.pep
Musa_itinerans_GCA_001649415.fasta.transdecoder.pep
Musa_itinerans_SRR28089652.fasta.transdecoder.pep
Musa_itinerans_SRR11085724.fasta.transdecoder.pep
Maranta_leuconeura_ERR2040759.fasta.transdecoder.pep
Thalia_dealbata_SRR23717304.fasta.transdecoder.pep
Thalia_dealbata_GCA_030544415.fasta.transdecoder.pep
Ischnosiphon_puberulus_SRR28549745.fasta.transdecoder.pep
Ischnosiphon_puberulus_GCA_040436625.fasta.transdecoder.pep
Canna_x_generalis_SRR26856490.fasta.transdecoder.pep
Canna_indica_GCA_034359265.fasta.transdecoder.pep
Canna_glauca_SRR26624636.fasta.transdecoder.pep
Canna_indica_SRR26152283.fasta.transdecoder.pep
Canna_glauca_SRR26624639.fasta.transdecoder.pep
Costus_pulverulentus_GCA_027562315.fasta.transdecoder.pep
Costus_allenii_GCA_047495955.fasta.transdecoder.pep
Costus_lasius_GCA_027563935.fasta.transdecoder.pep
Alpinia_galanga_GCA_037177395.fasta.transdecoder.pep
Etlingera_elatior_GCA_040939285.fasta.transdecoder.pep
Wurfbainia_villosa_GCA_025175895.fasta.transdecoder.pep
Curcuma_caesia_GCA_037834155.fasta.transdecoder.pep
Hedychium_gardnerianum_GCA_038050645.fasta.transdecoder.pep
Hedychium_flavescens_GCA_041731415.fasta.transdecoder.pep
Boesenbergia_rotunda_GCA_024364735.fasta.transdecoder.pep
44
Zingiber_zerumbet_GCA_037833815.fasta.transdecoder.pep
Zingiber_officinale_GCF_018446385.fasta.transdecoder.pep
Kaempferia_galanga_GCA_037974795.fasta.transdecoder.pep
Kaempferia_parviflora_GCA_037177295.fasta.transdecoder.pep
Kaempferia_rotunda_GCA_037177415.fasta.transdecoder.pep
Kaempferia_elegans_GCA_037974815.fasta.transdecoder.pep
1.0
0.8
0.6
0.4
0.2
0.0

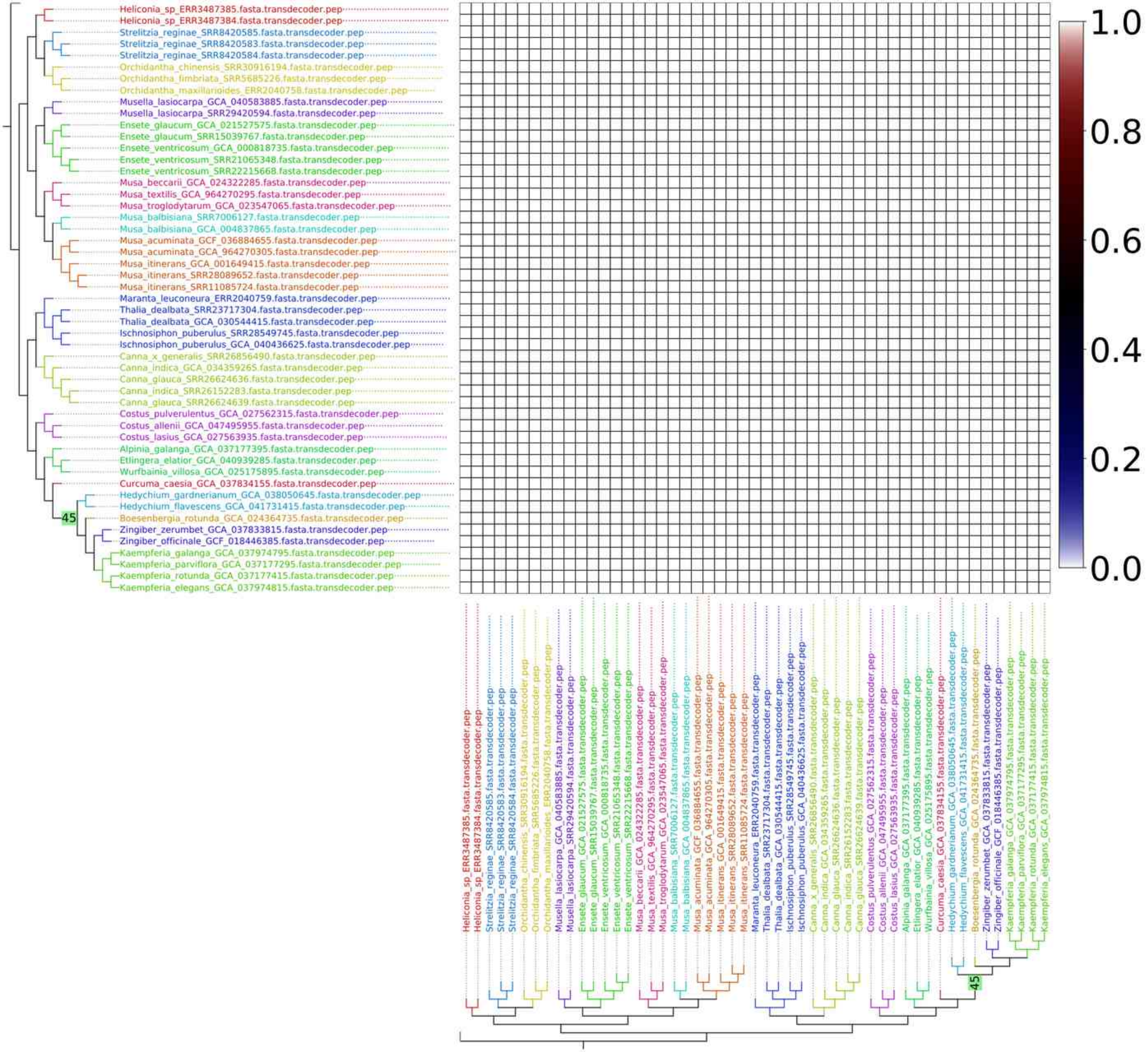
Heliconia_sp_ERR3487385.fasta.transdecoder.pep
Heliconia_sp_ERR3487384.fasta.transdecoder.pep
Strelitzia_reginae_SRR8420585.fasta.transdecoder.pep
Strelitzia_reginae_SRR8420583.fasta.transdecoder.pep
Strelitzia_reginae_SRR8420584.fasta.transdecoder.pep
Orchidantha_chinensis_SRR30916194.fasta.transdecoder.pep
Orchidantha_fimbriata_SRR5685226.fasta.transdecoder.pep
Orchidantha_maxillarioides_ERR2040758.fasta.transdecoder.pep
Musella_lasiocarpa_GCA_040583885.fasta.transdecoder.pep
Musella_lasiocarpa_SRR29420594.fasta.transdecoder.pep
Ensete_glaucum_GCA_021527575.fasta.transdecoder.pep
Ensete_glaucum_SRR15039767.fasta.transdecoder.pep
Ensete_ventricosum_GCA_000818735.fasta.transdecoder.pep
Ensete_ventricosum_SRR21065348.fasta.transdecoder.pep
Ensete_ventricosum_SRR22215668.fasta.transdecoder.pep
Musa_beccarii_GCA_024322285.fasta.transdecoder.pep
Musa_textilis_GCA_964270295.fasta.transdecoder.pep
Musa_troglodytarum_GCA_023547065.fasta.transdecoder.pep
Musa_balbisiana_SRR7006127.fasta.transdecoder.pep
Musa_balbisiana_GCA_004837865.fasta.transdecoder.pep
Musa_acuminata_GCF_036884655.fasta.transdecoder.pep
Musa_acuminata_GCA_964270305.fasta.transdecoder.pep
Musa_itinerans_GCA_001649415.fasta.transdecoder.pep
Musa_itinerans_SRR28089652.fasta.transdecoder.pep
Musa_itinerans_SRR11085724.fasta.transdecoder.pep
Maranta_leuconeura_ERR2040759.fasta.transdecoder.pep
Thalia_dealbata_SRR23717304.fasta.transdecoder.pep
Thalia_dealbata_GCA_030544415.fasta.transdecoder.pep
Ischnosiphon_puberulus_SRR28549745.fasta.transdecoder.pep
Ischnosiphon_puberulus_GCA_040436625.fasta.transdecoder.pep
Canna_x_generalis_SRR26856490.fasta.transdecoder.pep
Canna_indica_GCA_034359265.fasta.transdecoder.pep
Canna_glauca_SRR26624636.fasta.transdecoder.pep
Canna_indica_SRR26152283.fasta.transdecoder.pep
Canna_glauca_SRR26624639.fasta.transdecoder.pep
Costus_pulverulentus_GCA_027562315.fasta.transdecoder.pep
Costus_allenii_GCA_047495955.fasta.transdecoder.pep
Costus_lasius_GCA_027563935.fasta.transdecoder.pep
Alpinia_galanga_GCA_037177395.fasta.transdecoder.pep
Etlingera_elatior_GCA_040939285.fasta.transdecoder.pep
Wurfbainia_villosa_GCA_025175895.fasta.transdecoder.pep
Curcuma_caesia_GCA_037834155.fasta.transdecoder.pep
Hedychium_gardnerianum_GCA_038050645.fasta.transdecoder.pep
Hedychium_flavescens_GCA_041731415.fasta.transdecoder.pep
Boesenbergia_rotunda_GCA_024364735.fasta.transdecoder.pep
Zingiber_zerumbet_GCA_037833815.fasta.transdecoder.pep
Zingiber_officinale_GCF_018446385.fasta.transdecoder.pep
Kaempferia_galanga_GCA_037974795.fasta.transdecoder.pep
Kaempferia_parviflora_GCA_037177295.fasta.transdecoder.pep
Kaempferia_rotunda_GCA_037177415.fasta.transdecoder.pep
Kaempferia_elegans_GCA_037974815.fasta.transdecoder.pep
45
1.0
0.8
0.6
0.4
0.2
0.0

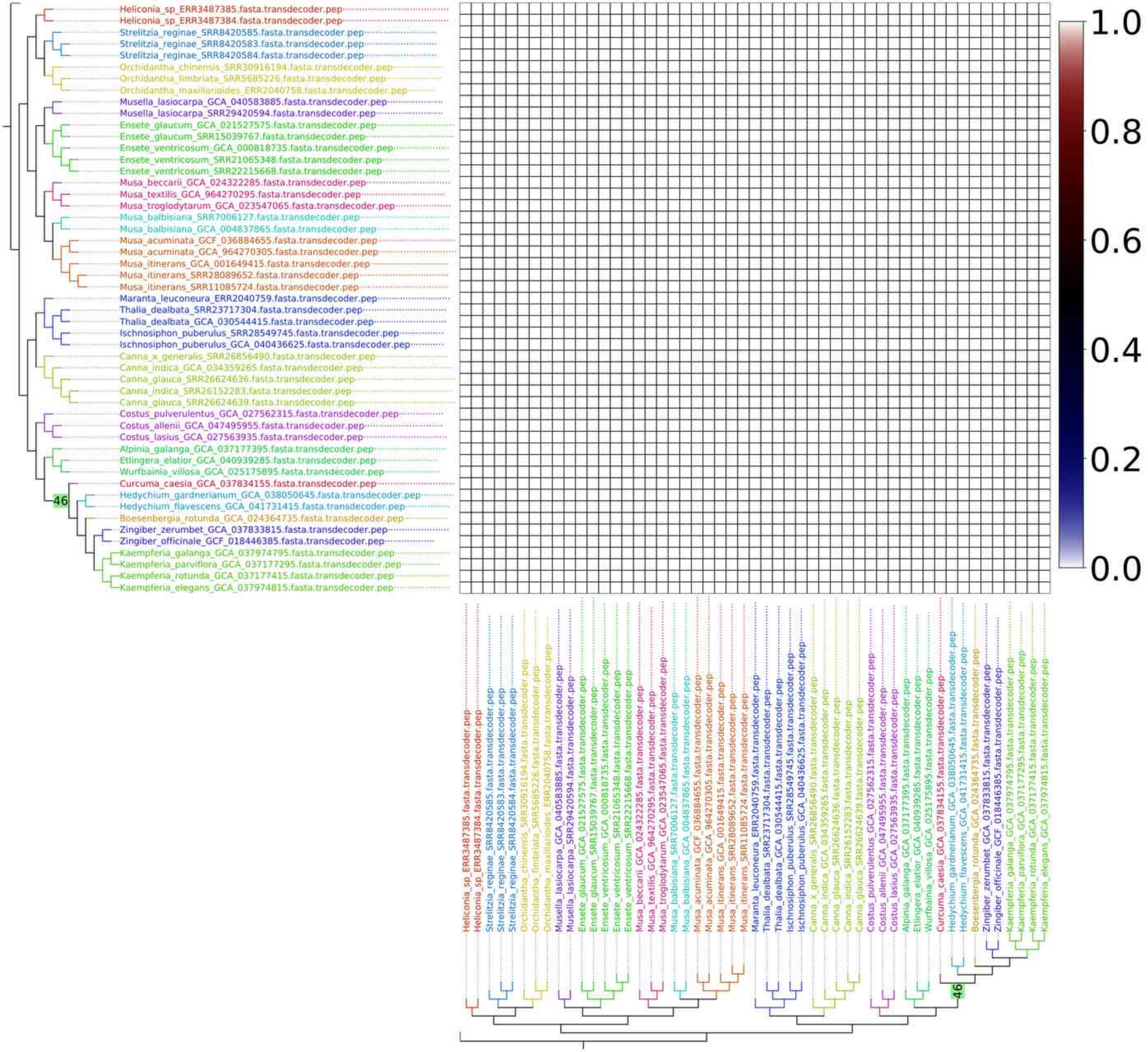

Heliconia_sp_ERR3487385.fasta.transdecoder.pep
Heliconia_sp_ERR3487384.fasta.transdecoder.pep
Strelitzia_reginae_SRR8420585.fasta.transdecoder.pep
Strelitzia_reginae_SRR8420583.fasta.transdecoder.pep
Strelitzia_reginae_SRR8420584.fasta.transdecoder.pep
Orchidantha_chinensis_SRR30916194.fasta.transdecoder.pep
Orchidantha_fimbriata_SRR5685226.fasta.transdecoder.pep
Orchidantha_maxillarioides_ERR2040758.fasta.transdecoder.pep
Musella_lasiocarpa_GCA_040583885.fasta.transdecoder.pep
Musella_lasiocarpa_SRR29420594.fasta.transdecoder.pep
Ensete_glaucum_GCA_021527575.fasta.transdecoder.pep
Ensete_glaucum_SRR15039767.fasta.transdecoder.pep
Ensete_ventricosum_GCA_000818735.fasta.transdecoder.pep
Ensete_ventricosum_SRR21065348.fasta.transdecoder.pep
Ensete_ventricosum_SRR22215668.fasta.transdecoder.pep
Musa_beccarii_GCA_024322285.fasta.transdecoder.pep
Musa_textilis_GCA_964270295.fasta.transdecoder.pep
Musa_troglodytarum_GCA_023547065.fasta.transdecoder.pep
Musa_balbisiana_SRR7006127.fasta.transdecoder.pep
Musa_balbisiana_GCA_004837865.fasta.transdecoder.pep
Musa_acuminata_GCF_036884655.fasta.transdecoder.pep
Musa_acuminata_GCA_964270305.fasta.transdecoder.pep
Musa_itinerans_GCA_001649415.fasta.transdecoder.pep
Musa_itinerans_SRR28089652.fasta.transdecoder.pep
Musa_itinerans_SRR11085724.fasta.transdecoder.pep
Maranta_leuconeura_ERR2040759.fasta.transdecoder.pep
Thalia_dealbata_SRR23717304.fasta.transdecoder.pep
Thalia_dealbata_GCA_030544415.fasta.transdecoder.pep
Ischnosiphon_puberulus_SRR28549745.fasta.transdecoder.pep
Ischnosiphon_puberulus_GCA_040436625.fasta.transdecoder.pep
Canna_x_generalis_SRR26856490.fasta.transdecoder.pep
Canna_indica_GCA_034359265.fasta.transdecoder.pep
Canna_glauca_SRR26624636.fasta.transdecoder.pep
Canna_indica_SRR26152283.fasta.transdecoder.pep
Canna_glauca_SRR26624639.fasta.transdecoder.pep
Costus_pulverulentus_GCA_027562315.fasta.transdecoder.pep
Costus_allenii_GCA_047495955.fasta.transdecoder.pep
Costus_lasius_GCA_027563935.fasta.transdecoder.pep
Alpinia_galanga_GCA_037177395.fasta.transdecoder.pep
Etlingera_elatior_GCA_040939285.fasta.transdecoder.pep
Wurfbainia_villosa_GCA_025175895.fasta.transdecoder.pep
Curcuma_caesia_GCA_037834155.fasta.transdecoder.pep
46
Hedychium_gardnerianum_GCA_038050645.fasta.transdecoder.pep
Hedychium_flavescens_GCA_041731415.fasta.transdecoder.pep
Boesenbergia_rotunda_GCA_024364735.fasta.transdecoder.pep
Zingiber_zerumbet_GCA_037833815.fasta.transdecoder.pep
Zingiber_officinale_GCF_018446385.fasta.transdecoder.pep
Kaempferia_galanga_GCA_037974795.fasta.transdecoder.pep
Kaempferia_parviflora_GCA_037177295.fasta.transdecoder.pep
Kaempferia_rotunda_GCA_037177415.fasta.transdecoder.pep
Kaempferia_elegans_GCA_037974815.fasta.transdecoder.pep
1.0
0.8
0.6
0.4
0.2
0.0

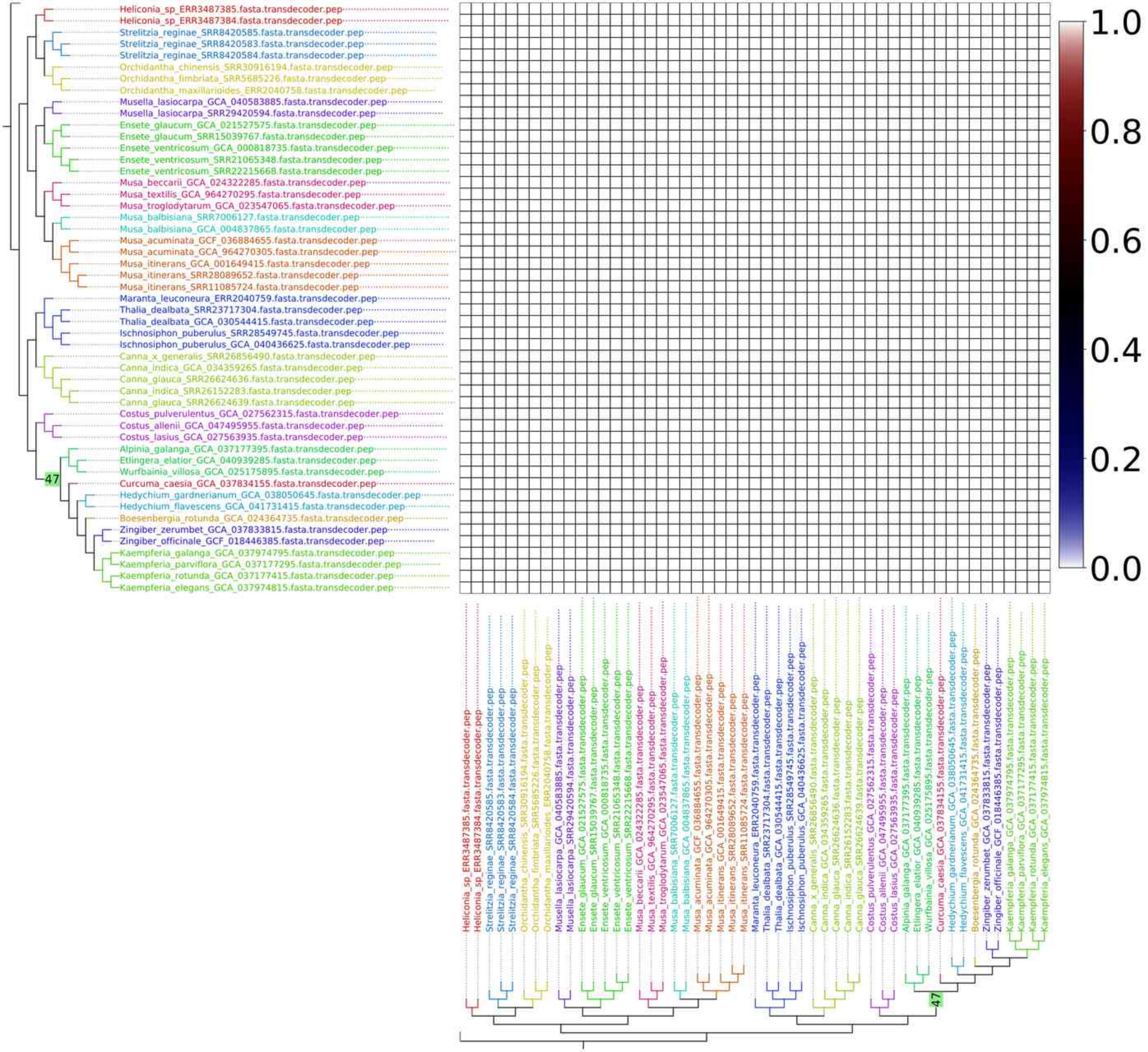
Heliconia_sp_ERR3487385.fasta.transdecoder.pep
Heliconia_sp_ERR3487384.fasta.transdecoder.pep
Strelitzia_reginae_SRR8420585.fasta.transdecoder.pep
Strelitzia_reginae_SRR8420583.fasta.transdecoder.pep
Strelitzia_reginae_SRR8420584.fasta.transdecoder.pep
Orchidantha_chinensis_SRR30916194.fasta.transdecoder.pep
Orchidantha_fimbriata_SRR5685226.fasta.transdecoder.pep
Orchidantha_maxillarioides_ERR2040758.fasta.transdecoder.pep
Musella_lasiocarpa_GCA_040583885.fasta.transdecoder.pep
Musella_lasiocarpa_SRR29420594.fasta.transdecoder.pep
Ensete_glaucum_GCA_021527575.fasta.transdecoder.pep
Ensete_glaucum_SRR15039767.fasta.transdecoder.pep
Ensete_ventricosum_GCA_000818735.fasta.transdecoder.pep
Ensete_ventricosum_SRR21065348.fasta.transdecoder.pep
Ensete_ventricosum_SRR22215668.fasta.transdecoder.pep
Musa_beccarii_GCA_024322285.fasta.transdecoder.pep
Musa_textilis_GCA_964270295.fasta.transdecoder.pep
Musa_troglodytarum_GCA_023547065.fasta.transdecoder.pep
Musa_balbisiana_SRR7006127.fasta.transdecoder.pep
Musa_balbisiana_GCA_004837865.fasta.transdecoder.pep
Musa_acuminata_GCF_036884655.fasta.transdecoder.pep
Musa_acuminata_GCA_964270305.fasta.transdecoder.pep
Musa_itinerans_GCA_001649415.fasta.transdecoder.pep
Musa_itinerans_SRR28089652.fasta.transdecoder.pep
Musa_itinerans_SRR11085724.fasta.transdecoder.pep
Maranta_leuconeura_ERR2040759.fasta.transdecoder.pep
Thalia_dealbata_SRR23717304.fasta.transdecoder.pep
Thalia_dealbata_GCA_030544415.fasta.transdecoder.pep
Ischnosiphon_puberulus_SRR28549745.fasta.transdecoder.pep
Ischnosiphon_puberulus_GCA_040436625.fasta.transdecoder.pep
Canna_x_generalis_SRR26856490.fasta.transdecoder.pep
Canna_indica_GCA_034359265.fasta.transdecoder.pep
Canna_glauca_SRR26624636.fasta.transdecoder.pep
Canna_indica_SRR26152283.fasta.transdecoder.pep
Canna_glauca_SRR26624639.fasta.transdecoder.pep
Costus_pulverulentus_GCA_027562315.fasta.transdecoder.pep
Costus_allenii_GCA_047495955.fasta.transdecoder.pep
Costus_lasius_GCA_027563935.fasta.transdecoder.pep
Alpinia_galanga_GCA_037177395.fasta.transdecoder.pep
Etlingera_elatior_GCA_040939285.fasta.transdecoder.pep
Wurfbainia_villosa_GCA_025175895.fasta.transdecoder.pep
Curcuma_caesia_GCA_037834155.fasta.transdecoder.pep
Hedychium_gardnerianum_GCA_038050645.fasta.transdecoder.pep
Hedychium_flavescens_GCA_041731415.fasta.transdecoder.pep
Boesenbergia_rotunda_GCA_024364735.fasta.transdecoder.pep
Zingiber_zerumbet_GCA_037833815.fasta.transdecoder.pep
Zingiber_officinale_GCF_018446385.fasta.transdecoder.pep
Kaempferia_galanga_GCA_037974795.fasta.transdecoder.pep
Kaempferia_parviflora_GCA_037177295.fasta.transdecoder.pep
Kaempferia_rotunda_GCA_037177415.fasta.transdecoder.pep
Kaempferia_elegans_GCA_037974815.fasta.transdecoder.pep
47
1.0
0.8
0.6
0.4
0.2
0.0

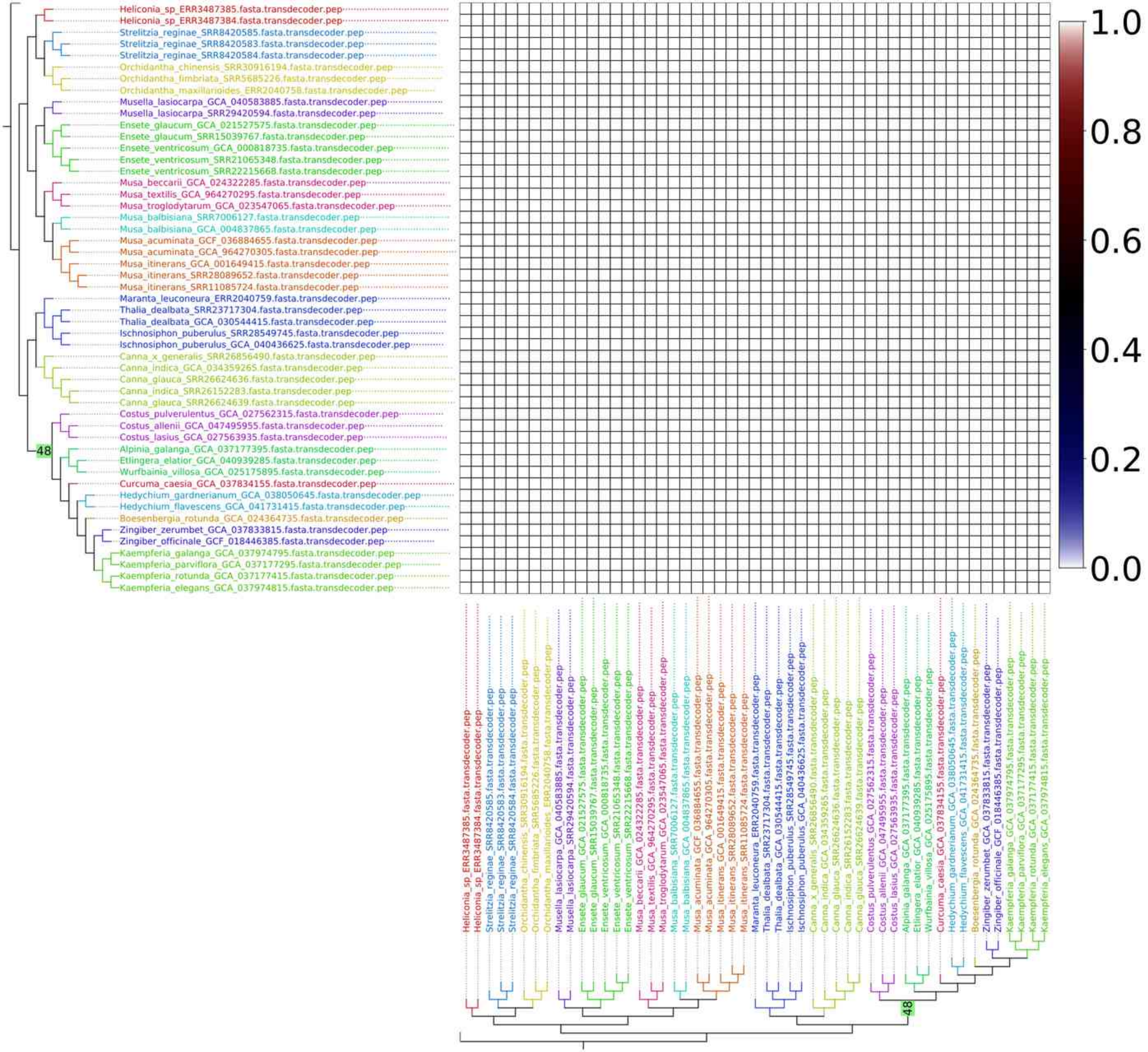
Heliconia_sp_ERR3487385.fasta.transdecoder.pep
Heliconia_sp_ERR3487384.fasta.transdecoder.pep
Strelitzia_reginae_SRR8420585.fasta.transdecoder.pep
Strelitzia_reginae_SRR8420583.fasta.transdecoder.pep
Strelitzia_reginae_SRR8420584.fasta.transdecoder.pep
Orchidantha_chinensis_SRR30916194.fasta.transdecoder.pep
Orchidantha_fimbriata_SRR5685226.fasta.transdecoder.pep
Orchidantha_maxillarioides_ERR2040758.fasta.transdecoder.pep
Musella_lasiocarpa_GCA_040583885.fasta.transdecoder.pep
Musella_lasiocarpa_SRR29420594.fasta.transdecoder.pep
Ensete_glaucum_GCA_021527575.fasta.transdecoder.pep
Ensete_glaucum_SRR15039767.fasta.transdecoder.pep
Ensete_ventricosum_GCA_000818735.fasta.transdecoder.pep
Ensete_ventricosum_SRR21065348.fasta.transdecoder.pep
Ensete_ventricosum_SRR22215668.fasta.transdecoder.pep
Musa_beccarii_GCA_024322285.fasta.transdecoder.pep
Musa_textilis_GCA_964270295.fasta.transdecoder.pep
Musa_troglodytarum_GCA_023547065.fasta.transdecoder.pep
Musa_balbisiana_SRR7006127.fasta.transdecoder.pep
Musa_balbisiana_GCA_004837865.fasta.transdecoder.pep
Musa_acuminata_GCF_036884655.fasta.transdecoder.pep
Musa_acuminata_GCA_964270305.fasta.transdecoder.pep
Musa_itinerans_GCA_001649415.fasta.transdecoder.pep
Musa_itinerans_SRR28089652.fasta.transdecoder.pep
Musa_itinerans_SRR11085724.fasta.transdecoder.pep
Maranta_leuconeura_ERR2040759.fasta.transdecoder.pep
Thalia_dealbata_SRR23717304.fasta.transdecoder.pep
Thalia_dealbata_GCA_030544415.fasta.transdecoder.pep
Ischnosiphon_puberulus_SRR28549745.fasta.transdecoder.pep
Ischnosiphon_puberulus_GCA_040436625.fasta.transdecoder.pep
Canna_x_generalis_SRR26856490.fasta.transdecoder.pep
Canna_indica_GCA_034359265.fasta.transdecoder.pep
Canna_glauca_SRR26624636.fasta.transdecoder.pep
Canna_indica_SRR26152283.fasta.transdecoder.pep
Canna_glauca_SRR26624639.fasta.transdecoder.pep
Costus_pulverulentus_GCA_027562315.fasta.transdecoder.pep
Costus_allenii_GCA_047495955.fasta.transdecoder.pep
Costus_lasius_GCA_027563935.fasta.transdecoder.pep
Alpinia_galanga_GCA_037177395.fasta.transdecoder.pep
Etlingera_elatior_GCA_040939285.fasta.transdecoder.pep
Wurfbainia_villosa_GCA_025175895.fasta.transdecoder.pep
Curcuma_caesia_GCA_037834155.fasta.transdecoder.pep
Hedychium_gardnerianum_GCA_038050645.fasta.transdecoder.pep
Hedychium_flavescens_GCA_041731415.fasta.transdecoder.pep
Boesenbergia_rotunda_GCA_024364735.fasta.transdecoder.pep
Zingiber_zerumbet_GCA_037833815.fasta.transdecoder.pep
Zingiber_officinale_GCF_018446385.fasta.transdecoder.pep
Kaempferia_galanga_GCA_037974795.fasta.transdecoder.pep
Kaempferia_parviflora_GCA_037177295.fasta.transdecoder.pep
Kaempferia_rotunda_GCA_037177415.fasta.transdecoder.pep
Kaempferia_elegans_GCA_037974815.fasta.transdecoder.pep
48
1.0
0.8
0.6
0.4
0.2
0.0

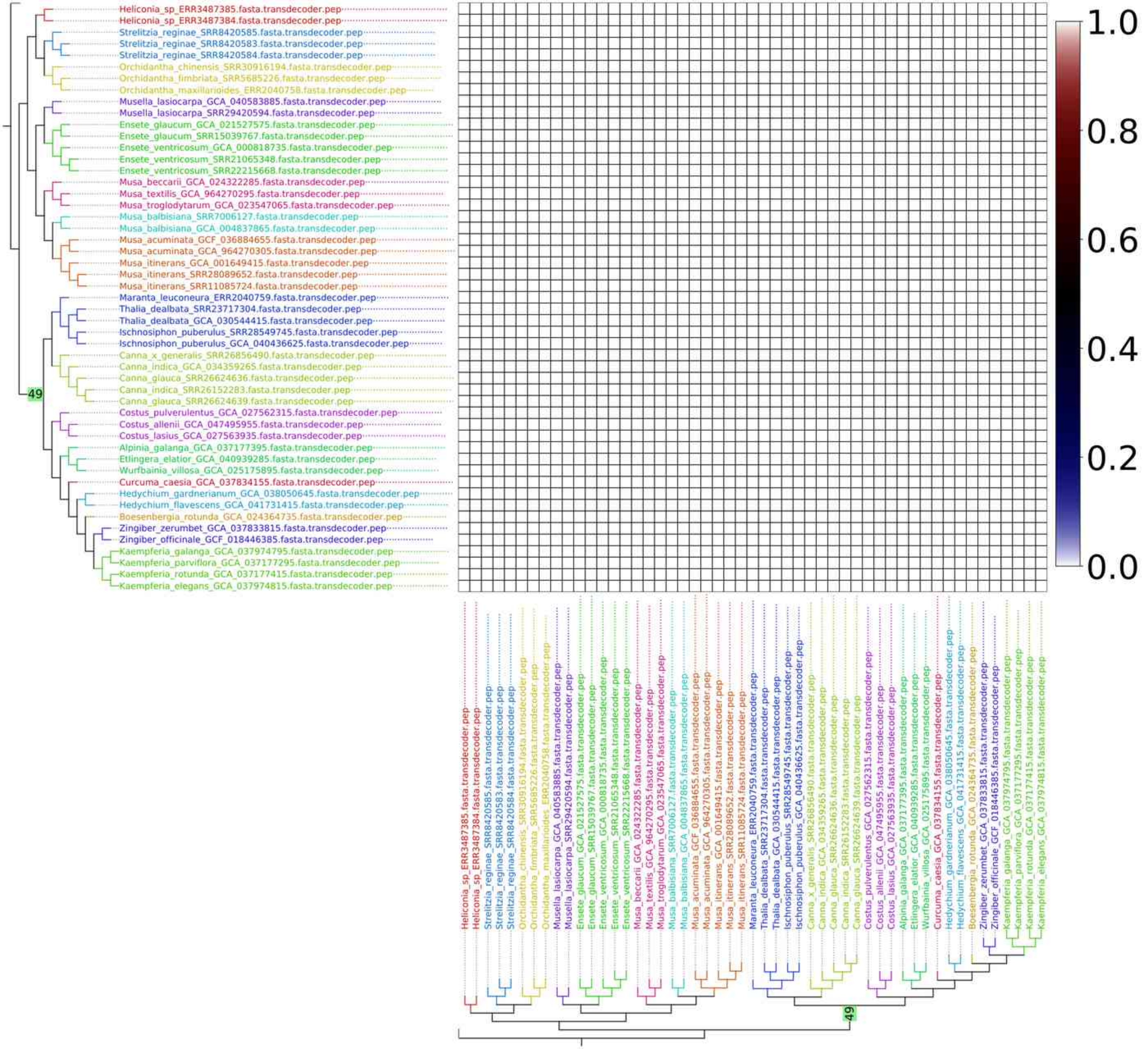
Heliconia_sp_ERR3487385.fasta.transdecoder.pep
Heliconia_sp_ERR3487384.fasta.transdecoder.pep
Strelitzia_reginae_SRR8420585.fasta.transdecoder.pep
Strelitzia_reginae_SRR8420583.fasta.transdecoder.pep
Strelitzia_reginae_SRR8420584.fasta.transdecoder.pep
Orchidantha_chinensis_SRR30916194.fasta.transdecoder.pep
Orchidantha_fimbriata_SRR5685226.fasta.transdecoder.pep
Orchidantha_maxillarioides_ERR2040758.fasta.transdecoder.pep
Musella_lasiocarpa_GCA_040583885.fasta.transdecoder.pep
Musella_lasiocarpa_SRR29420594.fasta.transdecoder.pep
Ensete_glaucum_GCA_021527575.fasta.transdecoder.pep
Ensete_glaucum_SRR15039767.fasta.transdecoder.pep
Ensete_ventricosum_GCA_000818735.fasta.transdecoder.pep
Ensete_ventricosum_SRR21065348.fasta.transdecoder.pep
Ensete_ventricosum_SRR22215668.fasta.transdecoder.pep
Musa_beccarii_GCA_024322285.fasta.transdecoder.pep
Musa_textilis_GCA_964270295.fasta.transdecoder.pep
Musa_troglodytarum_GCA_023547065.fasta.transdecoder.pep
Musa_balbisiana_SRR7006127.fasta.transdecoder.pep
Musa_balbisiana_GCA_004837865.fasta.transdecoder.pep
Musa_acuminata_GCF_036884655.fasta.transdecoder.pep
Musa_acuminata_GCA_964270305.fasta.transdecoder.pep
Musa_itinerans_GCA_001649415.fasta.transdecoder.pep
Musa_itinerans_SRR28089652.fasta.transdecoder.pep
Musa_itinerans_SRR11085724.fasta.transdecoder.pep
Maranta_leuconeura_ERR2040759.fasta.transdecoder.pep
Thalia_dealbata_SRR23717304.fasta.transdecoder.pep
Thalia_dealbata_GCA_030544415.fasta.transdecoder.pep
Ischnosiphon_puberulus_SRR28549745.fasta.transdecoder.pep
Ischnosiphon_puberulus_GCA_040436625.fasta.transdecoder.pep
Canna_x_generalis_SRR26856490.fasta.transdecoder.pep
Canna_indica_GCA_034359265.fasta.transdecoder.pep
Canna_glauca_SRR26624636.fasta.transdecoder.pep
Canna_indica_SRR26152283.fasta.transdecoder.pep
Canna_glauca_SRR26624639.fasta.transdecoder.pep
Costus_pulverulentus_GCA_027562315.fasta.transdecoder.pep
Costus_allenii_GCA_047495955.fasta.transdecoder.pep
Costus_lasius_GCA_027563935.fasta.transdecoder.pep
Alpinia_galanga_GCA_037177395.fasta.transdecoder.pep
Etlingera_elatior_GCA_040939285.fasta.transdecoder.pep
Wurfbainia_villosa_GCA_025175895.fasta.transdecoder.pep
Curcuma_caesia_GCA_037834155.fasta.transdecoder.pep
Hedychium_gardnerianum_GCA_038050645.fasta.transdecoder.pep
Hedychium_flavescens_GCA_041731415.fasta.transdecoder.pep
Boesenbergia_rotunda_GCA_024364735.fasta.transdecoder.pep
Zingiber_zerumbet_GCA_037833815.fasta.transdecoder.pep
Zingiber_officinale_GCF_018446385.fasta.transdecoder.pep
Kaempferia_galanga_GCA_037974795.fasta.transdecoder.pep
Kaempferia_parviflora_GCA_037177295.fasta.transdecoder.pep
Kaempferia_rotunda_GCA_037177415.fasta.transdecoder.pep
Kaempferia_elegans_GCA_037974815.fasta.transdecoder.pep
49
1.0
0.8
0.6
0.4
0.2
0.0

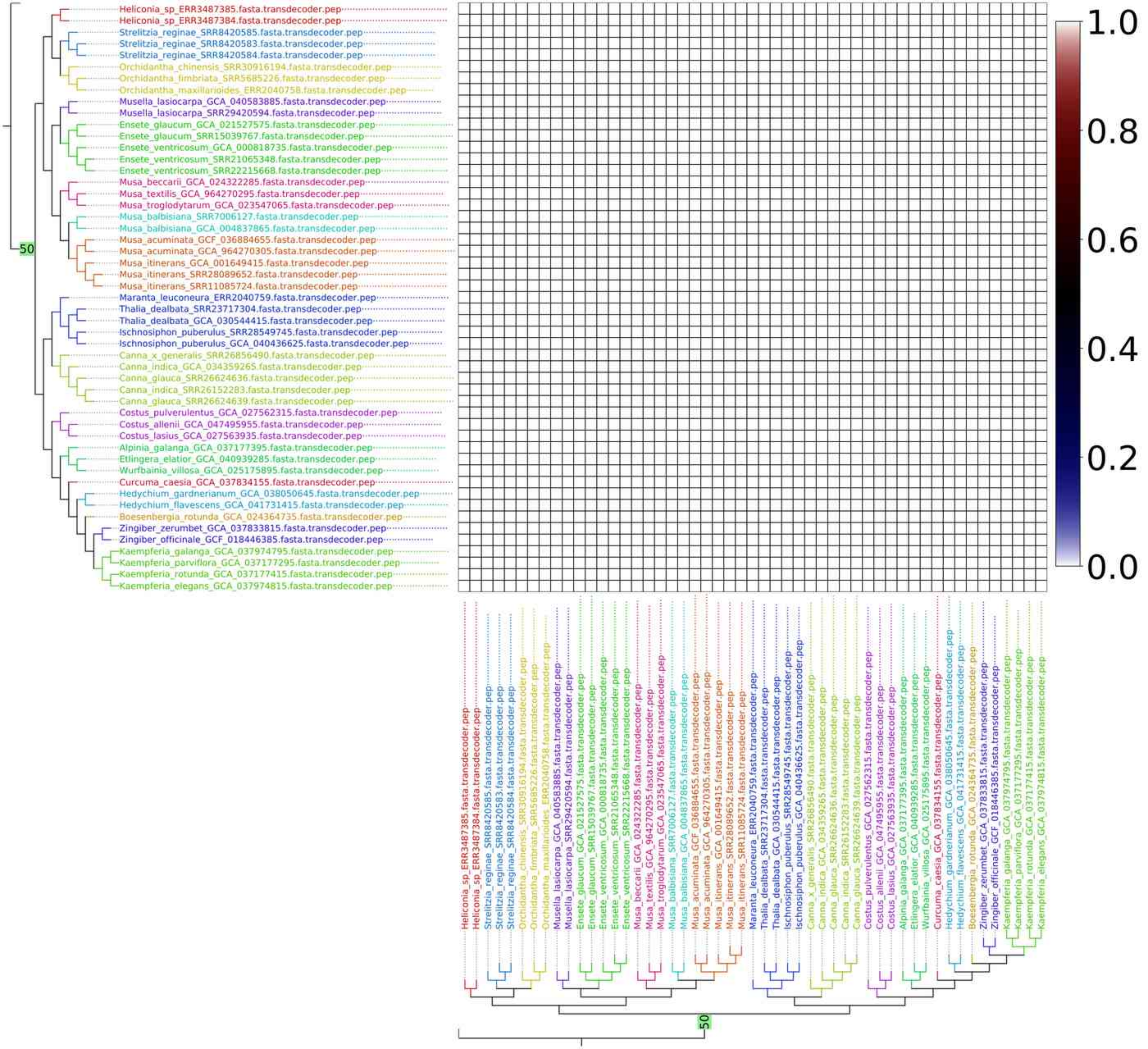

Heliconia_sp_ERR3487385.fasta.transdecoder.pep
Heliconia_sp_ERR3487384.fasta.transdecoder.pep
Strelitzia_reginae_SRR8420585.fasta.transdecoder.pep
Strelitzia_reginae_SRR8420583.fasta.transdecoder.pep
Strelitzia_reginae_SRR8420584.fasta.transdecoder.pep
Orchidantha_chinensis_SRR30916194.fasta.transdecoder.pep
Orchidantha_fimbriata_SRR5685226.fasta.transdecoder.pep
Orchidantha_maxillarioides_ERR2040758.fasta.transdecoder.pep
Musella_lasiocarpa_GCA_040583885.fasta.transdecoder.pep
Musella_lasiocarpa_SRR29420594.fasta.transdecoder.pep
Ensete_glaucum_GCA_021527575.fasta.transdecoder.pep
Ensete_glaucum_SRR15039767.fasta.transdecoder.pep
Ensete_ventricosum_GCA_000818735.fasta.transdecoder.pep
Ensete_ventricosum_SRR21065348.fasta.transdecoder.pep
Ensete_ventricosum_SRR22215668.fasta.transdecoder.pep
Musa_beccarii_GCA_024322285.fasta.transdecoder.pep
Musa_textilis_GCA_964270295.fasta.transdecoder.pep
Musa_troglodytarum_GCA_023547065.fasta.transdecoder.pep
Musa_balbisiana_SRR7006127.fasta.transdecoder.pep
Musa_balbisiana_GCA_004837865.fasta.transdecoder.pep
Musa_acuminata_GCF_036884655.fasta.transdecoder.pep
Musa_acuminata_GCA_964270305.fasta.transdecoder.pep
Musa_itinerans_GCA_001649415.fasta.transdecoder.pep
Musa_itinerans_SRR28089652.fasta.transdecoder.pep
Musa_itinerans_SRR11085724.fasta.transdecoder.pep
Maranta_leuconeura_ERR2040759.fasta.transdecoder.pep
Thalia_dealbata_SRR23717304.fasta.transdecoder.pep
Thalia_dealbata_GCA_030544415.fasta.transdecoder.pep
Ischnosiphon_puberulus_SRR28549745.fasta.transdecoder.pep
Ischnosiphon_puberulus_GCA_040436625.fasta.transdecoder.pep
Canna_x_generalis_SRR26856490.fasta.transdecoder.pep
Canna_indica_GCA_034359265.fasta.transdecoder.pep
Canna_glauca_SRR26624636.fasta.transdecoder.pep
Canna_indica_SRR26152283.fasta.transdecoder.pep
Canna_glauca_SRR26624639.fasta.transdecoder.pep
Costus_pulverulentus_GCA_027562315.fasta.transdecoder.pep
Costus_allenii_GCA_047495955.fasta.transdecoder.pep
Costus_lasius_GCA_027563935.fasta.transdecoder.pep
Alpinia_galanga_GCA_037177395.fasta.transdecoder.pep
Etlingera_elatior_GCA_040939285.fasta.transdecoder.pep
Wurfbainia_villosa_GCA_025175895.fasta.transdecoder.pep
Curcuma_caesia_GCA_037834155.fasta.transdecoder.pep
Hedychium_gardnerianum_GCA_038050645.fasta.transdecoder.pep
Hedychium_flavescens_GCA_041731415.fasta.transdecoder.pep
Boesenbergia_rotunda_GCA_024364735.fasta.transdecoder.pep
Zingiber_zerumbet_GCA_037833815.fasta.transdecoder.pep
Zingiber_officinale_GCF_018446385.fasta.transdecoder.pep
Kaempferia_galanga_GCA_037974795.fasta.transdecoder.pep
Kaempferia_parviflora_GCA_037177295.fasta.transdecoder.pep
Kaempferia_rotunda_GCA_037177415.fasta.transdecoder.pep
Kaempferia_elegans_GCA_037974815.fasta.transdecoder.pep
50
1.0
0.8
0.6
0.4
0.2
0.0
