## Supplementary material for "Nuclear phylogenomics clarifies the family-level backbone and gene-tree conflict in Zingiberales": Table S1

Table S1. Taxon sampling, data sources, data types, and NCBI accession numbers for Zingiberales and the outgroup sampled in this study.

| **Family** | **Species** | **Data type** | **Raw reads** | **NCBI accession** |
| --- | --- | --- | --- | --- |
| Musaceae | *Ensete ventricosum* | Whole-genome sequencing | - | GCA_000818735.3 |
| Musaceae | *Ensete glaucum* | Transcriptome sequencing | 31205420 | SRR15039767 |
| Musaceae | *Ensete glaucum* | Whole-genome sequencing | - | GCA_021527575.1 |
| Musaceae | *Ensete ventricosum* | Transcriptome sequencing | 25973466 | SRR21065348 |
| Musaceae | *Ensete ventricosum* | Transcriptome sequencing | 29551678 | SRR22215668 |
| Musaceae | *Musella lasiocarpa* | Whole-genome sequencing | - | GCA_040583885.1 |
| Musaceae | *Musella lasiocarpa* | Transcriptome sequencing | 33344831 | SRR29420594 |
| Musaceae | *Musa itinerans* | Whole-genome sequencing | - | GCA_001649415.1 |
| Musaceae | *Musa acuminata* | Whole-genome sequencing | - | GCA_964270305 |
| Musaceae | *Musa acuminata* | Whole-genome sequencing | - | GCF_036884655.1 |
| Musaceae | *Musa balbisiana* | Whole-genome sequencing | - | GCA_004837865.1 |
| Musaceae | *Musa balbisiana* | Transcriptome sequencing | 47714027 | SRR7006127 |
| Musaceae | *Musa itinerans* | Transcriptome sequencing | 33688798 | SRR11085724 |
| Musaceae | *Musa itinerans* | Transcriptome sequencing | 19318832 | SRR28089652 |
| Musaceae | *Musa beccarii* | Whole-genome sequencing | - | GCA_024322285.1 |
| Musaceae | *Musa textilis* | Whole-genome sequencing | - | GCA_964270295.1 |
| Musaceae | *Musa troglodytarum* | Whole-genome sequencing | - | GCA_023547065.1 |
| Strelitziaceae | *Strelitzia reginae* | Transcriptome sequencing | 21221372 | SRR8420583 |
| Strelitziaceae | *Strelitzia reginae* | Transcriptome sequencing | 20826586 | SRR8420585 |
| Strelitziaceae | *Strelitzia reginae* | Transcriptome sequencing | 23475068 | SRR8420584 |
| Lowiaceae | *Orchidantha chinensis* | Transcriptome sequencing | 24445020 | SRR30916194 |
| Lowiaceae | *Orchidantha fimbriata* | Transcriptome sequencing | 13111877 | SRR5685226 |
| Lowiaceae | *Orchidantha maxillarioides* | Transcriptome sequencing | 8690257 | ERR2040758 |
| Heliconiaceae | *Heliconia* sp. 1KP | Transcriptome sequencing | 15487342 | ERR3487385 |
| Heliconiaceae | *Heliconia* sp. 1KP | Transcriptome sequencing | 12796988 | ERR3487384 |
| Cannaceae | *Canna indica* | Whole-genome sequencing | - | GCA_034359265.1 |
| Cannaceae | *Canna indica* | Transcriptome sequencing | 25845082 | SRR26152283 |
| Cannaceae | *Canna* x *generalis* | Transcriptome sequencing | 26444021 | SRR26856490 |
| Cannaceae | *Canna glauca* | Transcriptome sequencing | 26680561 | SRR26624636 |
| Cannaceae | *Canna glauca* | Transcriptome sequencing | 27296035 | SRR26624639 |
| Marantaceae | *Thalia dealbata* | Whole-genome sequencing | - | GCA_030544415.1 |
| Marantaceae | *Thalia dealbata* | Transcriptome sequencing | 22028161 | SRR23717304 |
| Marantaceae | *Ischnosiphon puberulus* | Whole-genome sequencing | - | GCA_040436625.1 |
| Marantaceae | *Ischnosiphon puberulus* | Transcriptome sequencing | 34914725 | SRR28549745 |
| Marantaceae | *Maranta leuconeura* | Transcriptome sequencing | 7308231 | ERR2040759 |
| Costaceae | *Costus allenii* | Whole-genome sequencing | - | GCA_047495955.1 |
| Costaceae | *Costus lasius* | Whole-genome sequencing | - | GCA_027563935.2 |
| Costaceae | *Costus pulverulentus* | Whole-genome sequencing | - | GCA_027562315.1 |
| Zingiberaceae | *Alpinia galanga* | Whole-genome sequencing | - | GCA_037177395.1 |
| Zingiberaceae | *Etlingera elatior* | Whole-genome sequencing | - | GCA_040939285.1 |
| Zingiberaceae | *Boesenbergia rotunda* | Whole-genome sequencing | - | GCA_024364735.1 |
| Zingiberaceae | *Curcuma caesia* | Whole-genome sequencing | - | GCA_037834155.1 |
| Zingiberaceae | *Hedychium flavescens* | Whole-genome sequencing | - | GCA_041731415.1 |
| Zingiberaceae | *Hedychium gardnerianum* | Whole-genome sequencing | - | GCA_038050645.1 |
| Zingiberaceae | *Kaempferia elegans* | Whole-genome sequencing | - | GCA_037974815.1 |
| Zingiberaceae | *Kaempferia galanga* | Whole-genome sequencing | - | GCA_037974795.1 |
| Zingiberaceae | *Kaempferia parviflora* | Whole-genome sequencing | - | GCA_037177295.1 |
| Zingiberaceae | *Kaempferia rotunda* | Whole-genome sequencing | - | GCA_037177415.1 |
| Zingiberaceae | *Wurfbainia villosa* | Whole-genome sequencing | - | GCA_025175895.1 |
| Zingiberaceae | *Zingiber officinale* | Whole-genome sequencing | - | GCF_018446385.1 |
| Zingiberaceae | *Zingiber zerumbet* | Whole-genome sequencing | - | GCA_037833815.1 |
| Pontederiaceae | *Pontederia crassipes* | Whole-genome sequencing | - | GCA_030549335.1 |
