## Supplementary material for "Nuclear phylogenomics clarifies the family-level backbone and gene-tree conflict in Zingiberales": Table S2

Table S2. CDS recovery statistics for each sample after redundancy reduction.

| **Family** | **Species** | **NCBI accession** | **Number of CDSs** | **CDS N50 (bp)** |
| --- | --- | --- | --- | --- |
| Musaceae | *Ensete ventricosum* | GCA_000818735.3 | 39970 | 1002 |
| Musaceae | *Ensete glaucum* | SRR15039767 | 34336 | 1098 |
| Musaceae | *Ensete glaucum* | GCA_021527575.1 | 12622 | 1299 |
| Musaceae | *Ensete ventricosum* | SRR21065348 | 29138 | 1176 |
| Musaceae | *Ensete ventricosum* | SRR22215668 | 31559 | 753 |
| Musaceae | *Musella lasiocarpa* | GCA_040583885.1 | 12762 | 1311 |
| Musaceae | *Musella lasiocarpa* | SRR29420594 | 31236 | 1056 |
| Musaceae | *Musa itinerans* | GCA_001649415.1 | 13521 | 1329 |
| Musaceae | *Musa acuminata* | GCA_964270305 | 29086 | 1644 |
| Musaceae | *Musa acuminata* | GCF_036884655.1 | 36248 | 1671 |
| Musaceae | *Musa balbisiana* | GCA_004837865.1 | 29045 | 1587 |
| Musaceae | *Musa balbisiana* | SRR7006127 | 25801 | 1194 |
| Musaceae | *Musa itinerans* | SRR11085724 | 36891 | 1092 |
| Musaceae | *Musa itinerans* | SRR28089652 | 32967 | 1029 |
| Musaceae | *Musa beccarii* | GCA_024322285.1 | 13387 | 1347 |
| Musaceae | *Musa textilis* | GCA_964270295.1 | 28268 | 1596 |
| Musaceae | *Musa troglodytarum* | GCA_023547065.1 | 44760 | 1740 |
| Strelitziaceae | *Strelitzia reginae* | SRR8420583 | 29700 | 1113 |
| Strelitziaceae | *Strelitzia reginae* | SRR8420585 | 29193 | 1116 |
| Strelitziaceae | *Strelitzia reginae* | SRR8420584 | 30500 | 1107 |
| Lowiaceae | *Orchidantha chinensis* | SRR30916194 | 31667 | 996 |
| Lowiaceae | *Orchidantha fimbriata* | SRR5685226 | 15749 | 792 |
| Lowiaceae | *Orchidantha maxillarioides* | ERR2040758 | 25376 | 819 |
| Heliconiaceae | *Heliconia* sp. 1KP | ERR3487385 | 24370 | 1167 |
| Heliconiaceae | *Heliconia* sp. 1KP | ERR3487384 | 18255 | 714 |
| Cannaceae | *Canna indica* | GCA_034359265.1 | 25775 | 1569 |
| Cannaceae | *Canna indica* | SRR26152283 | 28290 | 1161 |
| Cannaceae | *Canna* x *generalis* | SRR26856490 | 29384 | 1161 |
| Cannaceae | *Canna glauca* | SRR26624636 | 29054 | 1191 |
| Cannaceae | *Canna glauca* | SRR26624639 | 29208 | 1197 |
| Marantaceae | *Thalia dealbata* | GCA_030544415.1 | 9906 | 1359 |
| Marantaceae | *Thalia dealbata* | SRR23717304 | 24743 | 1203 |
| Marantaceae | *Ischnosiphon puberulus* | GCA_040436625.1 | 10057 | 1398 |
| Marantaceae | *Ischnosiphon puberulus* | SRR28549745 | 24004 | 774 |
| Marantaceae | *Maranta leuconeura* | ERR2040759 | 23868 | 846 |
| Costaceae | *Costus allenii* | GCA_047495955.1 | 11660 | 1377 |
| Costaceae | *Costus lasius* | GCA_027563935.2 | 11261 | 1401 |
| Costaceae | *Costus pulverulentus* | GCA_027562315.1 | 11611 | 1341 |
| Zingiberaceae | *Alpinia galanga* | GCA_037177395.1 | 11166 | 1293 |
| Zingiberaceae | *Etlingera elatior* | GCA_040939285.1 | 9680 | 1314 |
| Zingiberaceae | *Boesenbergia rotunda* | GCA_024364735.1 | 12528 | 1386 |
| Zingiberaceae | *Curcuma caesia* | GCA_037834155.1 | 11320 | 1311 |
| Zingiberaceae | *Hedychium flavescens* | GCA_041731415.1 | 11017 | 1263 |
| Zingiberaceae | *Hedychium gardnerianum* | GCA_038050645.1 | 12141 | 1293 |
| Zingiberaceae | *Kaempferia elegans* | GCA_037974815.1 | 10623 | 1263 |
| Zingiberaceae | *Kaempferia galanga* | GCA_037974795.1 | 10436 | 1272 |
| Zingiberaceae | *Kaempferia parviflora* | GCA_037177295.1 | 11136 | 1296 |
| Zingiberaceae | *Kaempferia rotunda* | GCA_037177415.1 | 9863 | 1260 |
| Zingiberaceae | *Wurfbainia villosa* | GCA_025175895.1 | 11784 | 1389 |
| Zingiberaceae | *Zingiber officinale* | GCF_018446385.1 | 37753 | 1632 |
| Zingiberaceae | *Zingiber zerumbet* | GCA_037833815.1 | 11102 | 1308 |
| Pontederiaceae | *Pontederia crassipes* | GCA_030549335.1 | 18595 | 1377 |
